# Diversity of satellite glia in sympathetic and sensory ganglia

**DOI:** 10.1101/2021.05.25.445647

**Authors:** Aurelia A Mapps, Michael B Thomsen, Erica Boehm, Haiqing Zhao, Samer Hattar, Rejji Kuruvilla

## Abstract

Satellite glia are the major glial type found in ganglia of the peripheral nervous system and wrap around cell bodies of sympathetic and sensory neurons that are very diverse. Other than their close physical association with peripheral neurons, little is known about this glial population. Here, we performed single cell RNA sequencing analysis and identified five different populations of satellite glia from sympathetic and sensory ganglia. We identified three shared populations of satellite glia enriched in immune-response genes, immediate-early genes and ion channels/ECM-interactors, respectively. Sensory- and sympathetic-specific satellite glia are differentially enriched for modulators of lipid synthesis and metabolism. Sensory glia are also specifically enriched for genes involved in glutamate turnover. Further, satellite glia and Schwann cells can be distinguished by unique transcriptional signatures. This study reveals remarkable heterogeneity of satellite glia in the peripheral nervous system.

## INTRODUCTION

In the peripheral nervous system, the sensory arm is essential for transducing a wide variety of environmental stimuli from the outside world to the central nervous system, while the sympathetic division relays motor commands from the central nervous system to diverse peripheral organs/tissues to mobilize the “fight or flight” response and maintain body homeostasis in response to a continuously changing environment (Goldstein, 2013; Marmigere and Ernfors, 2007; Usoskin et al., 2015). Sympathetic and sensory neurons residing in their respective peripheral ganglia are remarkably diverse with respect to morphological, molecular, and electrophysiological properties, consistent with their distinct functions (Ernsberger et al., 2020; Liu and Ma, 2011). In addition to primary neurons, sympathetic and sensory ganglia contain two major glial cell types, satellite glia and Schwann cells, which are closely associated with their neuronal neighbors and influence a wide range of neuronal functions (Hanani and Spray, 2020; Jessen and Mirsky, 2005). While the diversity of neurons in sympathetic versus sensory ganglia is well-appreciated, it remains unknown if there are molecular differences between the associated glial cell types, particularly the satellite glia, that reflect ganglion-specific functions.

Satellite glial and Schwann cells are both derived from multipotent neural crest precursors during development (Jessen and Mirsky, 2005). Satellite glial cells wrap tightly around neuronal cell bodies, whereas Schwann cells ensheath axonal processes of peripheral neurons (Hanani and Spray, 2020; Pannese, 1981). While Schwann cells have been intensely investigated, and have well-documented roles in myelination, axon regeneration, and providing trophic and metabolic support to neurons (Jessen and Mirsky, 2005; Monk et al., 2015), the understanding of satellite glia has lagged far behind. Satellite glia have a unique architecture in that they form ring-like structures that completely wrap around individual neuronal cell bodies, with each neuron and associated satellite glia thought to form discrete structural and functional units (Hanani and Spray, 2020; Pannese, 1981). This intimate physical association implies that satellite glia can be critical regulators of neuronal connectivity, activity, homeostasis, and repair. Indeed, it has been proposed that satellite glia modulate ionic and neurotransmitter concentrations, promote neuronal morphogenesis during development, regulate synaptic transmission, and engulf dying neurons (Avraham et al., 2020; Enes et al., 2020; Hanani and Spray, 2020; Wu et al., 2009). However, knowledge about satellite glia functions, specifically *in vivo*, has been scarce. Several recent studies in sensory ganglia have suggested that satellite glia regulate chronic pain through modulating neuronal hyper-excitability and promote axon regeneration after peripheral nerve injury (Avraham et al., 2020; Kim et al., 2016).

Satellite glia wrap around sensory and sympathetic neurons, which are molecularly, functionally, and morphologically divergent populations. Sympathetic neurons, which are primarily noradrenergic, innervate peripheral organs and tissues to control various autonomic functions, including cardiac output, body temperature, regulation of blood glucose levels, and immune functions under basal conditions and in response to external stressors (Goldstein, 2013). Sensory neurons relay information about mechano-sensation, pain and temperature from the periphery to the brain (Liu and Ma, 2011; Usoskin et al., 2015). Morphologically, sympathetic neurons have an axon and multiple dendritic processes and form synapses with preganglionic neurons, whereas sensory neuron cell bodies extend a pseudo-unipolar axon which bifurcates to extend into the brain and periphery, respectively. In a key difference from sensory ganglia, satellite glia envelop the dendrites and synapses of neurons, in addition to cell bodies in sympathetic ganglia (Hanani, 2010). Despite the intimate association of satellite glial cells with peripheral neurons, there have been few investigations of whether differences exist in satellite glial cells from distinct peripheral ganglia.

Here, we utilized single cell transcriptome profiling and single molecule fluorescence *in situ* hybridization to comprehensively characterize satellite glial cell diversity in sensory and sympathetic ganglia. Our results reveal five types of satellite glia, including sensory-specific satellite glia, sympathetic-specific satellite glia, and three populations that are present in both ganglia. Sensory- and sympathetic-specific satellite glia are differentially enriched for modulators of lipid metabolism. Sensory glia are specifically enriched for genes involved in glutamate turnover, consistent with their association to glutamatergic neurons. We identify three populations of satellite glia that are present in both ganglia, each featuring enrichment of functionally distinct pathways, including immune-response genes, immediate-early genes, and ion channels/ECM-interactors, respectively. We also show that Schwann cells can be distinguished from satellite glia by hundreds of uniquely expressed genes. By providing a transcriptional atlas for satellite glia classification, this study reveals remarkable diversity in satellite glia across the peripheral nervous system and suggests that satellite glia are transcriptionally tuned to the functions of their respective ganglia.

## RESULTS

### Cell types in sympathetic and sensory ganglia

We prepared single cell suspensions from sympathetic (superior cervical ganglion, SCG) and sensory (dorsal root ganglion, DRG) ganglia from young adult mice and performed droplet-based high-throughput single cell RNA sequencing using the Drop-Seq protocol (Macosko et al., 2015) (Figures S1A-B). The Drop-Seq procedure was validated by 98.75% separation efficacy of mixed human and mouse cells (Figure S1B). Following quality control filtering, we obtained a total of 21,524 high quality single cell transcriptomes (13,435 SCG; 8,089 DRG) that were used as input for unsupervised clustering and dimensionality reduction with the Seurat package (Figure 1A). This analysis identified numerous transcriptionally distinct cell populations divided between sensory and sympathetic tissues (Figure 1B). Major cell types of the peripheral nervous system were separated into nine large clusters based on the expression of canonical marker genes: satellite glial cells (*Plp1*, *Fabp7*), Schwann cells (*Plp1*, *Ncmap*), sympathetic neurons (*Snap25*, *Th*), sensory neurons (*Snap25*, *Calca*), vascular endothelial cells (*Ly6c1*), macrophages (*C1qb*), T-cells (*Trb2*), fibroblasts (*Dcn*), and mural cells (*Rgs5*) (Figure 1C) (Saunders et al., 2018; Vanlandewijck et al., 2018; Zeisel et al., 2018). Thus, our methodology is robust enough to isolate all known cell types from sympathetic and sensory ganglia.

**Figure 1.**
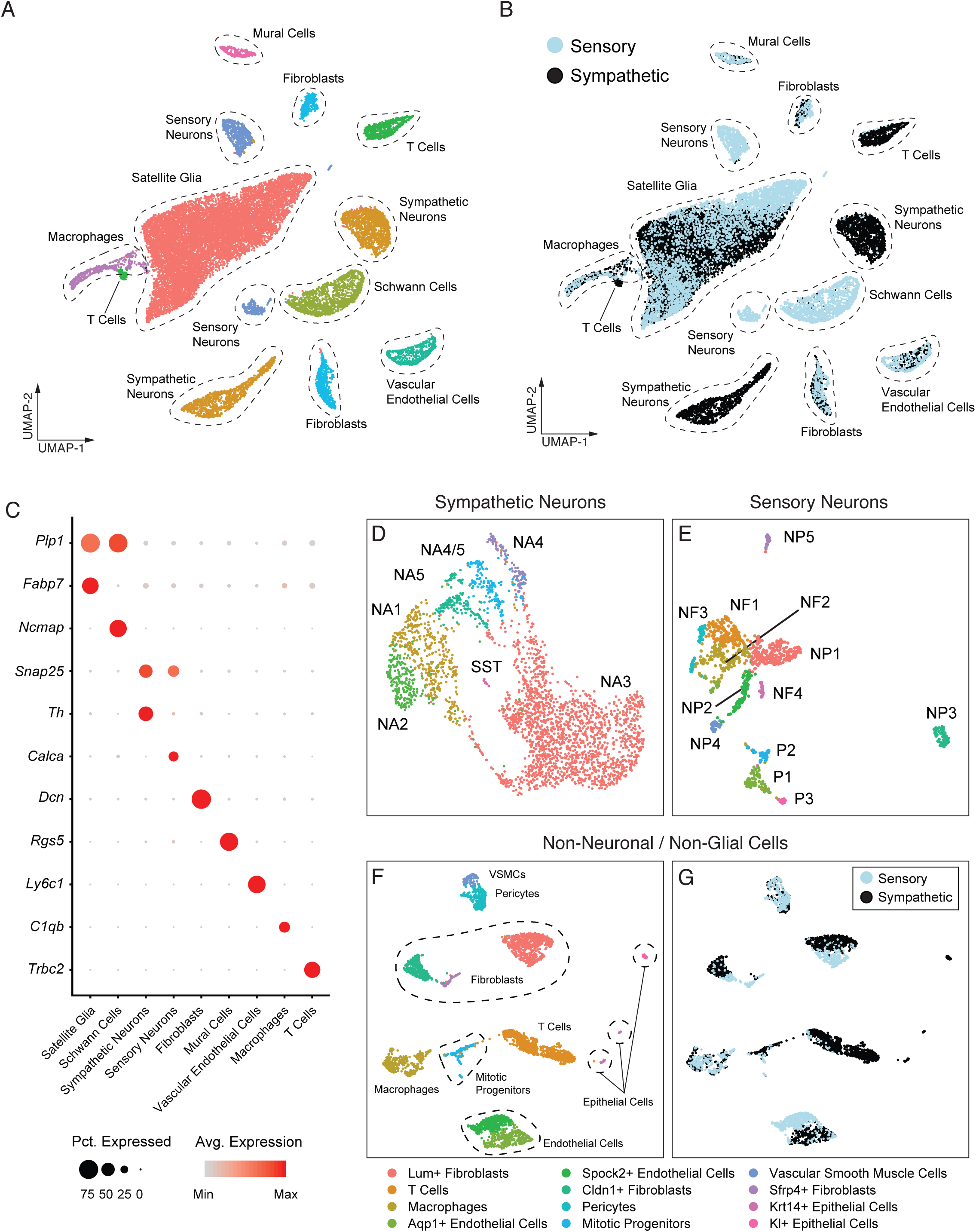
Cell type diversity in sensory and sympathetic ganglia. (A) UMAP of 21,524 single cells with major cell types highlighted (B) UMAP as in (A) overlaid with ganglia of origin (C) Dot plot showing differentially enriched genes for diverse cell types in peripheral ganglia. Dot size is proportional to the percentage of each cluster expressing the marker gene, and the color intensity is correlated with expression level. (D and E) UMAP plots of sympathetic and sensory neurons following isolation and re-clustering. Clusters are highlighted with literature-defined names based on published gene expression patterns (Furlan et al., 2016, Zeisel et al., 2018) (F) Non-neuronal and non-glial cell type clusters (G) Same as in (F) overlaid with ganglia of origin

We extracted sensory and sympathetic neurons and performed unsupervised clustering to identify cell type diversity contained within each class (Figures 1D-E). Within stellate and thoracic sympathetic ganglia, seven neuron types have been described to date, including five noradrenergic and two cholinergic populations (Furlan et al., 2016; Zeisel et al., 2018). Among SCG neurons, we identified all previously described five noradrenergic populations delineated by the expression of markers including *Th, Rarres1*, *Gfra2*, *Gfra3*, *Npy*, and *Enc1* (Figures 1D and S2A) (Furlan et al., 2016). However, we were unable to identify any cholinergic neurons, suggesting that cholinergic neurons are scarce in the SCG. Up to seventeen neuron types have been identified in sensory ganglia based on transcriptional profiles, and they are divided into three categories: peptidergic (8), non-peptidergic (6), and neurofilament-enriched (3) (Usoskin et al., 2015; Zeisel et al., 2018). We identified 12 types of sensory neurons corresponding to all three major categories including peptidergic (*Foxp2*, *Adcyap1*, *Mrgpra3*), non-peptidergic (*Th*, *Mrgprd*, *Nppb*), and neurofilament-enriched (*Nefh*, *Palm3*, *Trapp3cl*) (Figures 1E and S2B). In summary, our analyses of neuronal cell types from sensory and sympathetic ganglia are consistent with previous reports of neuronal diversity in the peripheral nervous system (Furlan et al., 2016; Usoskin et al., 2015; Zeisel et al., 2018).

### Non-neuronal/non glial cell types in peripheral ganglia

Non-neuronal cells were identified based on known gene expression signatures including vascular endothelial cells (*Ly6c1*, *Flt1*, *Cldn5*), mural cells (*Rgs5*, *Vtn*), fibroblasts (*Dcn, Lum*), T cells (*Trbc2*), macrophages (*C1qb*, *Ctss*), and epithelial cells (*Kl*, *Krt18*) (Zeisel et al., 2018) (Figures 1C, 1F **and** S1C). Epithelial cells and T-cells were found primarily in sympathetic samples, which may reflect a dissection artifact since these cell types reside outside of peripheral ganglia (Figure 1G). Pericytes, vascular smooth muscle cells, and macrophages showed limited or no differential gene expression between tissue of origin. We observed differences in gene expression between endothelial cells derived from sensory and sympathetic ganglia (Figure 1G), which may underlie observed differences in their vascular permeability (Kiernan, 1996).

Further, we identified a small cluster of cells that express several mitotic machinery components, including *Top2a* and *Mki67* (Figures 1F and S1C-D). Within this small cluster, we identified subsets of cells that contain gene expression signatures corresponding to T-cells, endothelial cells, erythrocytes, and satellite glial cells (Figure S1D). These results suggest that a subpopulation of satellite glia retain proliferative capacity in adult ganglia.

### Satellite glia and Schwann cells have distinct transcriptional signatures

Progress in understanding satellite glia functions has been hampered by the fact that they share many genetic markers with Schwann cells (Britsch et al., 2001; Hanani, 2010; Shi et al., 2008). While there are several genes that can be used to label satellite glial cells (e.g., *Glul, Gfap, S100b, Gja1, Plp1, Kcnj10*), most of these genes are also expressed in Schwann cells and/or other glial cell types in the central nervous system, most notably astrocytes (Hanani and Spray, 2020). To overcome this hurdle and isolate specific transcripts and functional pathways that distinguish these glial cell types, we independently clustered all glial cells (Figures 2A-B) and identified >450 genes that were differentially expressed (p<10^-50^) between Schwann cells and satellite glia (Figure 2C, **Table S1**). We found that the strongest and most significantly enriched transcript among satellite glial cells was fatty acid binding protein 7 (*Fabp7*) (Figure 2C), a fatty acid transporter that was previously identified as a marker of satellite glia (Avraham et al., 2020; Kurtz et al., 1994). In addition to *Fabp7*, satellite glia expressed high levels of genes associated with fatty acid synthesis including *ApoE*, an apolipoprotein which carries lipids and cholesterols to neurons, as well as *Dbi*, acyl-coA binding protein that regulates synapses, (**Table S1**), (Bouyakdan et al., 2019; Holtzman et al., 2012; Ioannou et al., 2019). Schwann cells were enriched for *Ncmap* (non-compact myelin associated protein) (Figure 2C), a glycoprotein that is found in myelin of the peripheral nervous system which is localized to paranodal regions flanking nodes of Ranvier (Ryu et al., 2008). *Ncmap* and other genes involved in myelination (*Mag, Mog*) were uniformly expressed within the Schwann cell cluster (**Table S1**), suggesting that we primarily isolated myelinating Schwann cells.

**Figure 2.**
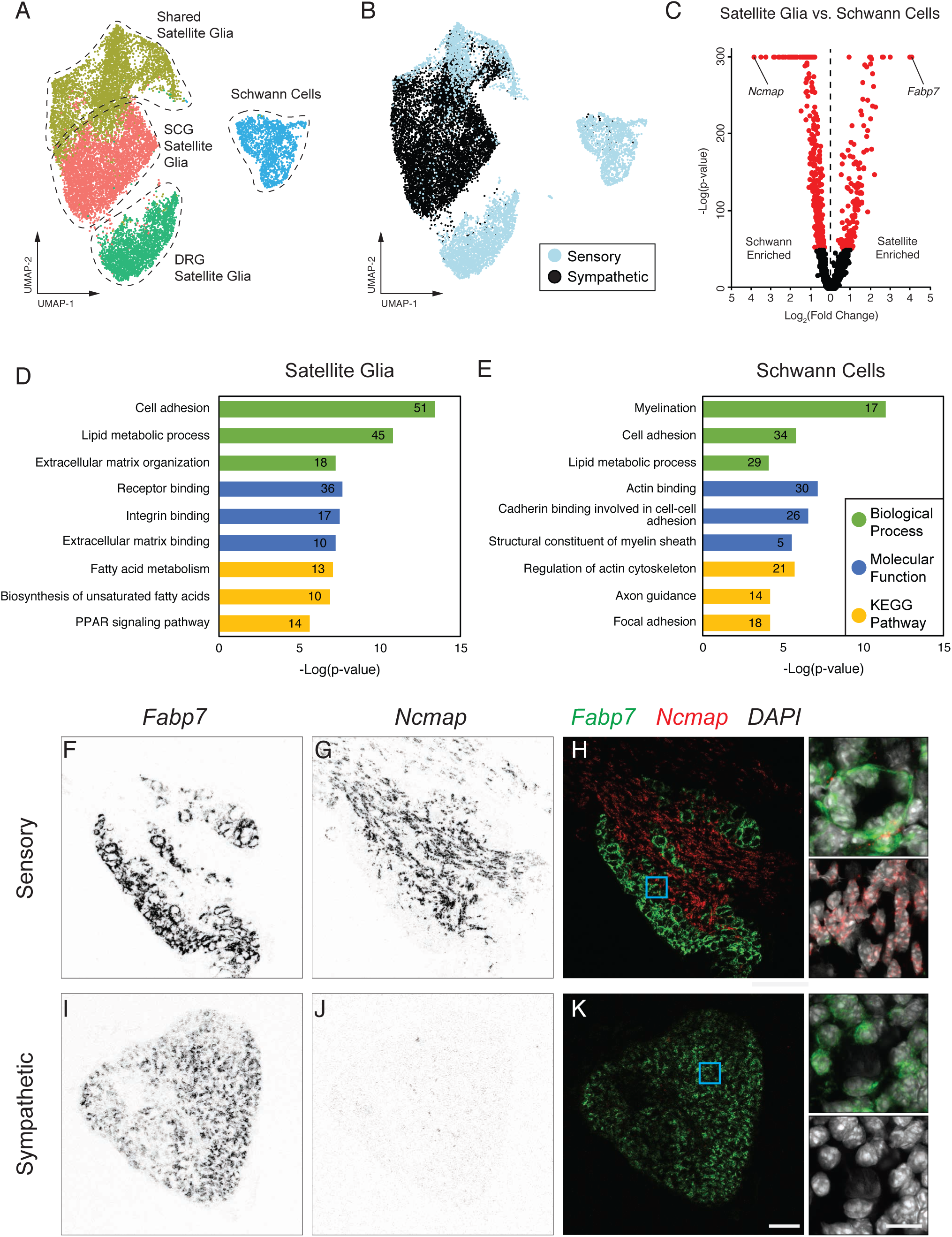
Comparison of Schwann cell and satellite glial cell transcriptional identity. (A) UMAP of glial cell populations in sensory and sympathetic tissue. (B) Same as in (A) overlaid with ganglia of origin (C) Volcano plot of all differentially expressed genes between Schwann cells and satellite glia (red dots: p<10^-50^) (D-E) Gene ontology and KEGG pathway analysis of satellite glia- and Schwann cell-enriched transcripts. Top 3 biological process terms, molecular function terms, and KEGG pathways are shown. Labels denote the number of genes identified in each category. (F-H) smFISH of *Fabp7* and *Ncmap* mRNA in sensory ganglia (I-K) smFISH of *Fabp7* and *Ncmap* mRNA in sympathetic ganglia Scale bars are 100 μm and 15 μm (insets)

Schwann cells are readily separated from satellite glia in sensory ganglia, consistent with recent studies (Avraham et al., 2020). Due to their scarcity in sympathetic ganglia, Schwann cells could only be differentiated from satellite glia when we performed an integrated analysis with sensory ganglia (Figures 2A-B) (Satija et al., 2015; Stuart et al., 2019). These results suggest that Schwann cells are prevalent in sensory, but not sympathetic ganglia, a difference that may have important functional implications for modulation of neuronal connectivity, activity, and repair in these two systems.

To identify functional pathways that distinguish satellite glia and Schwann cells, we performed gene ontology analysis of all significantly enriched genes within each group (Figures 2D-E **and Table S1**). Satellite glia were enriched for genes associated with fatty acid metabolism including chaperone proteins (*Fabp5, Fabp7*), elongases (*Elovl2, Elovl5*), desaturases (*Fads1, Fads2, Fads6, Scd1, Scd2*), and fatty acid synthase, *Fasn* (Figure 2D, **Table S1**). Furthermore, satellite glia specifically expressed genes involved in mitochondrial beta-oxidation (*Acaa2, Acadl, Acadm, Acsbg1, Eci1*) (Figure 2D, **Table S1**), which produces energy from fatty acids (Barber and Raben, 2019). Schwann cells were strongly enriched for genes involved in myelination (Figure 2E), including genes specific for sphingolipid synthesis (*Fa2h, Samd8, Sptlc2, Ugt8a*) (**Table S1**), which are abundant myelin components in both central and peripheral nervous systems (Giussani et al., 2021). Schwann cells also expressed fatty acid elongase genes (*Elovl1, Elovl7*) that were distinct from those expressed in satellite glia (**Table S1**). Combined, these data suggest that lipid synthesis in satellite glia and Schwann cell is specialized for neuronal compartments contacted by glial cells.

Satellite glia uniquely ensheathe cell bodies of peripheral neurons, while Schwann cells tightly wrap around axons, yet it is unknown if these contacts are driven by distinct cell adhesion molecules. We found that satellite glia were specifically enriched for 46 cell adhesion molecules, including known neuron-glia interactors (*Atp1b2, Ncam1, Chl1, Cadm1, Cadm2, and Cdh10*), and two genes (*Fbln5, Vcam1*) involved in endothelial cell adhesion (Albig and Schiemann, 2004; Batiuk et al., 2020; Hillenbrand et al., 1999; Kokovay et al., 2012; Sukhanov et al., 2021) (Figure 2D **and Table S1**). Some of the cell adhesion molecules (*Cadm2, Cdh10, Chl1, Megf10, Tnc*) have known roles in regulating synapse formation, neurite outgrowth, and axon pathfinding (Batiuk et al., 2020; Chung et al., 2013; Frei et al., 2014; Joester and Faissner, 2001). Interestingly, some cell adhesion genes (*Chl1, Ncam1, Postn*) expressed in satellite glia are upregulated in Schwann cells in response to neuronal injury (Allard et al., 2018; Zhang et al., 2000).

Satellite glia also expressed specific integrin proteins (*Itga7, Itgav, Itgb3*), collagen proteins (*Col14a1, Col16a1, Col28a1*), and other genes involved in extracellular matrix (ECM) binding (Figure 2D **and Table S1**). The enrichment of ECM-related cell adhesion molecules is consistent with a role for satellite glia in maintaining a healthy and organized neuronal microenvironment in peripheral ganglia. Compared to satellite glia, Schwann cells expressed 29 unique cell adhesion molecules, including genes involved in myelination (*Mag, Mog, Nfasc, Mpdz*), cell-cell adhesion (*Cadm3, Ctnna3*), and cell-ECM adhesion (Cd9, Cd47), as well as several specific integrin proteins (*Itga6, Itgb1, Itgb4, Itgb5*) (Figure 2E, **Table S1**).

Finally, we confirmed the specificity of expression of the satellite glia marker, *Fabp7*, and the Schwann cell marker, *Ncmap,* in sympathetic and sensory ganglia using RNAscope single molecule Fluorescence *in situ* Hybridization (smFISH) (Figures 2F-K). Consistent with the sequencing analysis (Figures 2A-C), *Fabp7* was enriched in both sympathetic and sensory ganglia (Figures 2F and 2I), while *Ncmap* was abundant in sensory ganglia (Figure 2G), but not detected in sympathetic ganglia (Figure 2J).

### Transcriptional heterogeneity of satellite glia

Satellite glia could be separated into five transcriptionally distinct populations, when analyzed independent of all other cell types (Figures 3A-B). Satellite glia are differentiated into individual clusters specific to sensory or sympathetic ganglia, or three shared clusters which are distinguished by enriched expression of immediate early genes (“IEG”), immune response genes (“Immune response”), or cell adhesion and extracellular matrix-related genes (“General resident”).

**Figure 3.**
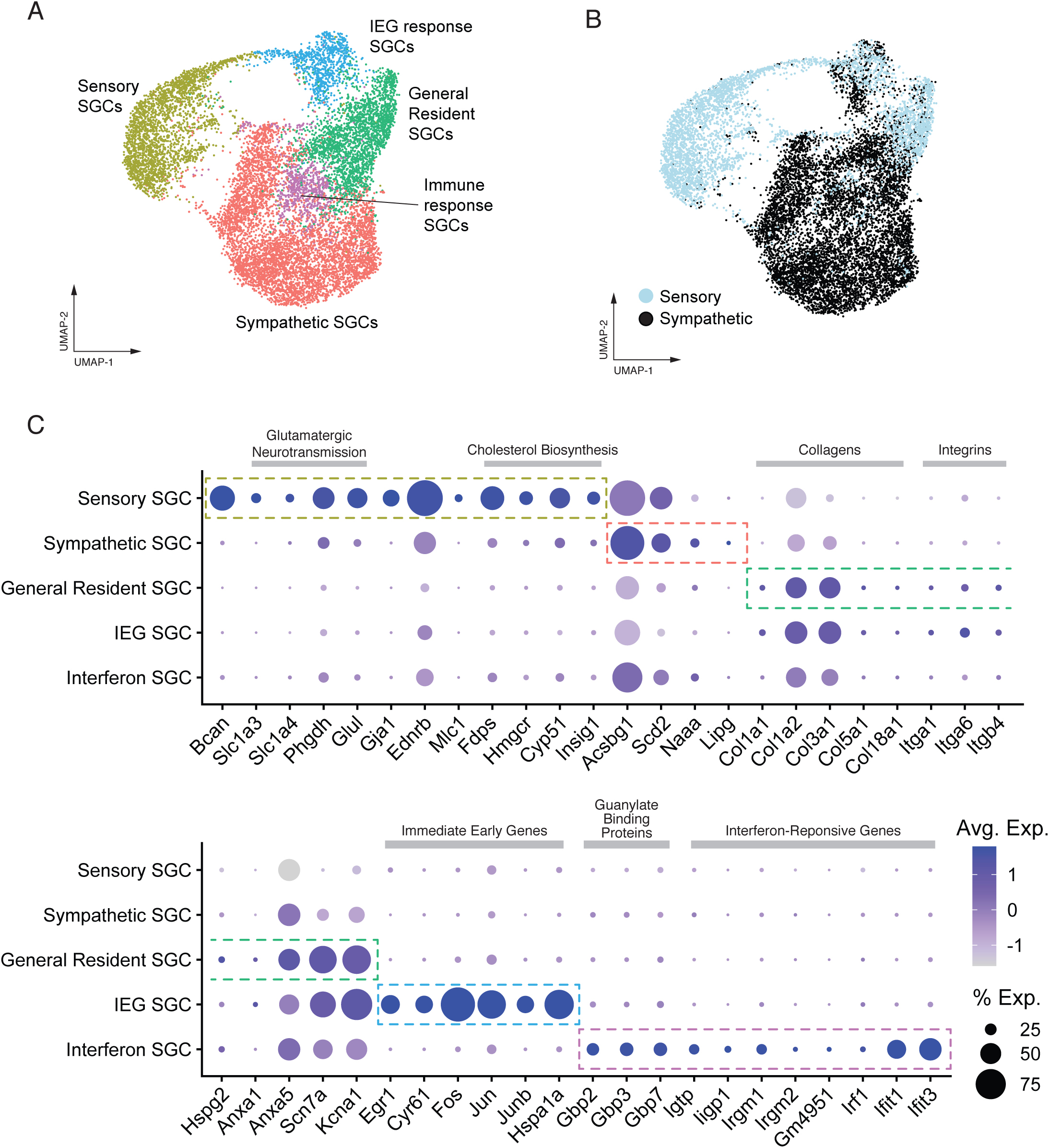
Transcriptional heterogeneity of satellite glial cells. (A) UMAP of satellite glial cell populations in sensory and sympathetic ganglia (B) Same as in (A) overlaid with ganglia of origin (C) Dot plots of satellite glial cell cluster marker genes and enriched transcripts. Dot size is proportional to the percentage of each cluster expressing the marker gene, and the color intensity is correlated with expression level.

### Sensory satellite glia

Consistent with their association to glutamatergic neurons, one of the strongest marker genes of sensory satellite glia was the excitatory amino acid transporter *Slc1a3* (Figures 3C and S3A-B). *Slc1a3* is an established glutamate transporter that acts to uptake excess glutamate from the extracellular space (Kanai and Hediger, 2004). Other enriched glutamatergic machinery in sensory satellite glia included *Slc1a4*, glutamate/neutral amino acid transporter (Kanai and Hediger, 2004), *Phgdh*, an enzyme in serine biosynthesis which influences glutamate receptor activity (Neame et al., 2019), and *Glul*, a glutamine synthetase which catalyzes the conversion of glutamate to glutamine (Rose et al., 2013) (Figure 3C). *Glul* mRNA is also found in non-glial cell types in the sensory ganglia, as previously reported (Avraham et al., 2020).

In addition to glutamate-associated genes, we also found striking enrichment of *Gja1*, a gap junction protein that mediates sensory neuron coupling and chronic pain in response to injury (Kim et al., 2016); *Ednrb* endothelin receptor B, which is involved in Schwann cell maturation and myelination (Brennan et al., 2000; Swire et al., 2019); and *Mlc1*, a putative membrane protein of unknown function that has been linked to a neurological disorder, megalencephalic leukoencephalopathy (Schmitt et al., 2003) (Figure 3C). Using smFISH, we confirmed that *Mlc1* transcript is more enriched in sensory satellite glia compared to sympathetic glia (Figures 4A-B). The most specific and highly expressed marker of sensory satellite glia was the proteoglycan, *Bcan* (Brevican), a structural component of the brain ECM which has been implicated in regulating neurite outgrowth, glioma cell motility, synaptic plasticity, and axon regeneration (Frischknecht and Seidenbecher, 2012) (Figure 3C). Of note, astrocyte-derived brevican inhibits outgrowth in cultured cerebellar granule neurons in the brain (Yamada et al., 1997). Since sensory satellite glia inhibit neurite outgrowth in neuron-glia co-cultures (De Koninck et al., 1993) and sensory neurons do not bear dendritic processes, it is tempting to speculate that satellite glia-derived Brevican may play a specific role in suppressing dendrite morphogenesis in sensory ganglia.

**Figure 4.**
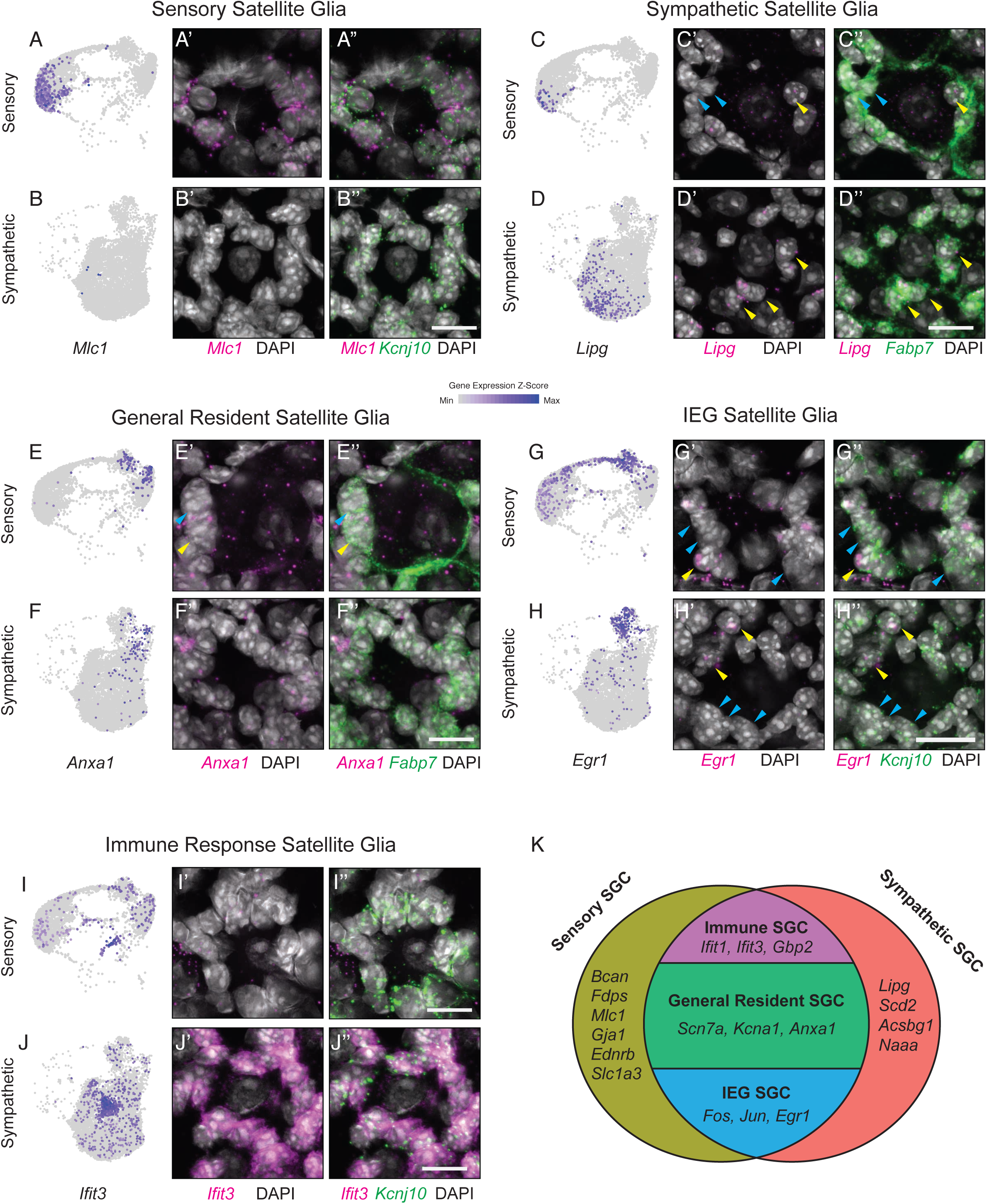
Histological validation of satellite glial cell type markers. (A-B) UMAP expression plot of *Mlc1* mRNA in sensory (A) and sympathetic ganglia (B) (A’-B’’) smFISH of *Mlc1* (magenta) and *Kcnj10* (green) mRNA in sensory (A’-A’’) and sympathetic (B’-B’’) ganglia. (C-D) UMAP expression plot of *Lipg* mRNA in sensory (C) and sympathetic ganglia (D) (C’-D’’) smFISH of *Lipg* (magenta) and *Fabp7* (green) mRNA in sensory (C’-C’’) and sympathetic (D’-D’’) ganglia. (E-F) UMAP expression plot of *Anxa1* mRNA in sensory (E) and sympathetic ganglia (F) (E’-F’’) smFISH of *Anxa1* (magenta) and *Fabp7* (green) mRNA in sensory (E’-E’’) and sympathetic (F’-F’’) ganglia. (G-H) UMAP expression plot of *Egr1* mRNA in sensory (G) and sympathetic ganglia (H) (G’-H’’) smFISH of *Egr1* (magenta) and *Kcnj10* (green) mRNA in sensory (G’-G’’) and sympathetic (H’-H’’) ganglia. (I-J) UMAP expression plot of *Ifit3* mRNA in sensory (I) and sympathetic ganglia (J). (I’-J’’) smFISH of *Ifit3* (magenta) and *Kcnj10* (green) mRNA in sensory (I’-I’’) and sympathetic (J’-J’’) ganglia. Yellow arrows indicate satellite glia positive for the gene of interest, while cyan arrows indicate satellite glia negative for the gene of interest Scale bars are 15 μm. (K) Transcriptional classification of sympathetic and sensory satellite glia subtypes.

Notably, sensory satellite glia were enriched in transcripts associated with cholesterol biosynthesis and turnover including *Hmgcr, Fdps*, *Cyp51*, and *Insig1* (Li et al., 2021; Luo et al., 2020) (Figure 3C). The enrichment of cholesterol synthetic machinery in sensory ganglia suggests a greater need for exogenously supplied lipids than in sympathetic ganglia. Altogether, we identified genes in sensory satellite glia that are involved in neuronal maintenance, signal transduction, and neurotransmission.

### Sympathetic satellite glia

We found few transcripts that are unique to sympathetic satellite glia. Among the few enriched transcripts within sympathetic satellite glia, we detected components of fatty acid biosynthesis, activation, beta-oxidation, and lipoprotein uptake/metabolism, including *Scd2, Acsbg1*, *Naaa*, and *Lipg* (McCoy et al., 2002; Miyazaki et al., 2005; Pei et al., 2003) (Figure 3C). Using smFISH we confirmed that one of these genes, *Lipg* (endothelial lipase G), was enriched in sympathetic glia (Figures 4C-D **and** S3C-D). Although, sympathetic satellite glia are in direct contact with cholinergic synapses between pre- and post-ganglionic neurons (Hanani, 2010), we were unable to identify transcripts involved in cholinergic neurotransmission, in contrast to the specific glutamatergic neurotransmission-related transcripts detected in sensory ganglia.

### General resident satellite glia

General resident satellite glia were found in both sensory and sympathetic ganglia and were distinguished by the expression of many genes related to ECM interactions. These genes included collagen proteins (*Col1a1, Col1a2, Col3a1, Col5a1, Col18a1*), integrins (*Itga1, Itga6, Itgb4*), two extracellular annexin proteins (*Anxa1, Anxa5*), and the proteoglycan *Hspg2* (Figure 3C). In addition to ECM-related transcripts, this cluster also featured the ion channel genes, *Scn7a* and *Kcna1* (Figure 3C), which are likely key players in the regulation of extracellular ion concentrations. Interestingly, mutations in two genes, *Kcna1* and *Hspg2*, have been linked to aberrant neuronal excitability and peripheral neuropathies (Bangratz et al., 2012; Browne et al., 1994), though these effects have been primarily attributed to their expression in neurons or Schwann cells (Bangratz et al., 2012; Browne et al., 1994).

Annexin A1 and A5 (*Anxa1 and Anxa5*), which were enriched in the General resident glial population (Figure 3C), are members of the Annexin family of Ca^2+^- and phospholipid-regulated proteins that mediate membrane trafficking events (Gerke et al., 2005). Anxa5 binds with high affinity to phosphatidylserine expressed on the surface of apoptotic cells (Koopman et al., 1994). In the DRG, satellite glia precursors, and not the canonical macrophages, are responsible for clearing neuronal corpses during naturally occurring cell death (Wu et al., 2009), raising the possibility that the General resident satellite glial population performs this function in both sensory and sympathetic ganglia. Further, *Anxa1* is upregulated in sensory ganglia in response to inflammation and attenuates nociceptive responses (Chen et al., 2014). Using smFISH, we detected *Anxa1* expression in a subset of satellite glial cells in both sensory and sympathetic ganglia (Figures 4E-F **and** S3E-F), suggesting that this population may have a general role in anti-inflammatory responses.

### Immediate early gene-expressing satellite glia

A shared population of satellite glia between sensory and sympathetic ganglia expressed many classical immediate early genes, including *Egr1, Cyr61, Fos*, *Jun*, *Junb*, and *Hspa1a* (Figure 3C). The upregulation of immediate early genes is a hallmark of neuronal activity (Guzowski et al., 2005), and in some cases of neuronal injury (Arthur-Farraj et al., 2012). In particular, satellite glia up-regulate c-Fos and c-Jun immediately after sciatic nerve injury (Soares et al., 2001). We observed satellite glia-specific *Egr1* expression in both sensory and sympathetic ganglia using smFISH (Figures 4G-H, S3G-H). Detection of *Egr1* in ganglia by smFISH suggests that IEG-expressing satellite glia are present under physiological conditions, and are not a result of dissociation-induced up-regulation in the course of isolating cells for sequencing.

### Immune-responsive satellite glia

A separate shared cluster of satellite glia was demarcated by the specific expression of a cohort of guanylate binding proteins (*Gbp2, Gbp3, Gbp7*) and related interferon-inducible GTPases (*Igtp, Iigp1, Irgm1, Irgm2, Gm4951*), which have established roles in intracellular defense against pathogens (Ngo and Man, 2017; Tretina et al., 2019) (Figure 3C). The enrichment of these transcripts, in addition to other interferon-inducible genes (*Irf1, Ifit1, Ifit3*) (Figure 3C) that have anti-viral properties (Fensterl and Sen, 2015), suggests that this population defends against viral and bacterial pathogens. Further, these cluster was also enriched for transcription factors *Irf1* and *Stat1* (Figure 3C), which are induced by interferon B signaling (Tretina et al., 2019). Strikingly, we found that Immune-responsive satellite glia were disproportionately enriched in sympathetic ganglia (Figures 4I-J, S3I-J), in contrast to the other two shared clusters of satellite glia that were more evenly distributed between both ganglia. We used smFISH to validate one of the genes, *Ifit3*, and found that *Ifit3*-positive satellite glia were indeed more abundant in sympathetic ganglia (Figures 4I-J, S3I-J). Interestingly, we noticed that *Ifit3*^+^ satellite glial cells were concentrated in specific regions within the SCG (Figure S3J). These findings suggest that despite the presence of resident macrophages (Pirzgalska et al., 2017), satellite glia are primed to defend the peripheral nervous system against bacterial or viral infections.

## DISCUSSION

In this study, we describe a high-resolution transcriptional comparison of glial cell types between functionally and anatomically distinct ganglia in the peripheral nervous system. Compared to the well-characterized diversity in neuronal populations in peripheral ganglia, little is known about the cellular and molecular diversity of satellite glia that are intimately associated with cell bodies of peripheral neurons. Our analysis reveals five molecularly defined satellite glial cell types (Figure 4K), provides a rich resource of gene expression in satellite glia and Schwann cells, and suggests that satellite glia are transcriptionally tuned to their resident ganglia.

Our analyses revealed that satellite glia in sensory and sympathetic ganglia exhibit similarities, but also significant differences in gene expression, which may reflect ganglia-specific functions. Both sensory and sympathetic satellite glial cells express >70 genes involved in fatty acid/cholesterol synthesis and metabolism, including highly expressed transcripts such as *ApoE, Fabp7, and Fasn*. In general, neurons are inefficient in lipid synthesis, and rely on the uptake of lipids from extrinsic sources (Pfrieger and Ungerer, 2011). Thus, satellite glia may be an important source for the production and release of lipids to their neuronal neighbors. In the mouse brain, astrocyte-derived ApoE regulates neuronal cholesterol biosynthesis and epigenetic mechanisms to promote memory consolidation (Li et al., 2021). *Fabp7* deletion in mice elicits deficits in pre-pulse inhibition, dendrite complexity, and spine density (Bruce et al., 2017; Ebrahimi et al., 2016; Watanabe et al., 2007). Loss of *Fasn* in sensory satellite glia impairs axonal regeneration following injury (Avraham et al., 2020). Together, these results imply critical functions for peripheral satellite glia in regulating neuronal metabolism, epigenetic states, morphology, synaptic function, and nerve repair, through lipid-mediated communication. However, we also observed differences in the lipid pathways that were enriched in sensory *versus* sympathetic satellite glia, suggesting specialized functions within each ganglia. Sensory satellite glia were specifically enriched in genes associated with cholesterol biosynthesis, important for maintenance of neuronal processes, myelination, vesicle formation, and synaptic transmission (Barber and Raben, 2019; Dietschy and Turley, 2004). The majority of cholesterol (∼70-80%) in the adult brain is in myelin sheaths that insulate axons (Dietschy and Turley, 2004). Since sensory, but not sympathetic ganglia, contain myelinating Schwann cells which are primarily engaged in membrane synthesis, it is likely that sensory satellite glia provide an additional source for extrinsic cholesterol supply. Consistent with this notion, a substantial fraction of lipids for CNS myelin formation is contributed by astrocytes in addition to the canonical role of oligodendrocytes (Camargo et al., 2017).

Within sensory satellite glia, we also observed specific enrichment of genes involved in glutamate uptake and recycling, consistent with the presence of glutamatergic neurons in sensory, but not sympathetic ganglia. We also observed enriched expression of Connexin 43 (*Gja1*), which is up-regulated following injury in sensory satellite glia and promotes gap junction-mediated coupling between adjacent neurons, a form of neuronal plasticity that contributes to neuropathic pain (Kim et al., 2016). Compared to sensory satellite glia, we found fewer transcripts specific to sympathetic satellite glia, and none that were obviously involved in regulation of cholinergic neurotransmission. Sympathetic satellite glia were largely distinguished by genes involved in fatty acid biosynthesis and metabolism, e.g., *Scd2*, *Acsbg1*, and *Lipg*, which could be play a role in supplying building blocks for dendrite and synapse morphogenesis, as well as modulate neurotransmission in sympathetic ganglia.

Our work is consistent with recent transcriptional studies of satellite glial cells (Avraham et al., 2020; Tasdemir-Yilmaz et al., 2020; van Weperen et al., 2021). However, we expand the repertoire of sensory satellite glial cell markers, and reveal new clusters of shared satellite glial cells that are present in functionally disparate peripheral ganglia. Specifically, we define general resident, IEG-expressing, and immune-responsive satellite glia populations and we confirmed the expression of markers by smFISH. One of the shared satellite glial cell types contains signatures of immune responsiveness, including interferon-induced transcripts and genes involved in antiviral defense (Ngo and Man, 2017; Tretina et al., 2019). Although shared, these glia were more abundant in sympathetic ganglia and may have a specific role in protecting peripheral neurons against viral or bacterial pathogens. Notably, while we focused on the SCG, it remains unclear whether the 5 identified sympathetic satellite glial sub-types exist across the entire sympathetic chain, or if each ganglion has its own unique set of glial populations. A recent analysis of the stellate ganglion identified 5 satellite glial populations, including a subtype enriched for *Kcna1* and *Scn7a* (van Weperen et al., 2021), which corresponds well with the General resident satellite glia in our data, as well as a subtype enriched for genes involved in “interferon signaling”, which is similar to the immune-response satellite glia revealed by our analysis. Additionally, the authors noted a population of satellite glia which expressed high levels of immediate early genes, though it was excluded from their analysis. Thus, the 3 shared populations of satellite glia that we identified may be present throughout the sympathetic chain ganglia.

In summary, we provide a resource for satellite glia in functionally distinct peripheral ganglia, which provides a framework for future investigations of ganglion-specific satellite glia populations during development, injury, and disease. The identification of new markers for satellite glia in sensory and sympathetic ganglia can be used to genetically target these cells for imaging, ablation, and optogenetic studies. Recent studies suggest that satellite glia could serve as important targets for interventions in chronic pain (Hanani and Spray, 2020; Kim et al., 2016; Xie et al., 2017) or heart disease, and the knowledge gained from this study can be used to generate relevant animal models to study human disease.

## QUANTIFICATION AND STATISTICAL ANALYSES

Information for statistical analyses for all experiments are provided in figure and table legends.

## Acknowledgements

We thank all members of the Kuruvilla, Zhao, and Hattar labs for helpful comments on the project, and the NIMH Microarray Core Facility for providing library quality control and sequencing. This work was supported by NIH R01 awards, NS073751 and NS107342, to R.K., DC016065 and EY027202 to H.Z, and NIMH intramural research funds (ZIAMH002964) to S.H., a NSF GRFP award to A.M., and NIH Training grant T32GM007231 to the JHU CMDB graduate program for A.M, M.T, and E.B.

## Author Contributions

A.M and M.T contributed to study design, investigations, data analyses, and writing and editing the manuscript. E.B. assisted with experiments. R.K, H.Z, and S.H contributed to writing/editing the manuscript and funding acquisition.

---

**Supplemental Figure S1.**
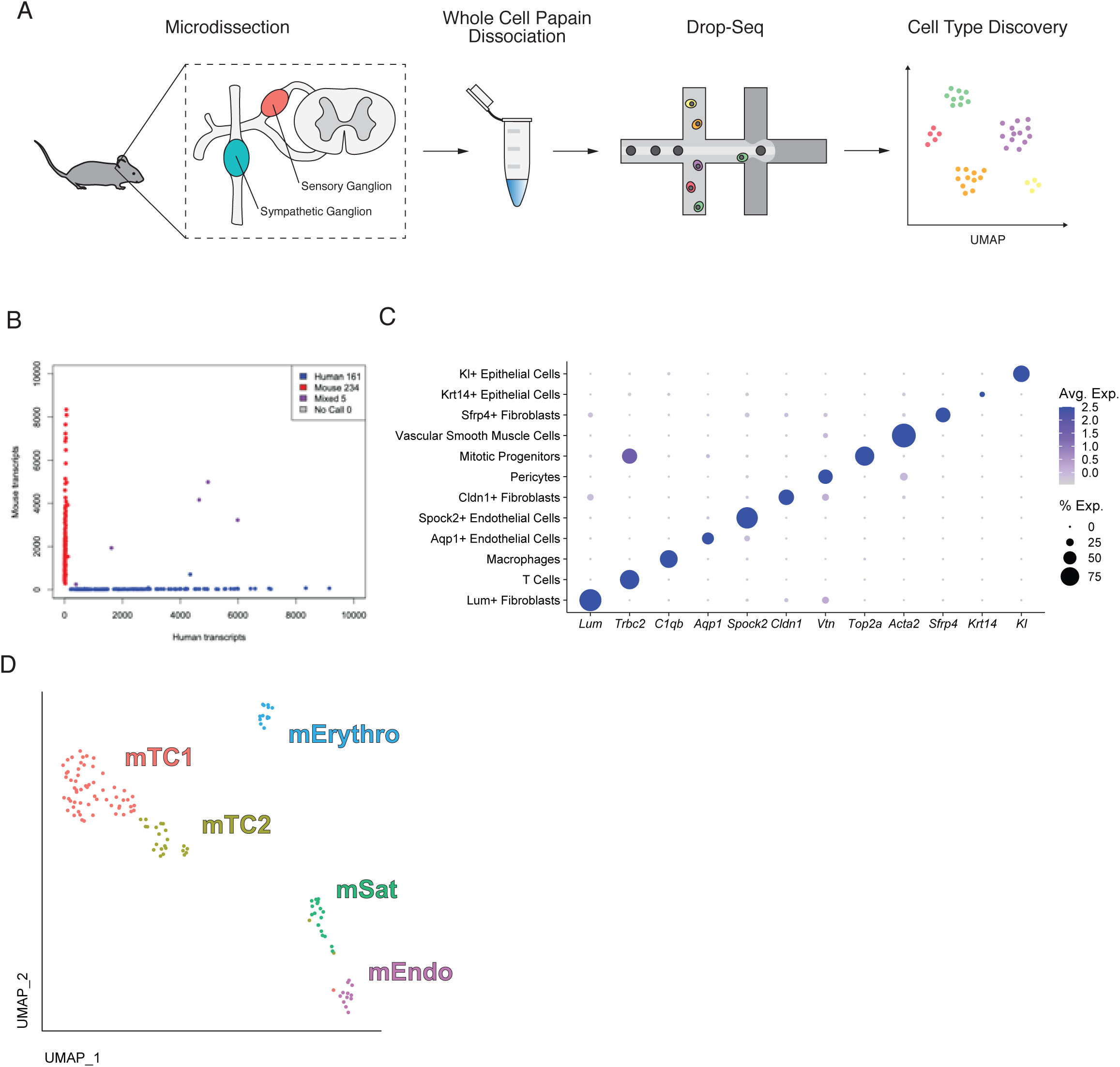
Drop-Seq workflow, validation, and marker genes, Related to Figure 1. (A) Workflow of Drop-seq single cell RNA sequencing (B) Barnyard plot of mixed-species Drop-seq. NIH-3T3 (mouse) and HEK293 (human) cells sequenced in equal mixture results in 5/400 (1.25%) detectable cell-cell doublets. (C) Dot plot of marker genes identified in non-neuronal and non-glial cells (D) UMAP plot of proliferating cells within sensory and sympathetic ganglia (mTC1/2: T-cells, mErythro: erythrocytes, mEndo: endothelial cells, mSat: satellite glia)

**Supplemental Figure S2.**
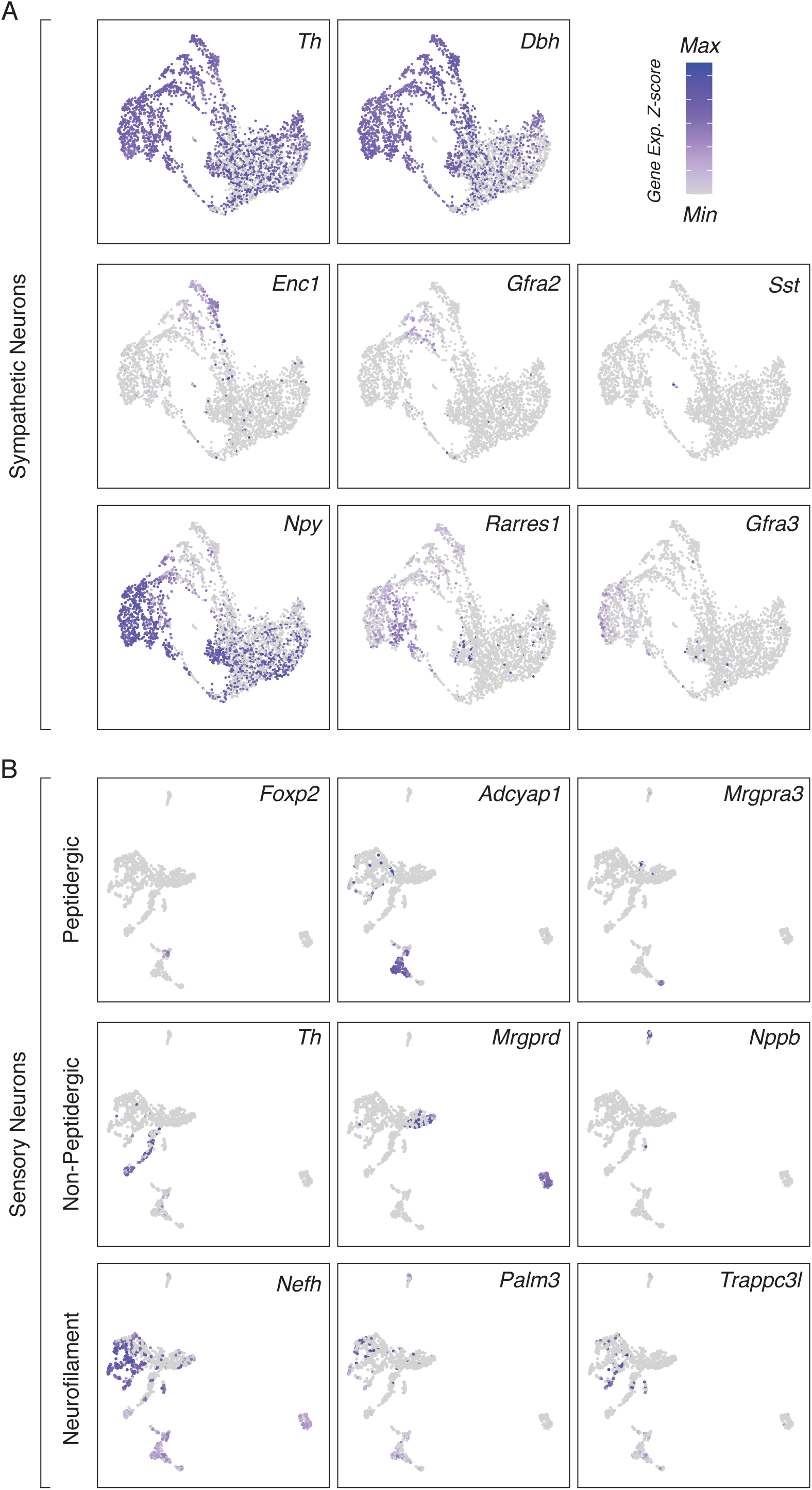
Heterogeneity of peripheral neurons, Related to Figure 2. (A) UMAP expression plots of subtypes of adrenergic sympathetic neurons (B) UMAP expression plots of subtypes of peptidergic, non-peptidergic, and neurofilament-expressing sensory neurons

**Supplemental Figure S3.**
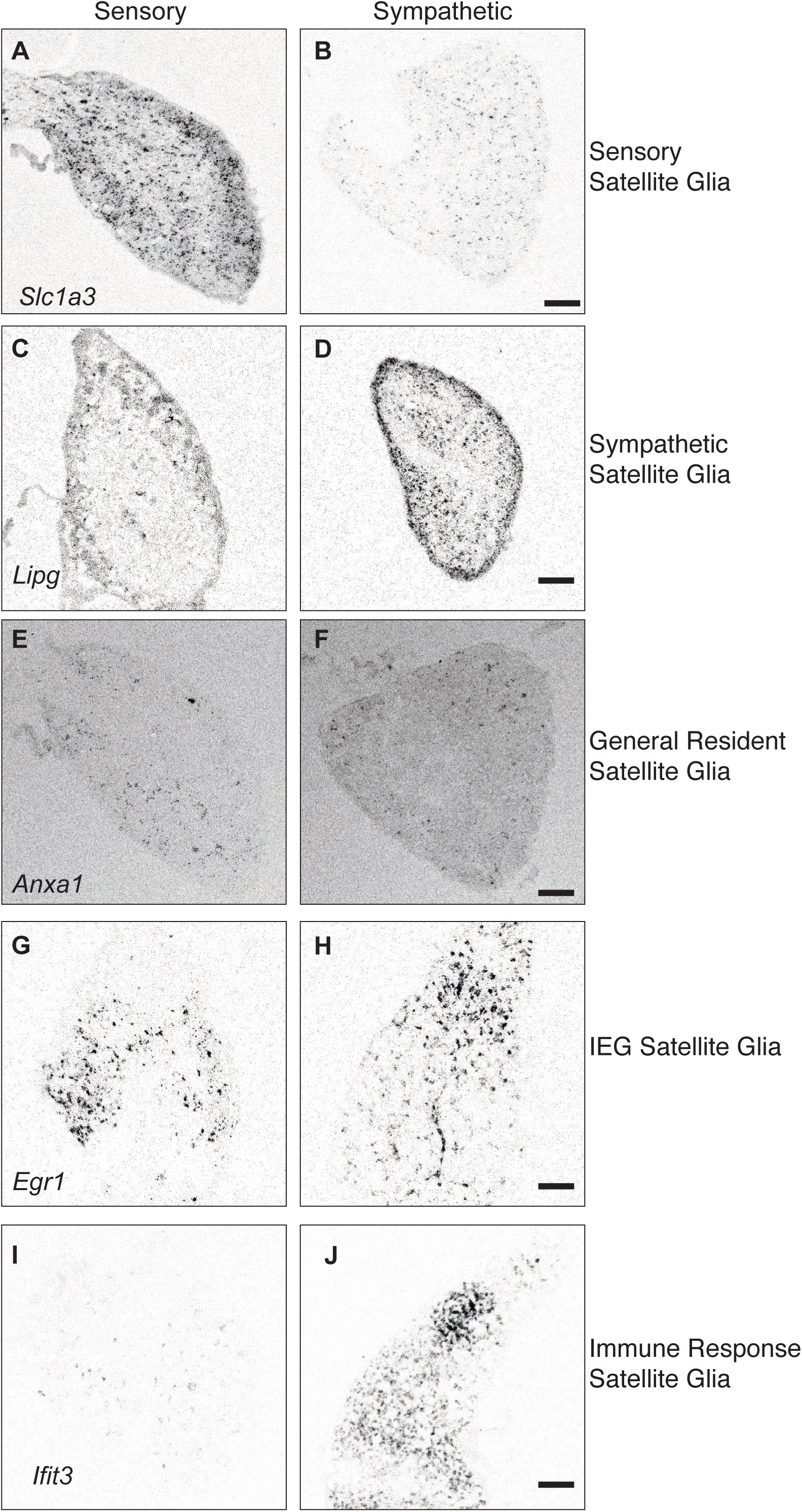
Satellite glia heterogeneity between sympathetic and sensory ganglia, Related to Figure 3. (A-B) smFISH of *Slc1a3* mRNA in sensory (A) and sympathetic ganglia (B). (C-D) smFISH of *Lipg* mRNA in sensory (C) and sympathetic ganglia (D). (E-F) smFISH of *Anxa1* mRNA in sensory (E) and sympathetic ganglia (F). (G-H) smFISH of *Egr1* mRNA in sensory (G) and sympathetic ganglia (H). (I-J) smFISH of *Ifit3* mRNA in sensory (I) and sympathetic ganglia (J). Scale bars are 100 µm.

## STAR METHODS

### KEY RESOURCES TABLE

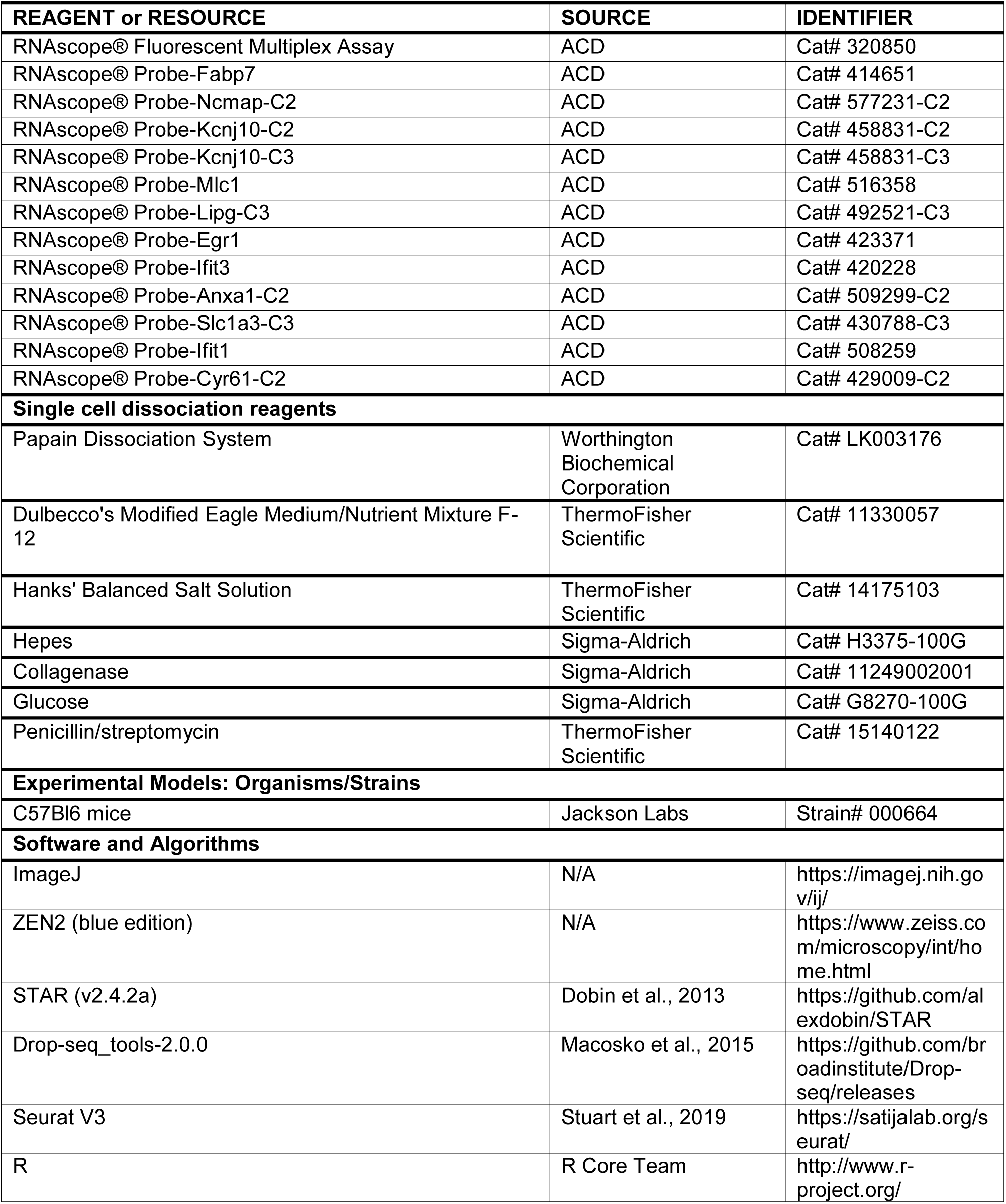
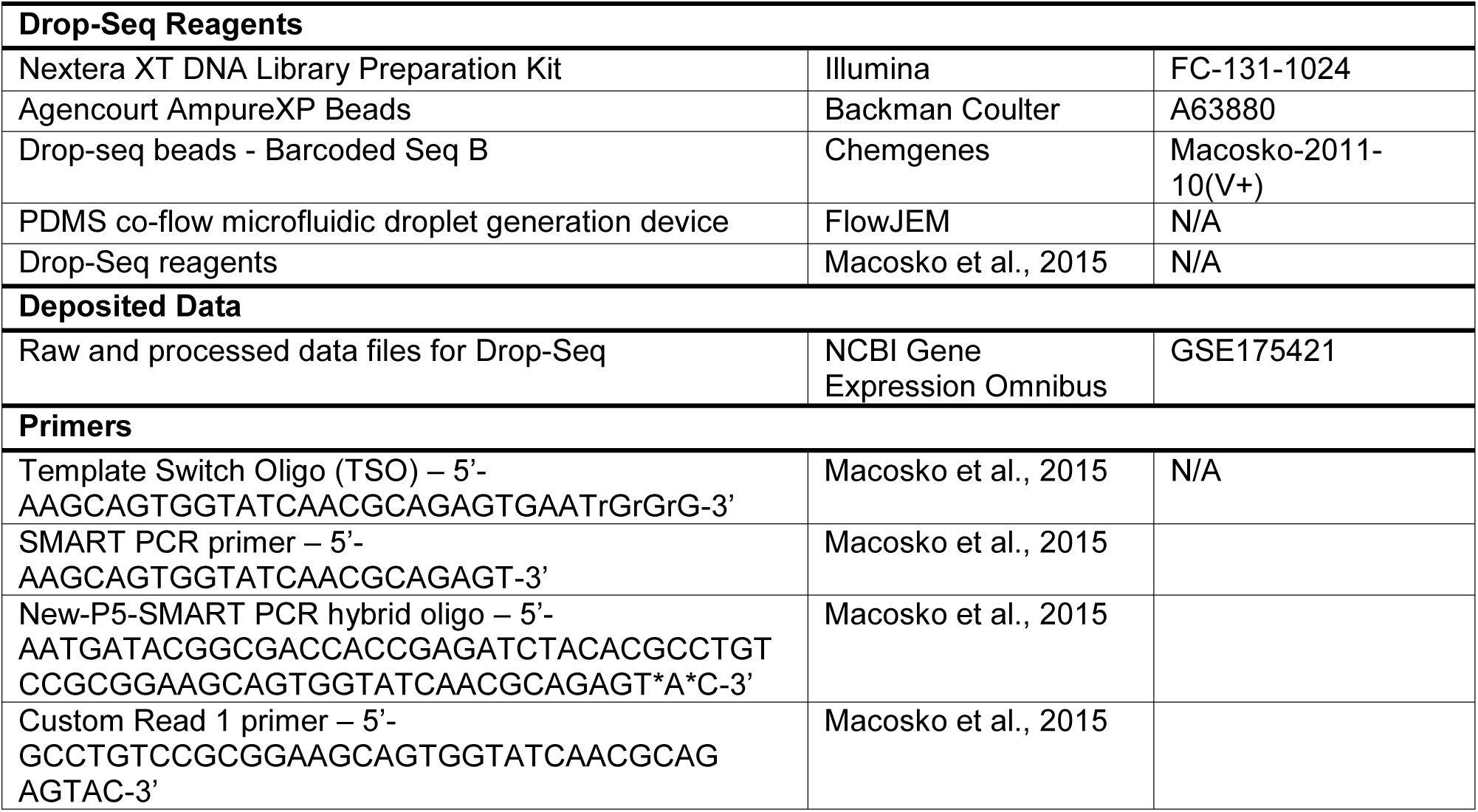

### CONTACT FOR REAGENT AND RESOURCE SHARING

Further information and requests for resources and reagents should be directed to and will be fulfilled by the Lead Contact, Rejji Kuruvilla.

## EXPERIMENTAL MODEL AND SUBJECT DETAILS

### MATERIALS & METHODS

#### Animal care and housing

Adult *C57Bl6* mice (stock no. 000664; Jackson Labs), aged between postnatal days P30-P45, were used for all experiments. Male and female mice were included in all experiments. Mice were housed under standard conditions with access to food and water *ad* libitum. All experimental procedures were performed in accordance with guidelines of the Animal Care and Use Committee of Johns Hopkins University.

#### Drop-Seq mixed species validation

HEK293 (human) and NIH-3T3 (mouse) cells were grown separately in culture until nearly confluent, then dissociated with Trypsin and resuspended to a concentration of approximately 50,000 cells/ml. An equal amount of mouse and human cells were combined in a single tube which was used as input to the Drop-Seq setup using the parameters described in the ‘Drop-Seq single cell partitioning, library preparation’ section below. For mixed species experiments, completed sequencing libraries were run on a MiSeq and reads were demultiplexed and aligned using the Drop-Seq Tools 1.0 pipeline. Barnyard plots for mixed species experiments were generated in R.

#### SCG/DRG tissue isolation and preparation of single cell suspensions

Dissections and single cell collections were performed between 9 a.m-2 p.m. Multiple sympathetic or sensory ganglia were pooled together for each round of Drop-Seq. Ganglia were incubated in 50 units Papain diluted in HBSS plus HEPES for 20 min at 37°C then washed with HBSS plus HEPES and incubated for an additional 20 min at 37°C with 1.5mg/mL collagenase in HBSS plus HEPES. Following the second incubation, ganglia were washed again and then suspended and triturated in DMEM AIR (DMEM F12 supplemented with 12.5mM glucose and 1U/ml penicillin/streptomycin). Cells were centrifuged and resuspended in fresh DMEM AIR.

#### Drop-Seq single cell partitioning, library preparation, and sequencing

Single cell suspensions were diluted to a concentration of 100 cells/μl and processed with the Drop-Seq protocol as previously described (Macosko et al., 2015). Flow rates for cells, beads, and oil were optimized for aquapel-treated PDMS devices purchased from FlowJem (cells and beads: 2,300 μl/hour, oil: 13,000 μl/hour). Up to two samples were processed in series, with single cell suspensions and stable emulsions held on ice until all collections were completed (no more than 1 hour) before proceeding immediately with reverse transcription. cDNA amplification was performed using 4000 beads/reaction with a total of 15 cycles of PCR. Sequencing libraries were generated from amplified cDNA with the Illumina Nextera XT Library Prep kit and up to 6 libraries were multiplexed for sequencing on an Illumina NextSeq500 platform. Sequencing was performed by the NIMH Microarray Core Facility (Bethesda, MD).

#### Drop-Seq data preprocessing

Reads were demultiplexed and aligned to the mouse genome (mm10), and digital gene expression matrices were generated using the Drop-Seq Tools 2.0.0 pipeline (https://github.com/broadinstitute/Drop-seq/releases).

#### PCA, clustering, and differential gene expression analysis

Digital gene expression matrices for all samples were imported and processed with the Seurat package (Satija et al., 2015; Stuart et al., 2019). Cells containing fewer than 200 genes or greater than 7.5% mitochondrial gene content were removed prior to data normalization and scaling. Principal component analysis was used to identify major sources of variation within the dataset, and gene loadings within each PC were manually inspected to ensure the capture of biologically relevant signals. For the initial round of clustering, 20 PCs were included as input to clustering and dimensionality reduction, resulting in the identification of 9 major cell classes. Cell types were identified by the overlay of canonical marker genes. Following cell type identification, each major cluster was isolated and analyzed using iterative rounds of PCA to identify finer substructure among each class. Differential gene expression testing (Wilcox rank sum test) was used to identify cell type specific marker genes.

#### Gene Ontology analysis

Gene ontology (GO) analysis was performed using the Database for Annotation, Visualization and Integrated Discovery (DAVID) (Huang da et al., 2009).

#### Data and code availability

The accession number for raw and processed data from Drop-Seq experiments reported in this paper is GEO: GSE175421. Data from this study are available from the corresponding author upon request.

#### RNAscope

Superior cervical ganglia and L3-L5 dorsal root ganglia were dissected from P30-45 *C57Bl6* mice, cryo-protected in 30% sucrose in PBS for one hour and embedded in OCT prior to being frozen at −80°C. Ganglia were cryo-sectioned at 14 µm and kept at −80°C until RNAscope was performed. Target mRNA was probed using RNAscope^®^ Multiplex Fluorescent Reagent Kit v2 Assay. Tissues were incubated in fresh 4% paraformaldehyde for five minutes, washed twice in 1xPBS, and dehydrated with increasing concentrations of ethanol. Subsequently, the tissues were treated with hydrogen peroxide for 10 minutes and protease treatment for 15 minutes. The RNAscope assay was performed following the manufacturer’s instructions. *Kcnj10* and *Fabp7* are two genes known to be specific to satellite glia and were used to identify subtypes of satellite glia (Hanani and Spray, 2020).

## Supplemental Text and Figures

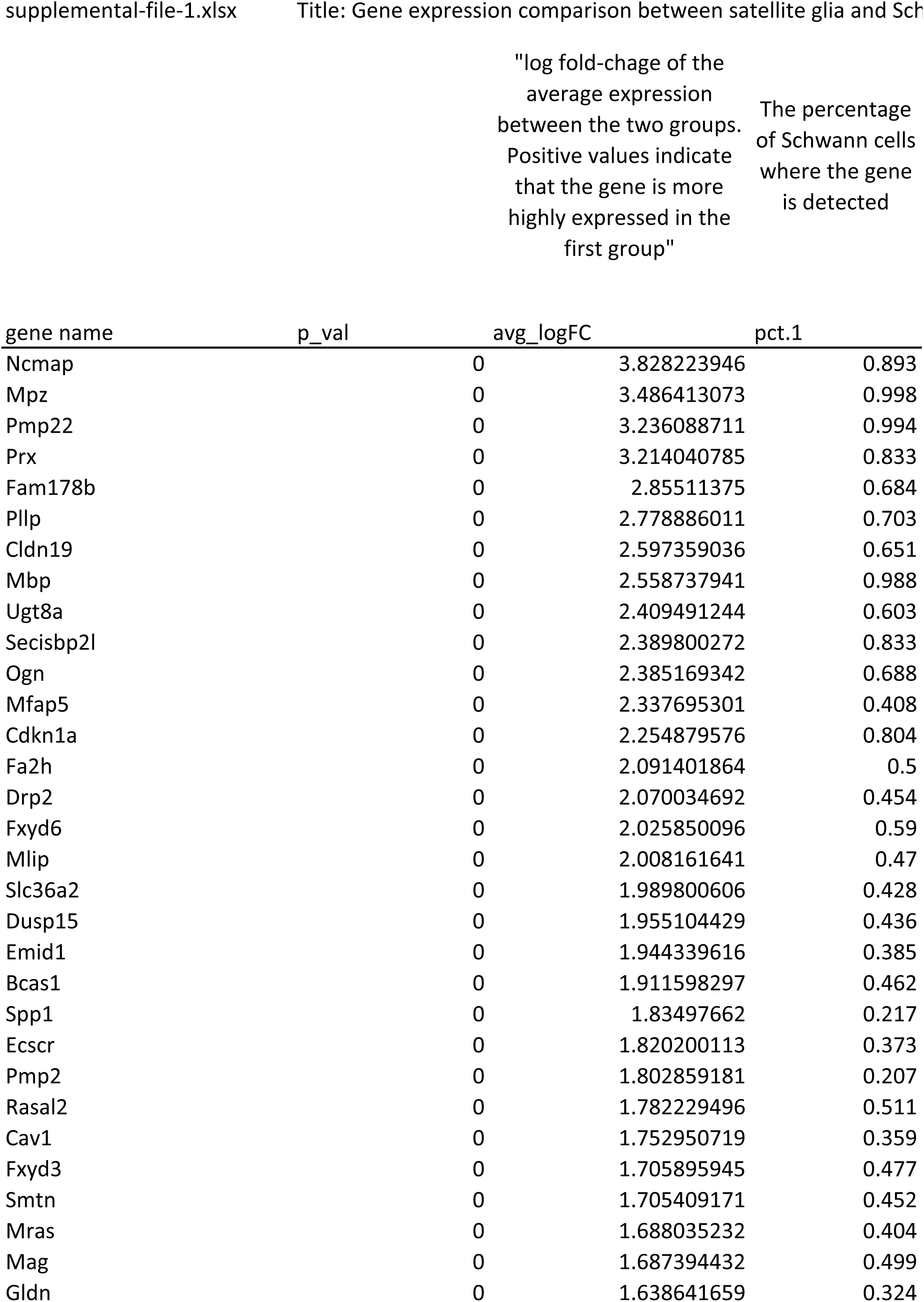
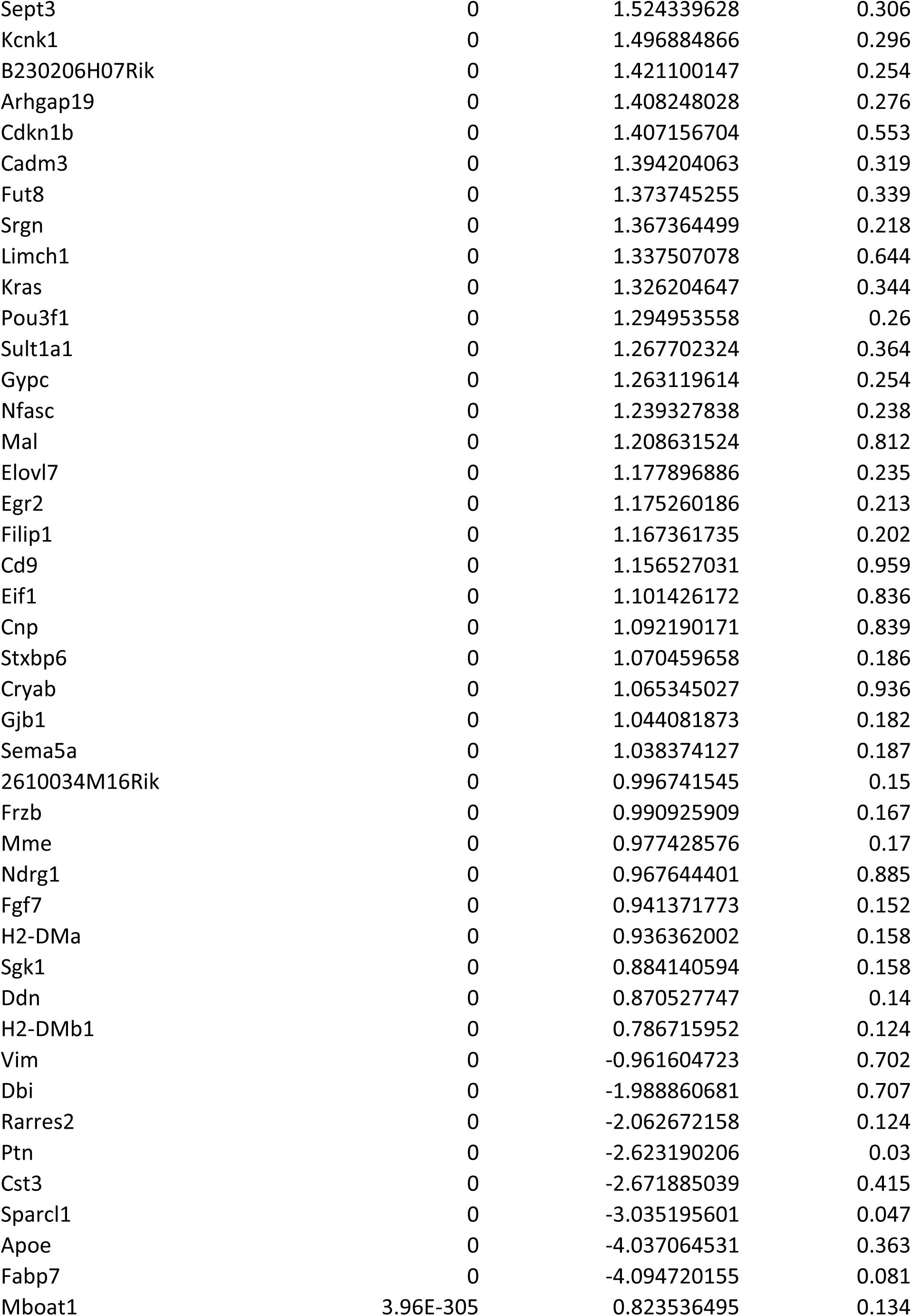
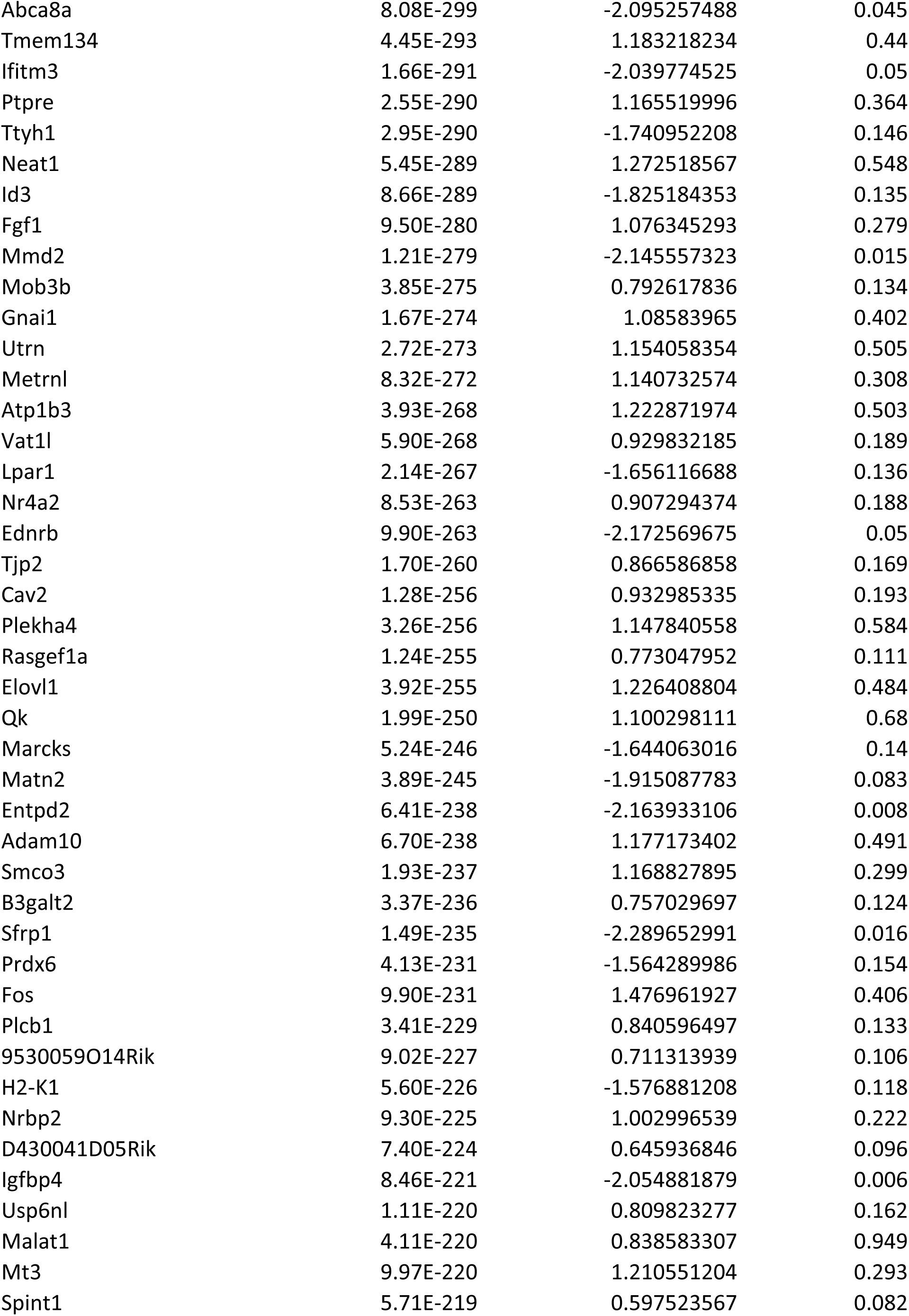
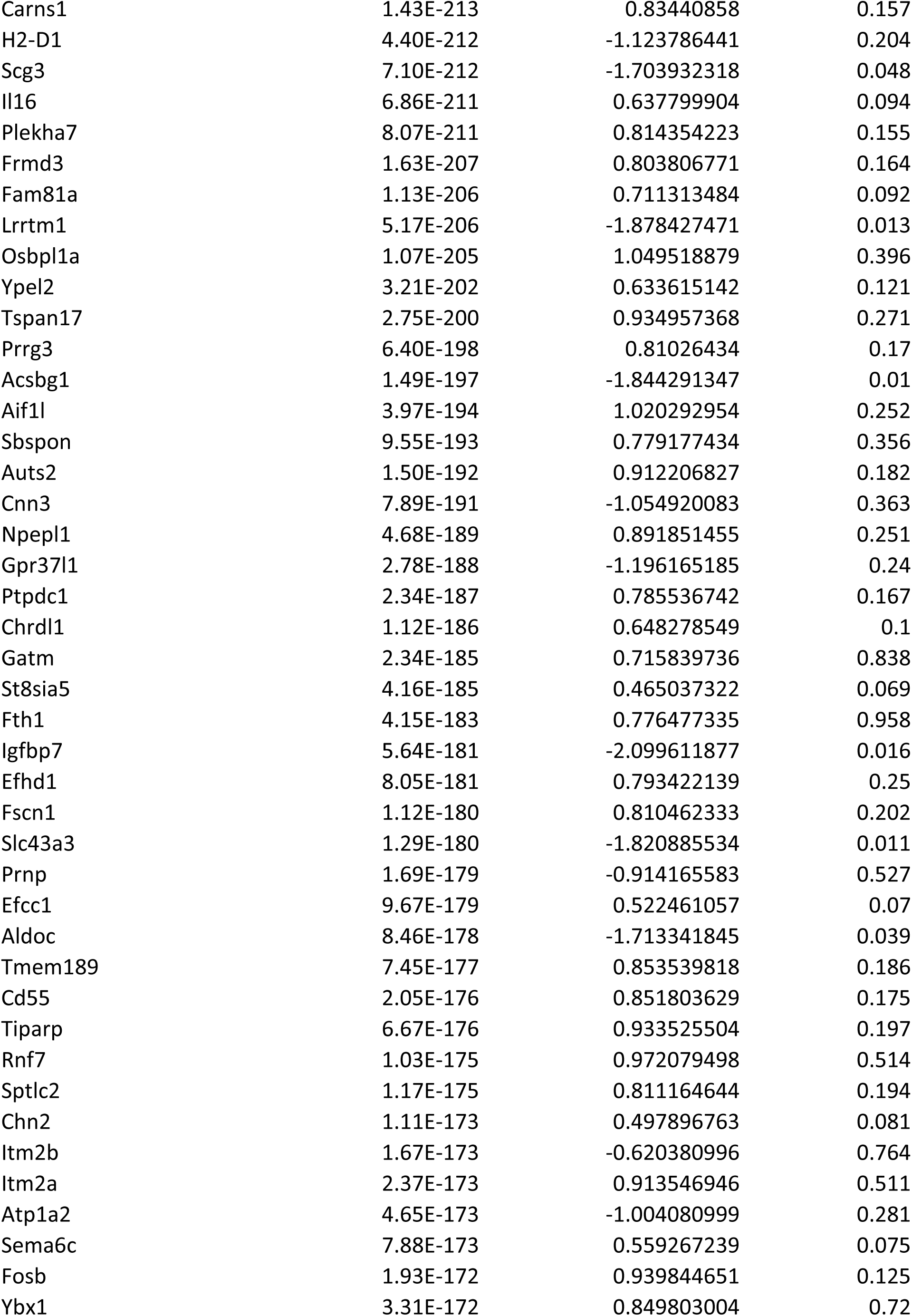
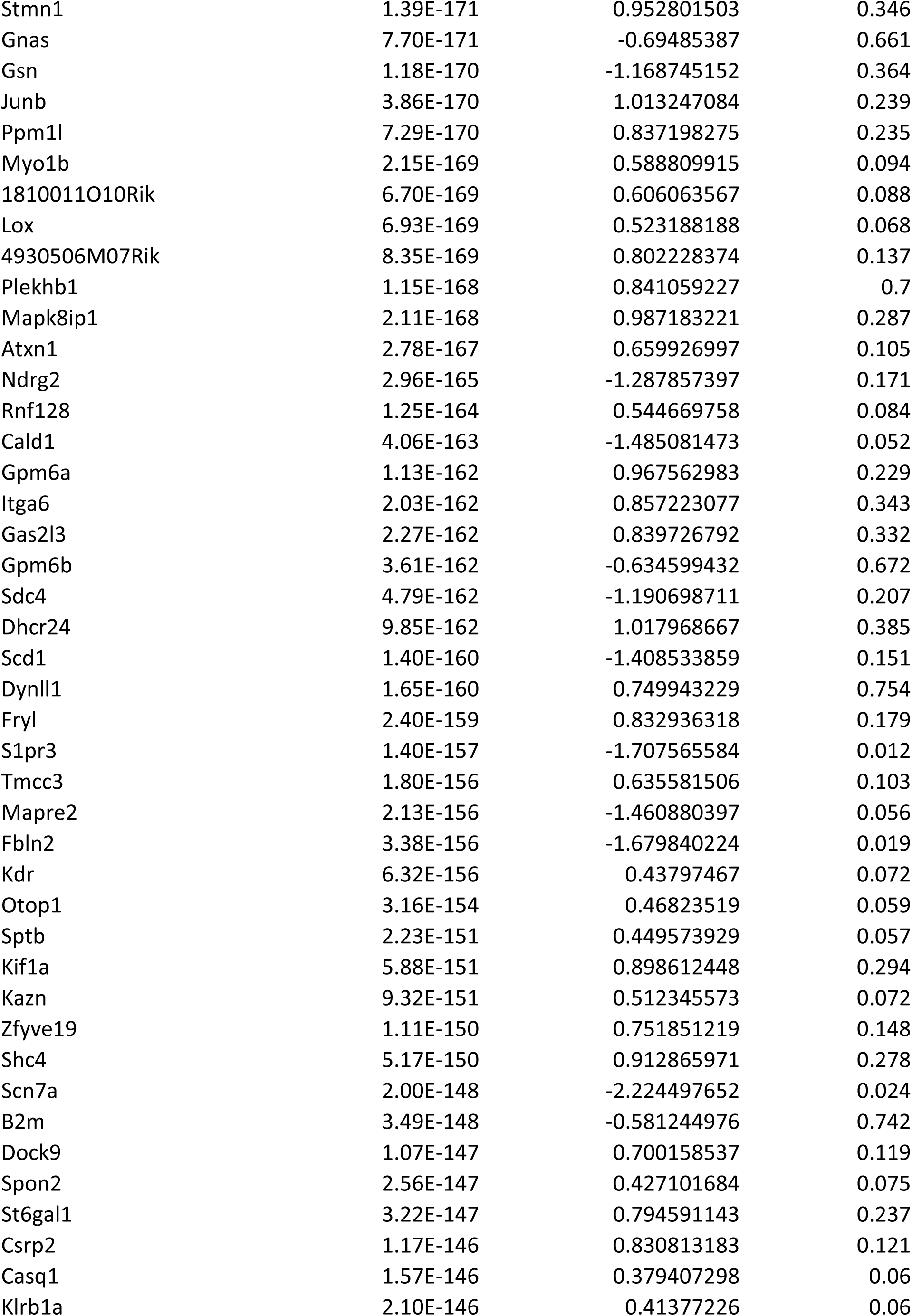
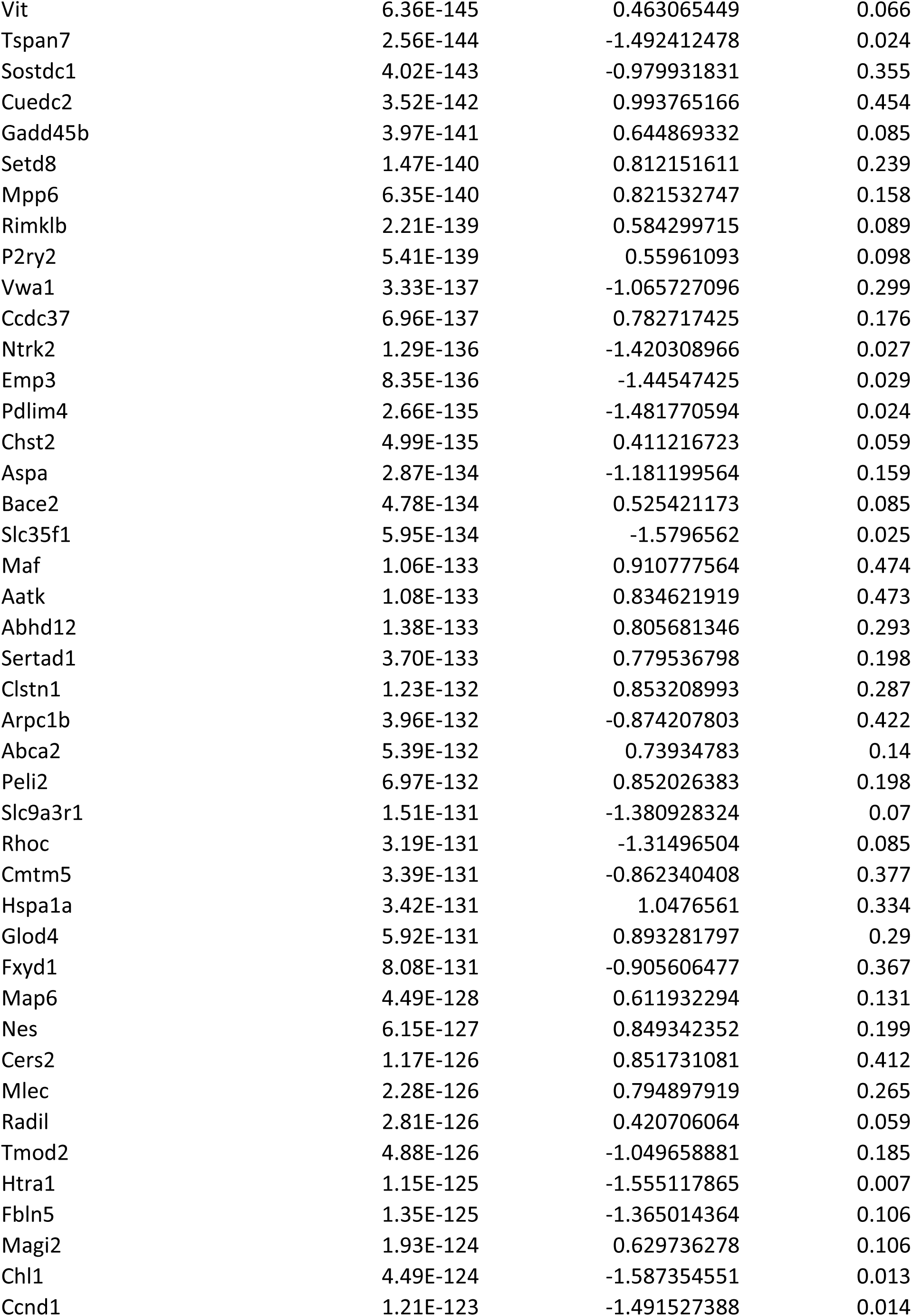
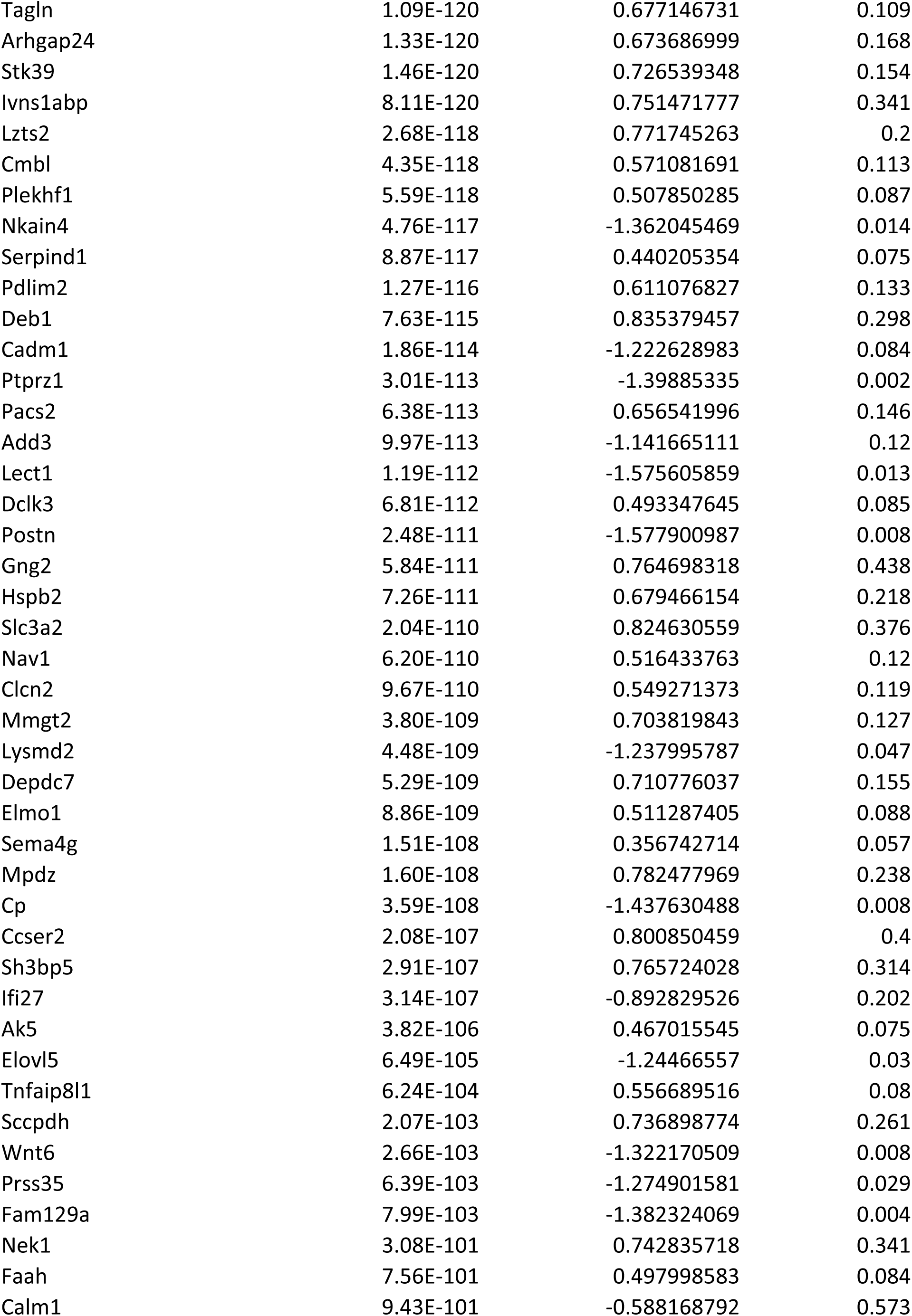
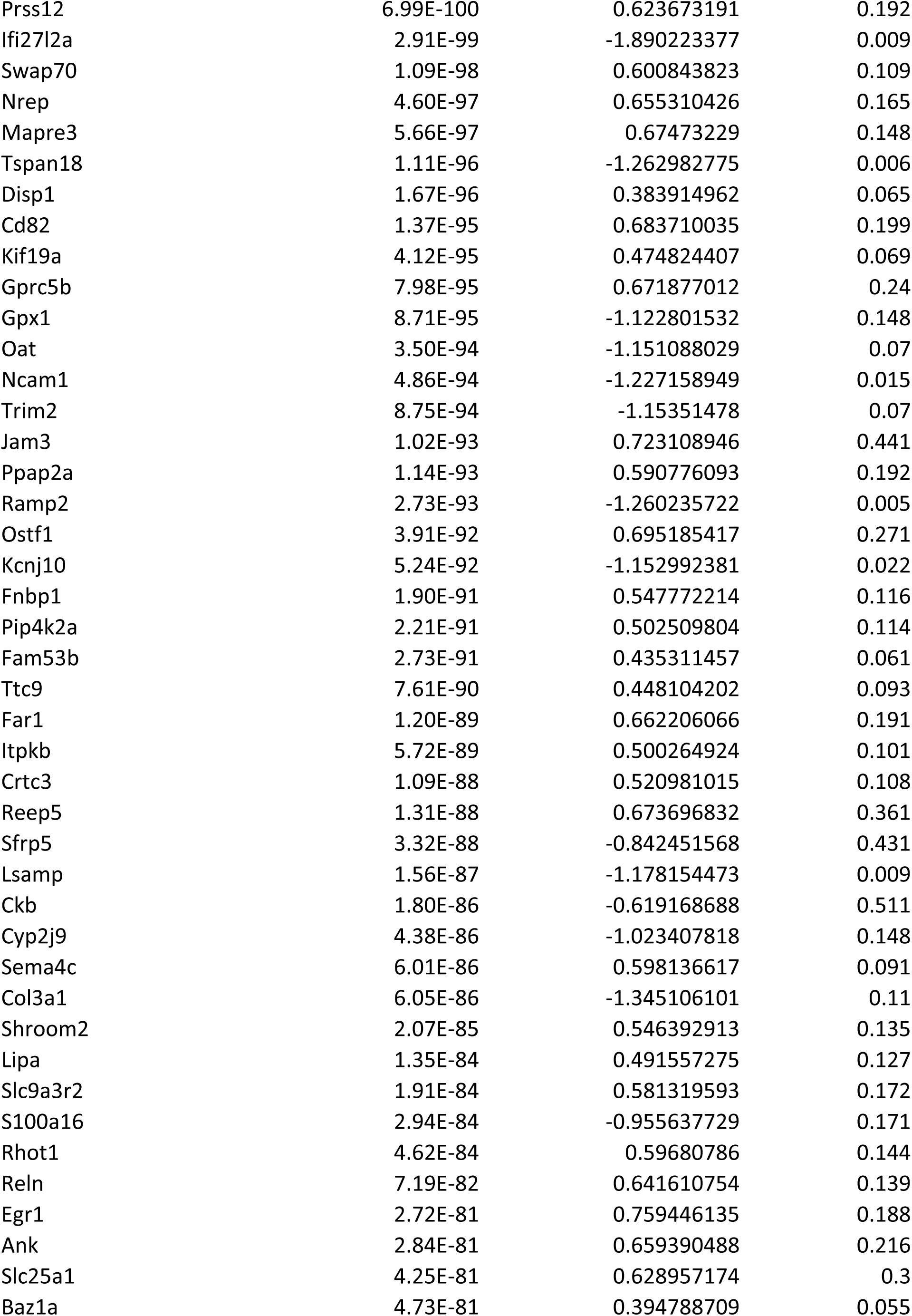
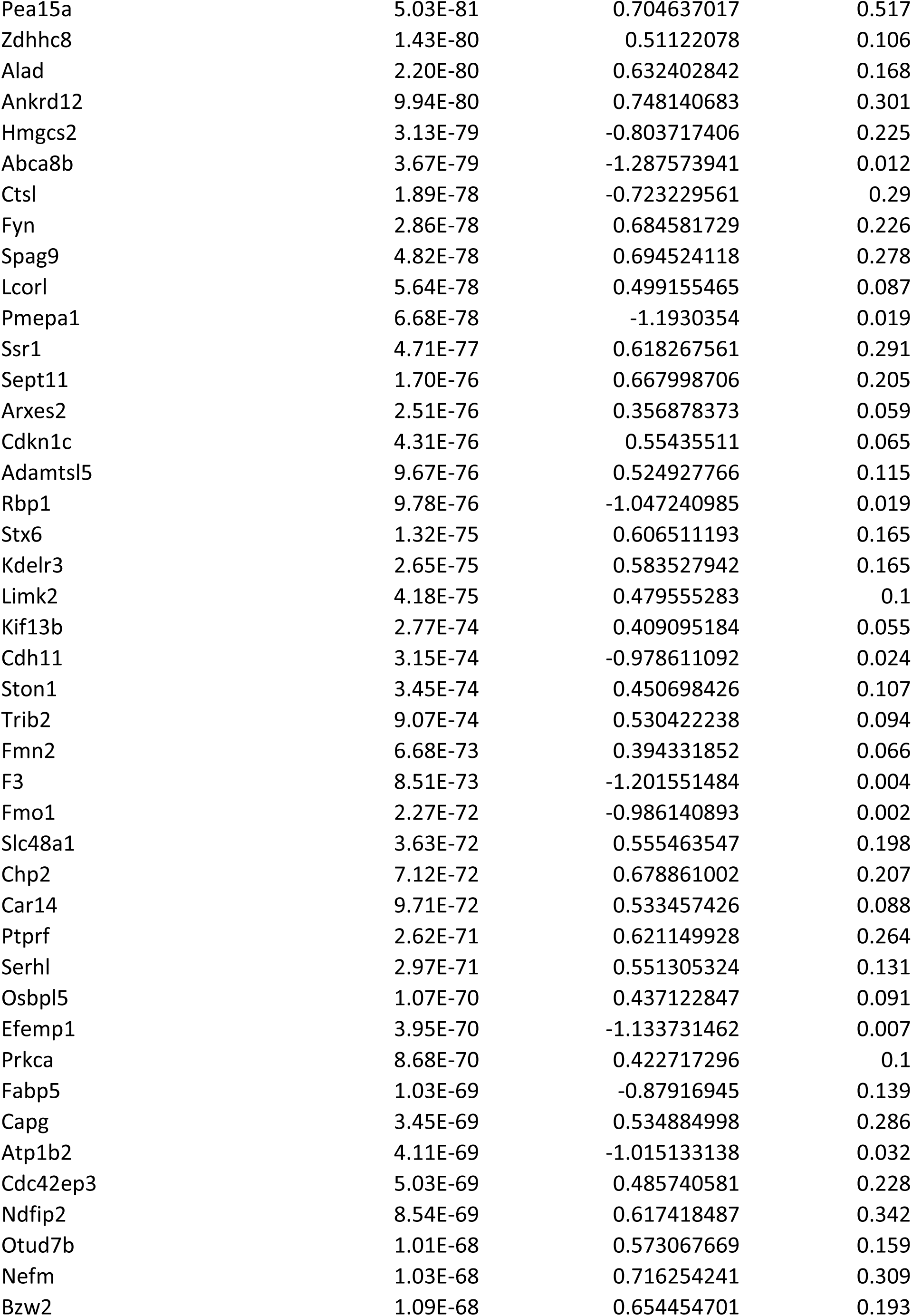
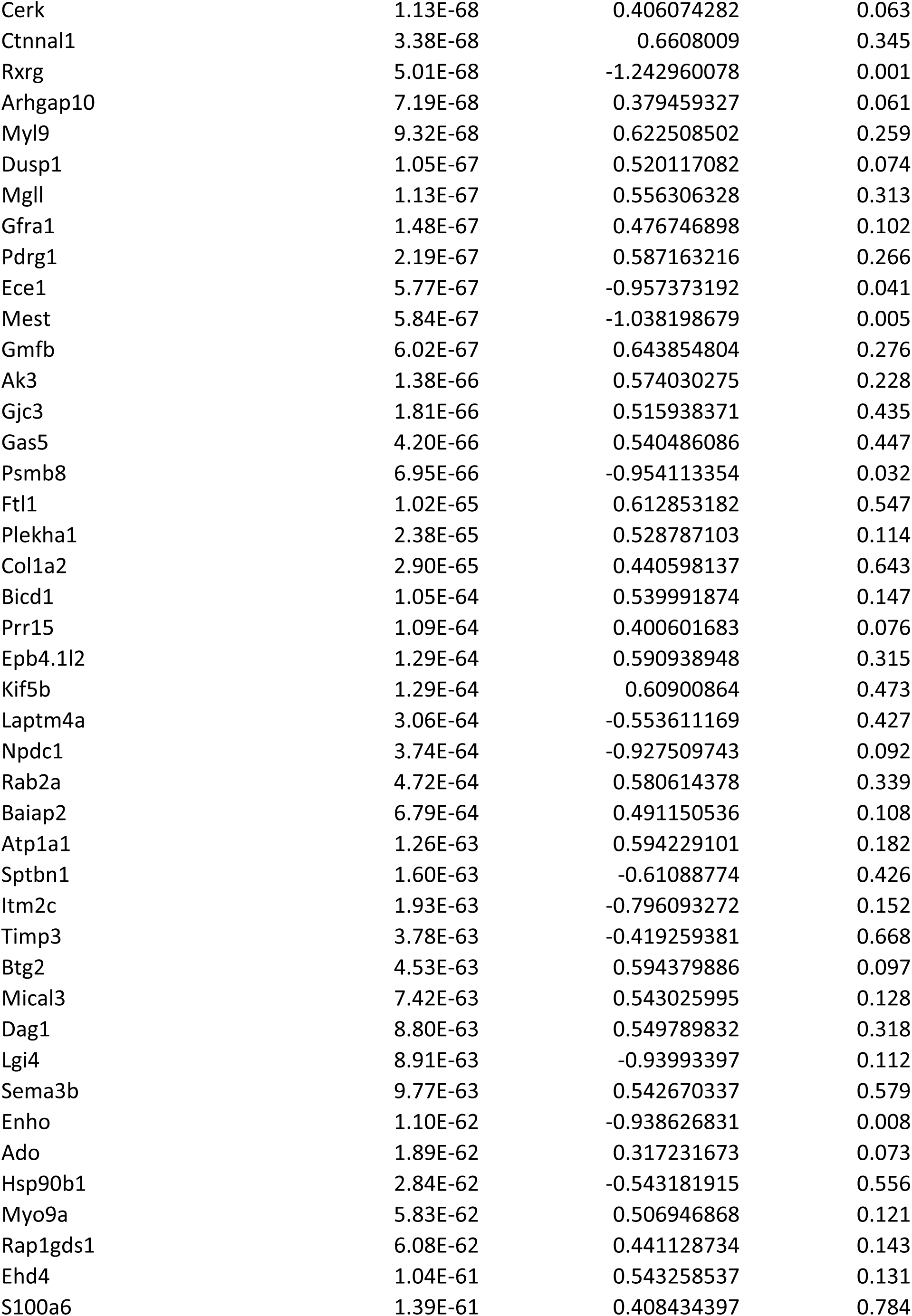
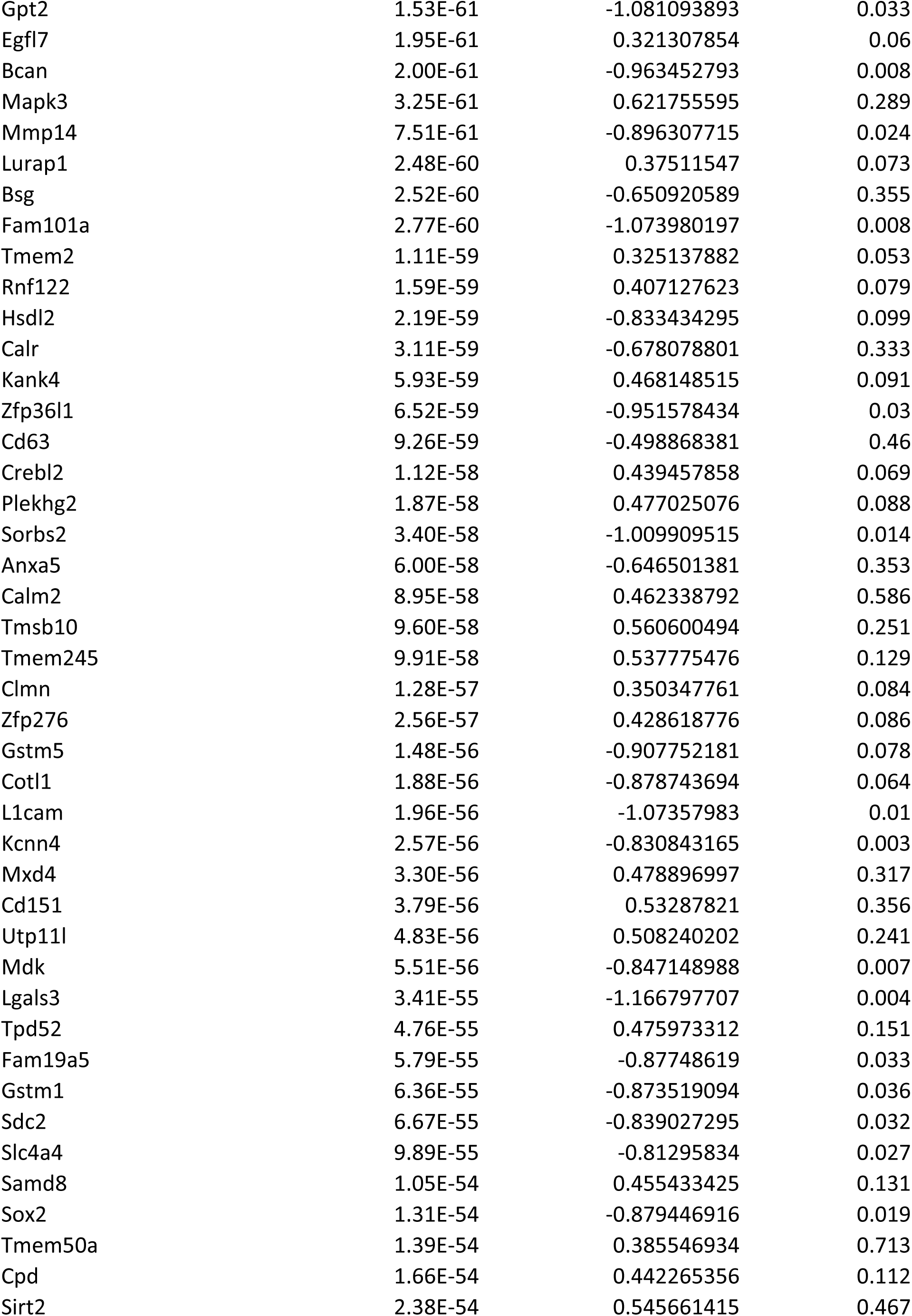
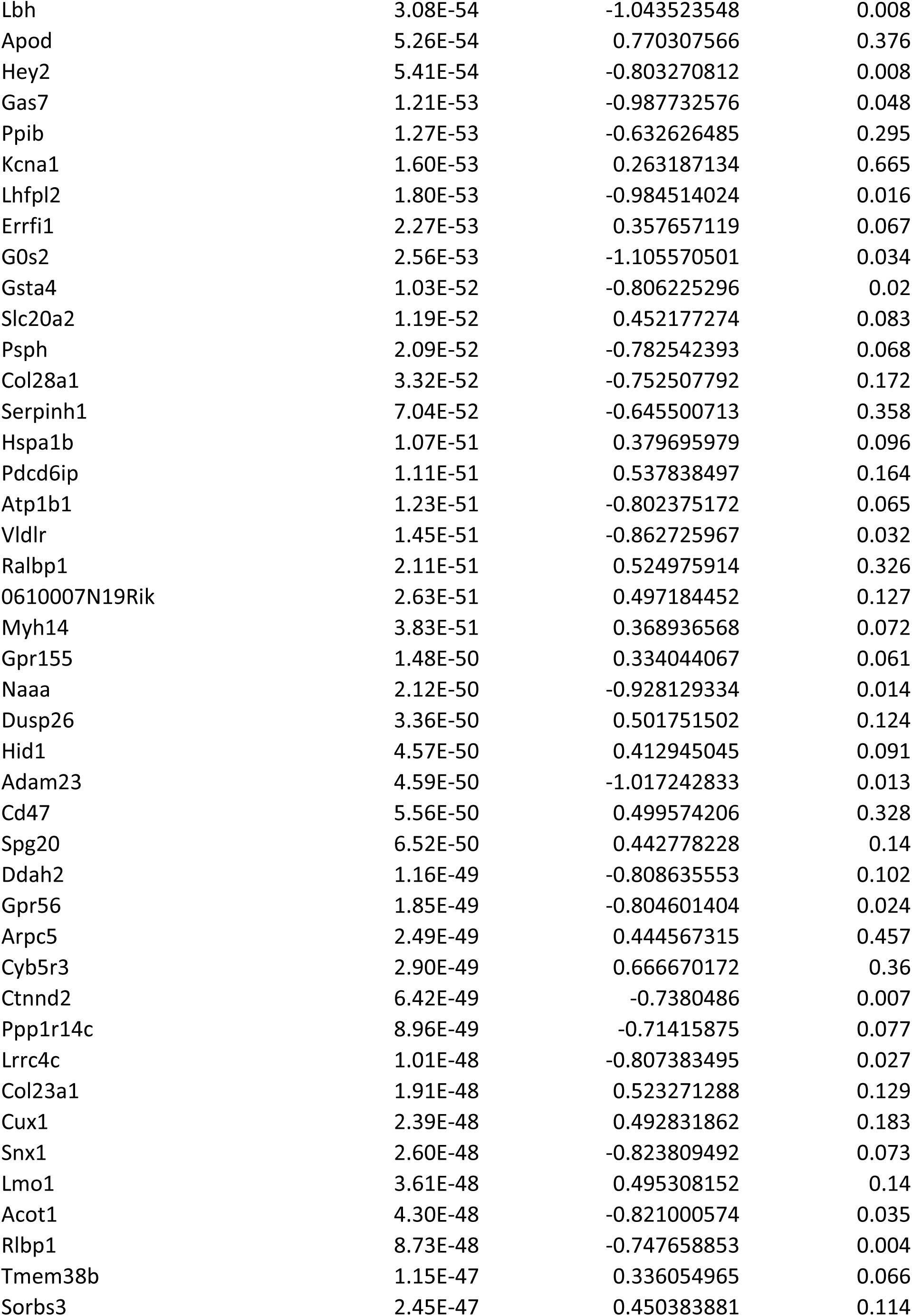
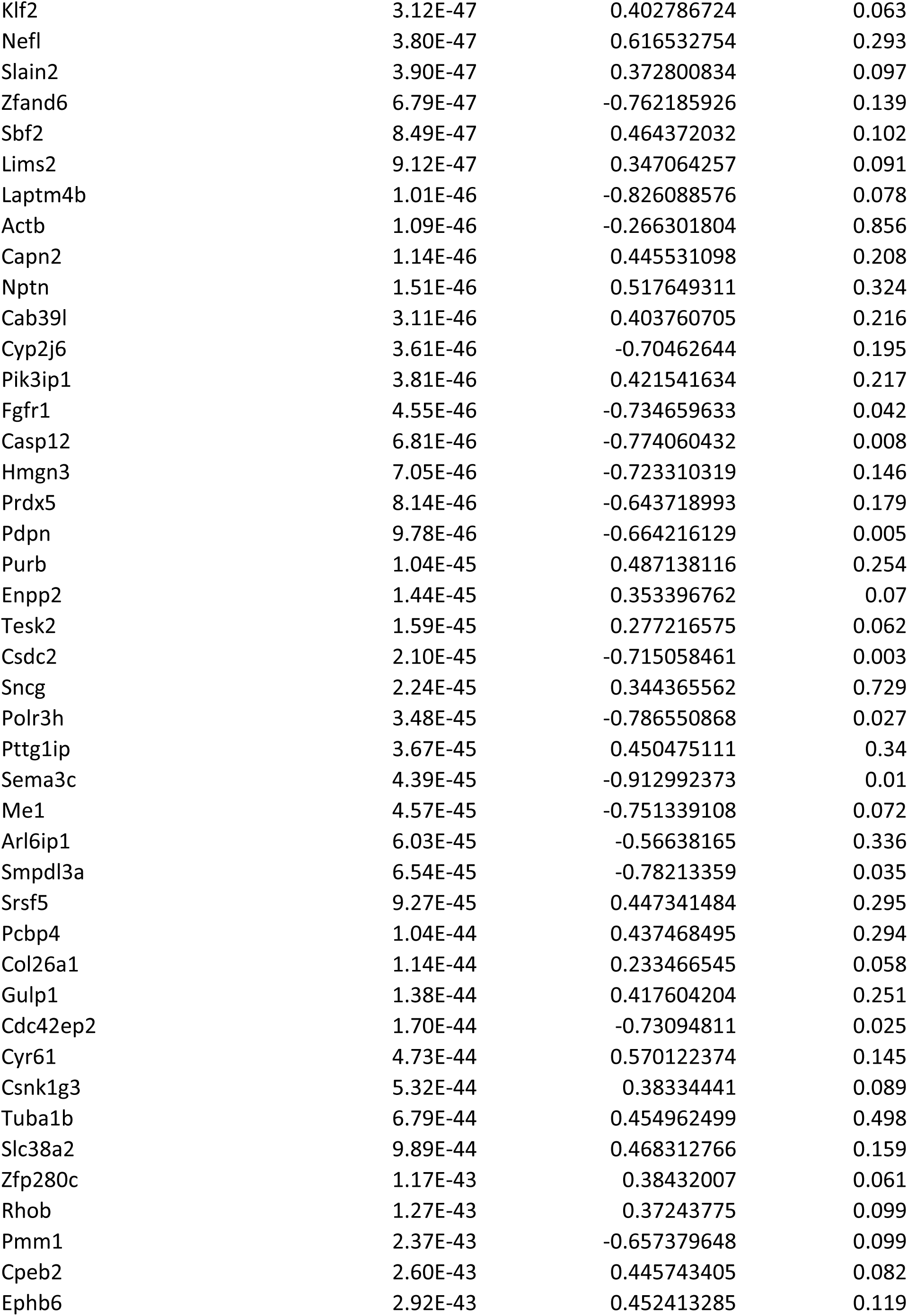
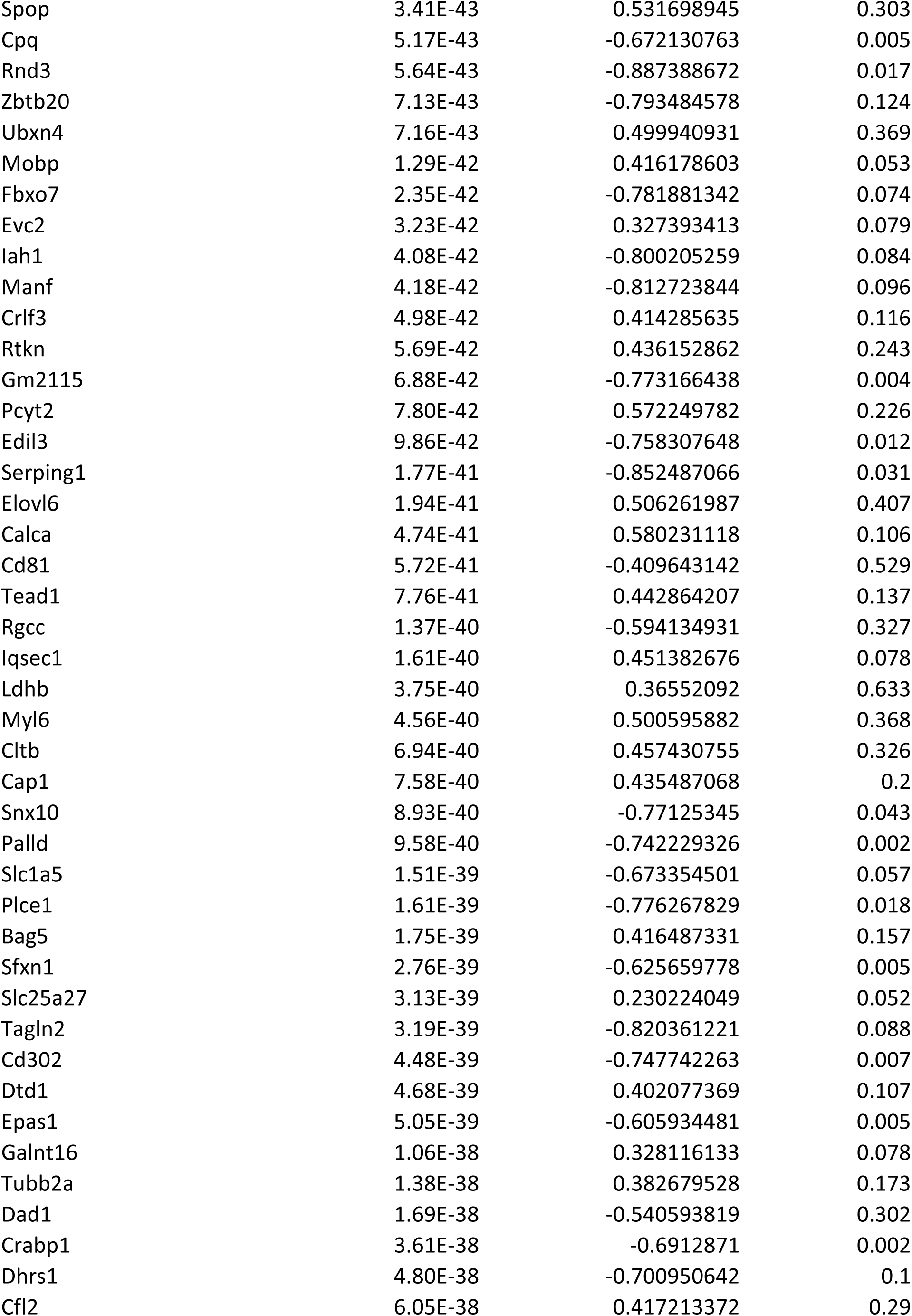
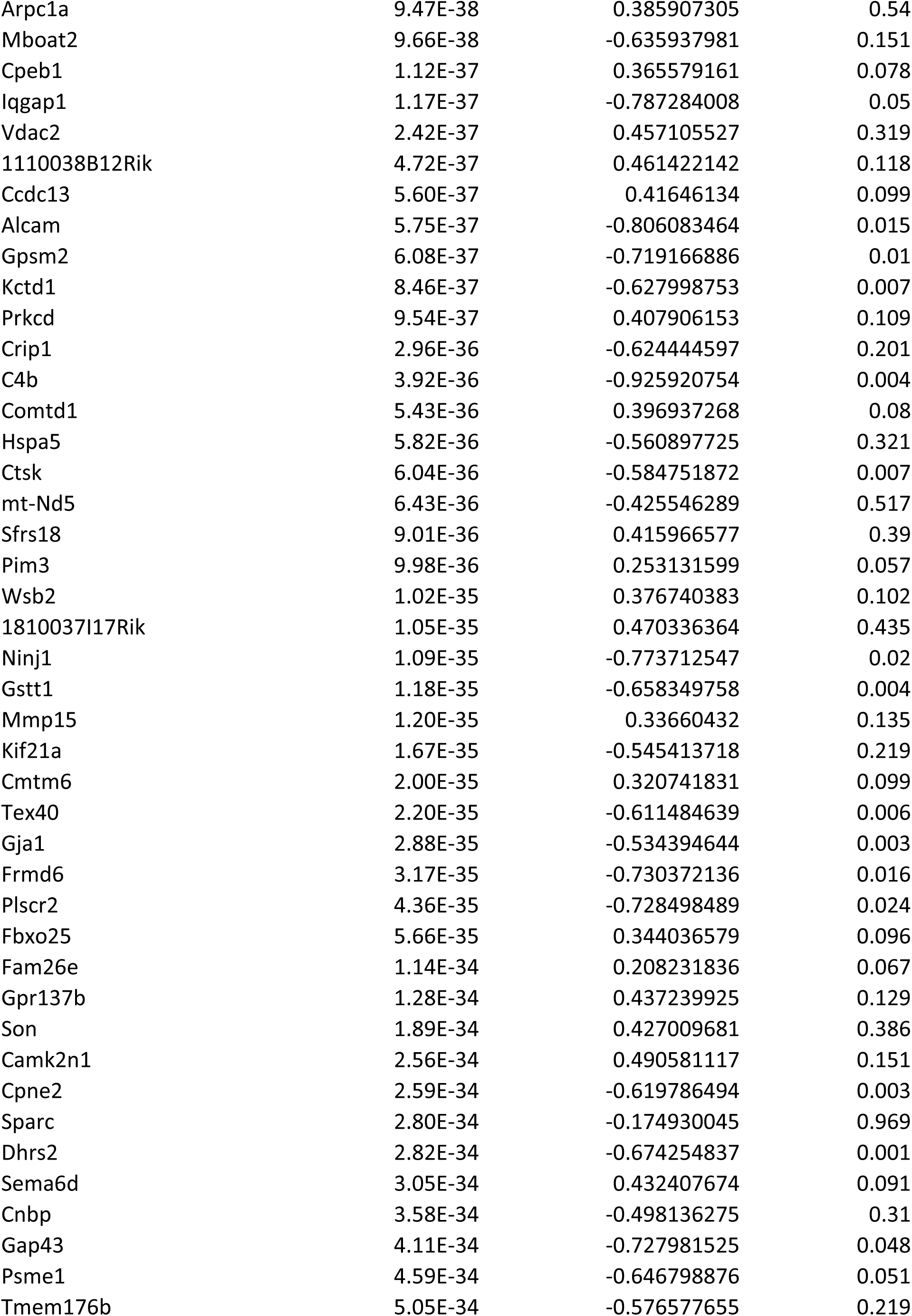
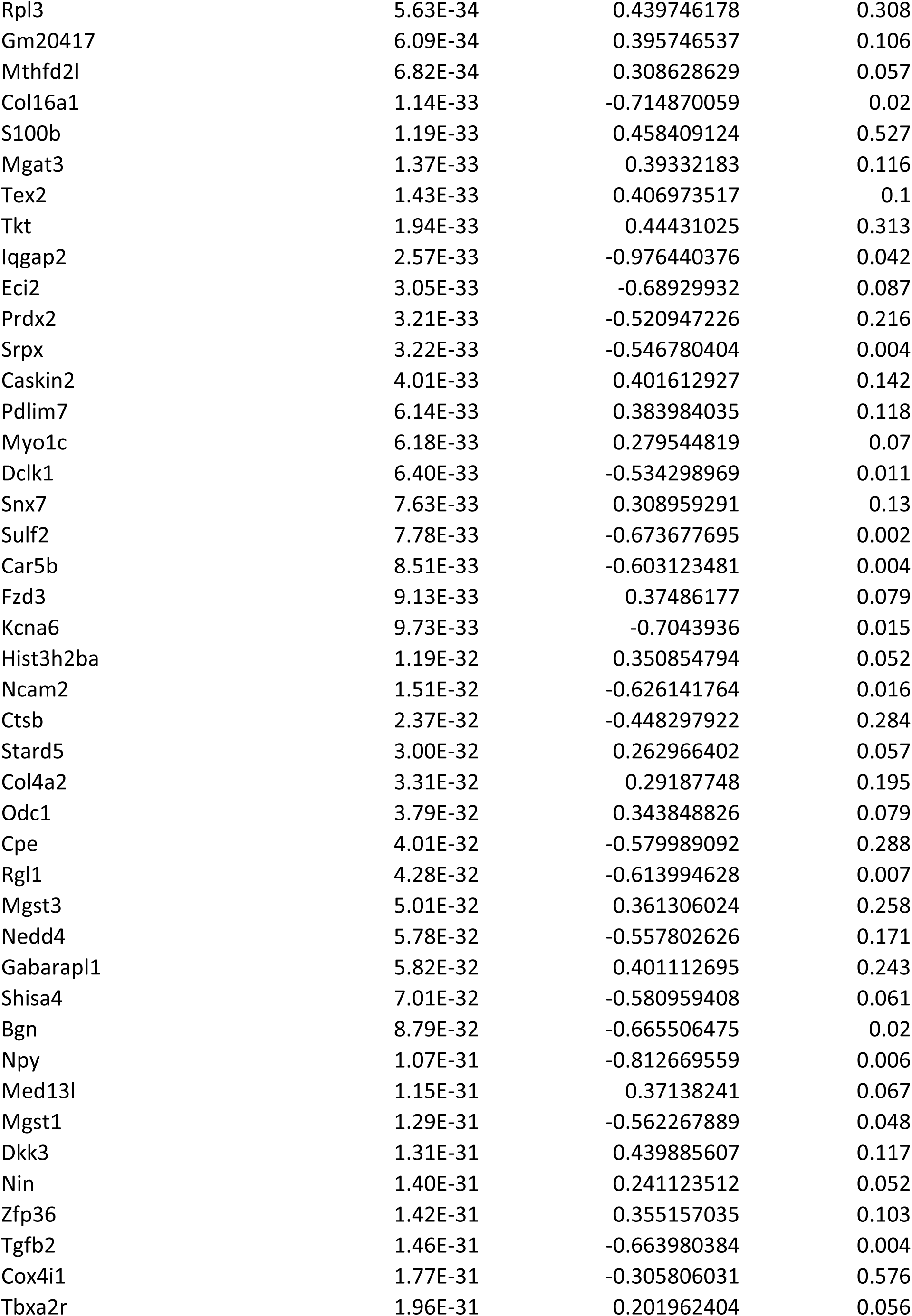
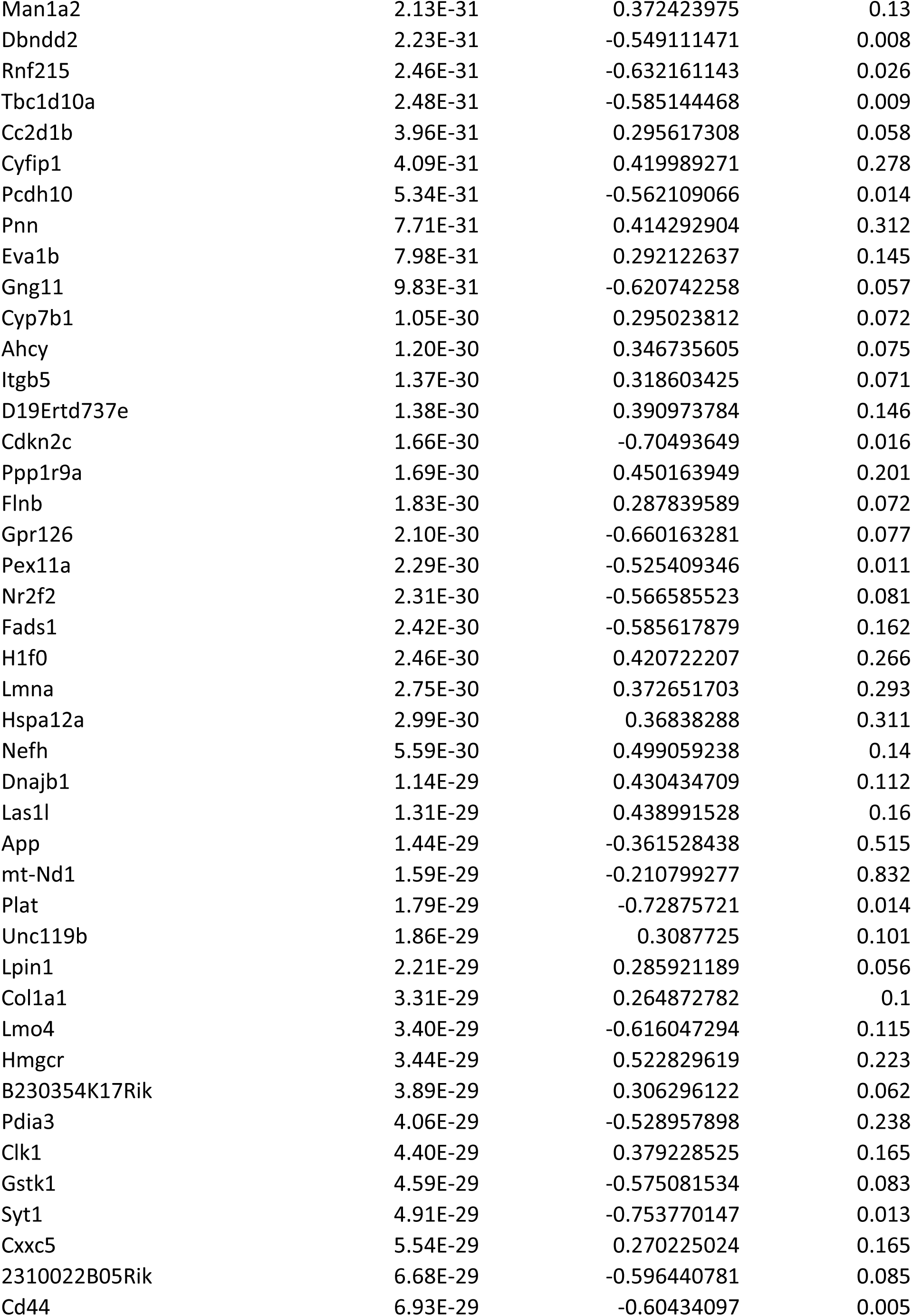
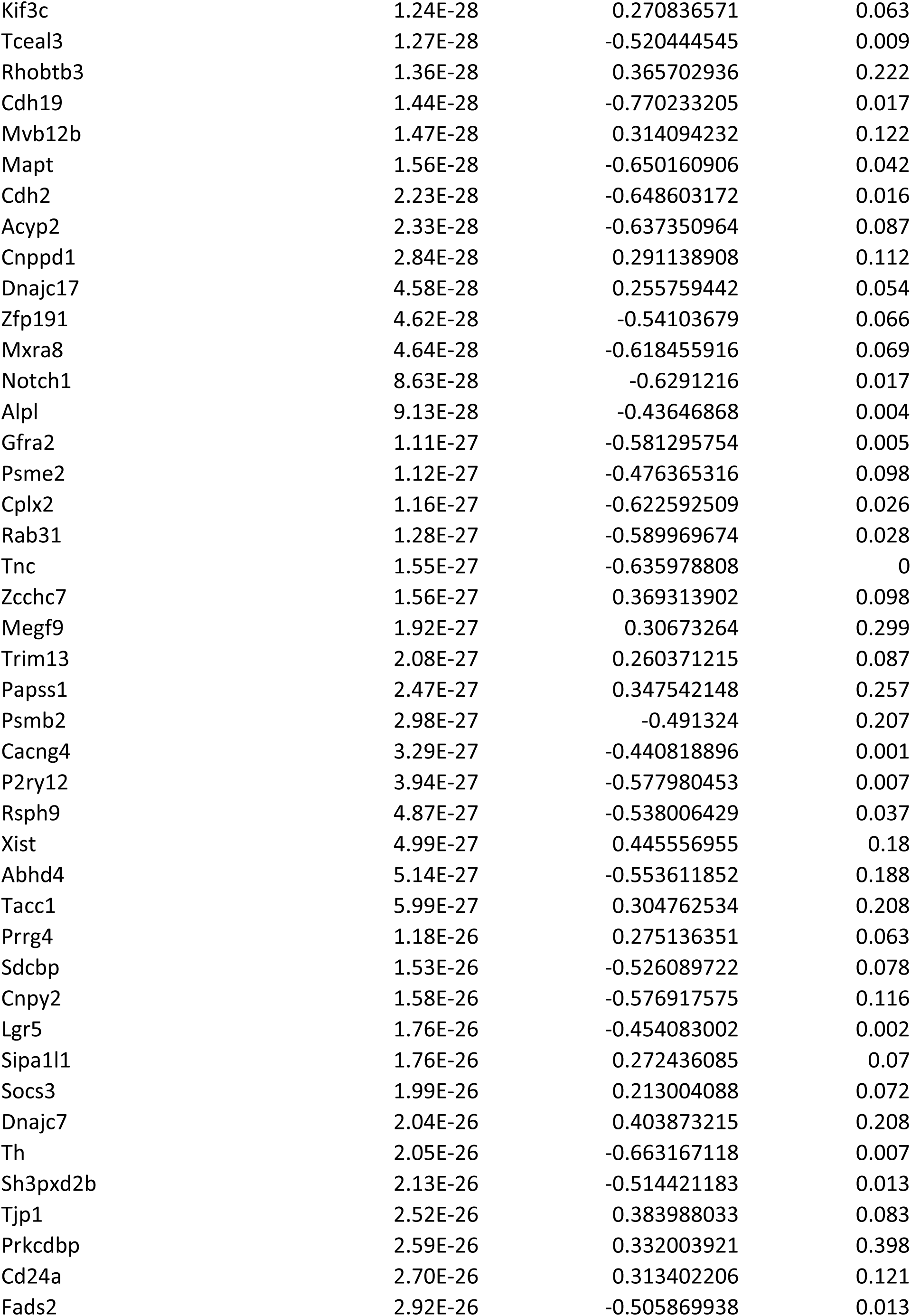
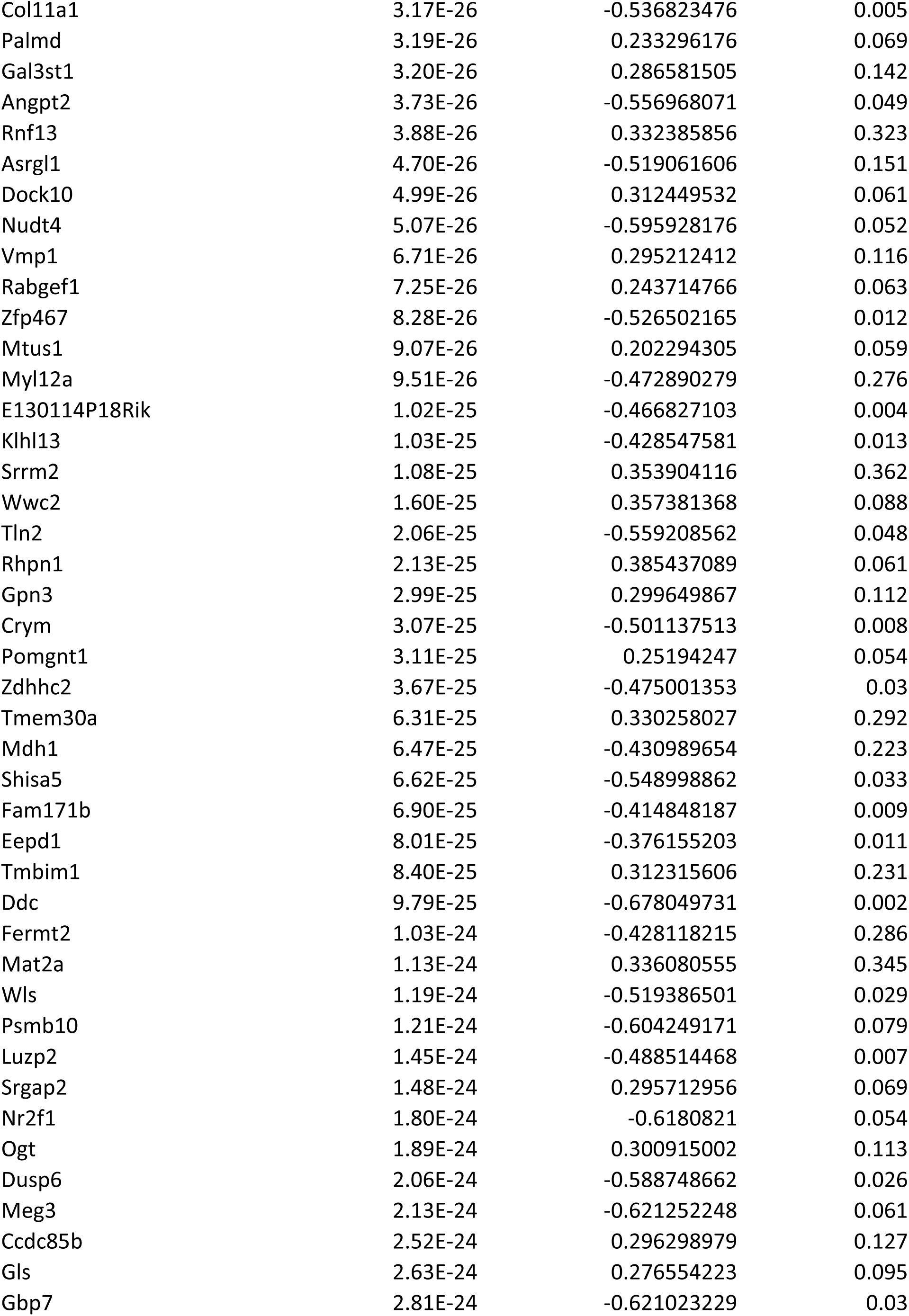
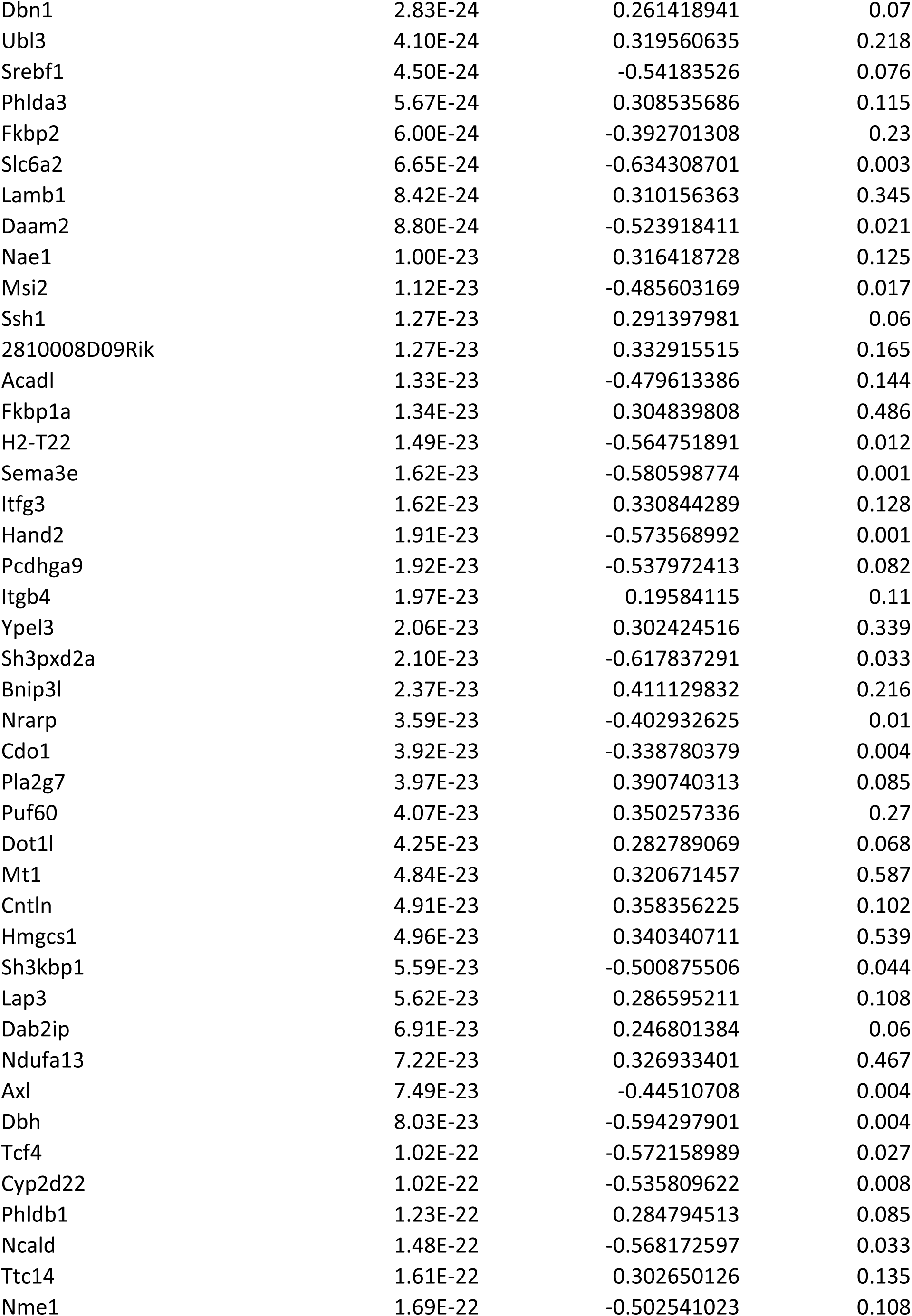
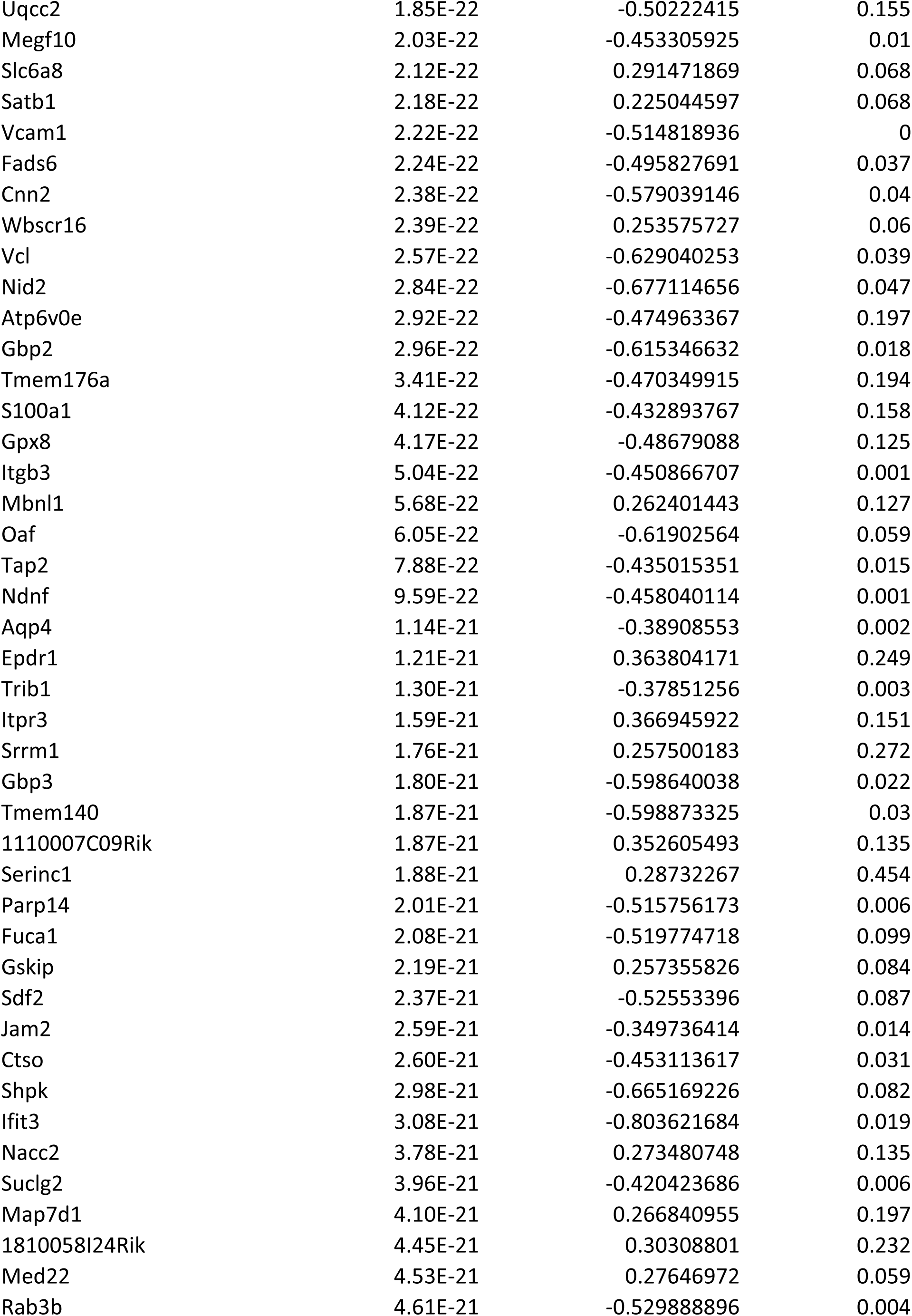
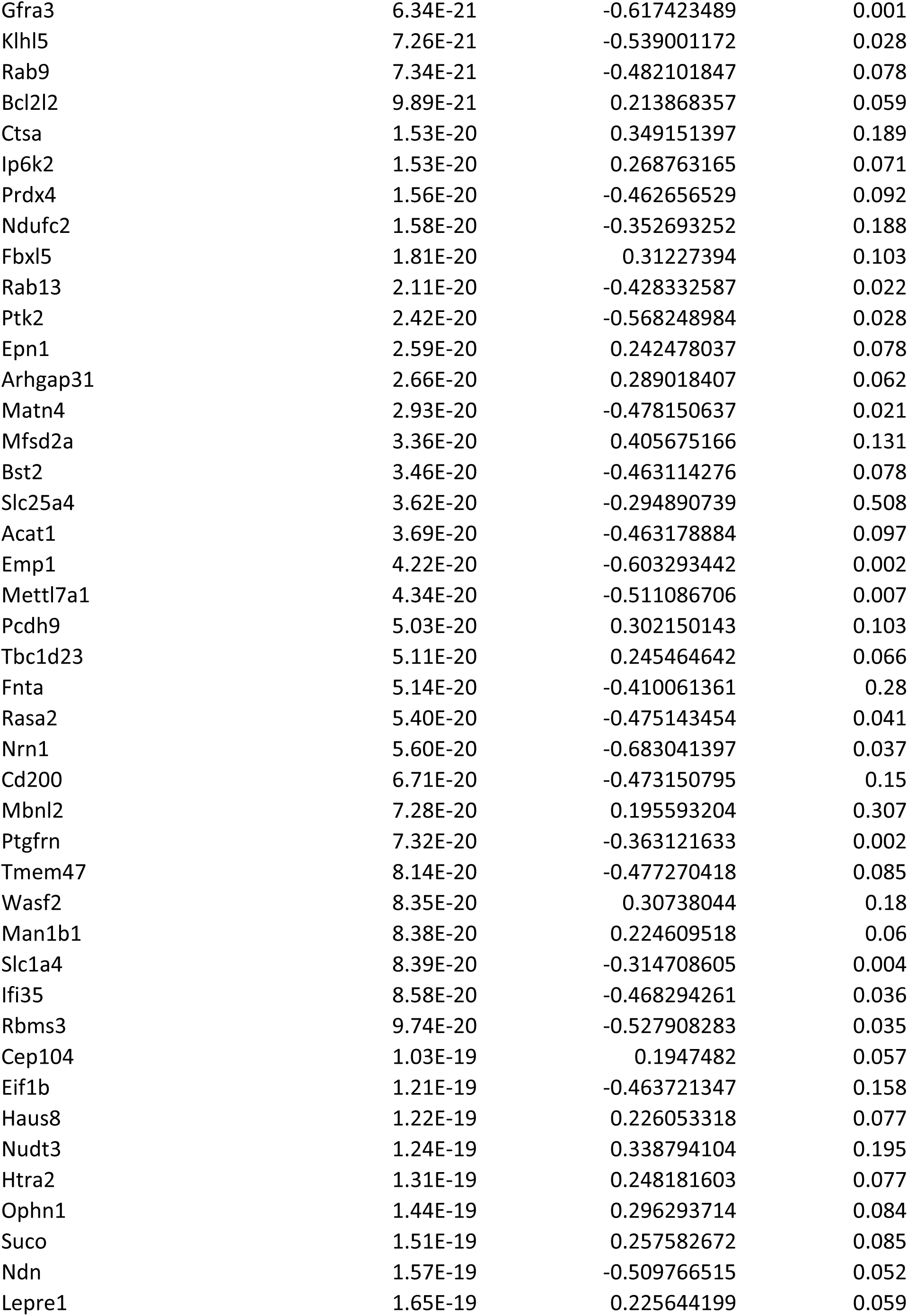
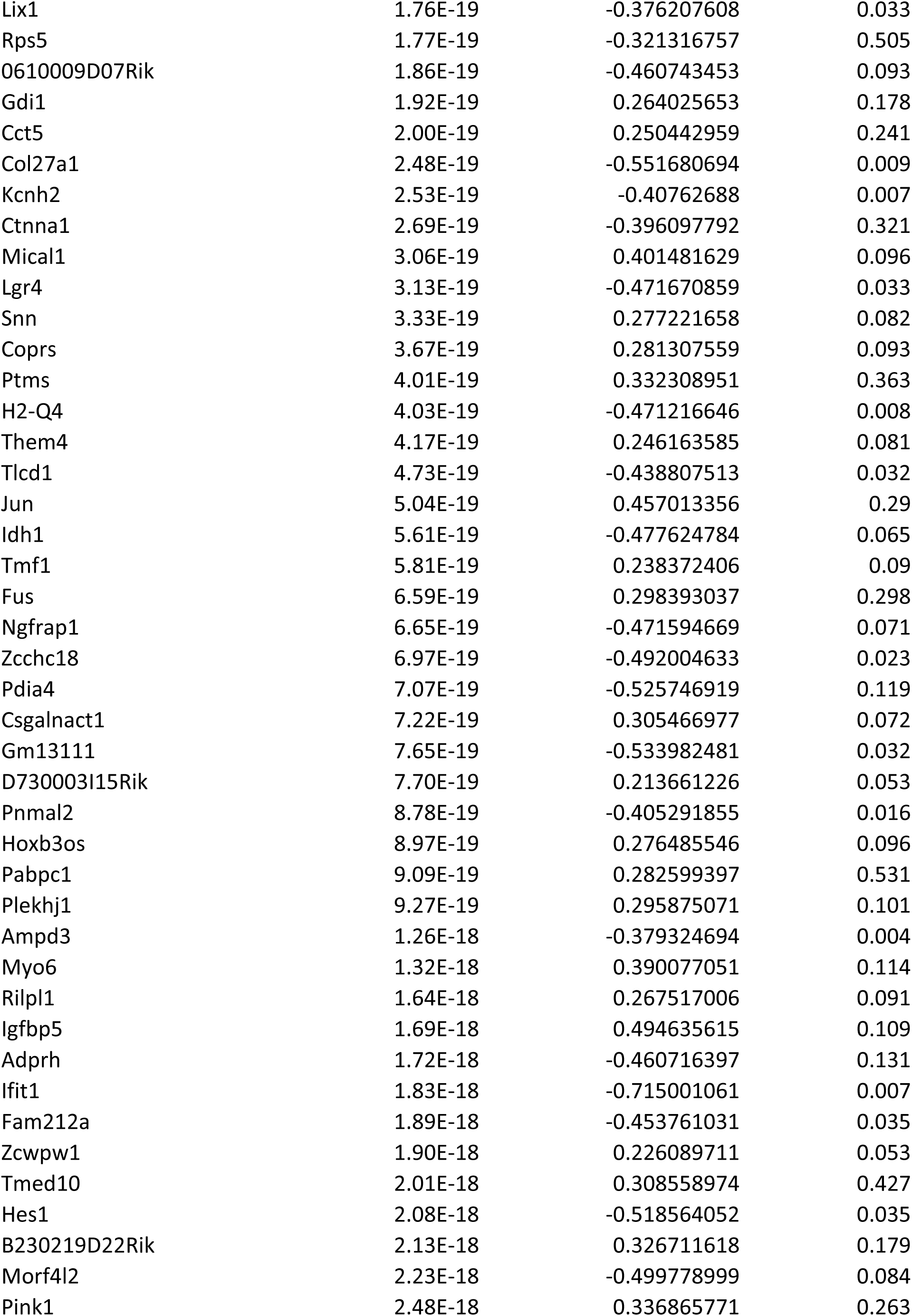
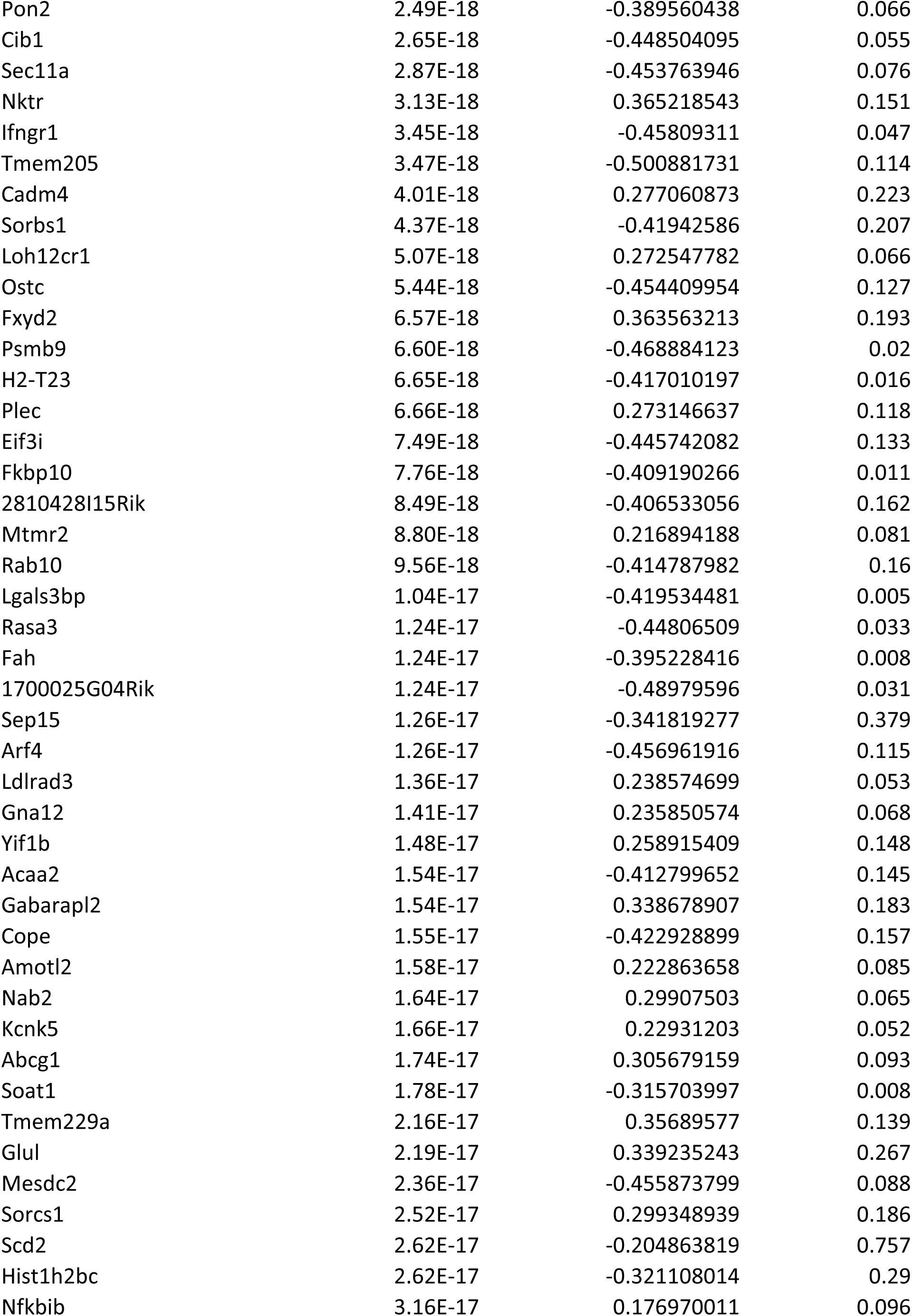
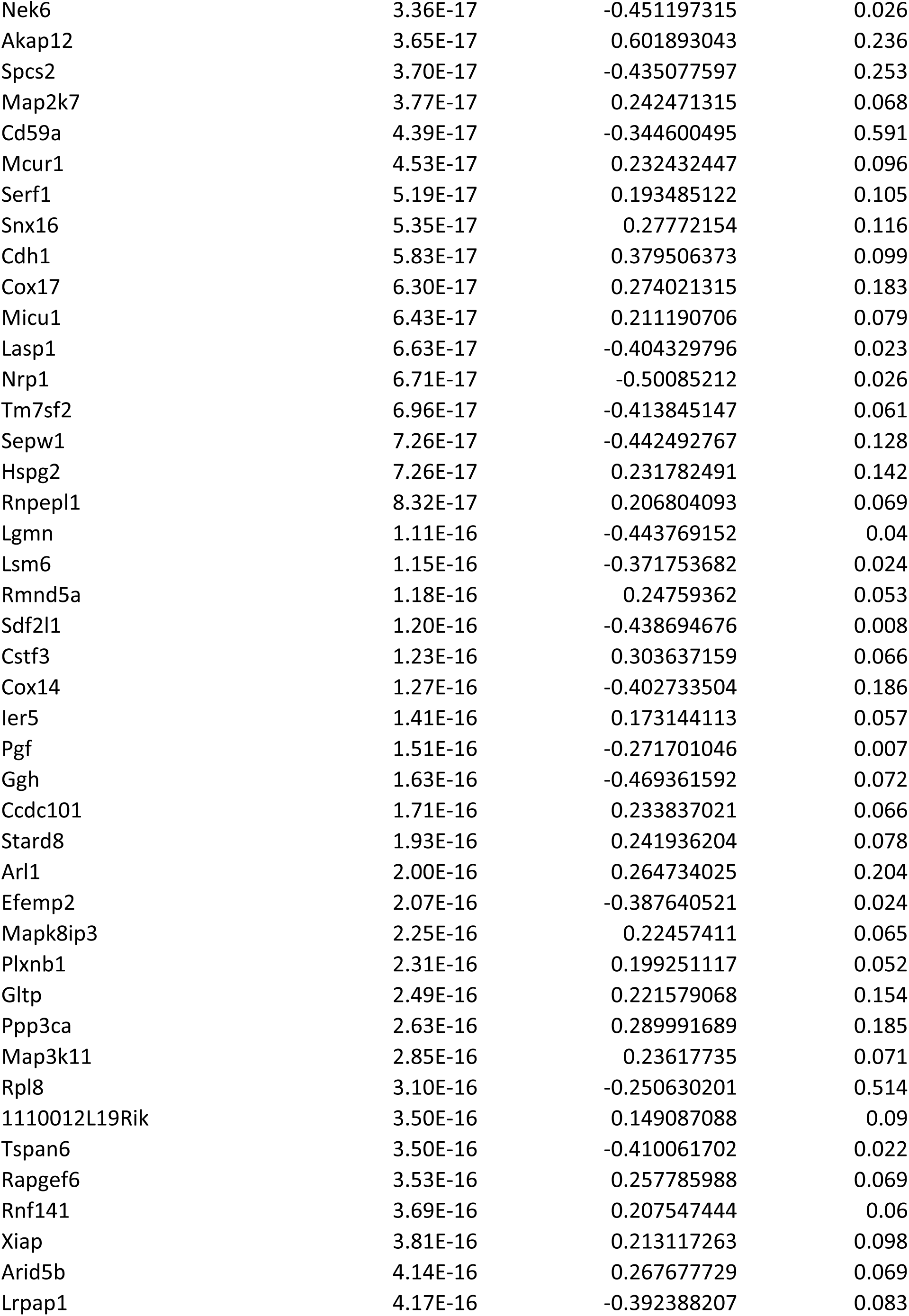
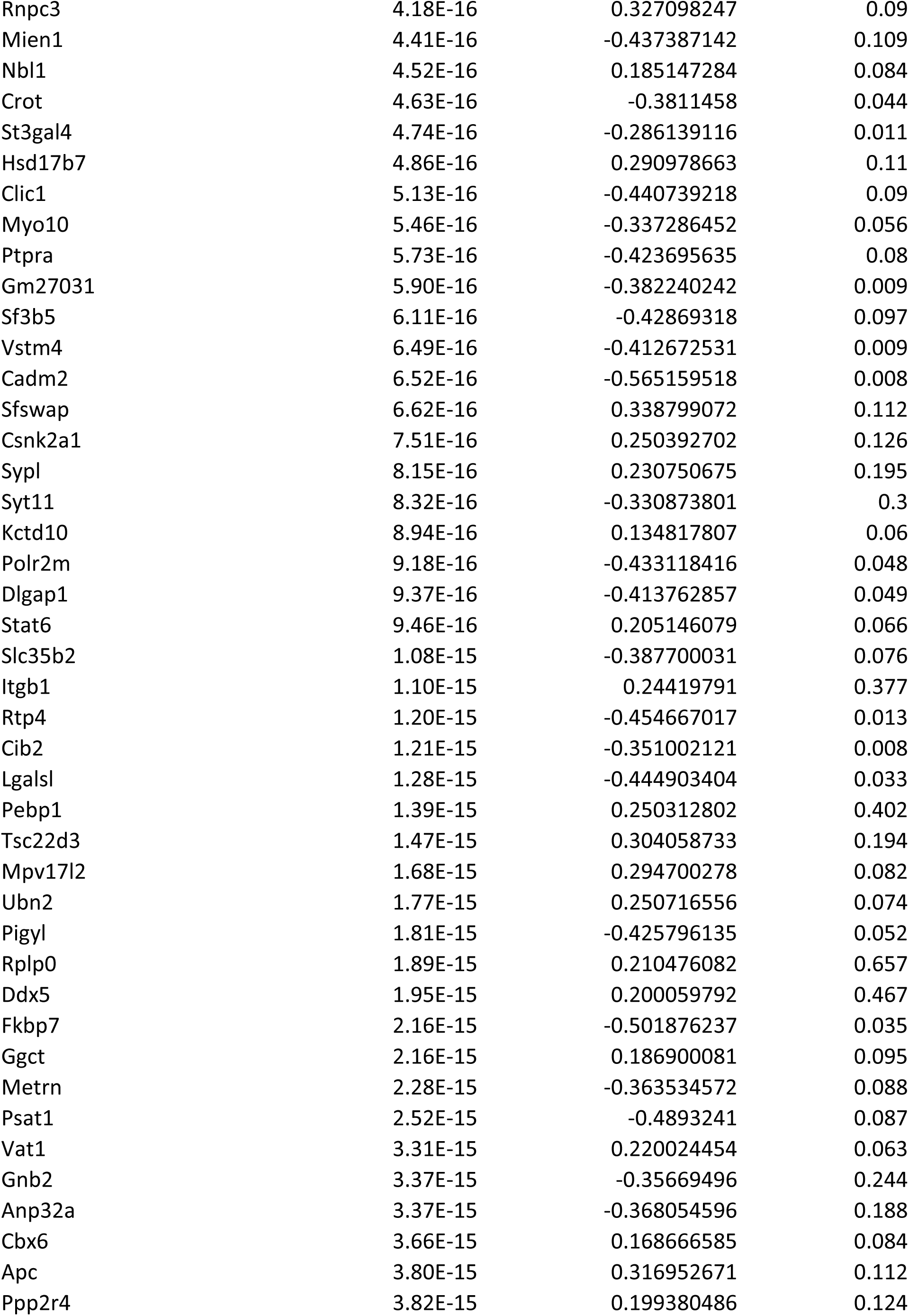
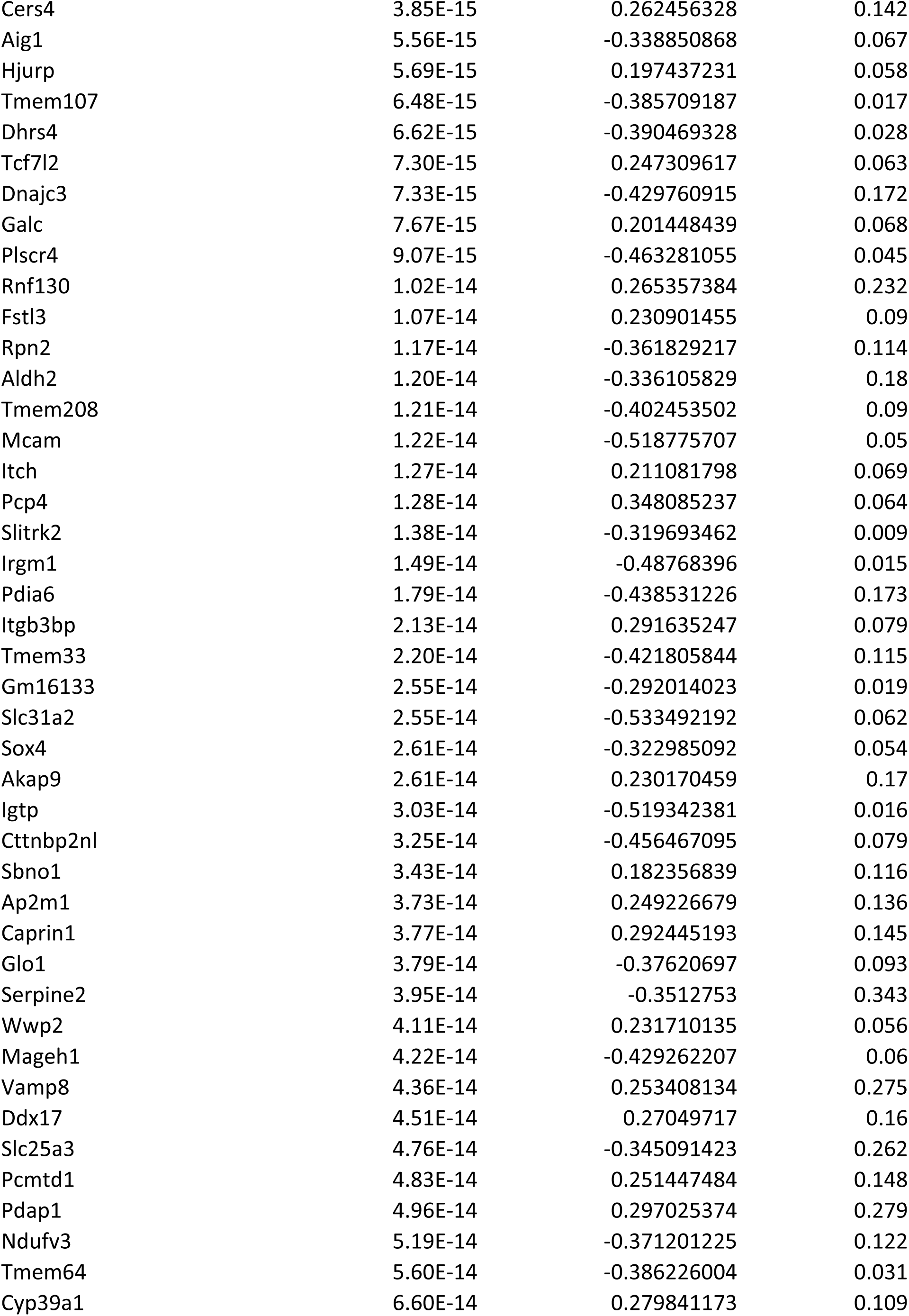
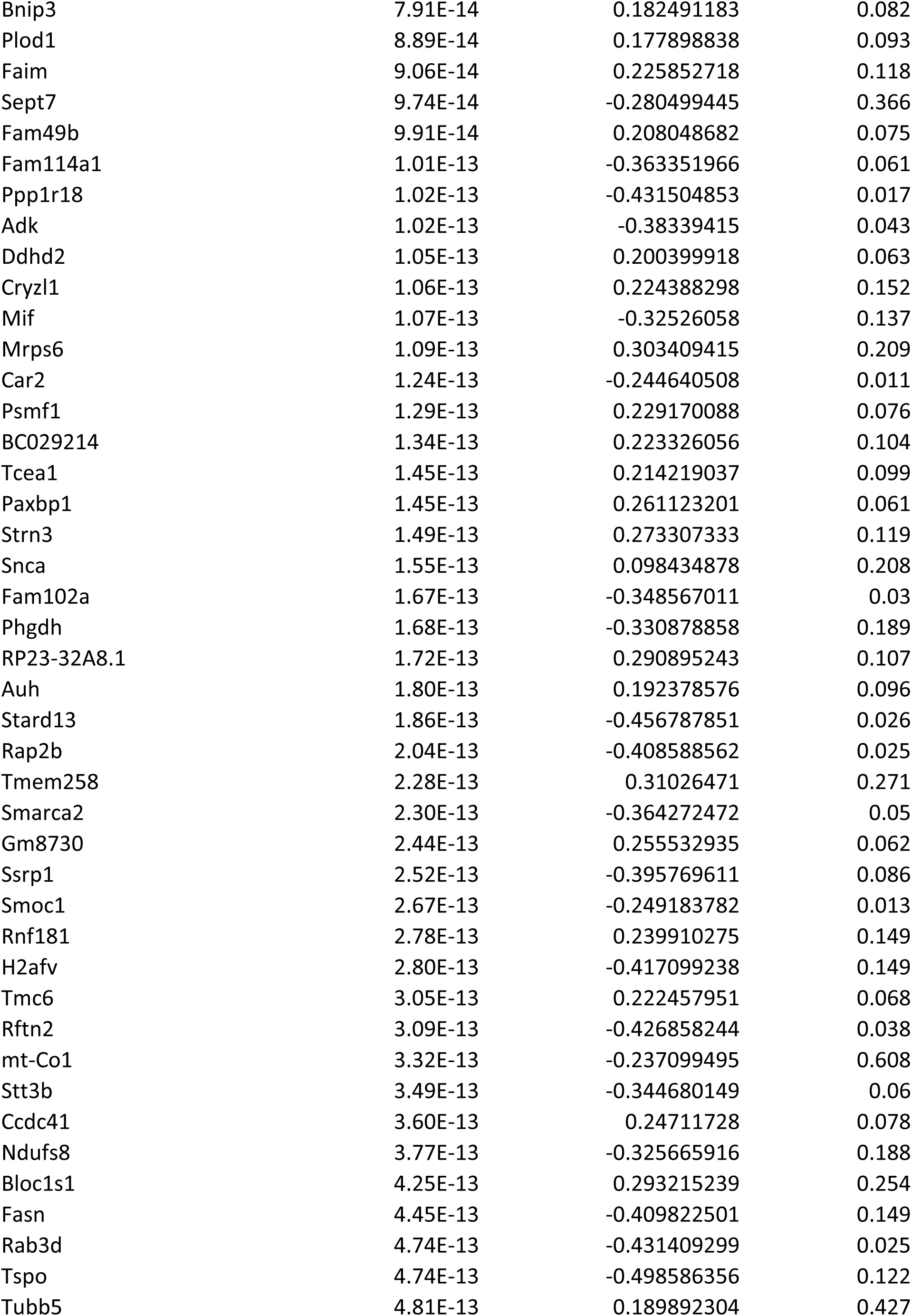
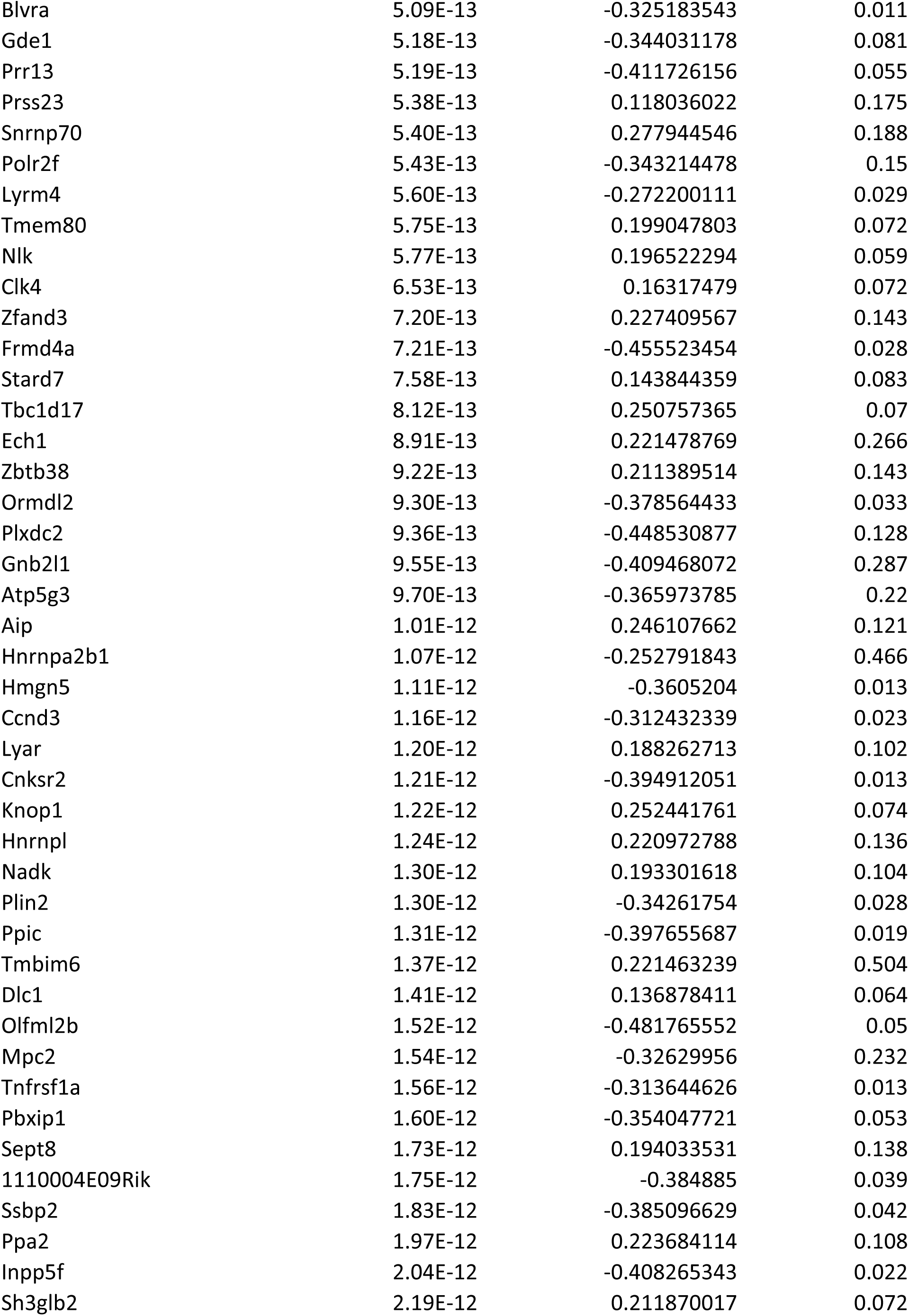
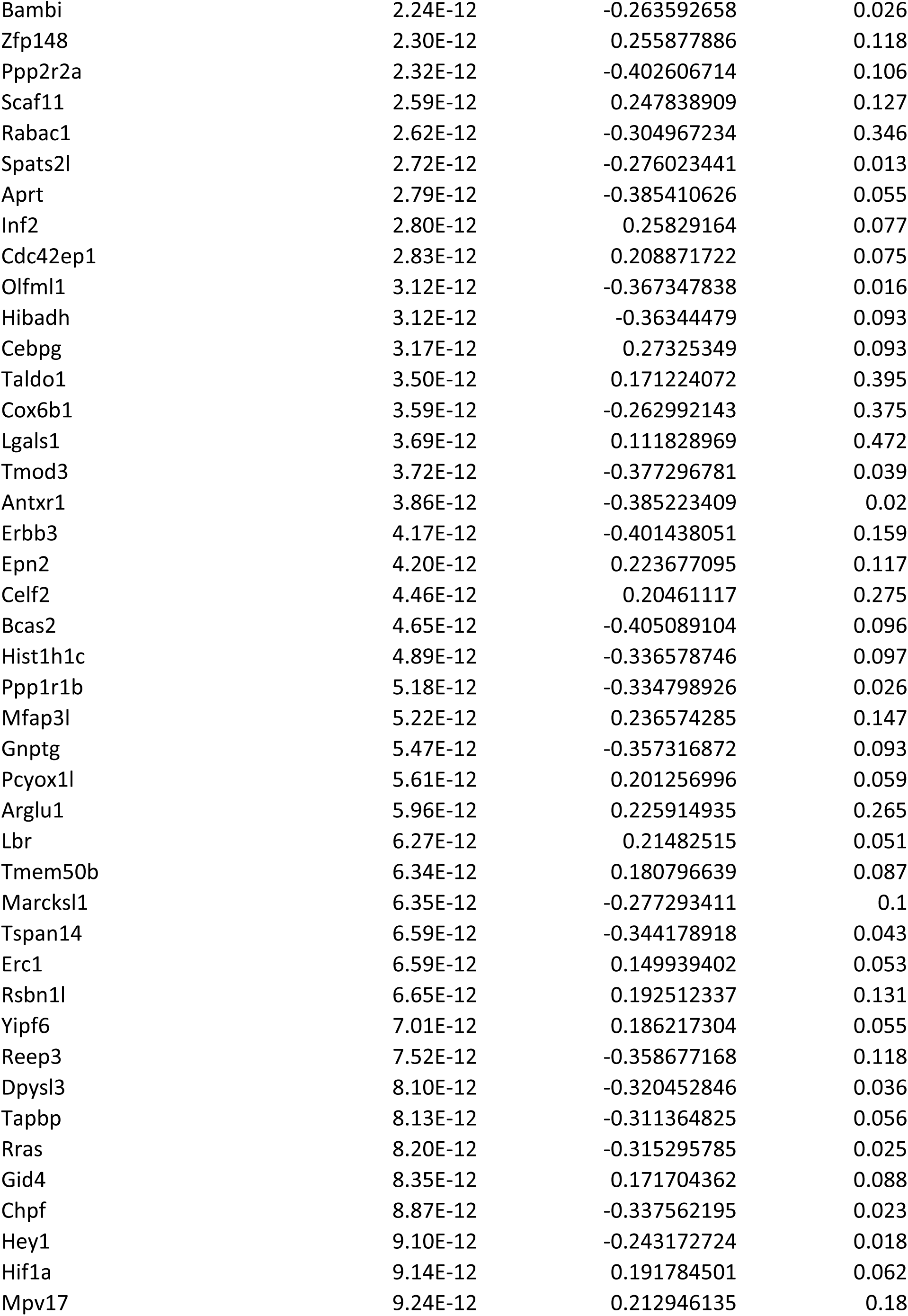
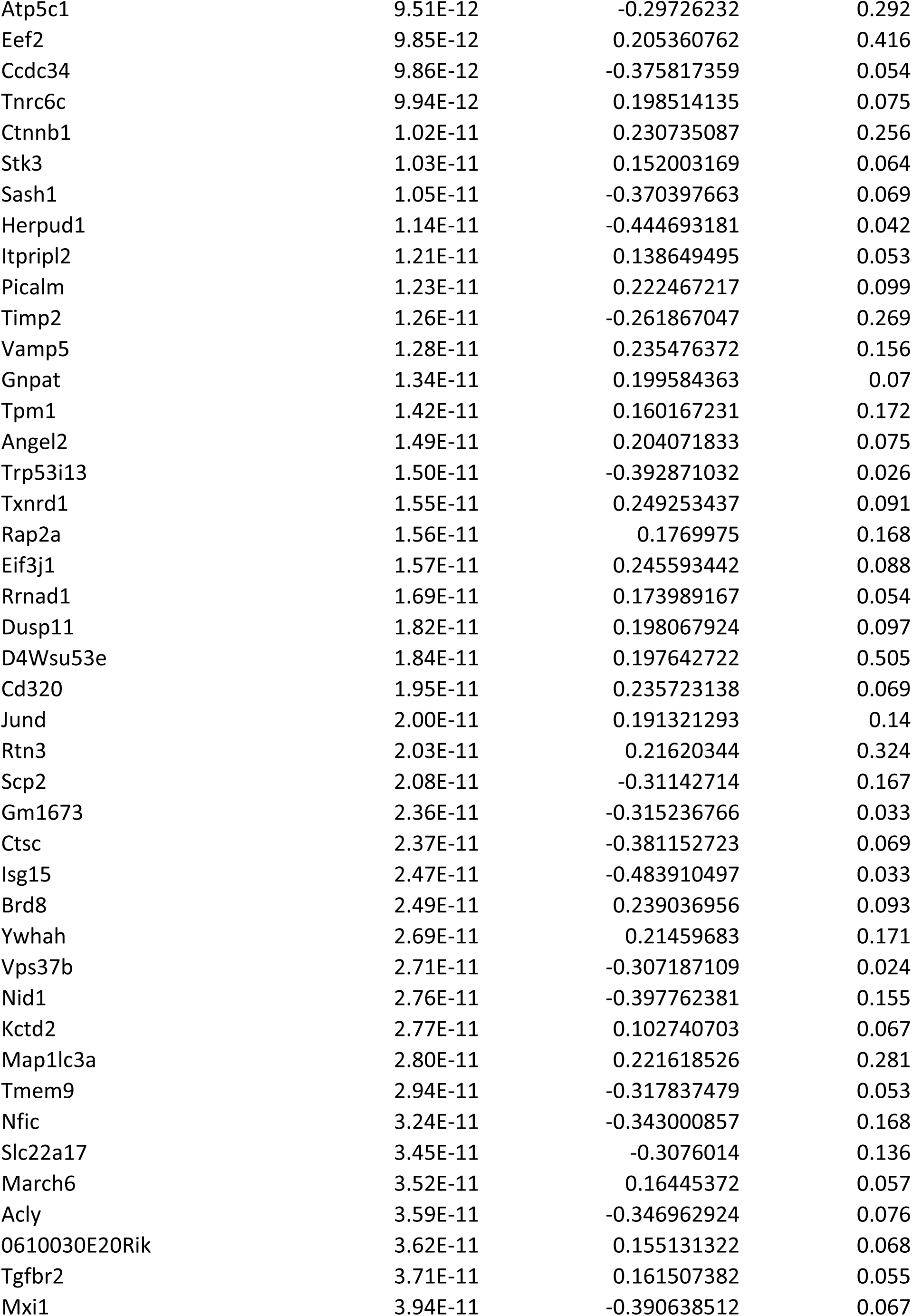
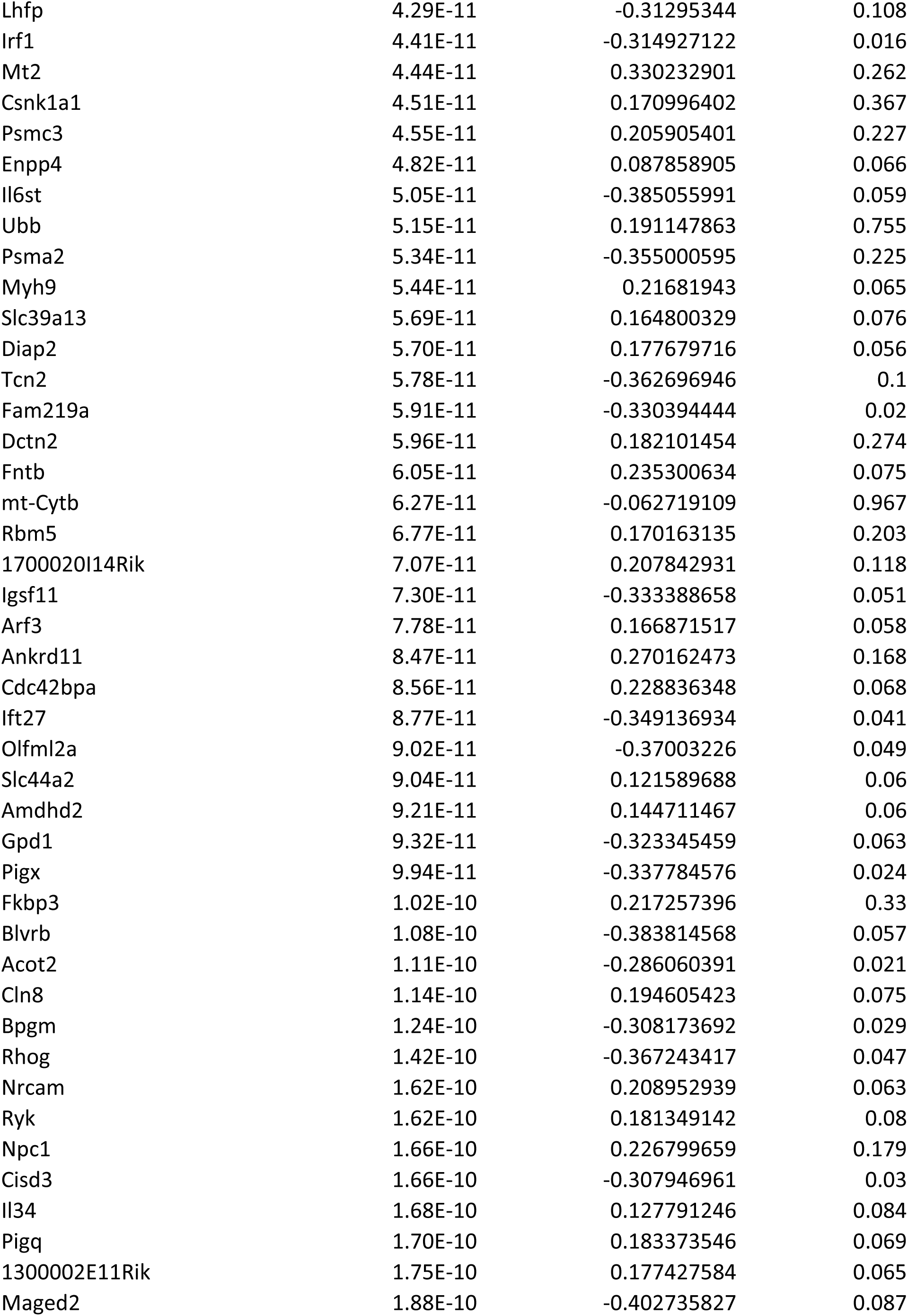
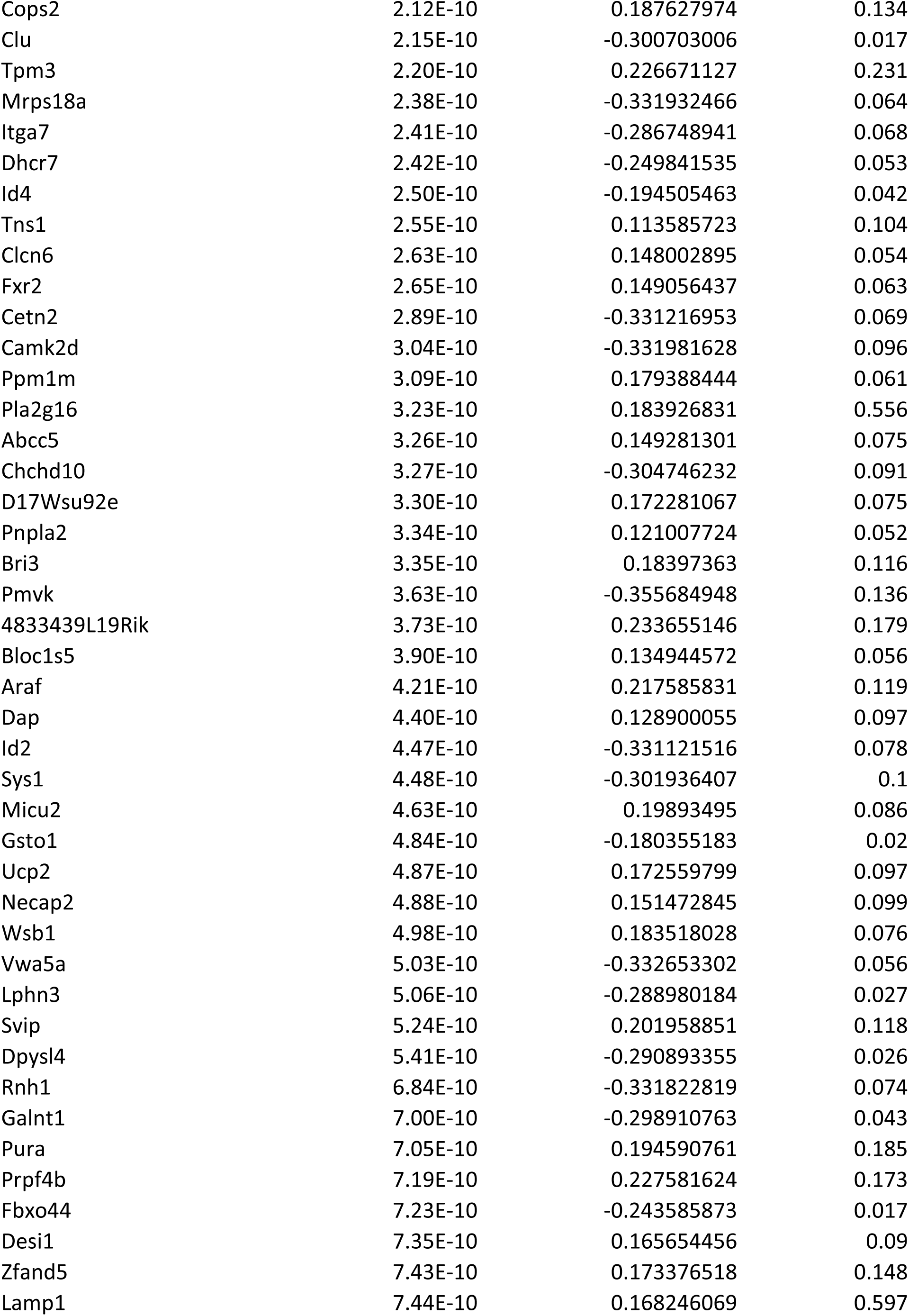
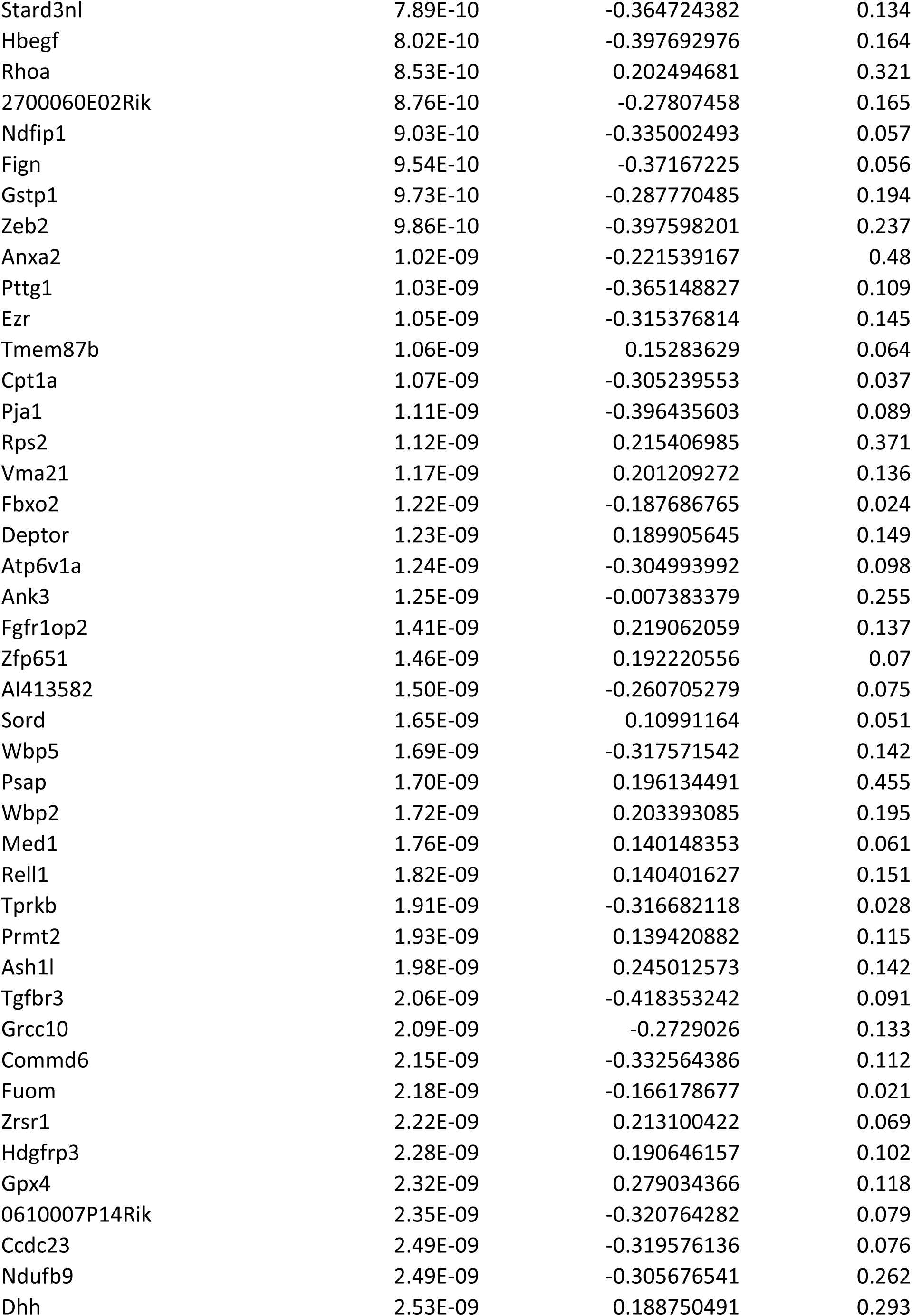
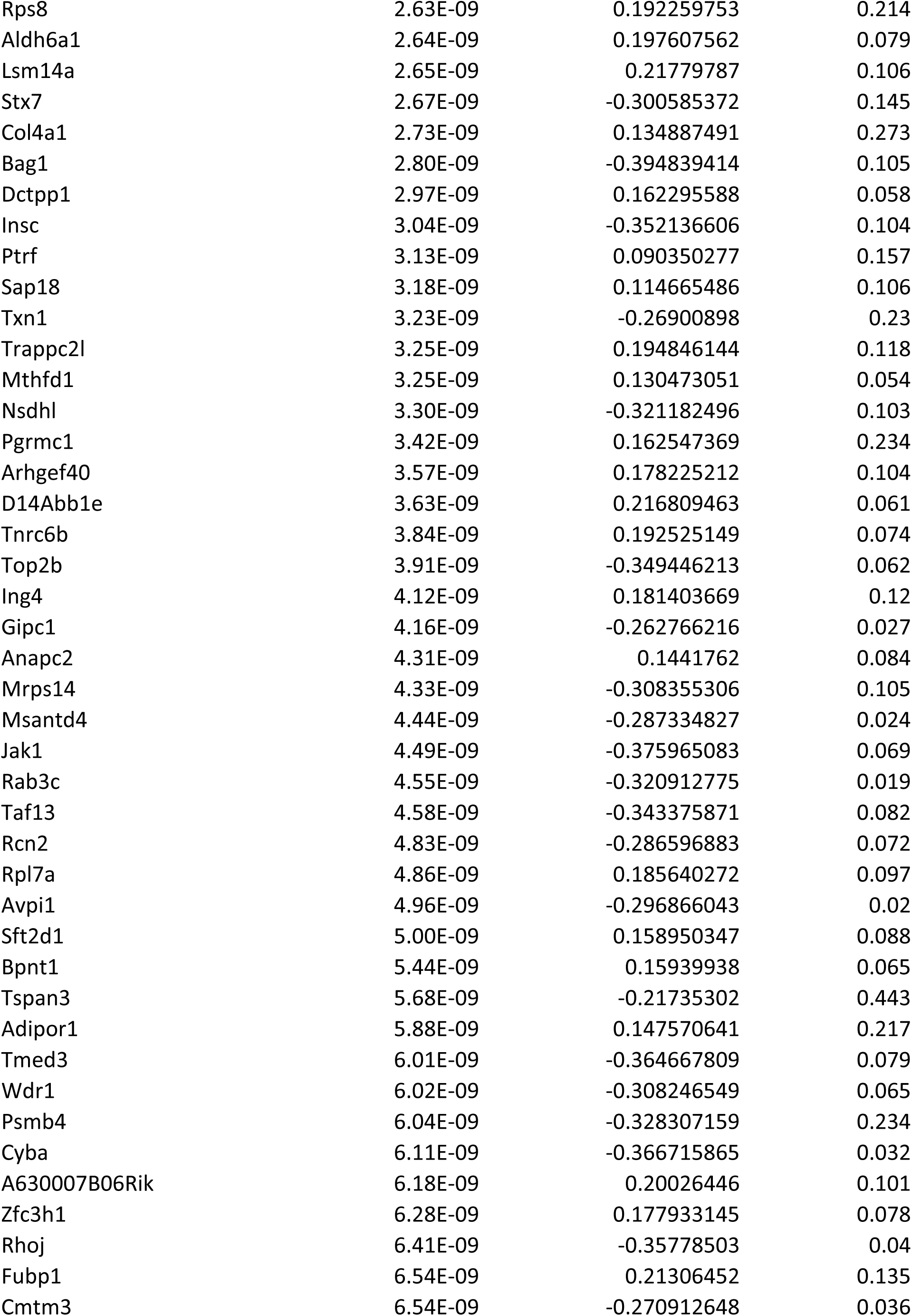
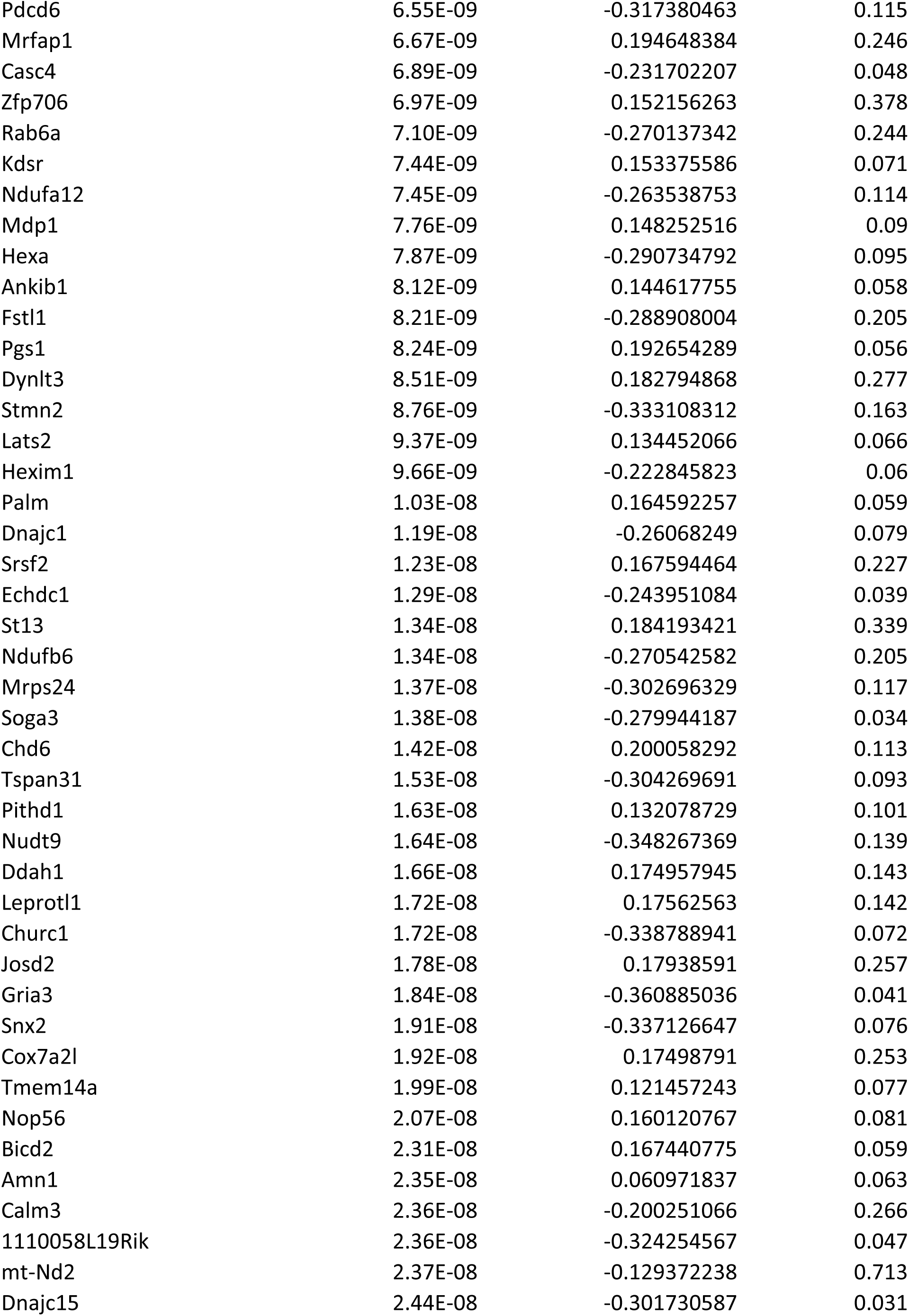
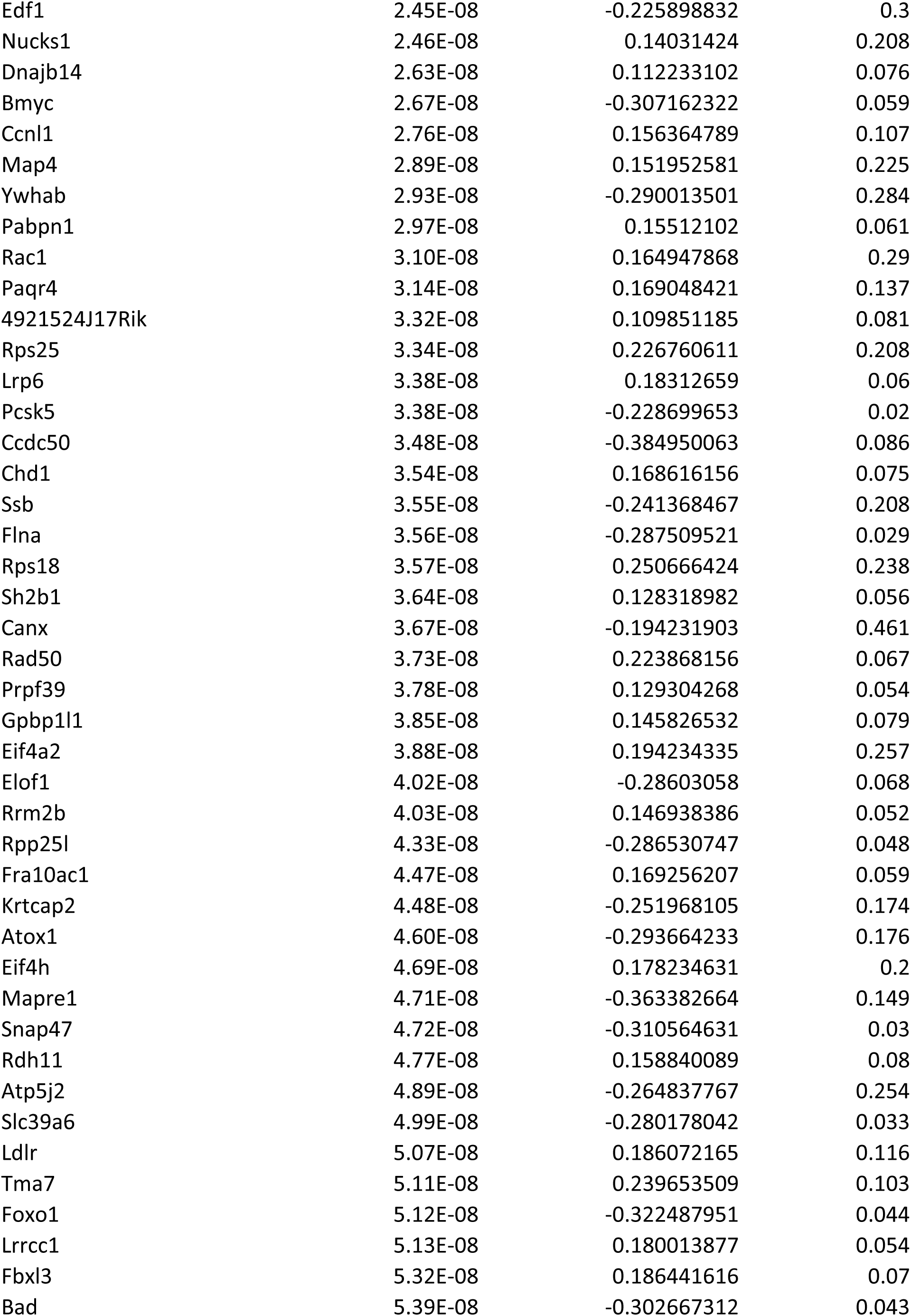
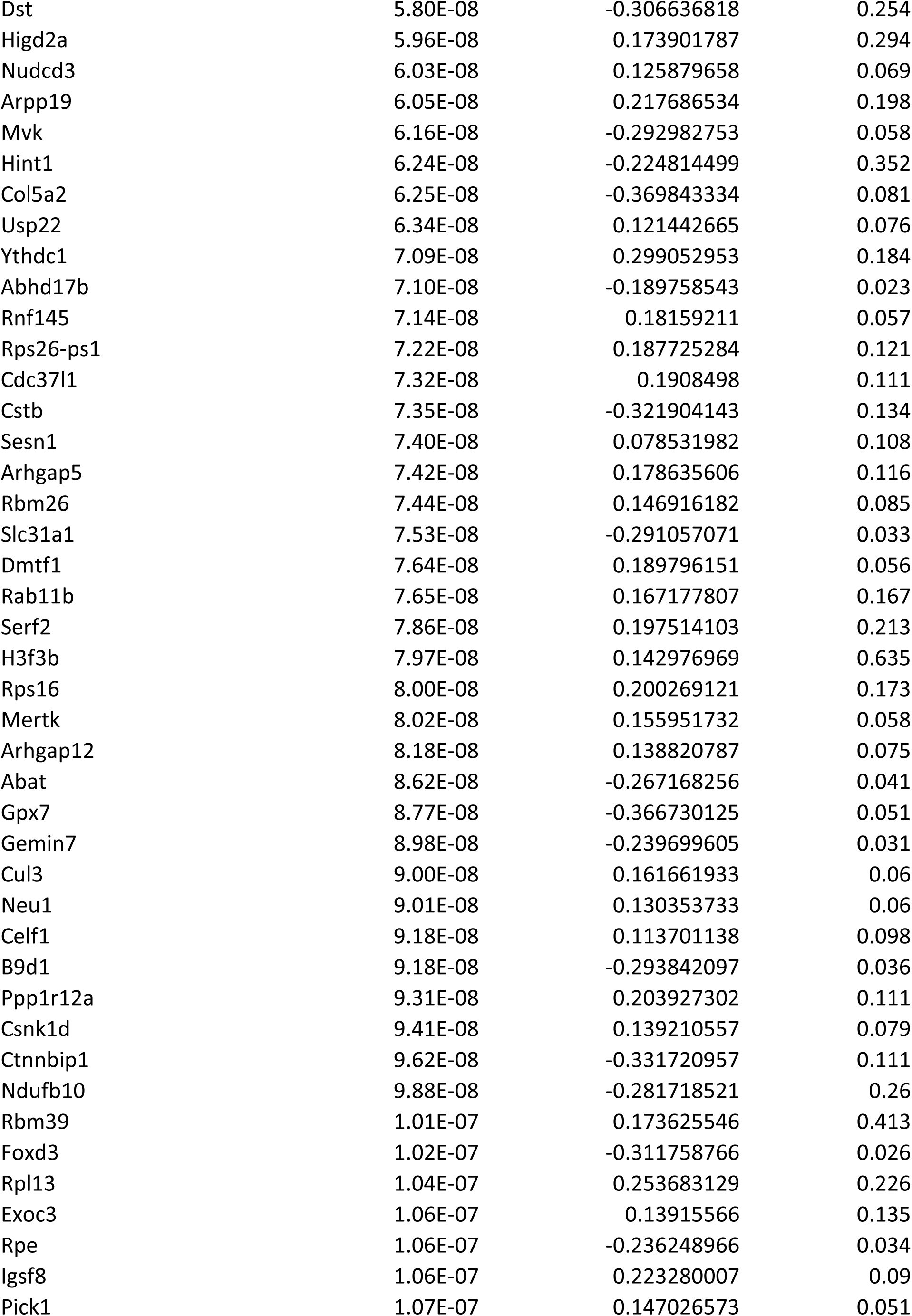
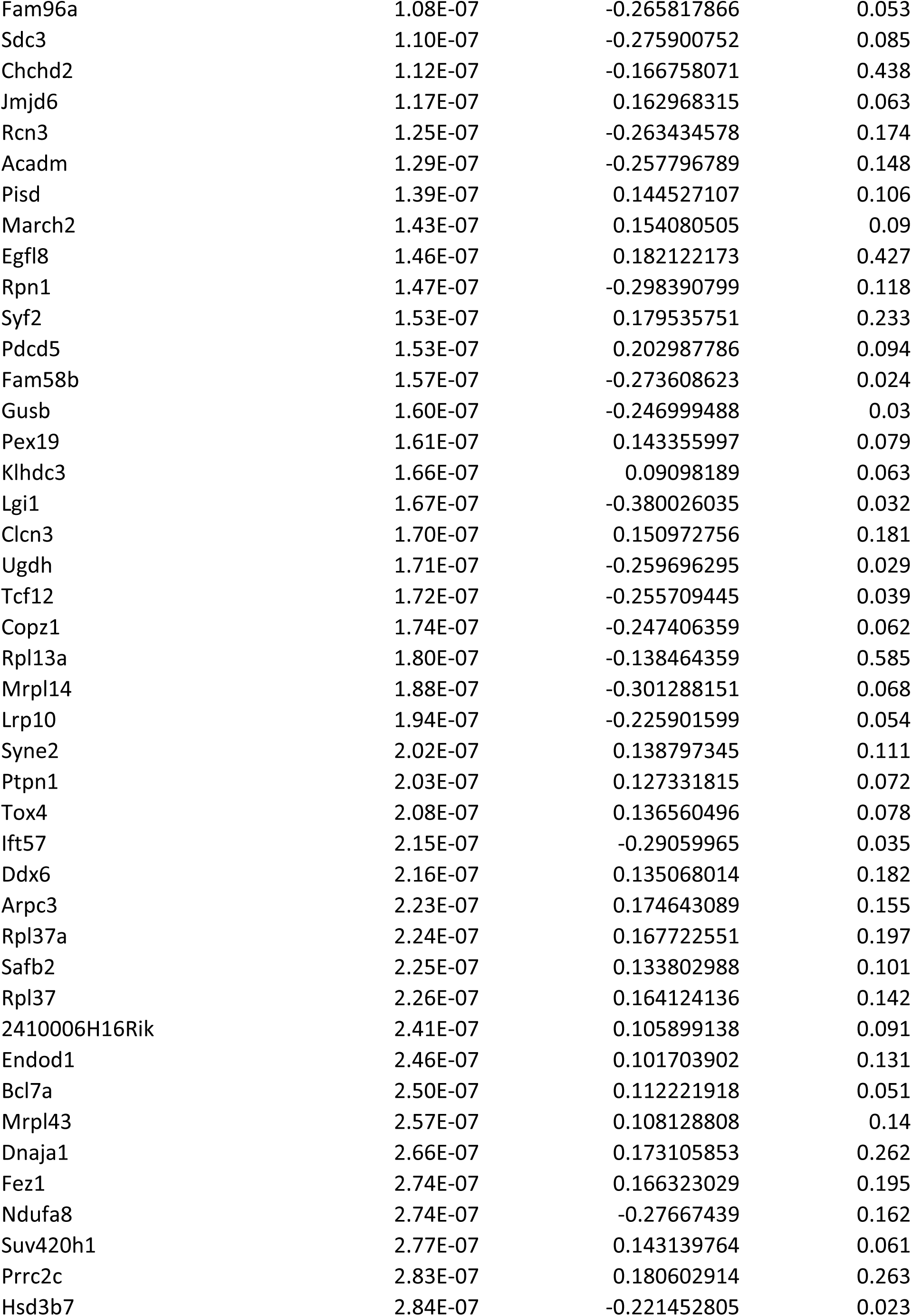
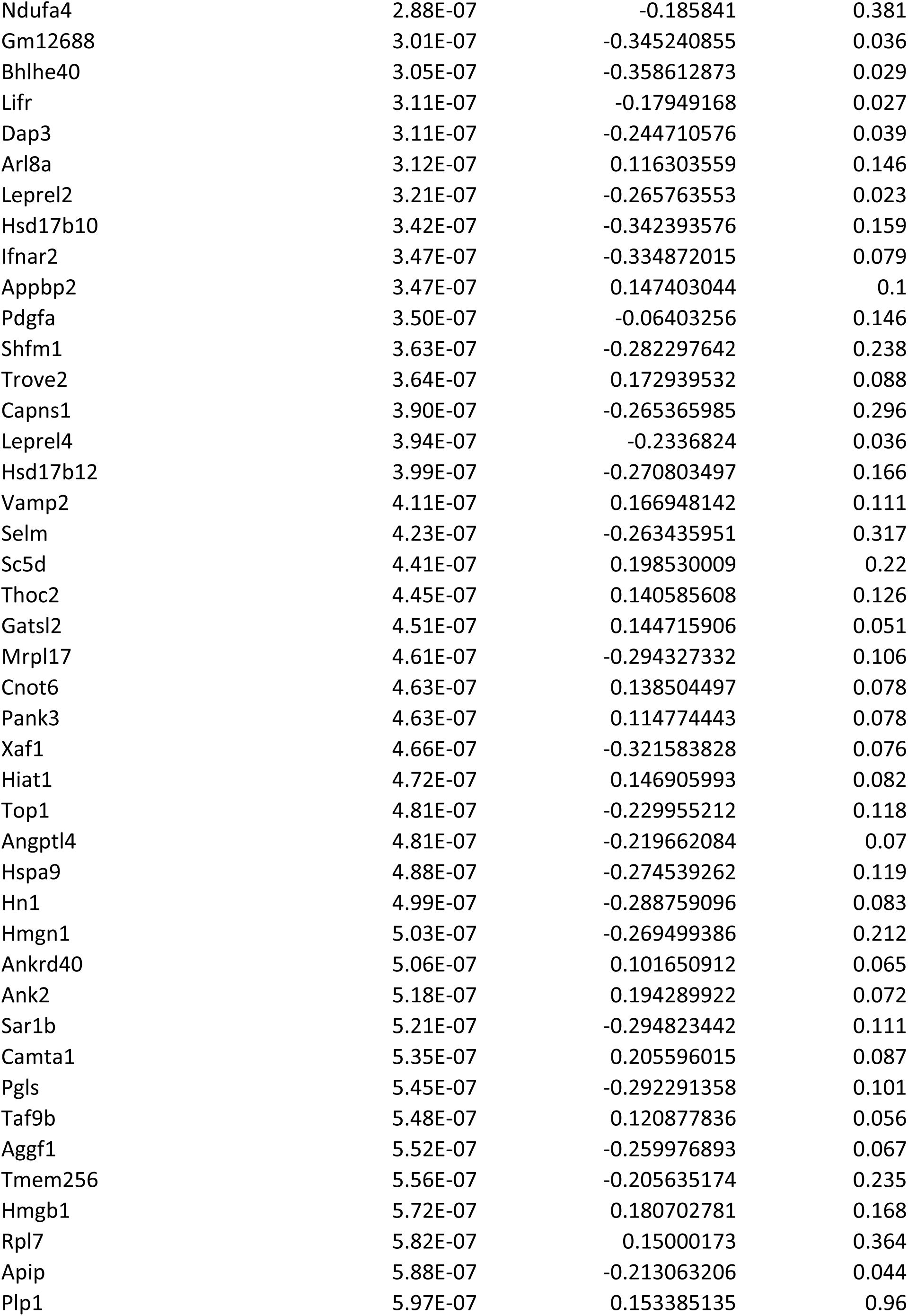
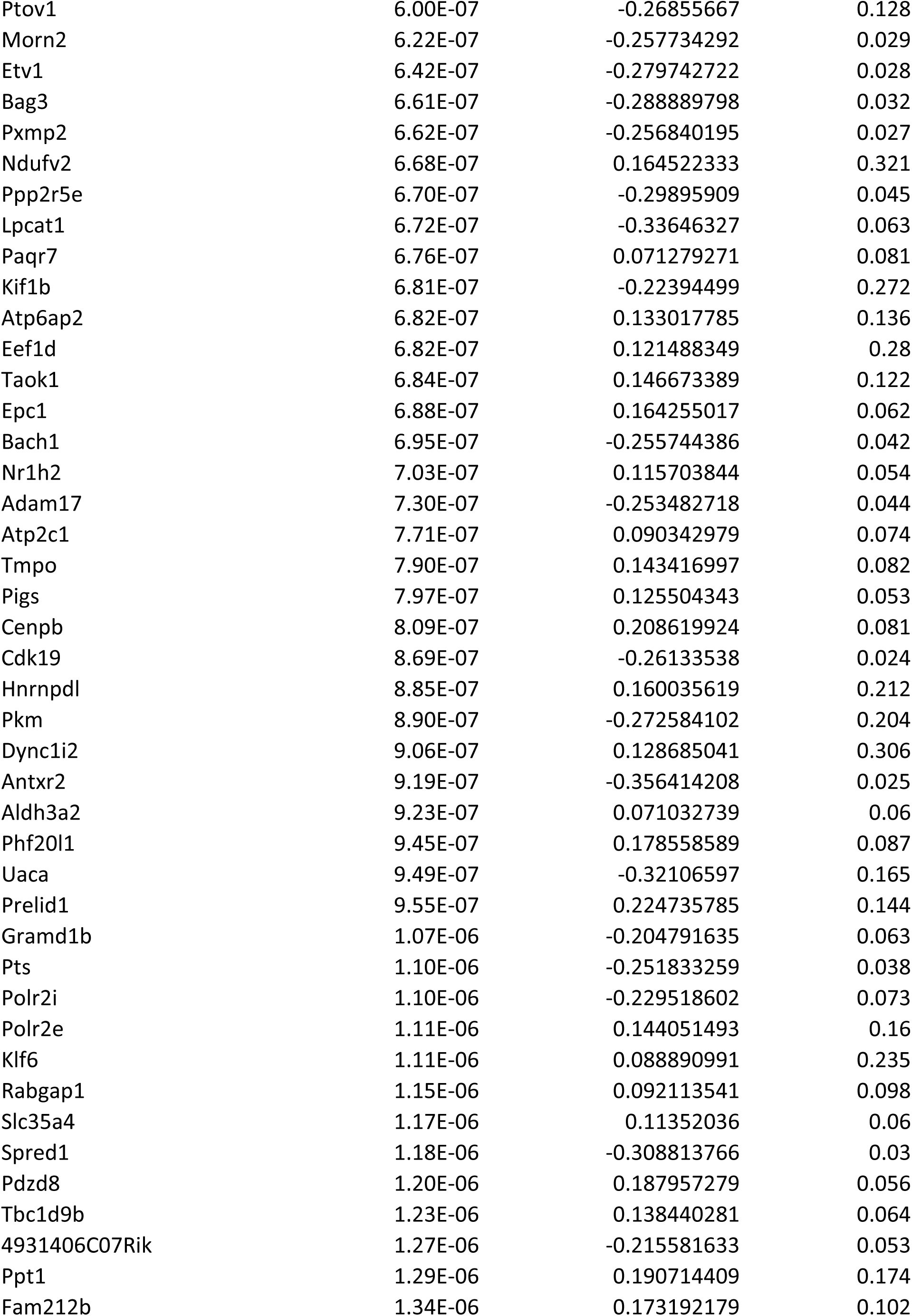
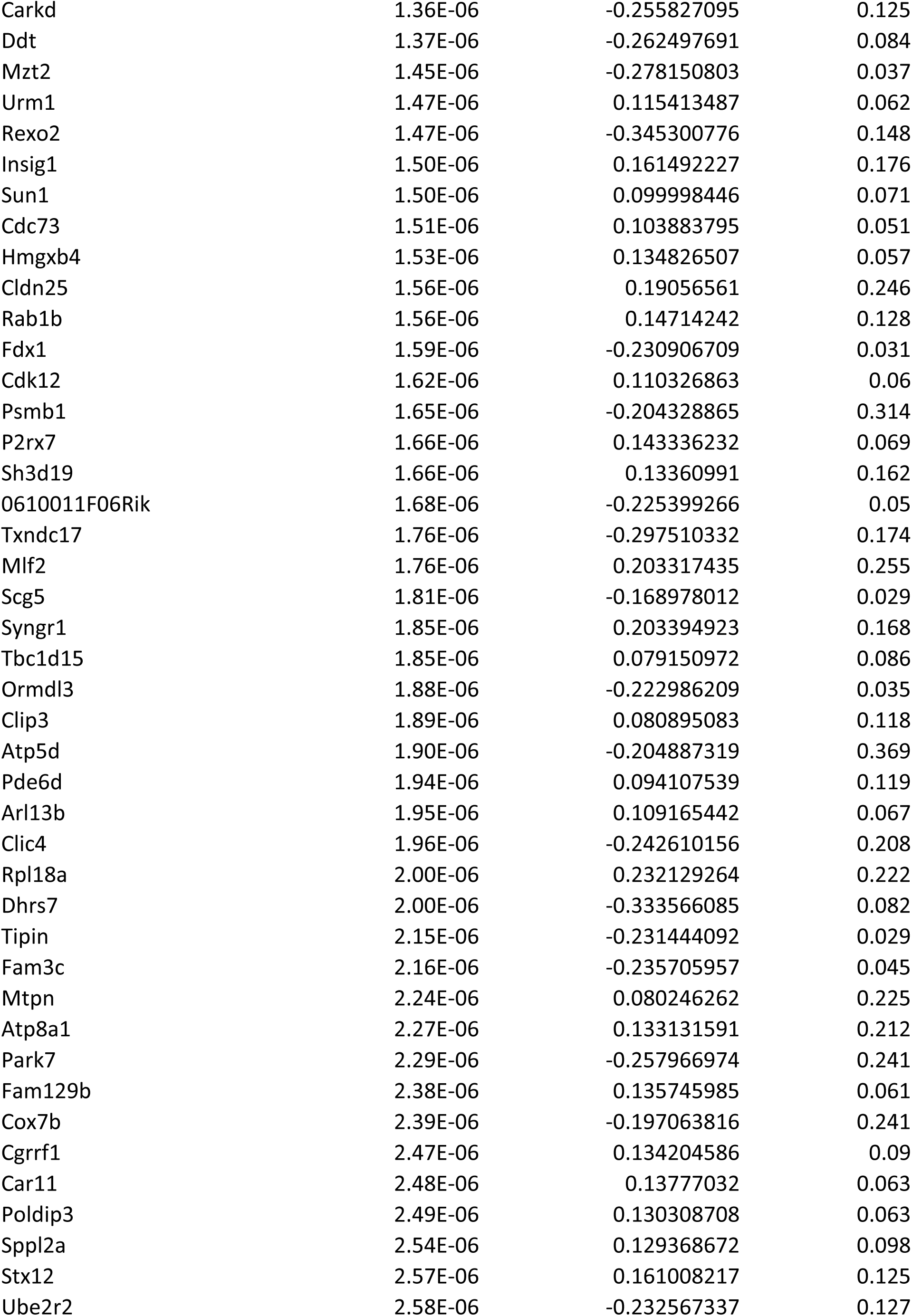
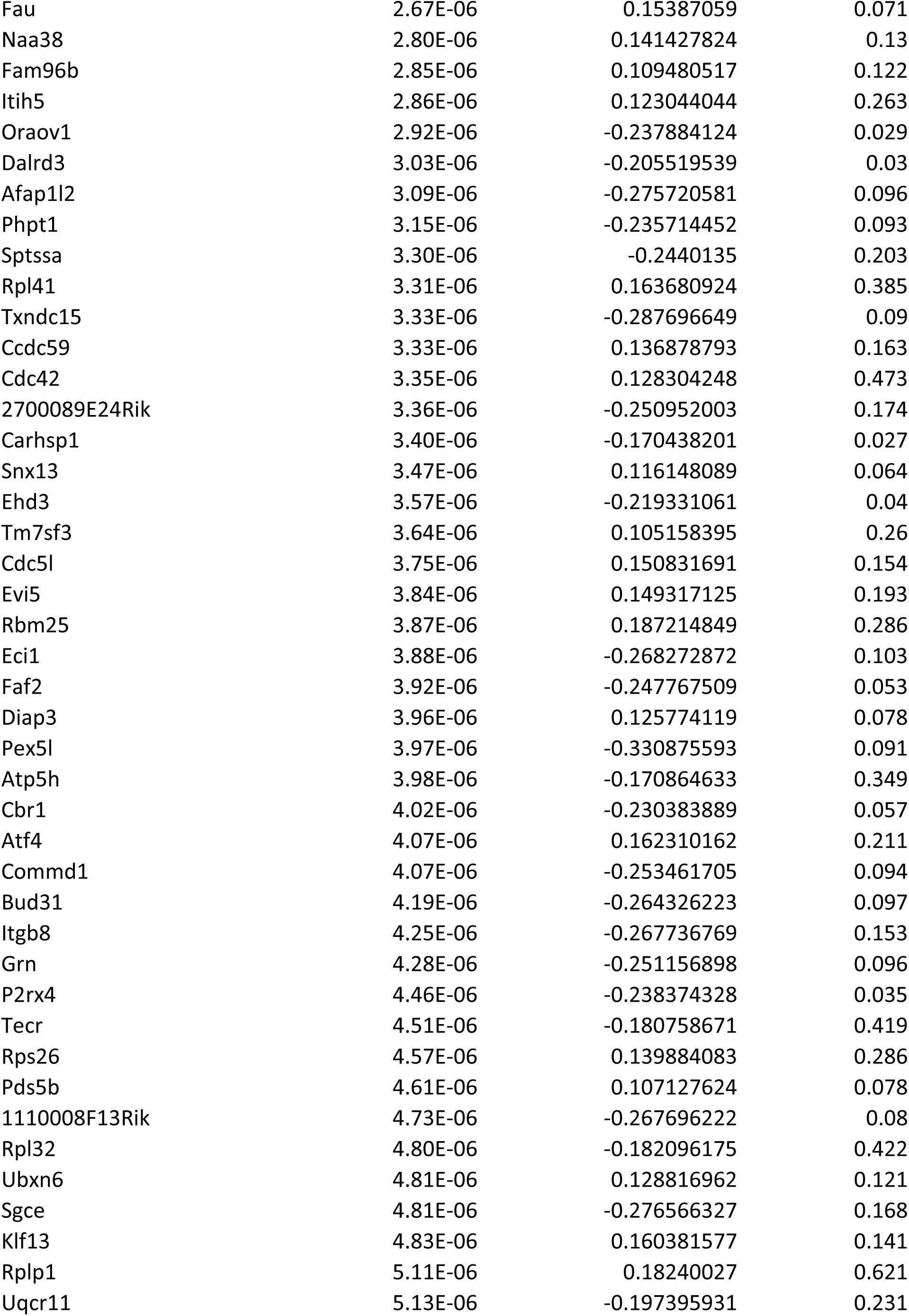
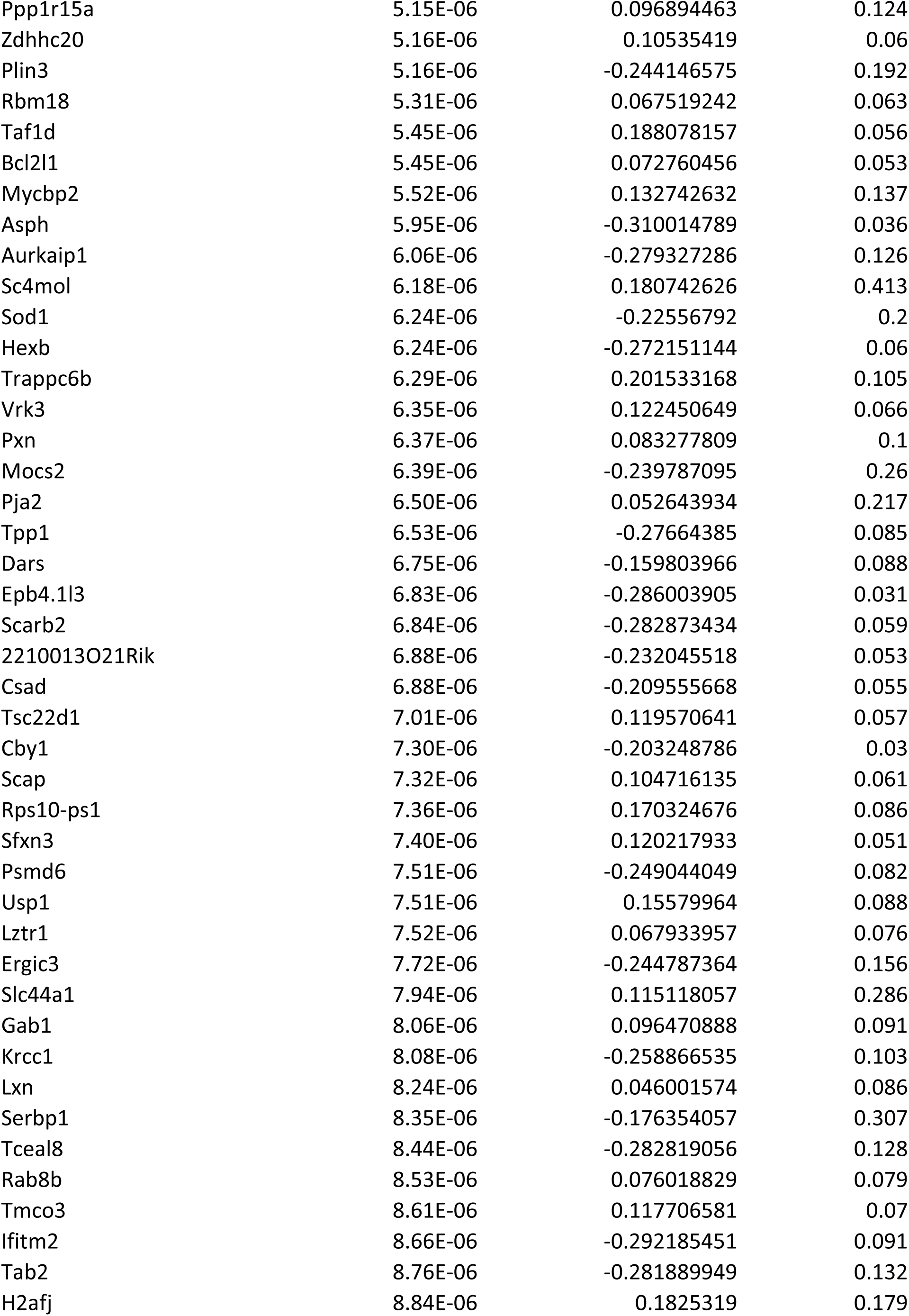
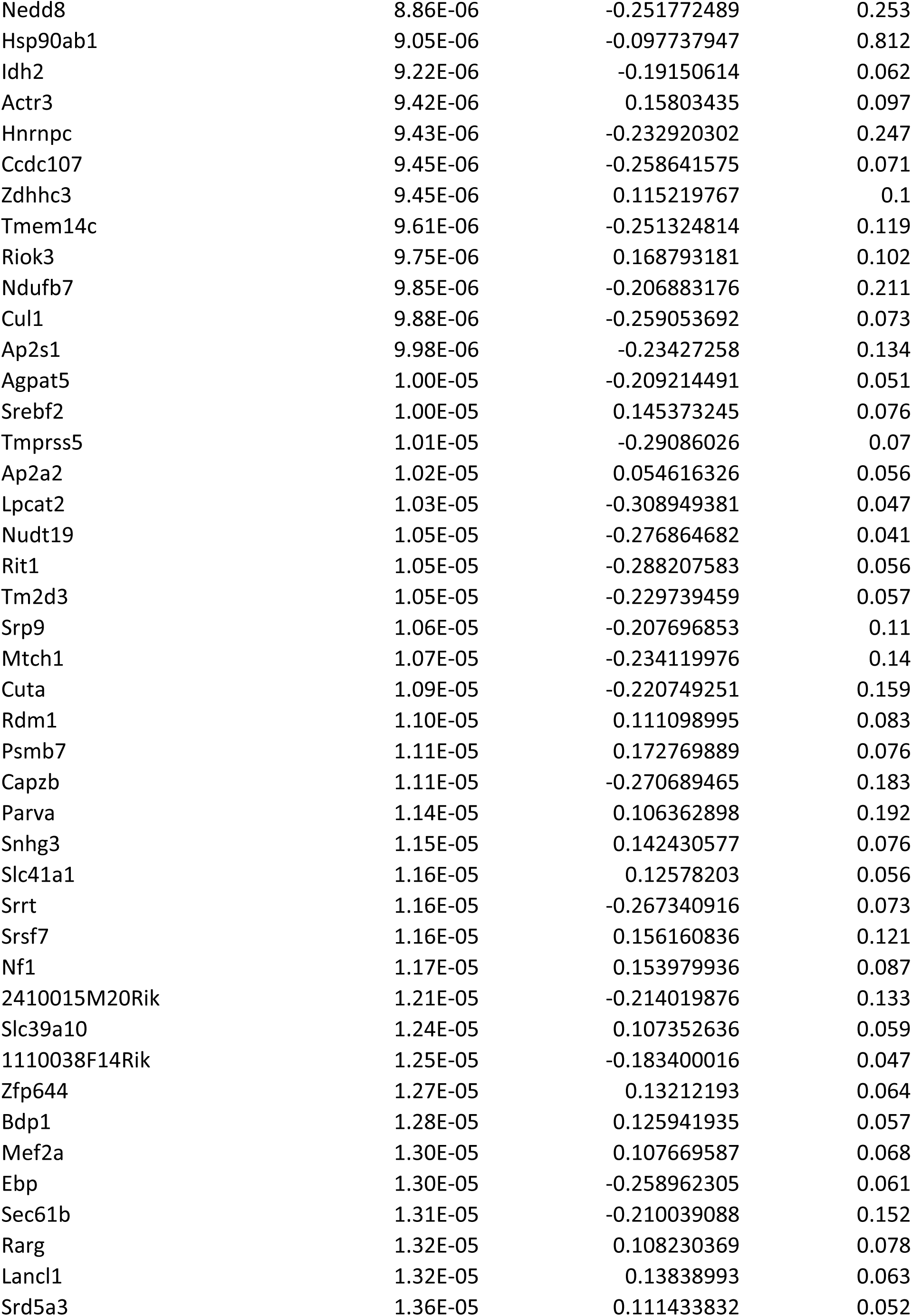
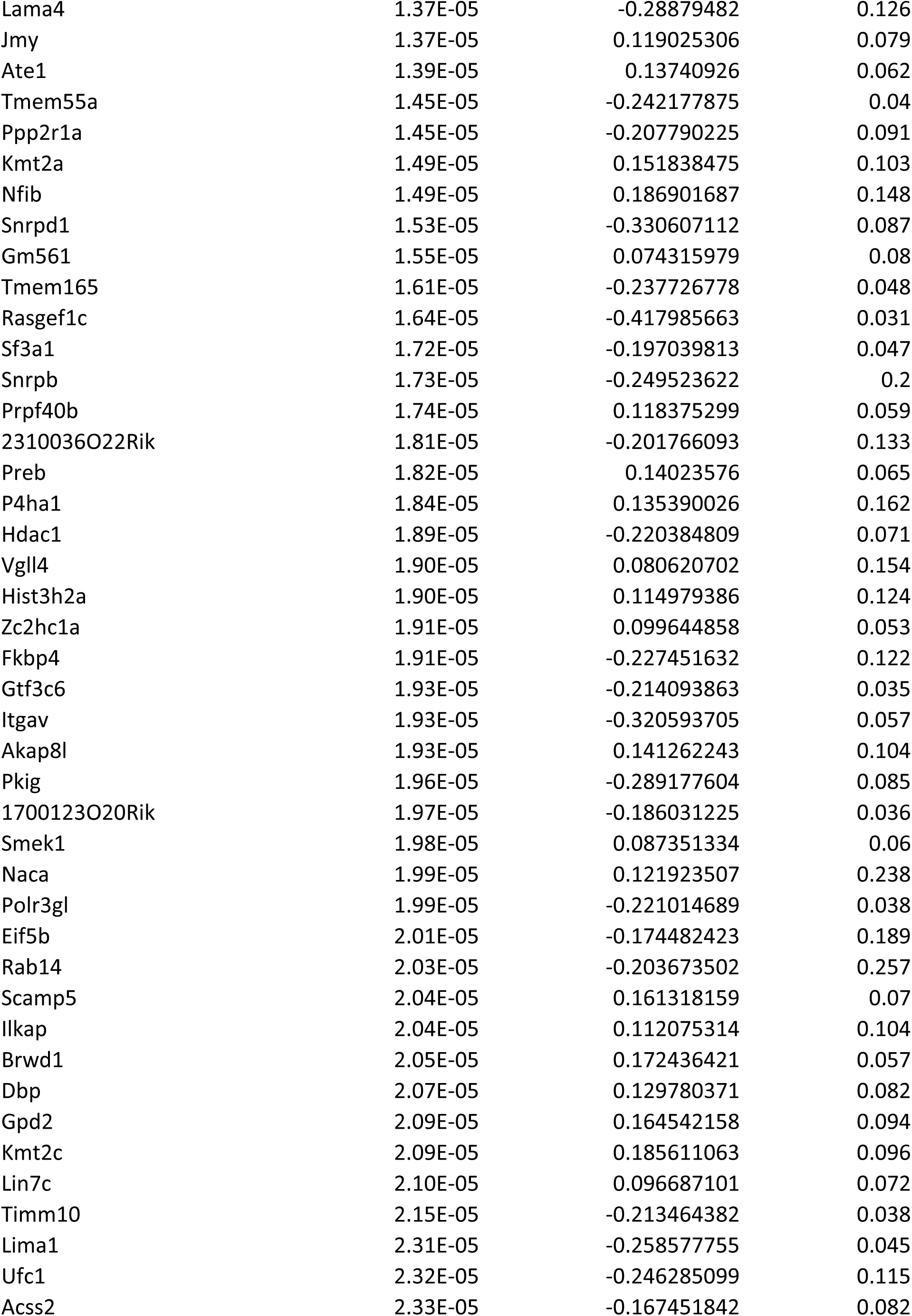
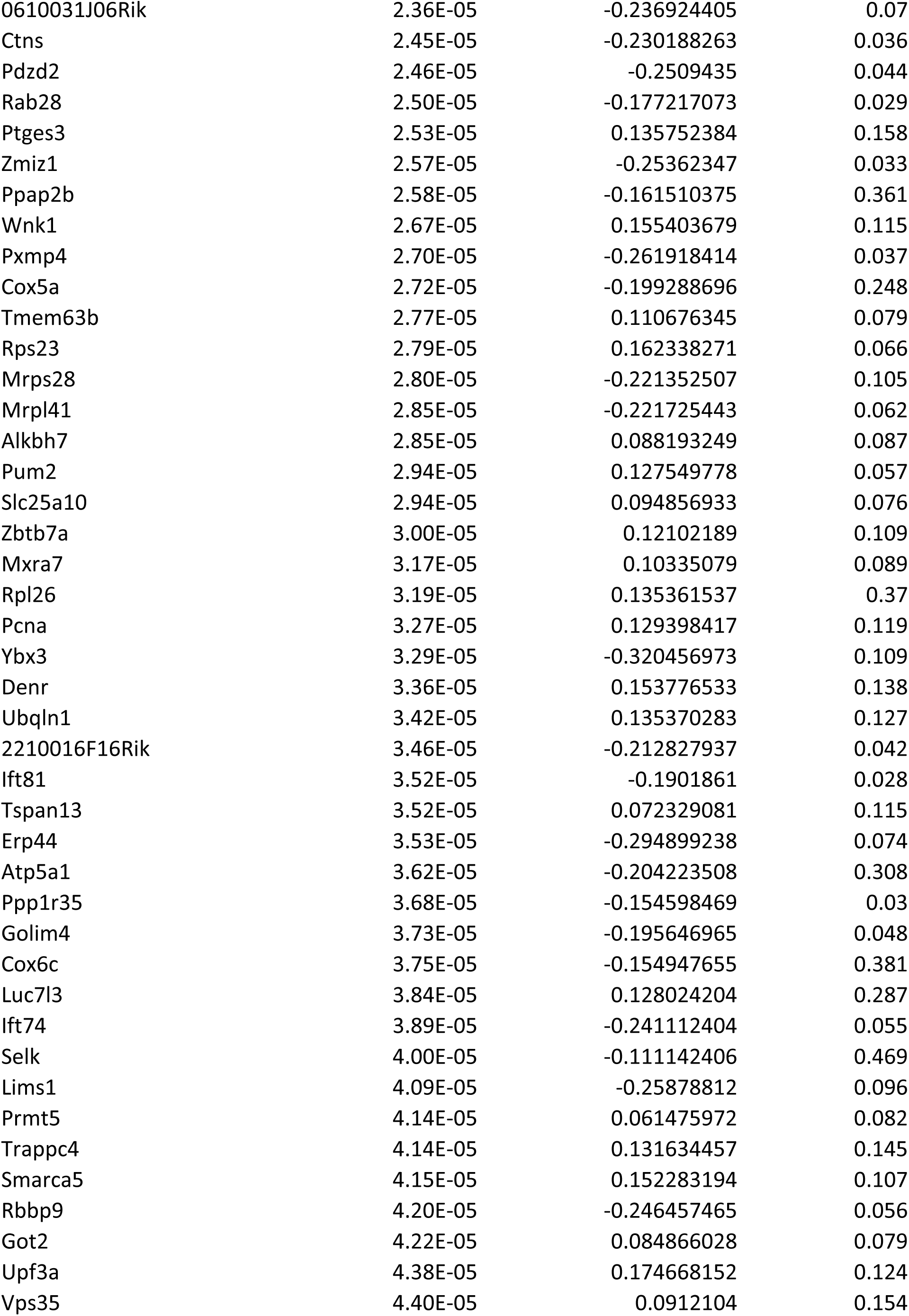
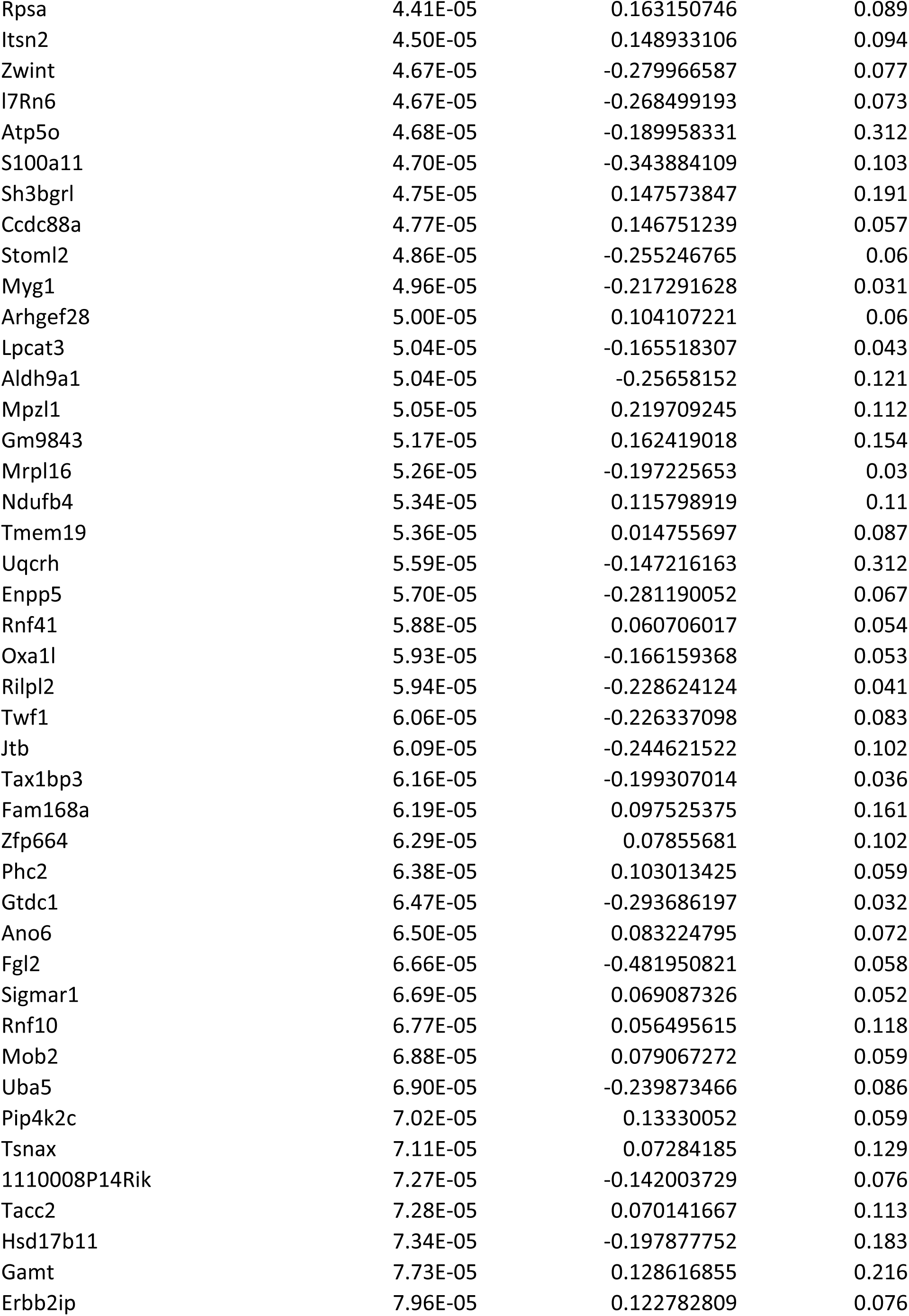
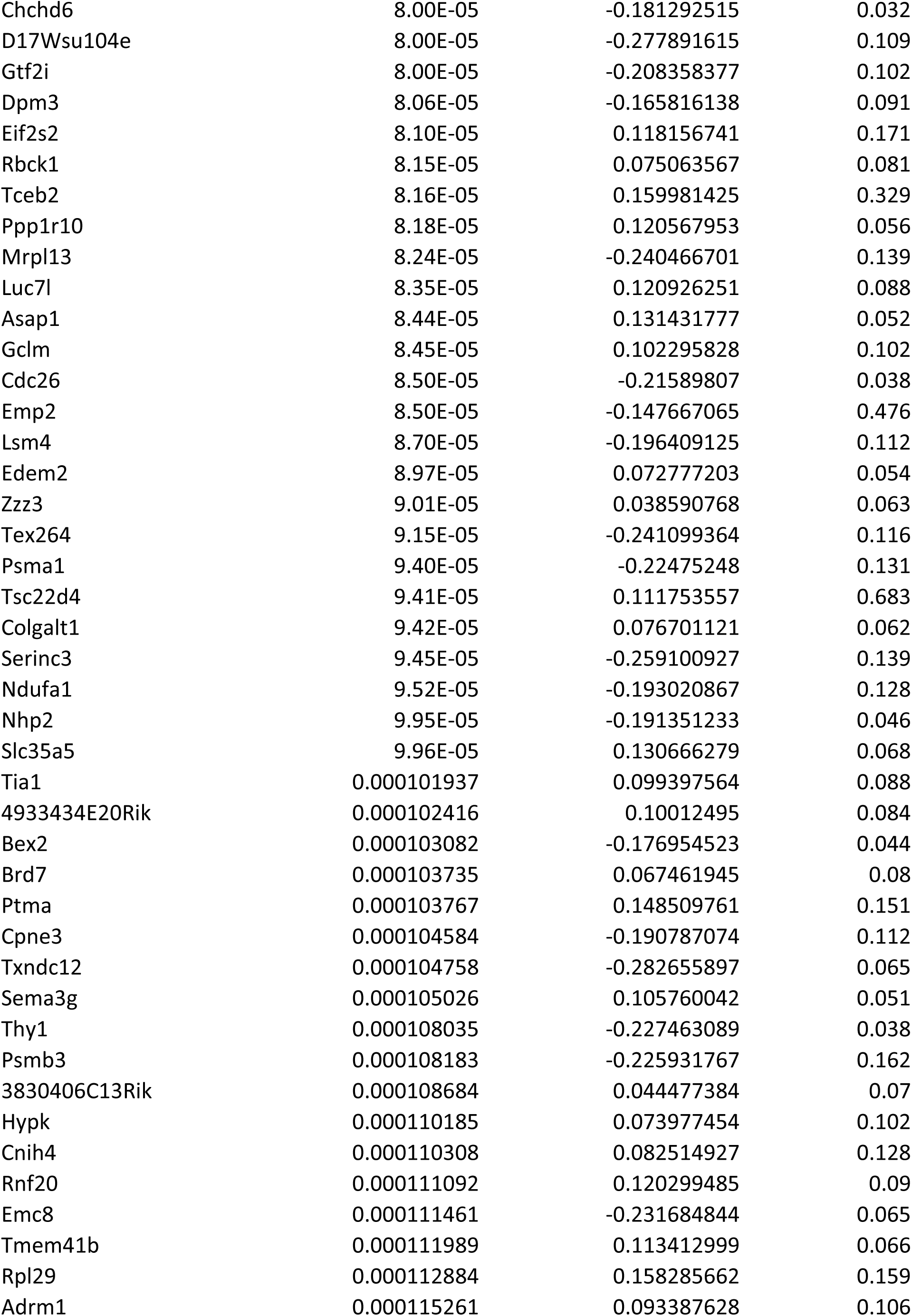
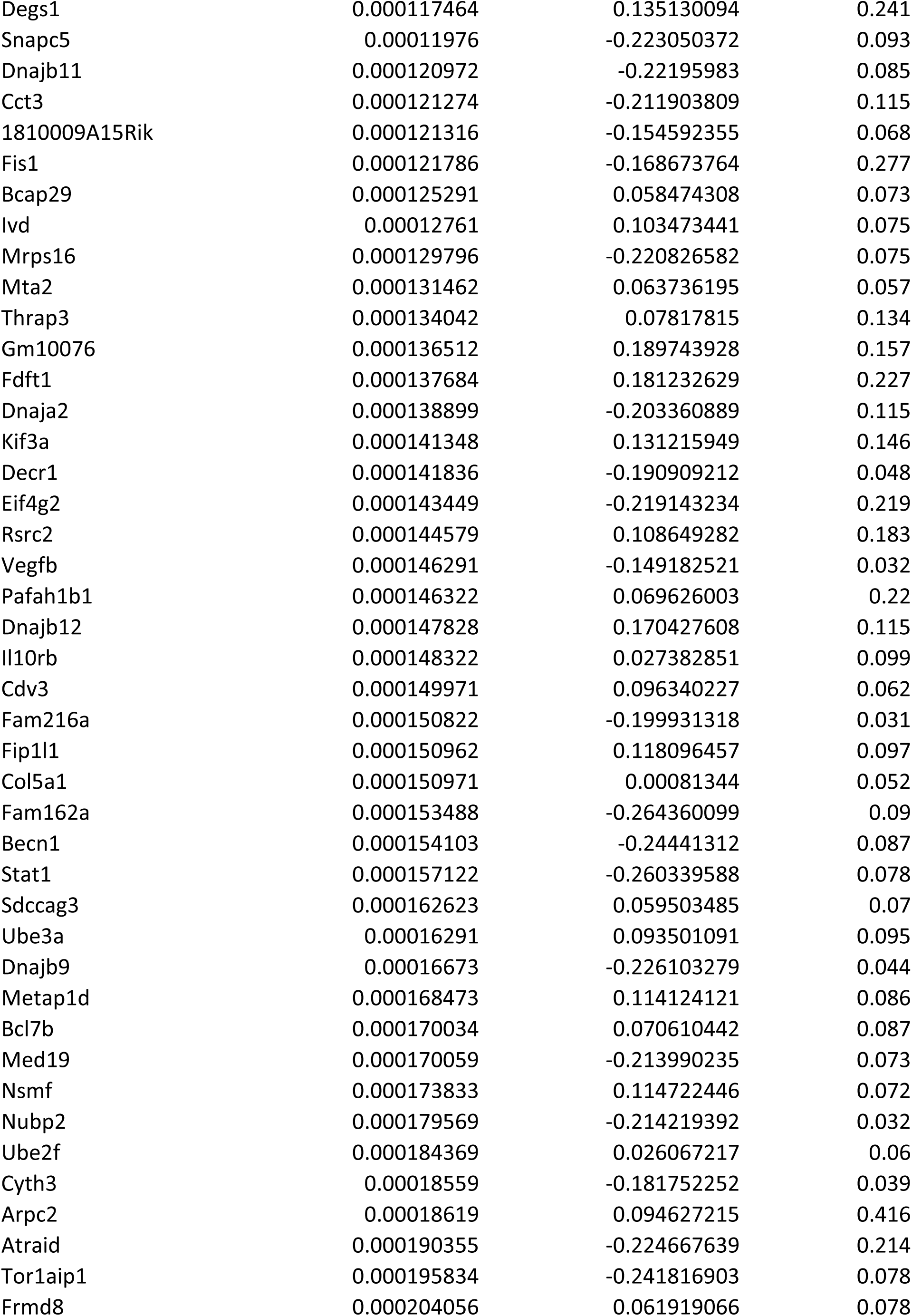
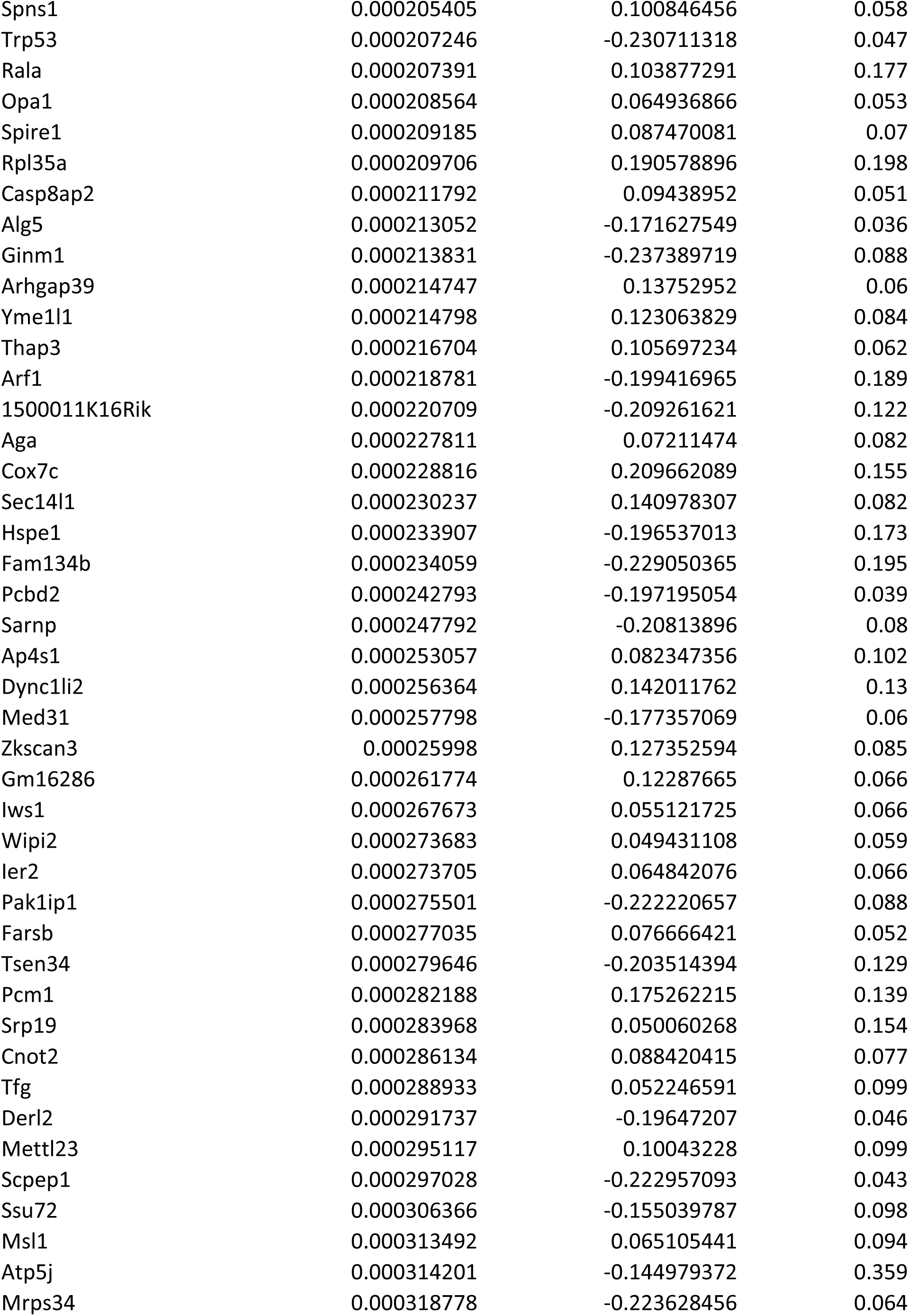
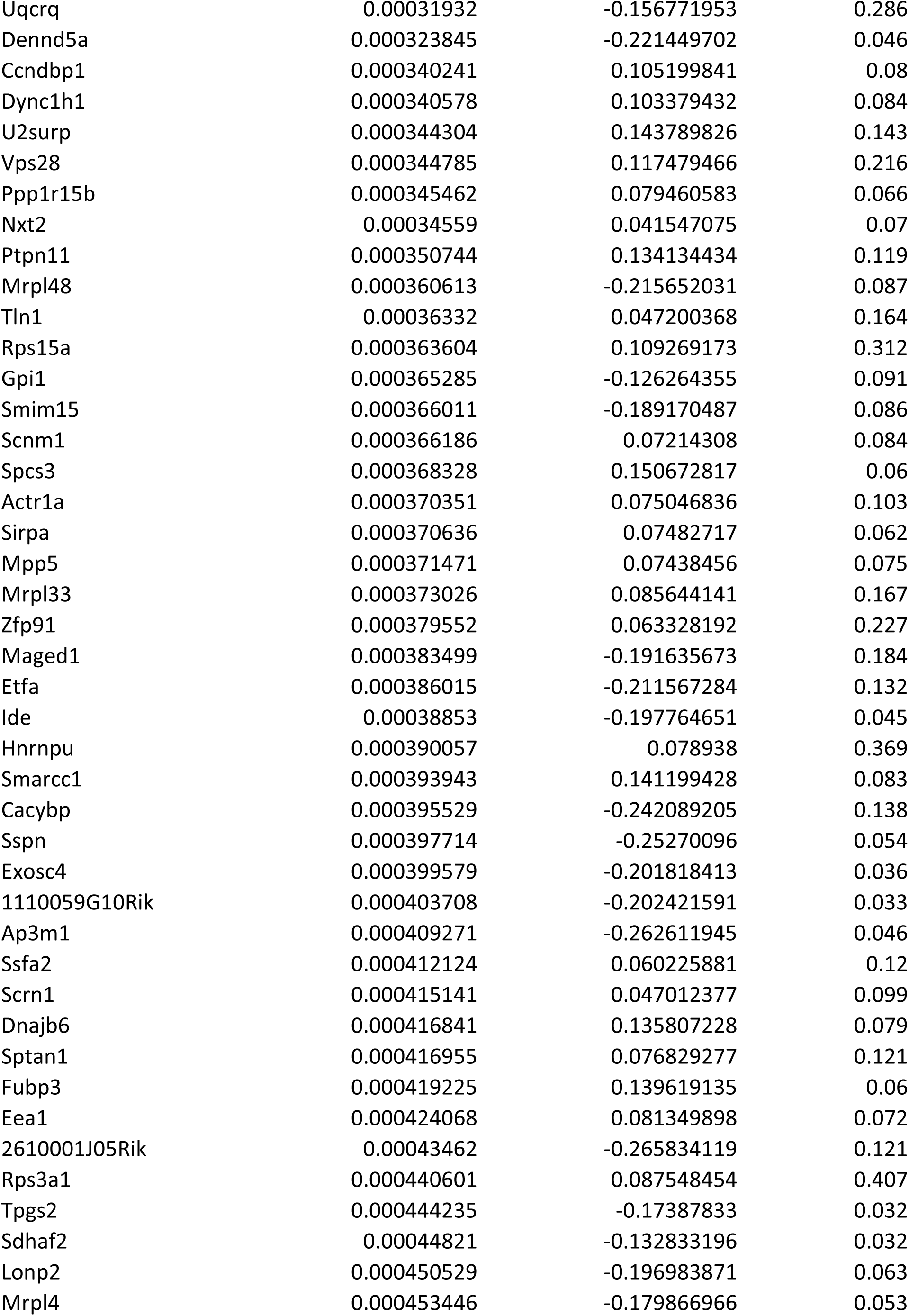
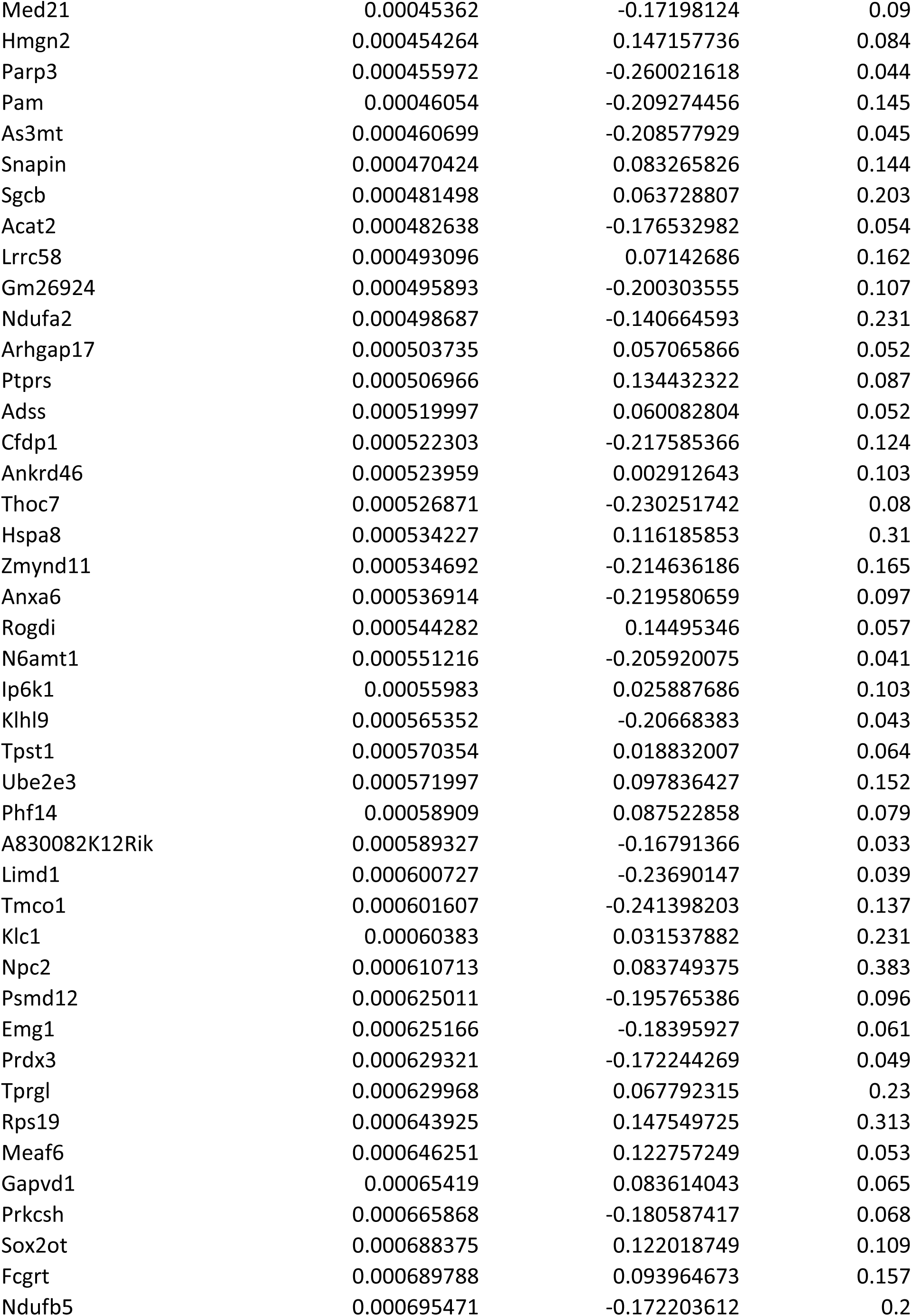
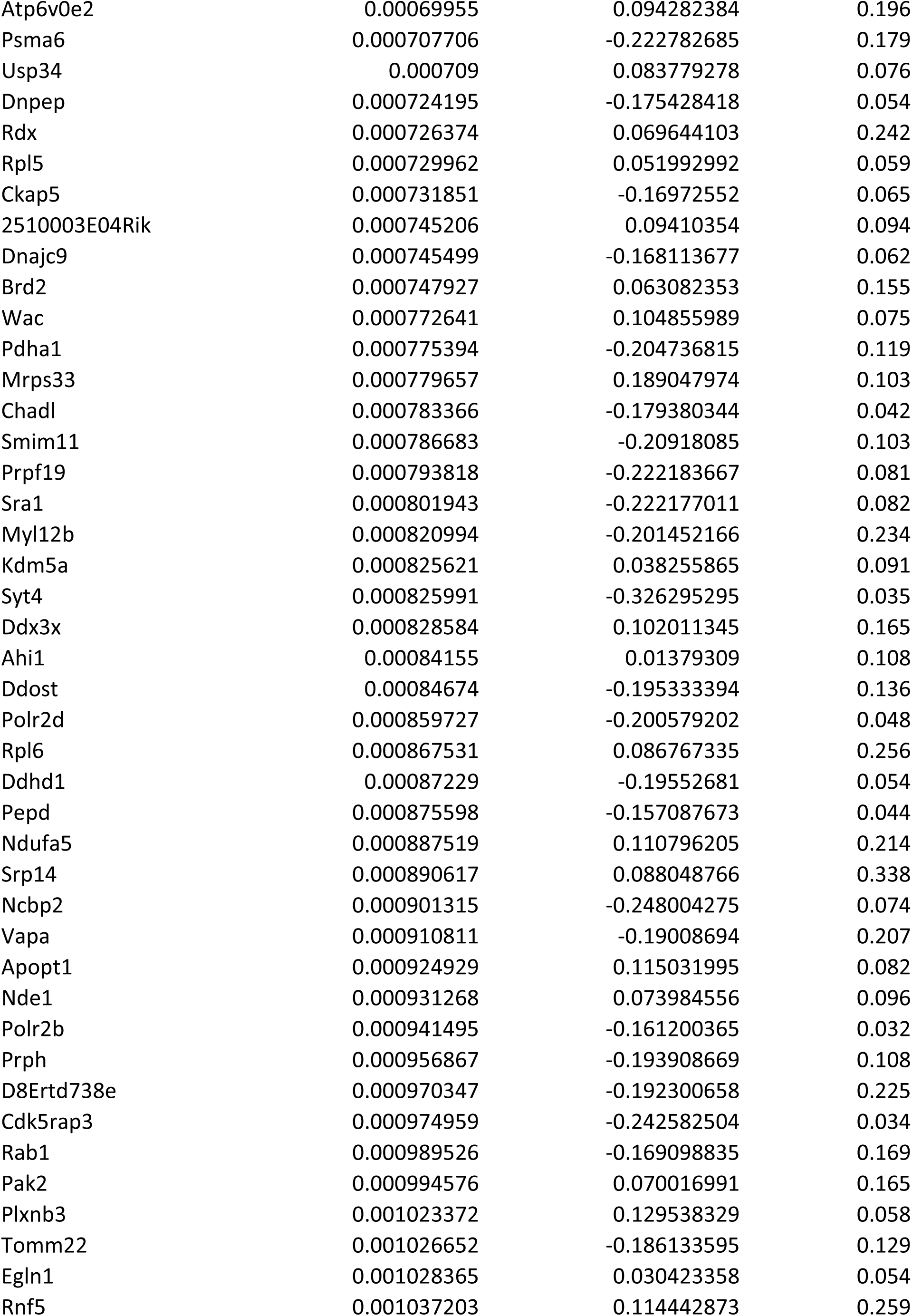
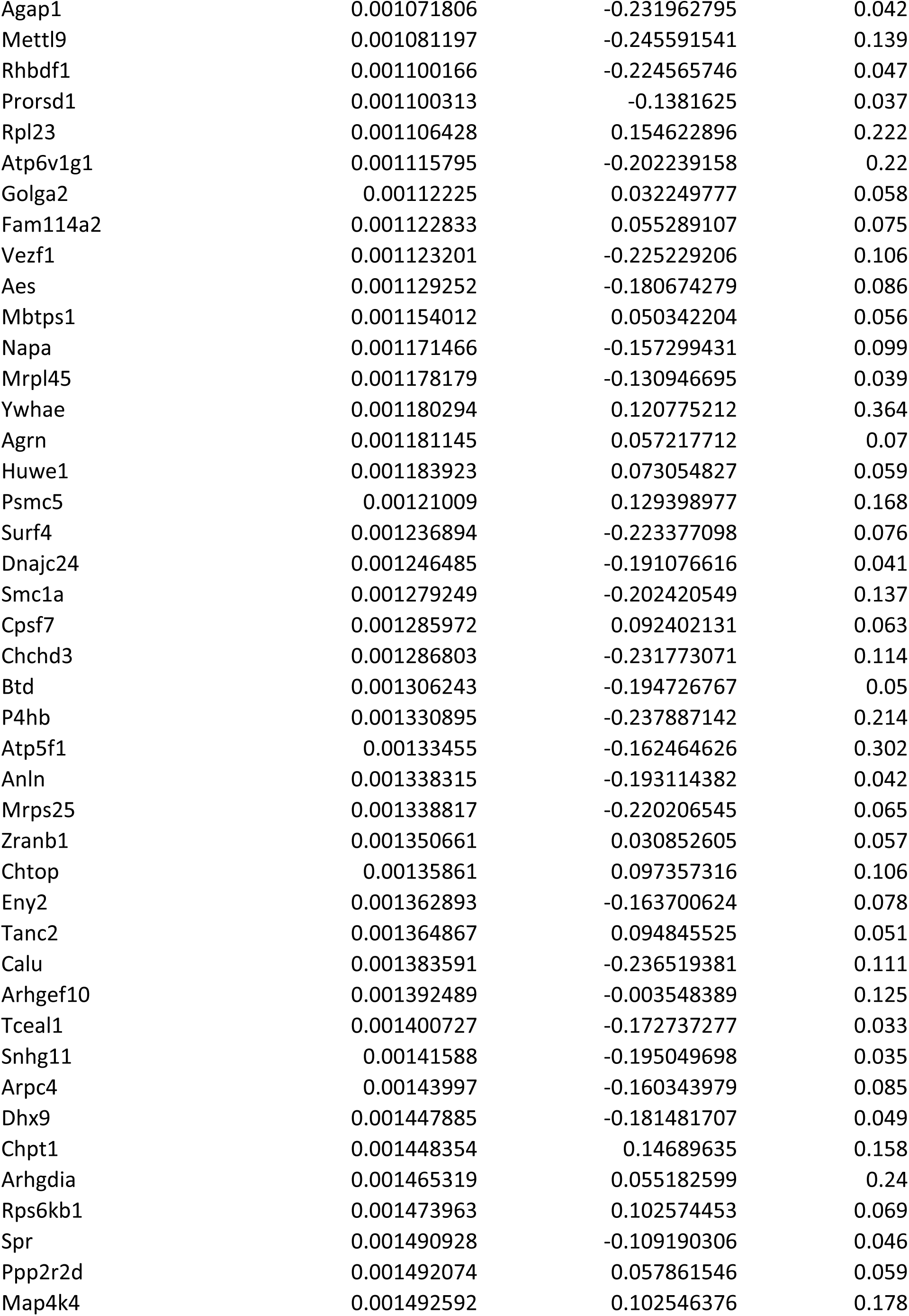
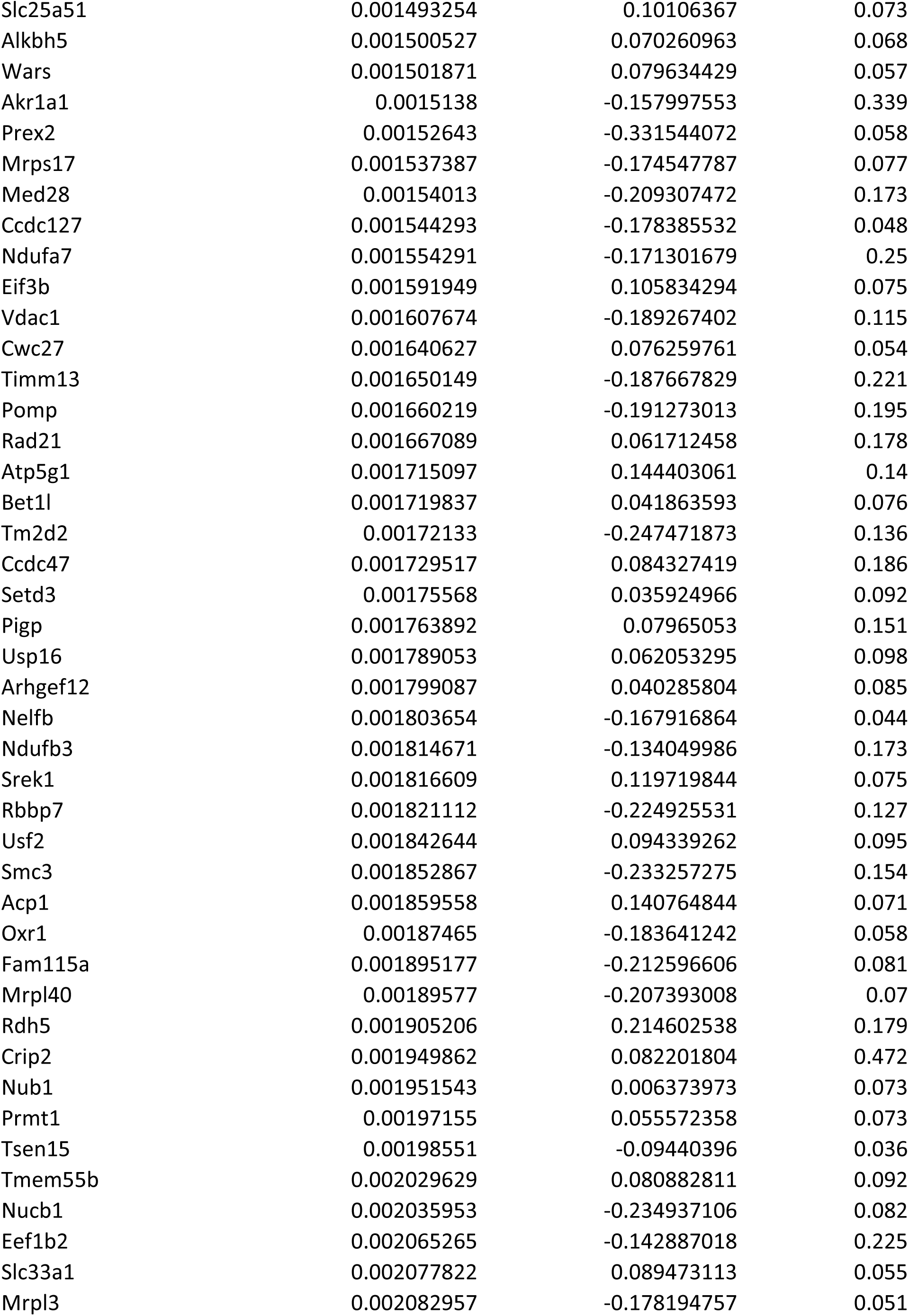
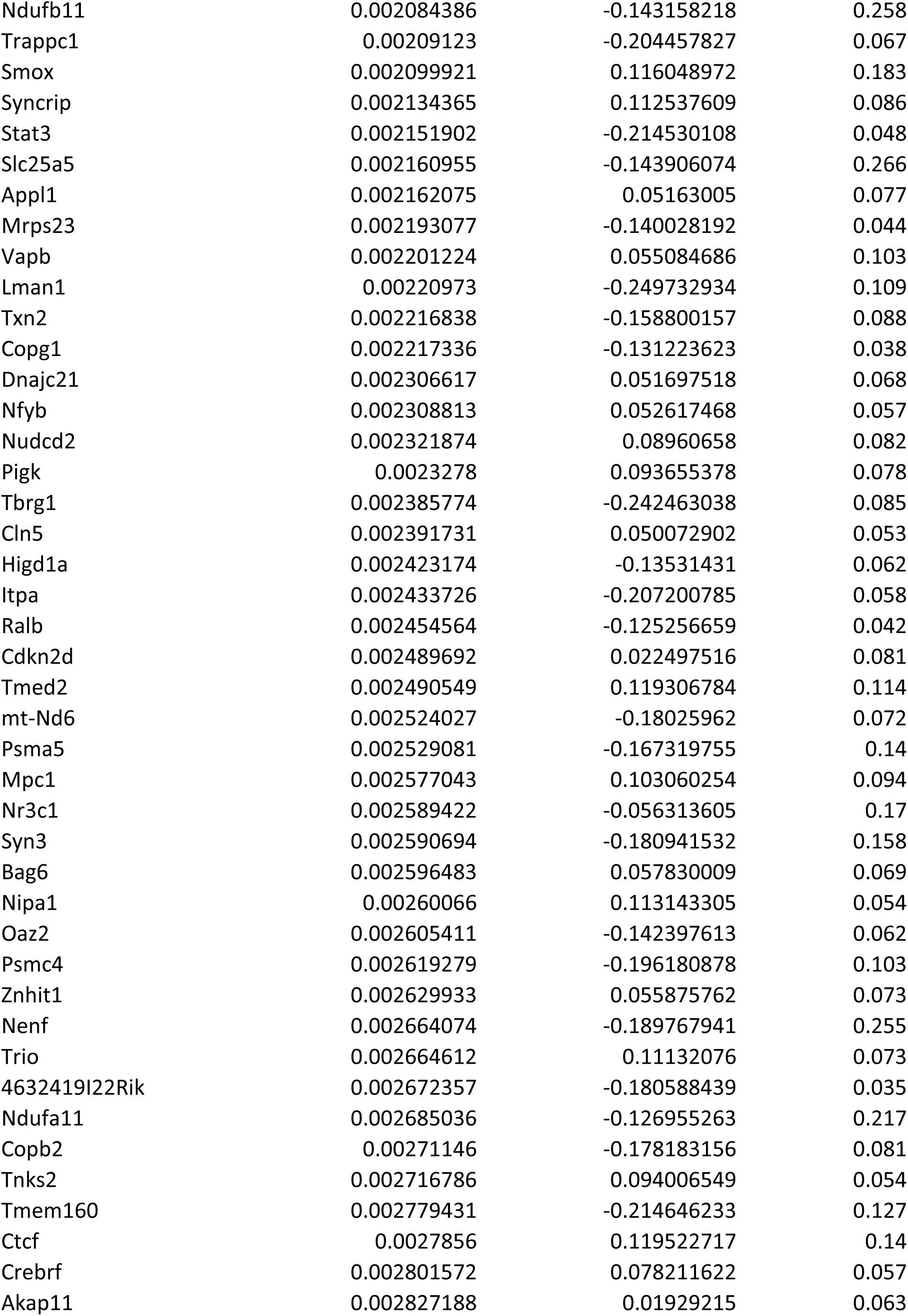
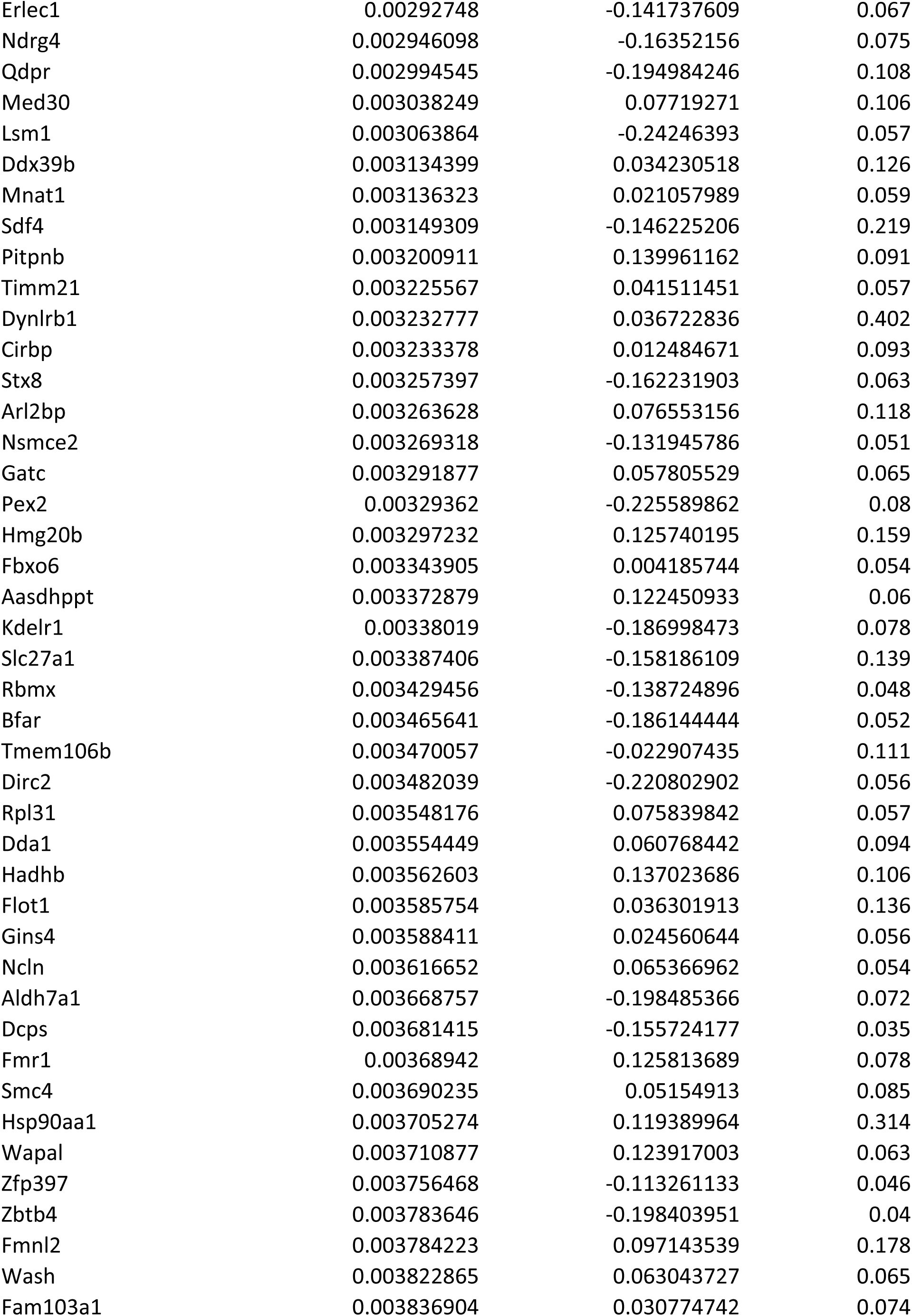
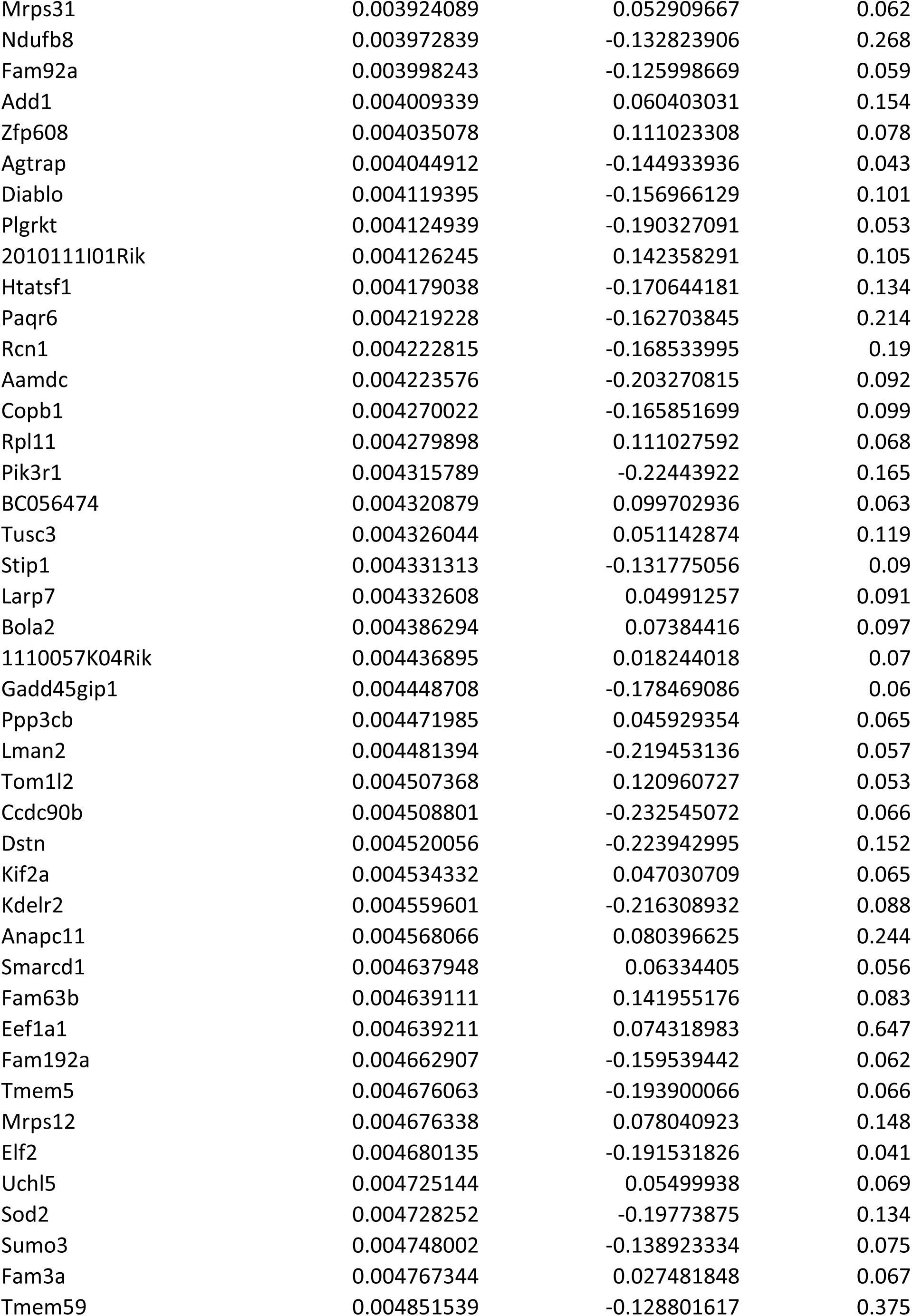
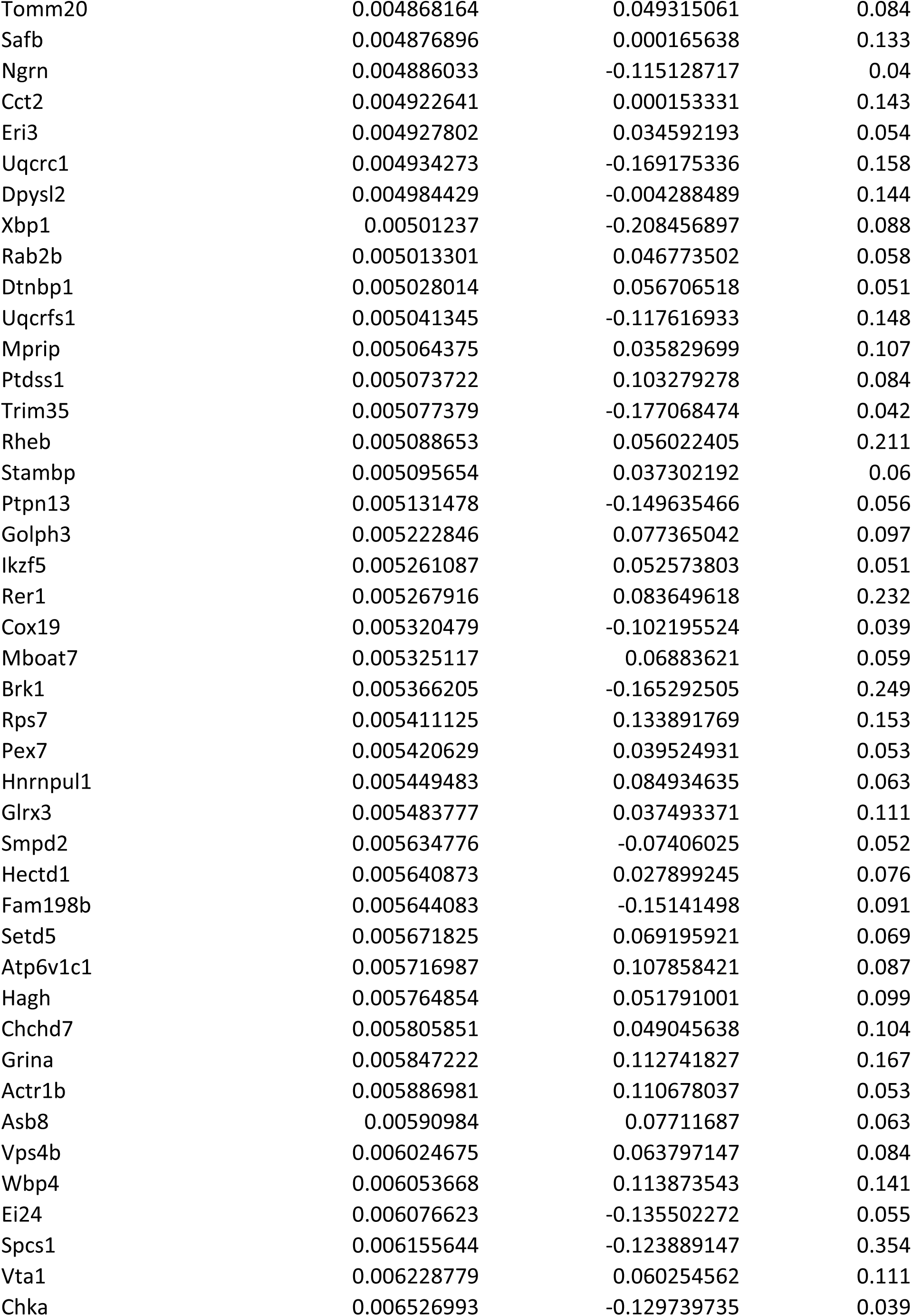
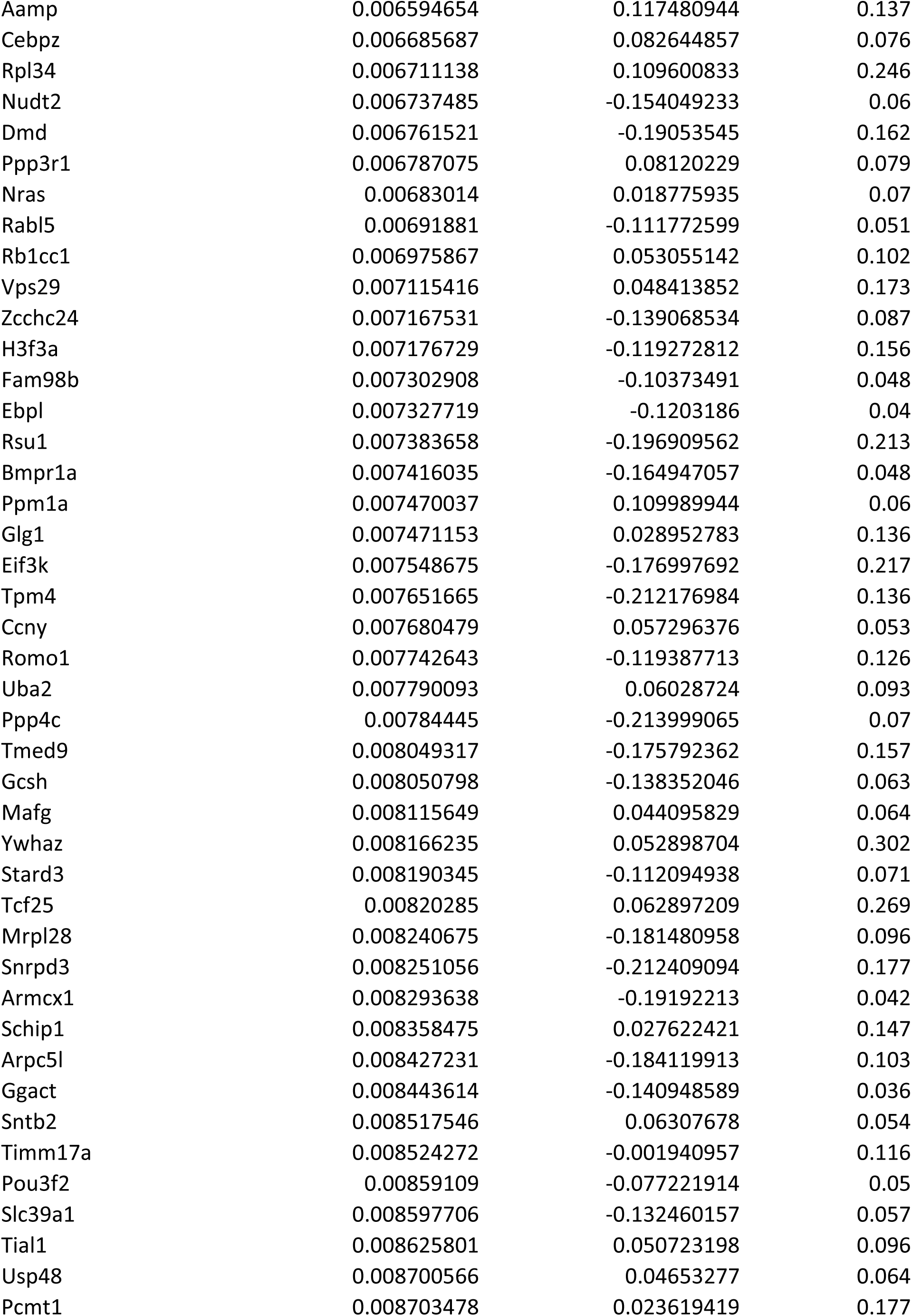
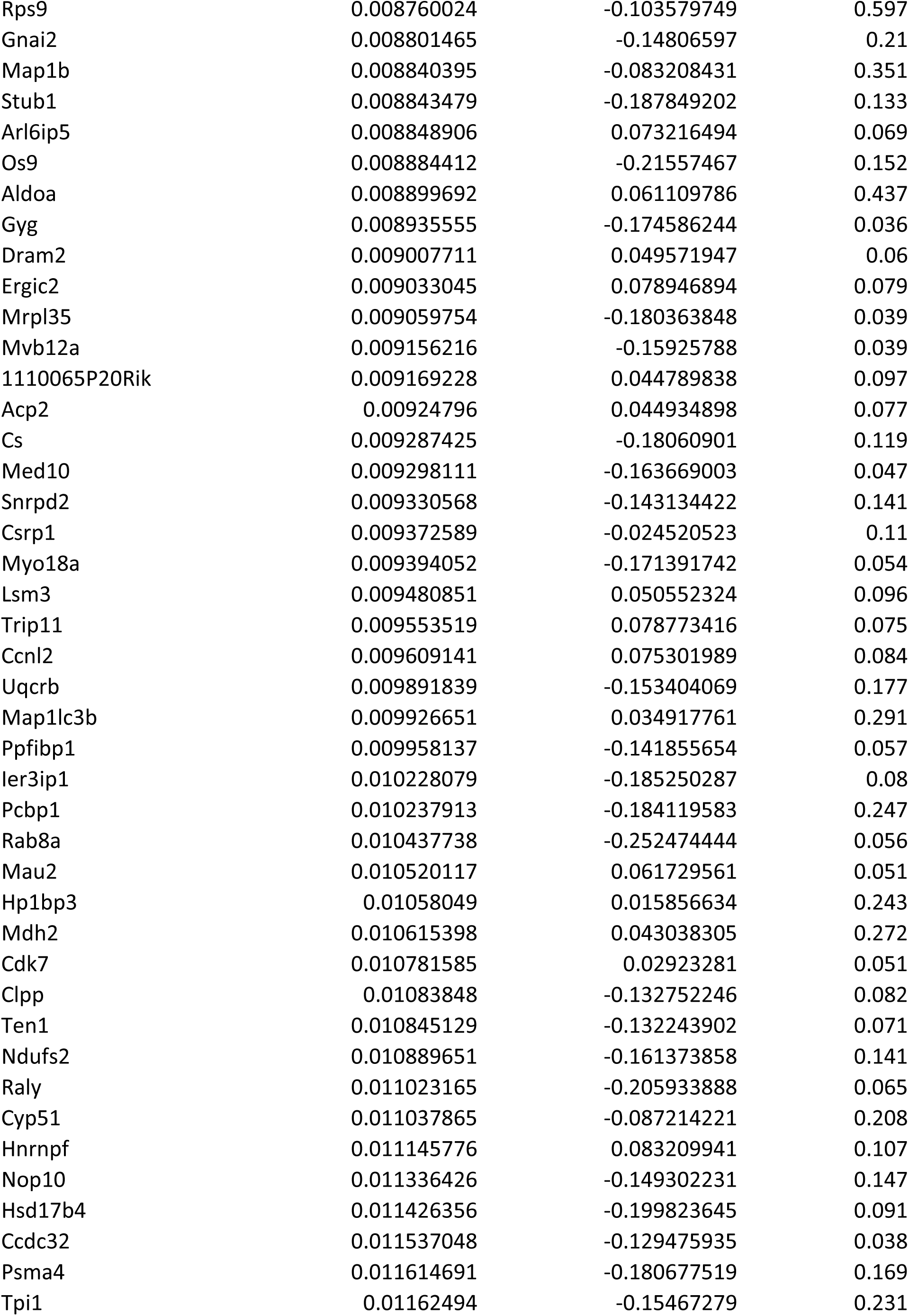
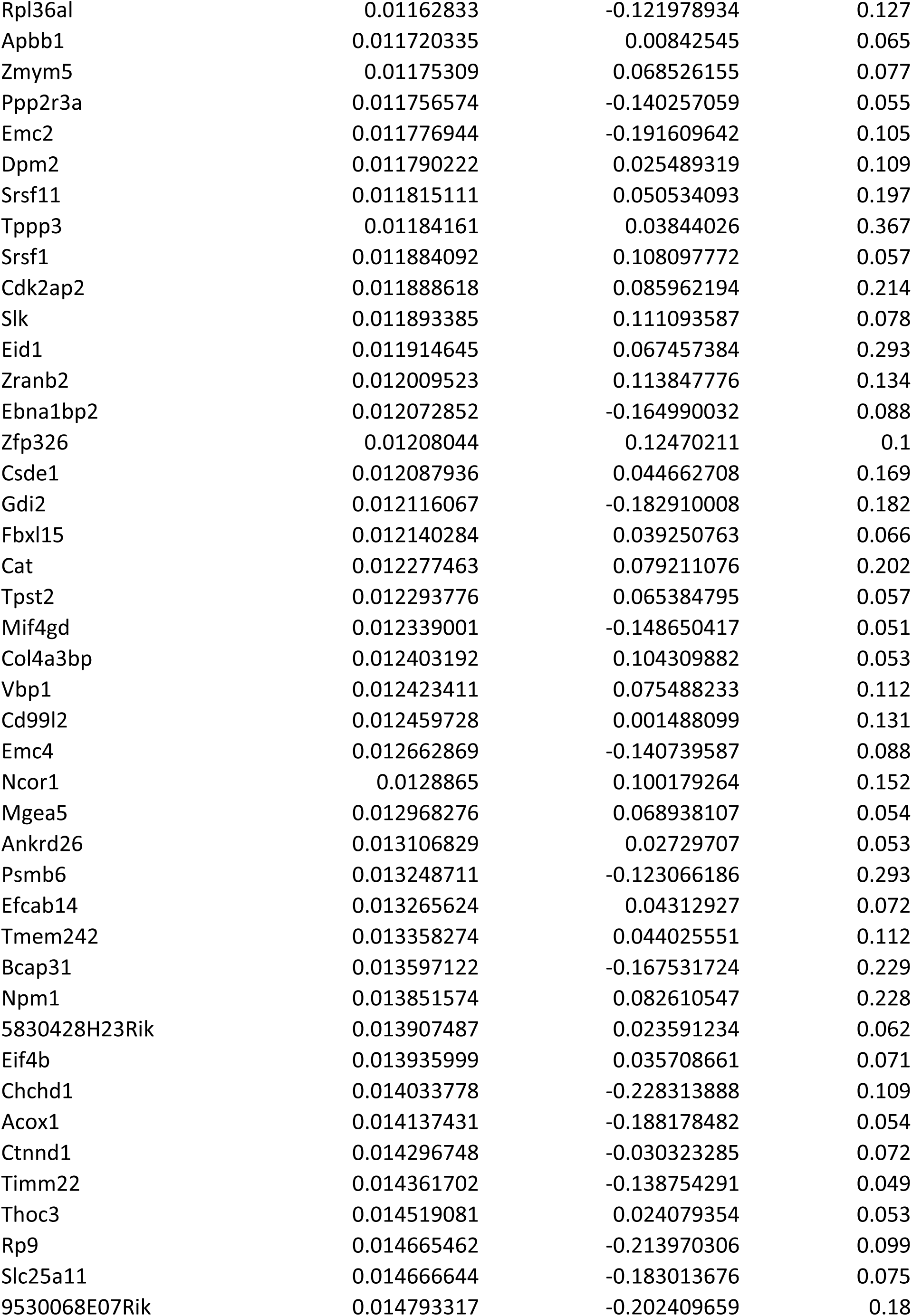
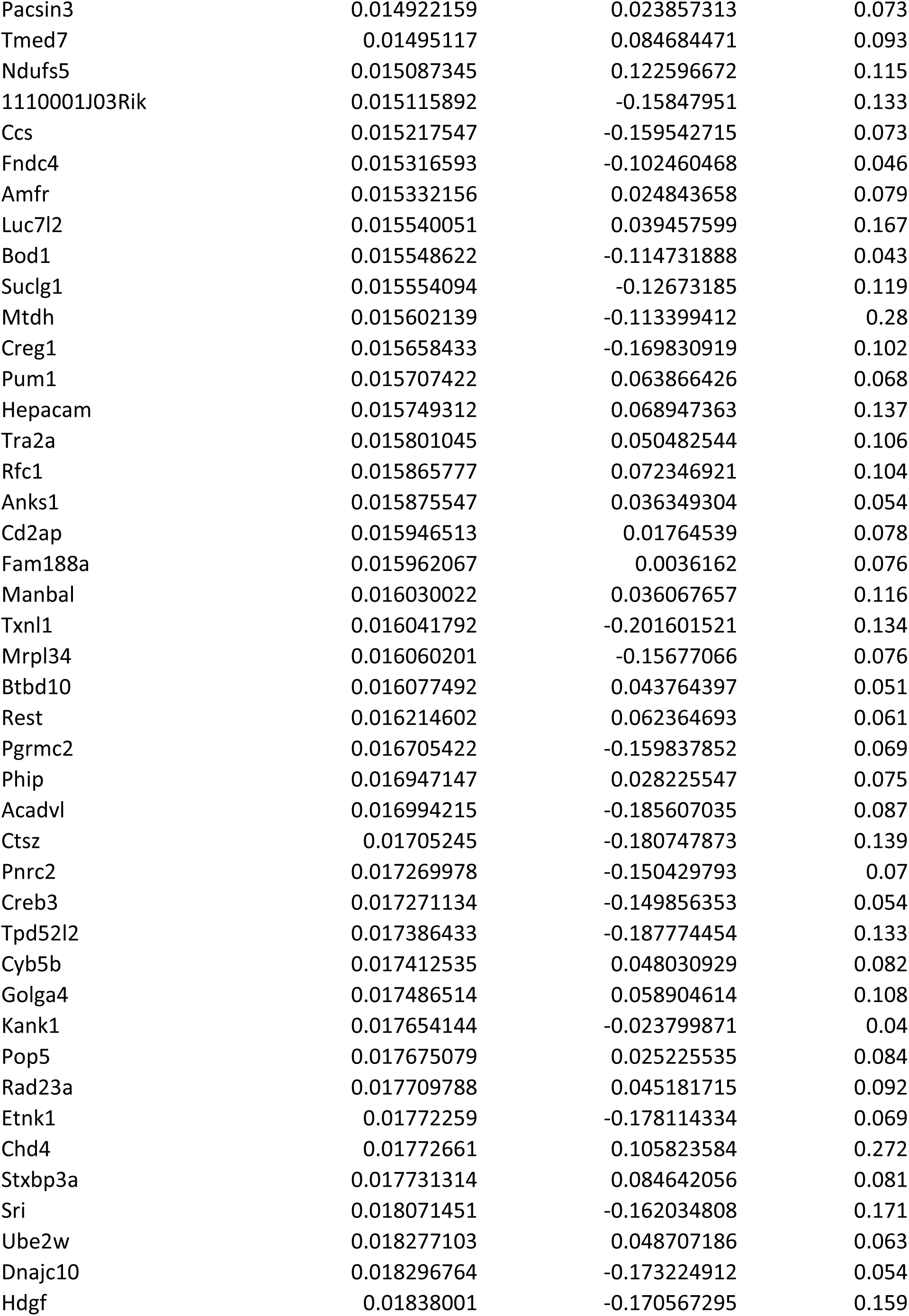
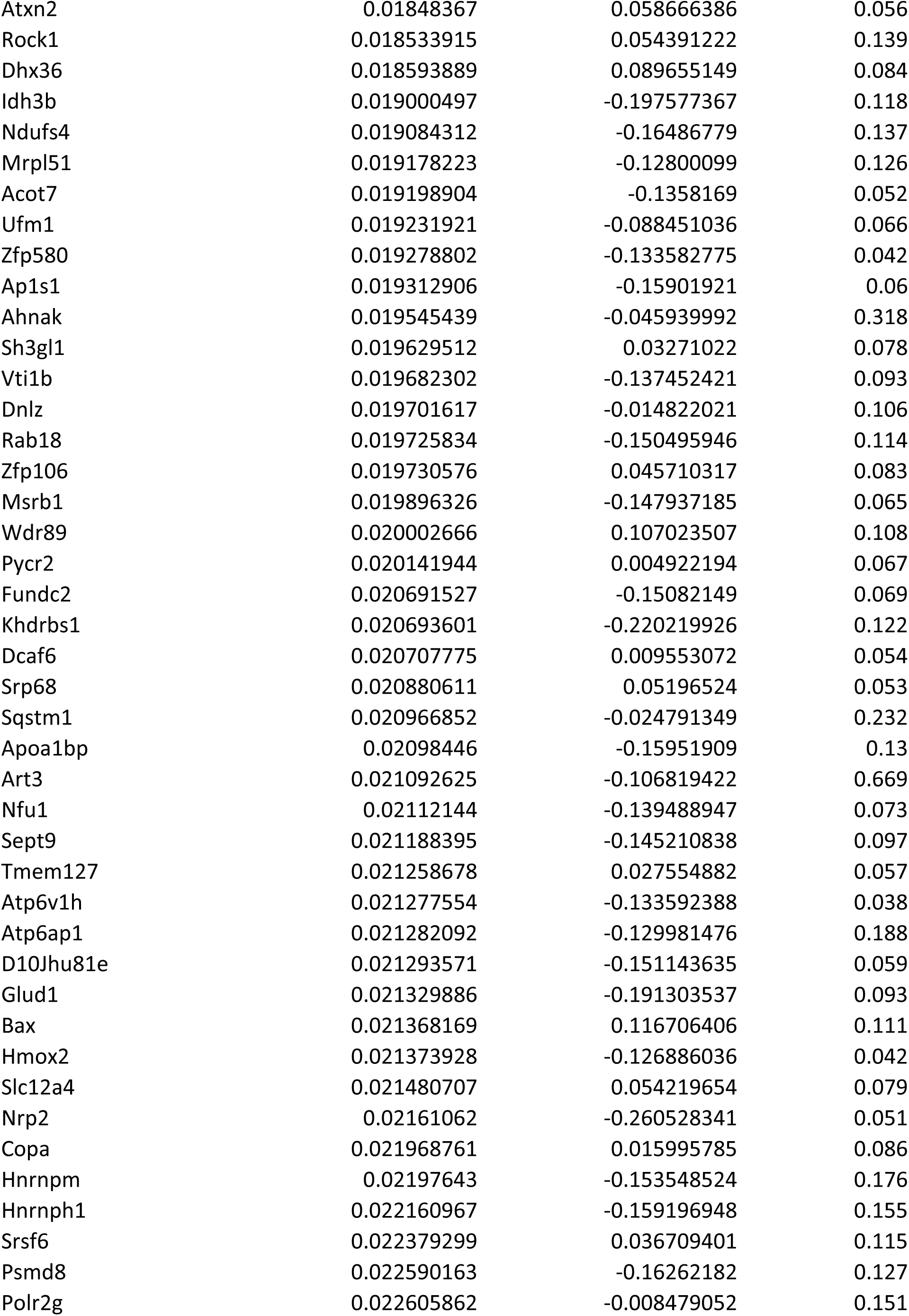
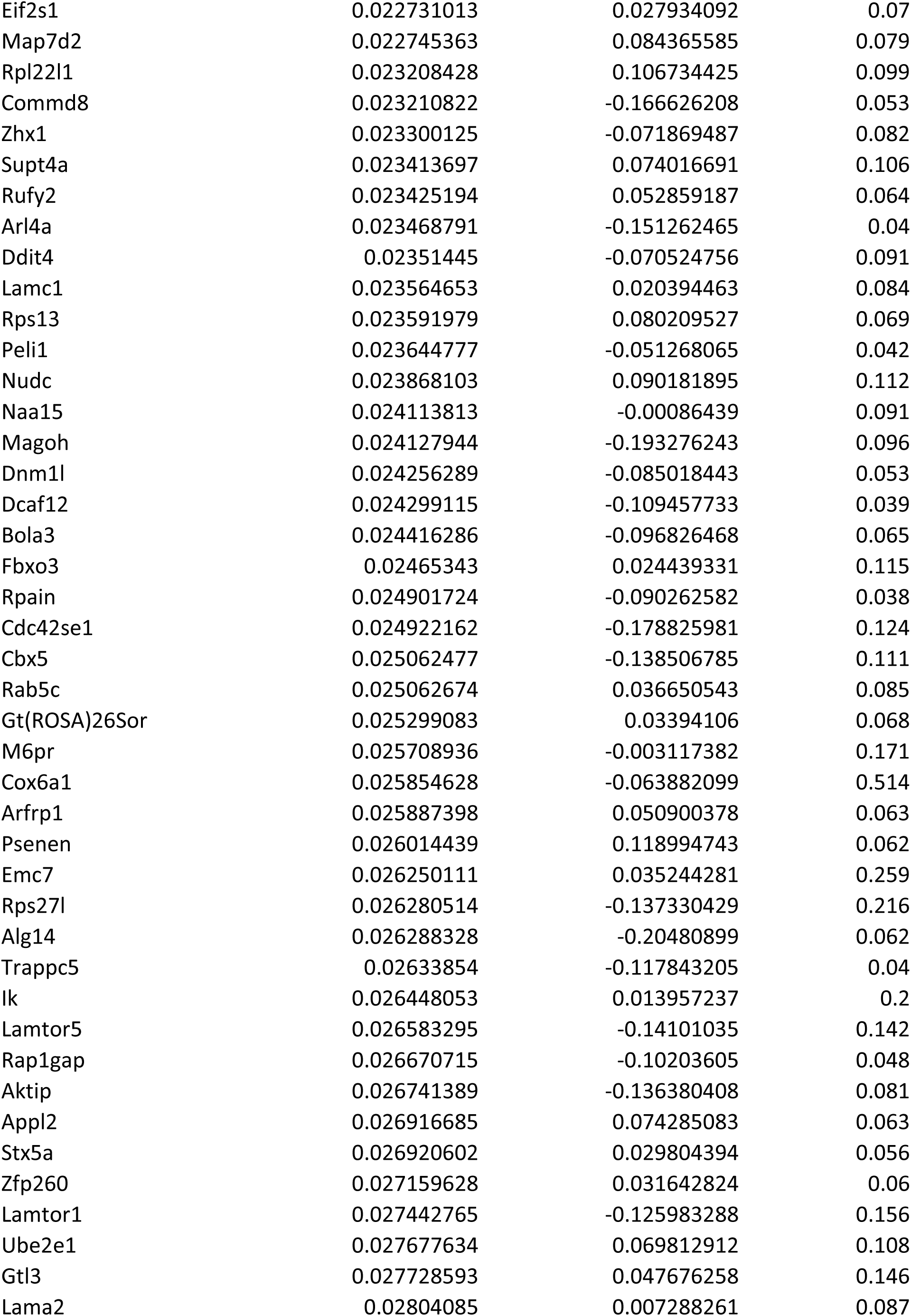
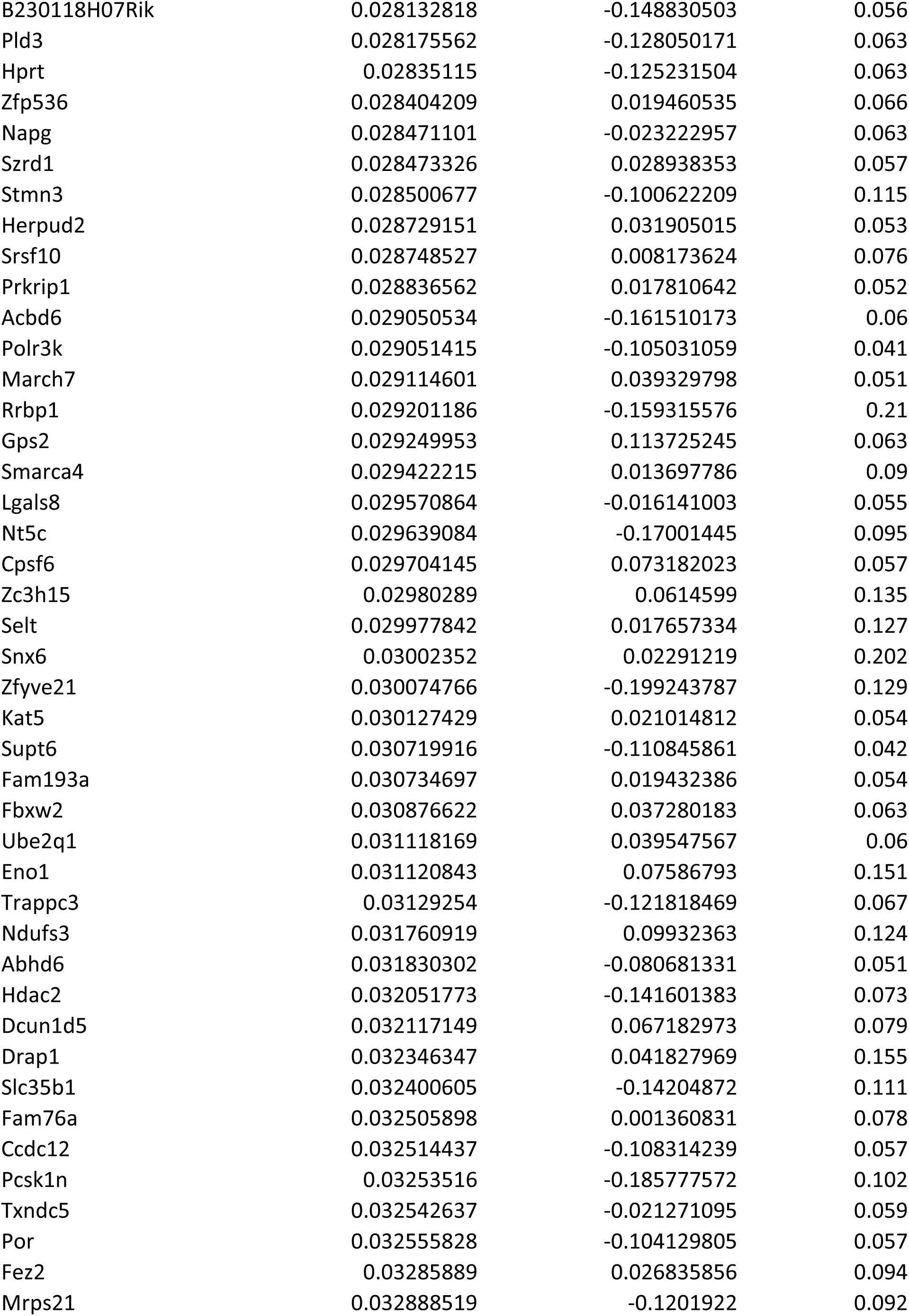
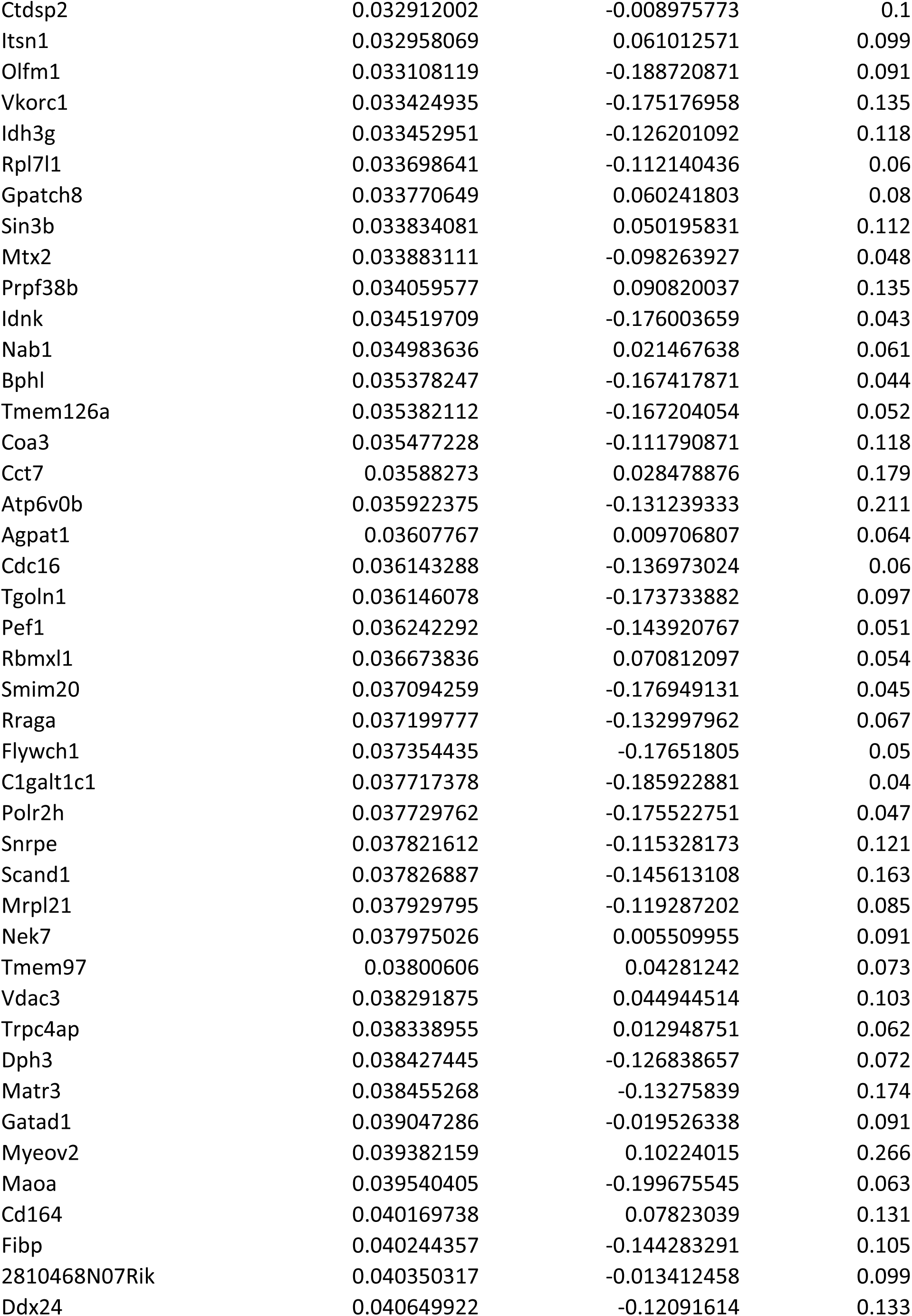
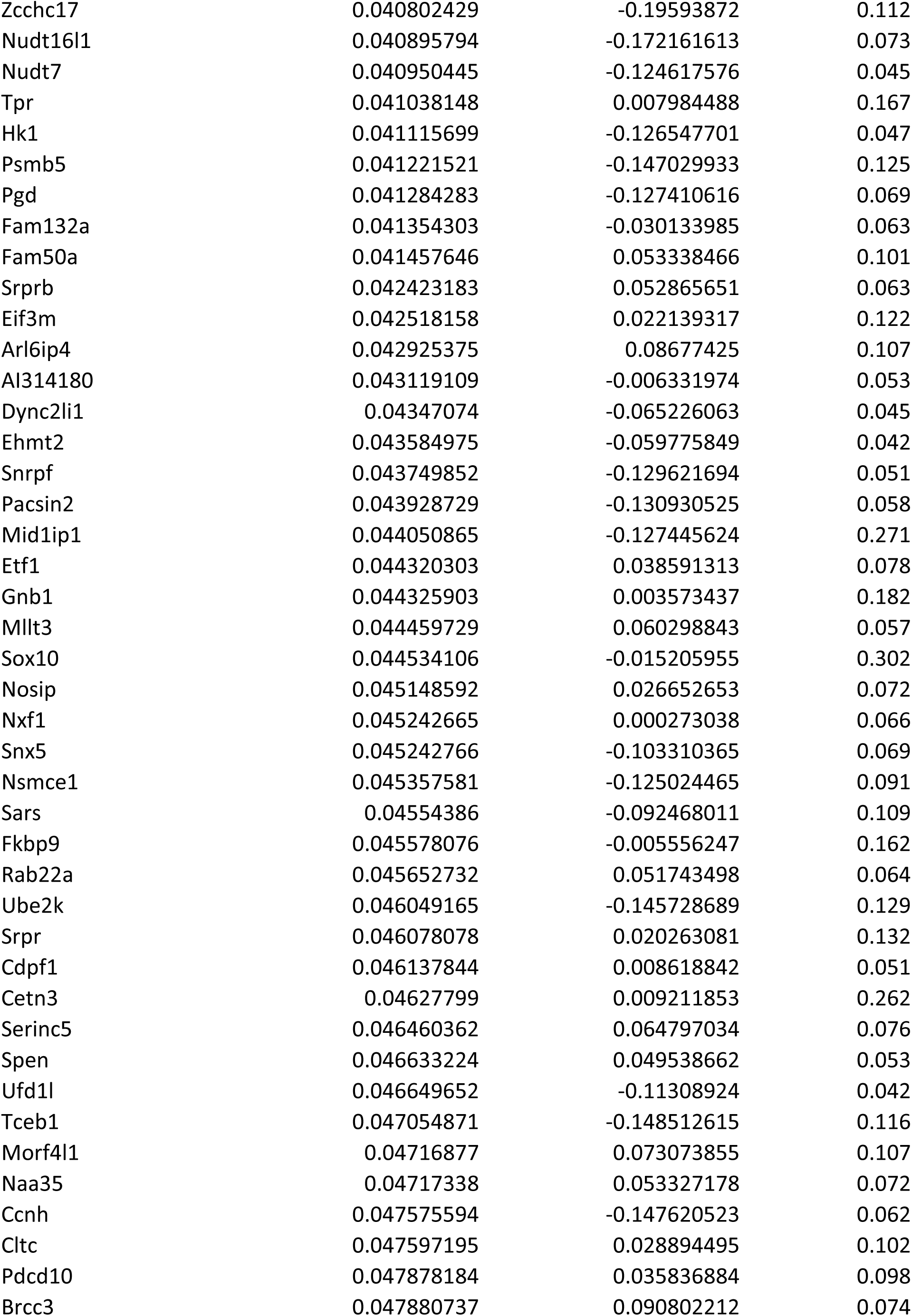
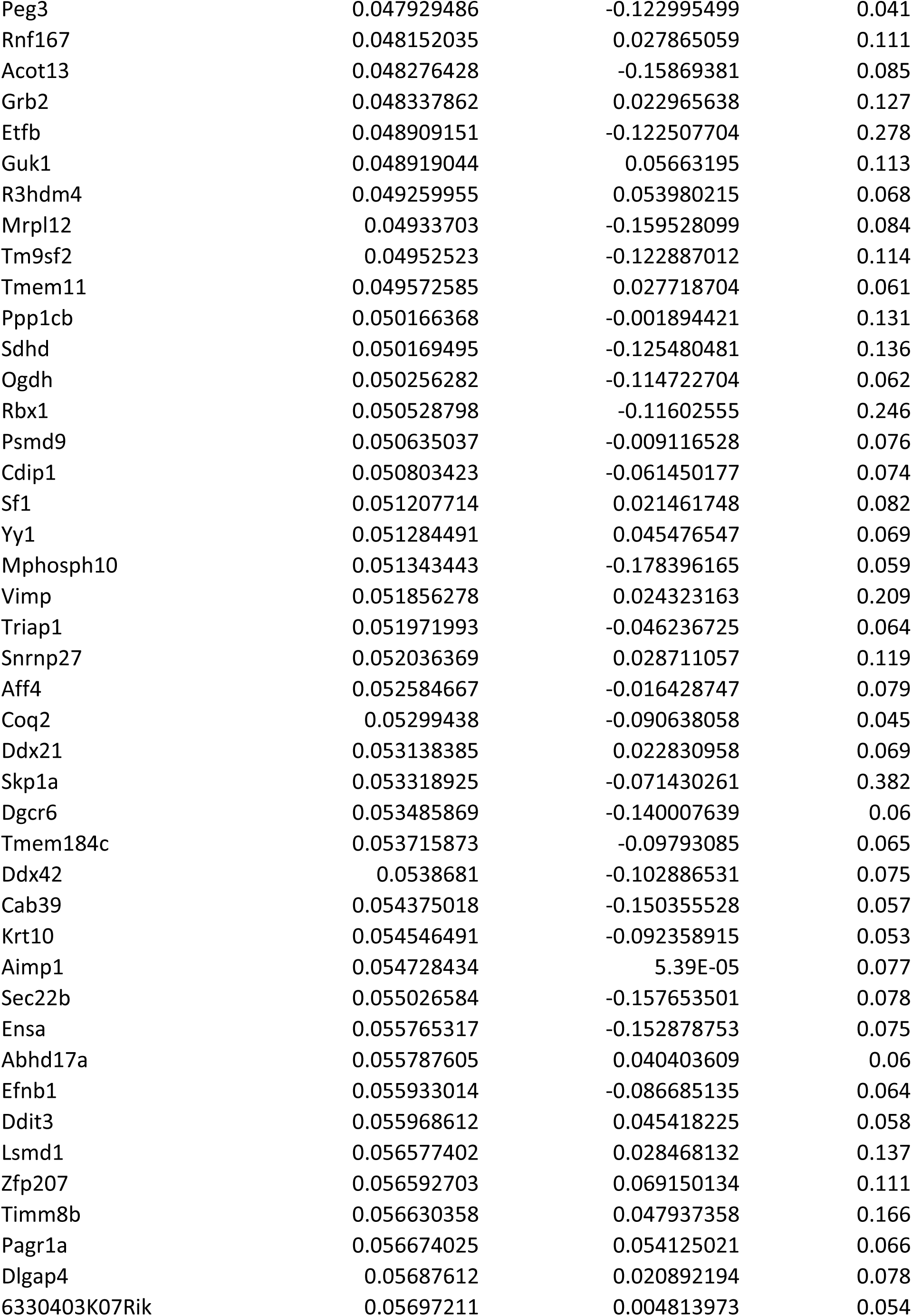
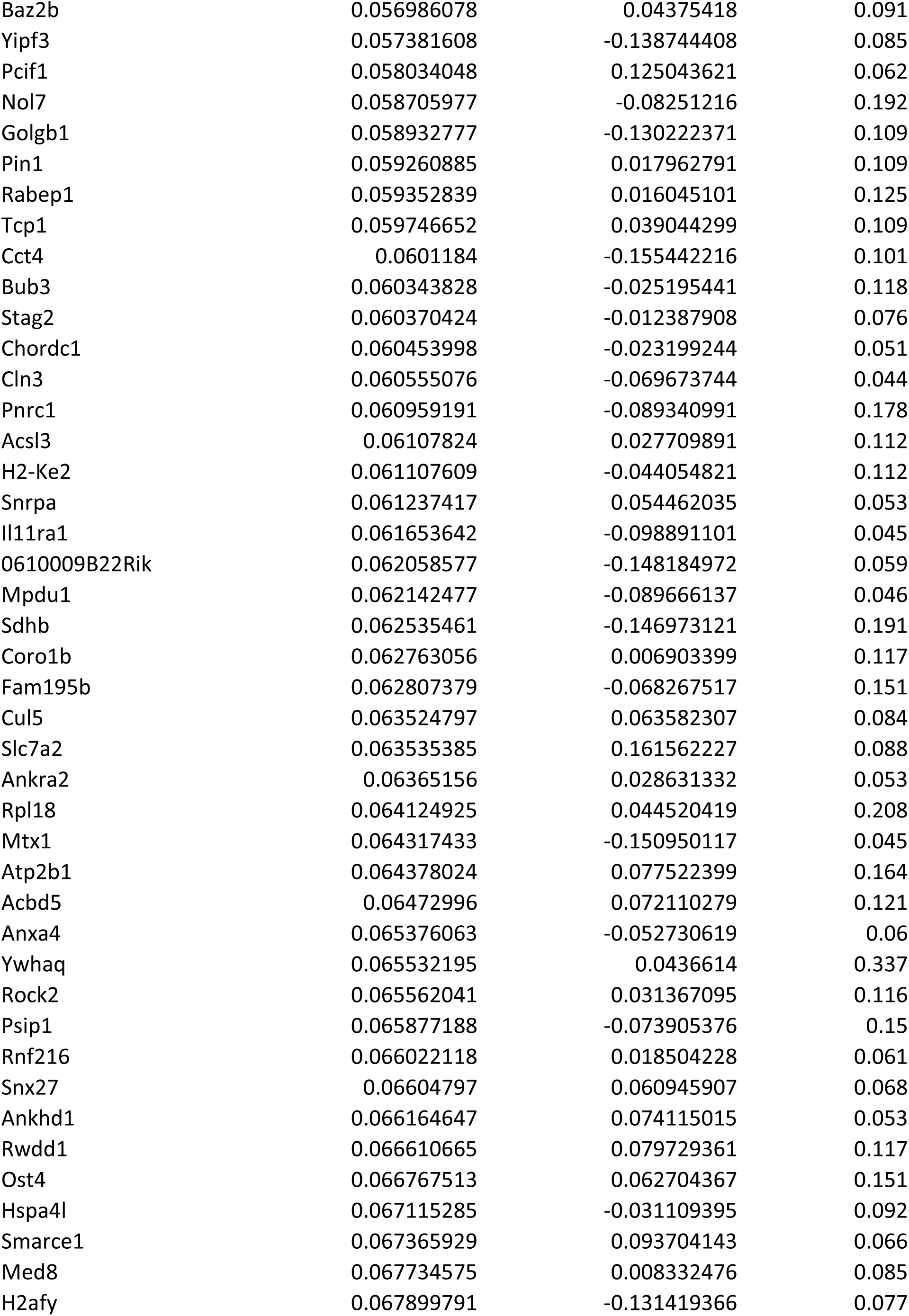
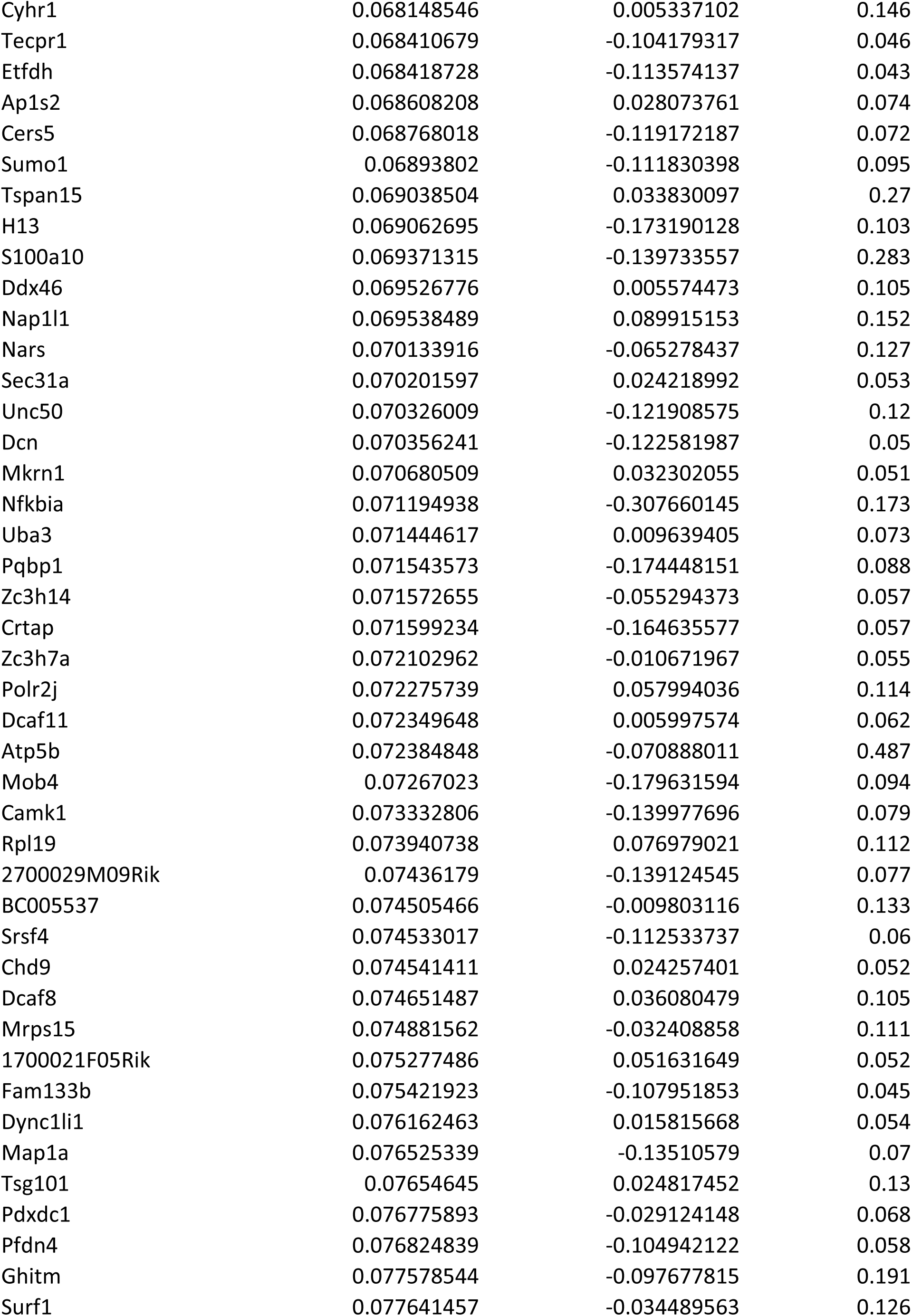
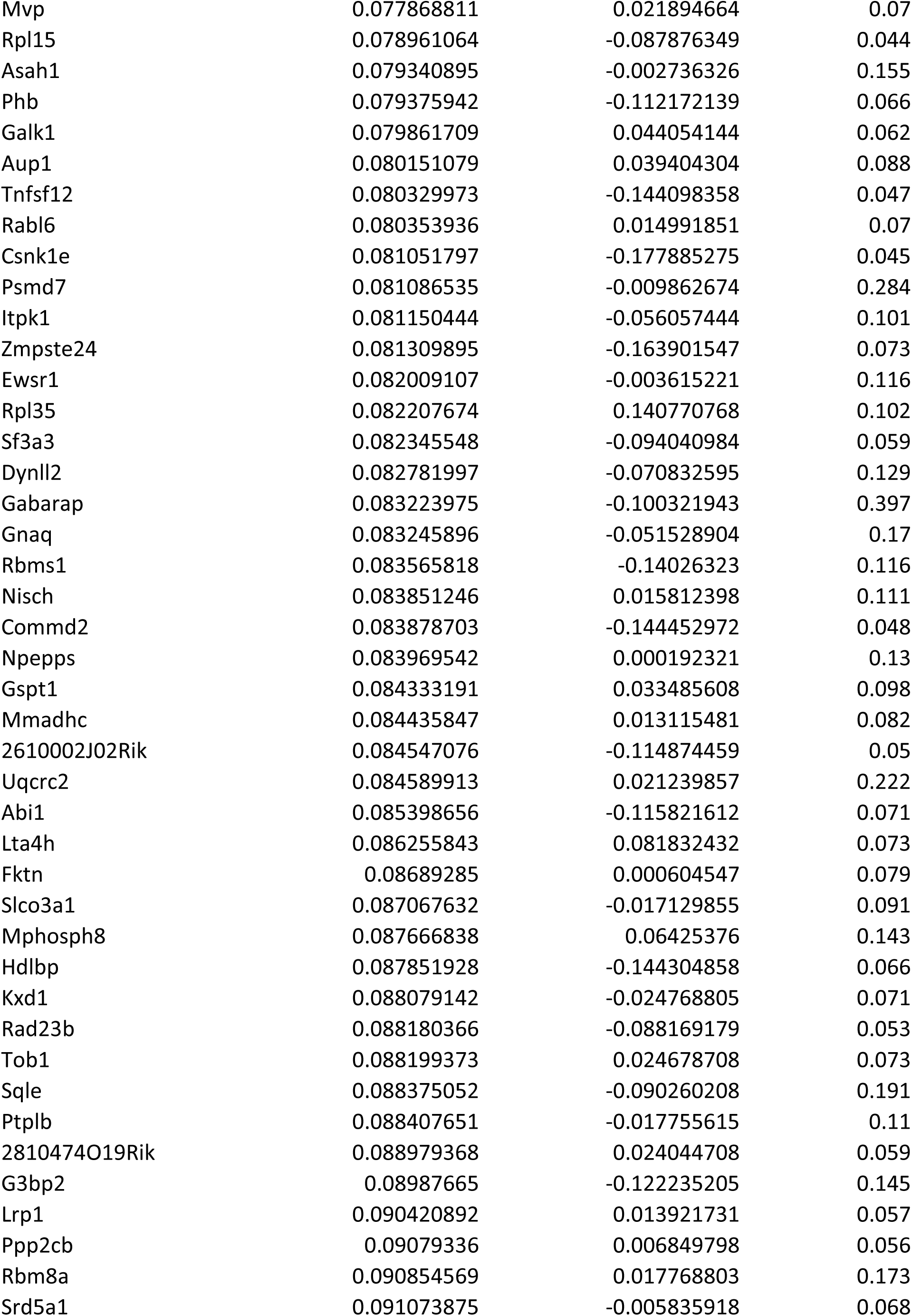
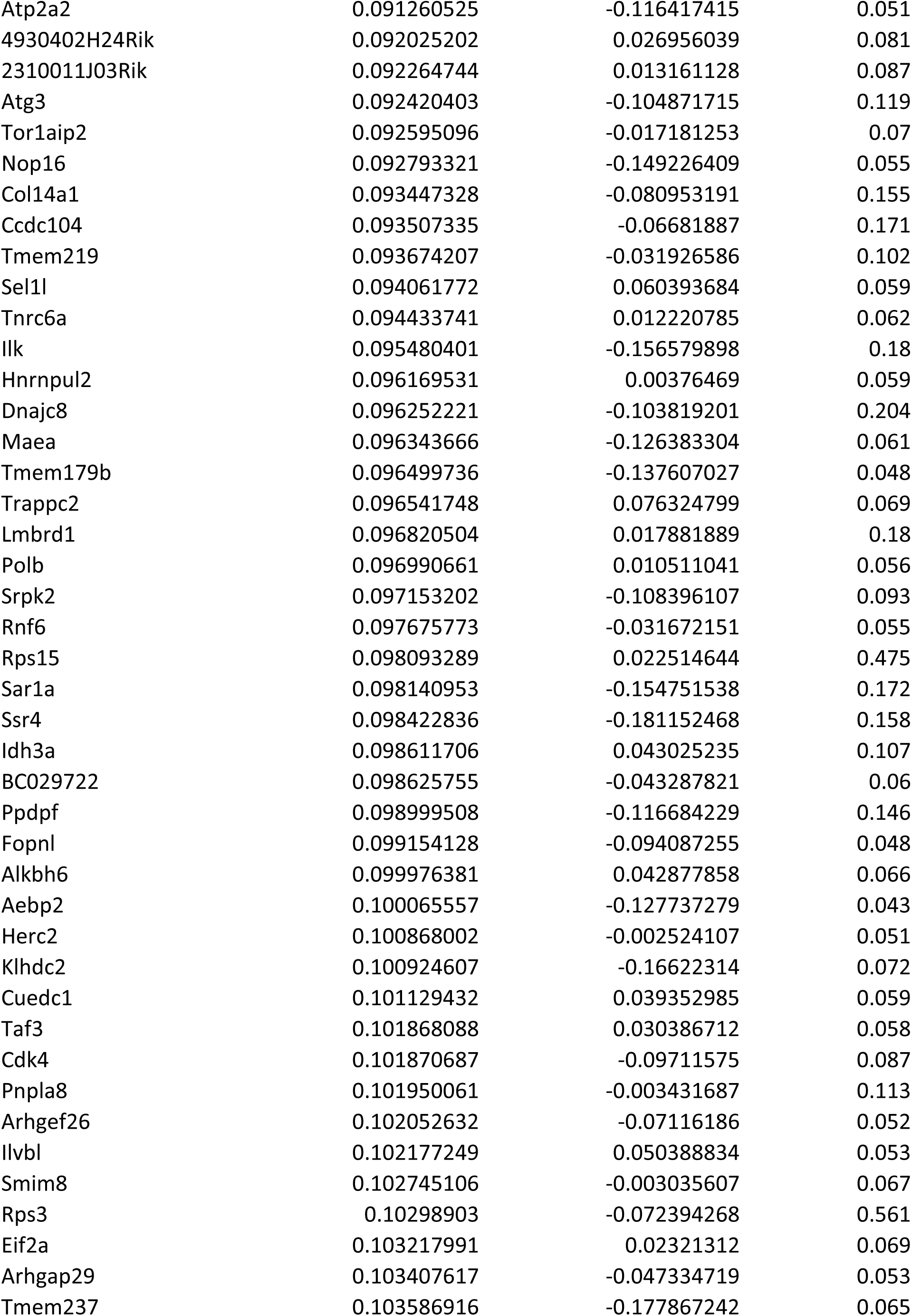
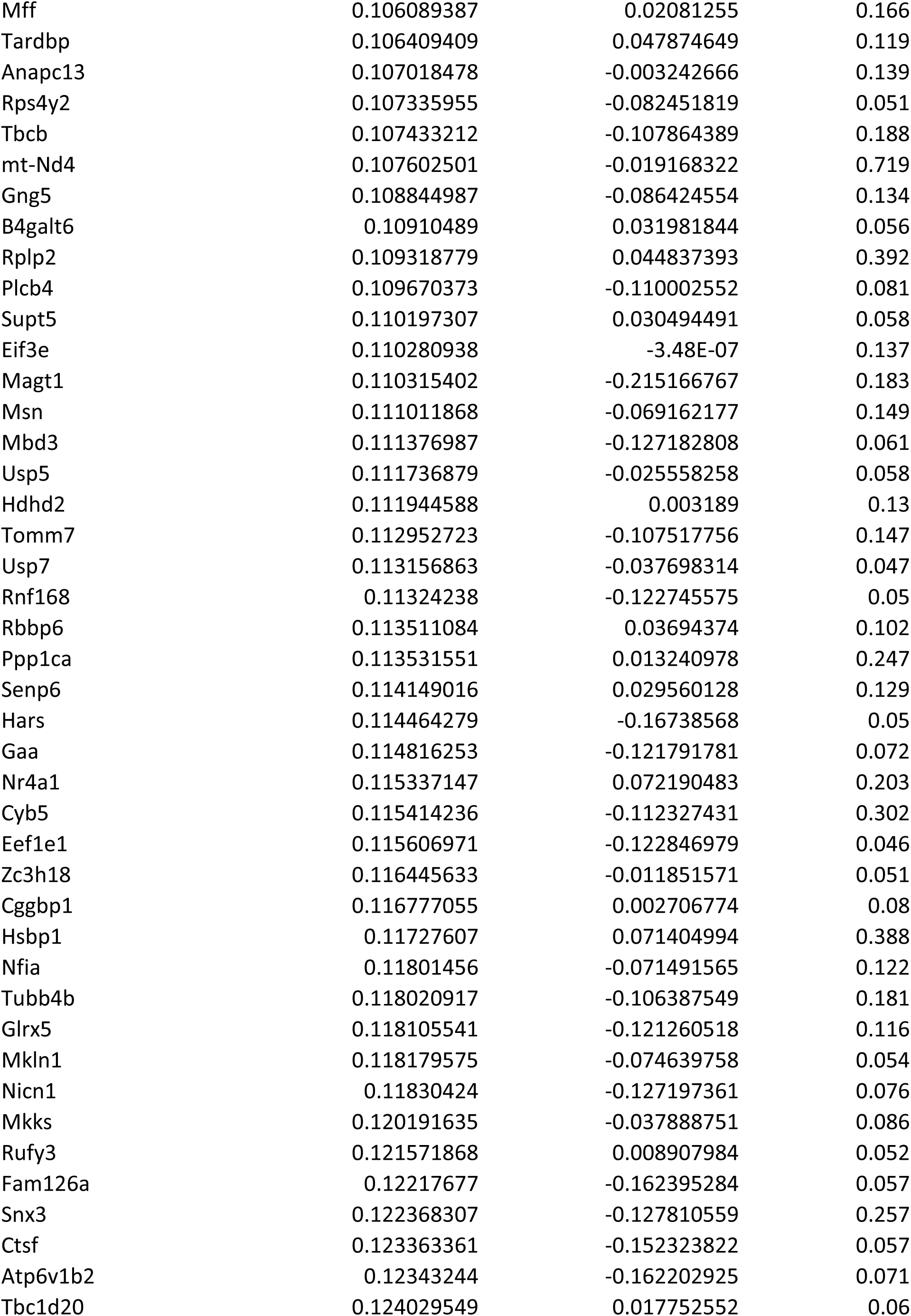
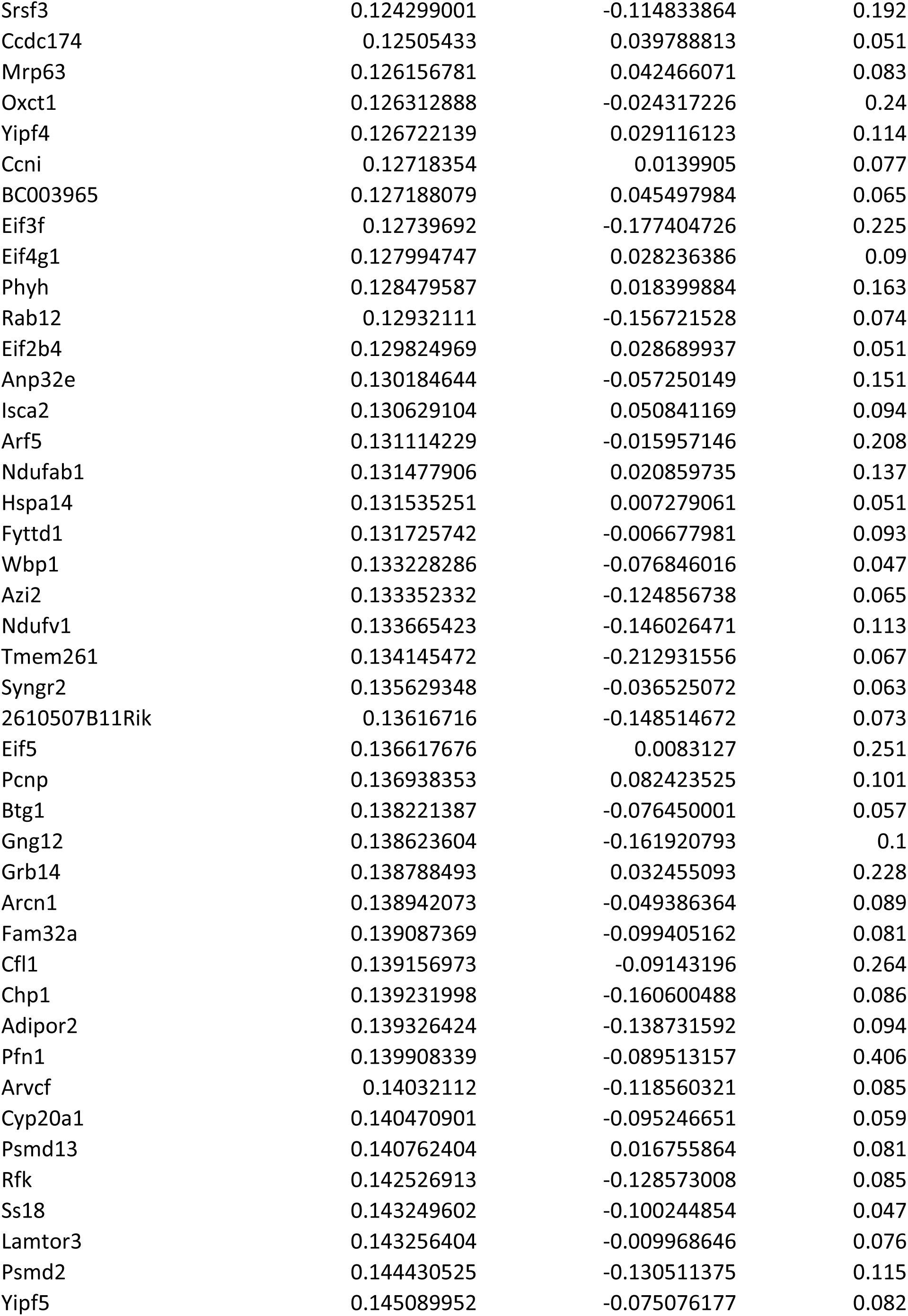
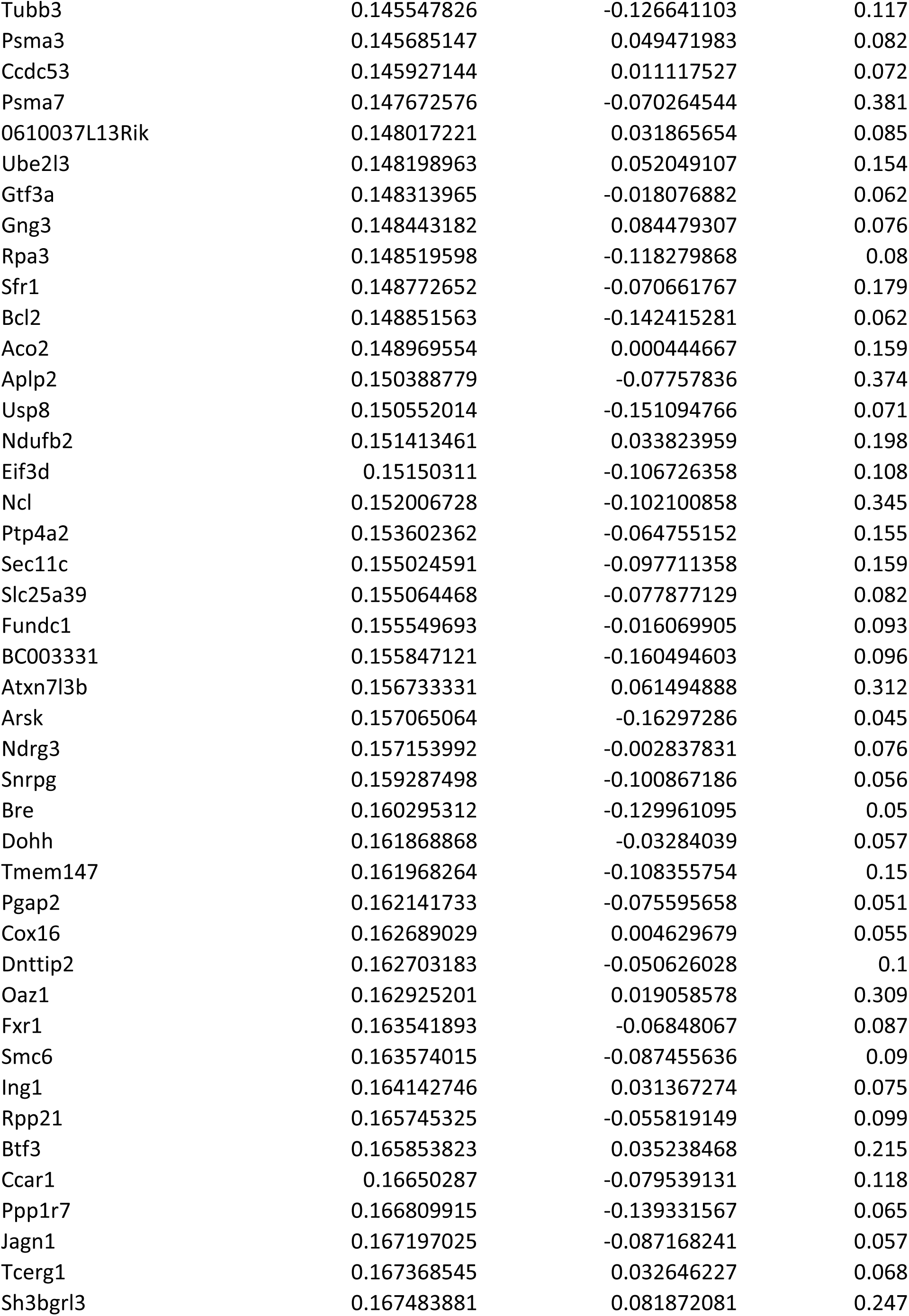
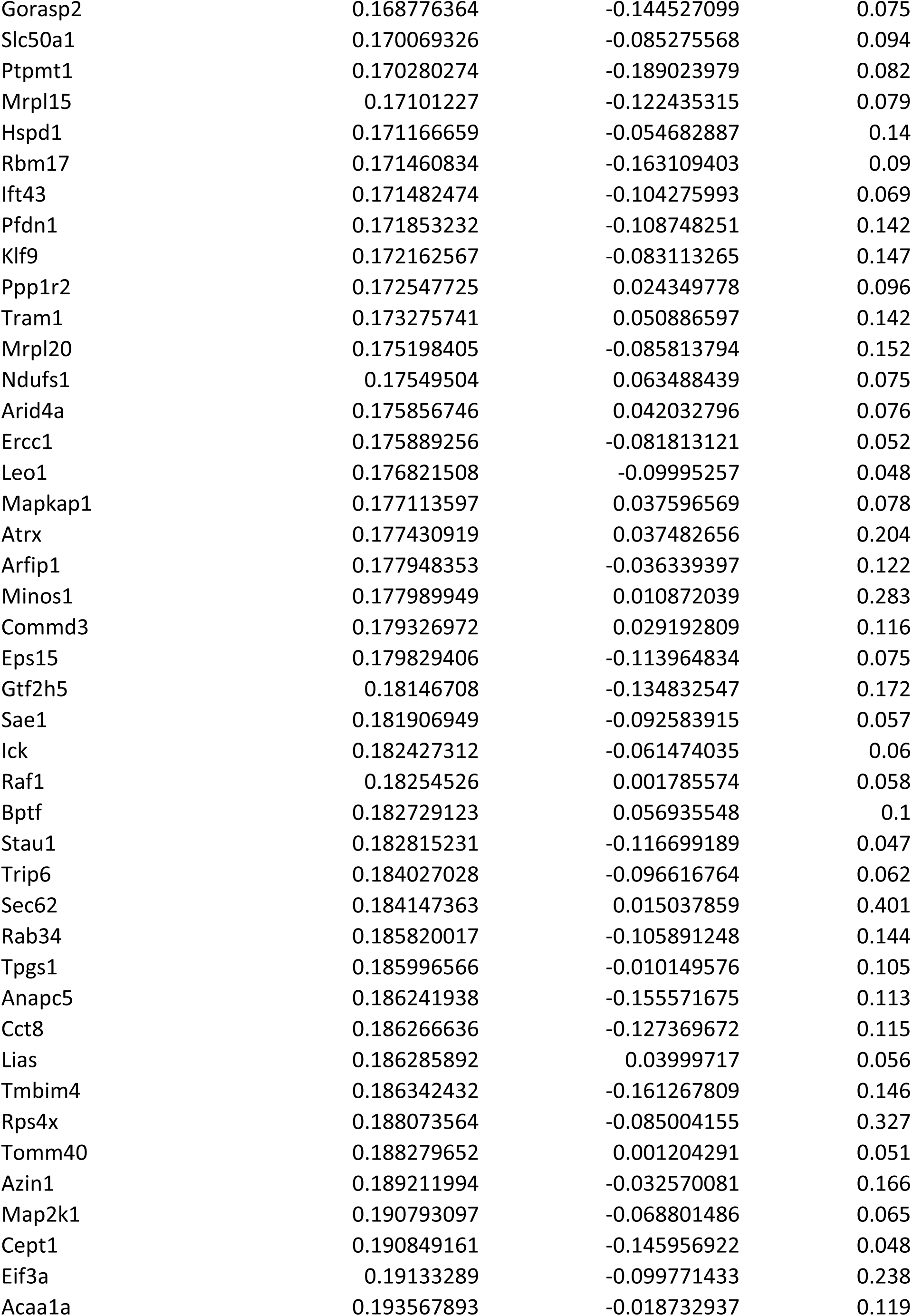
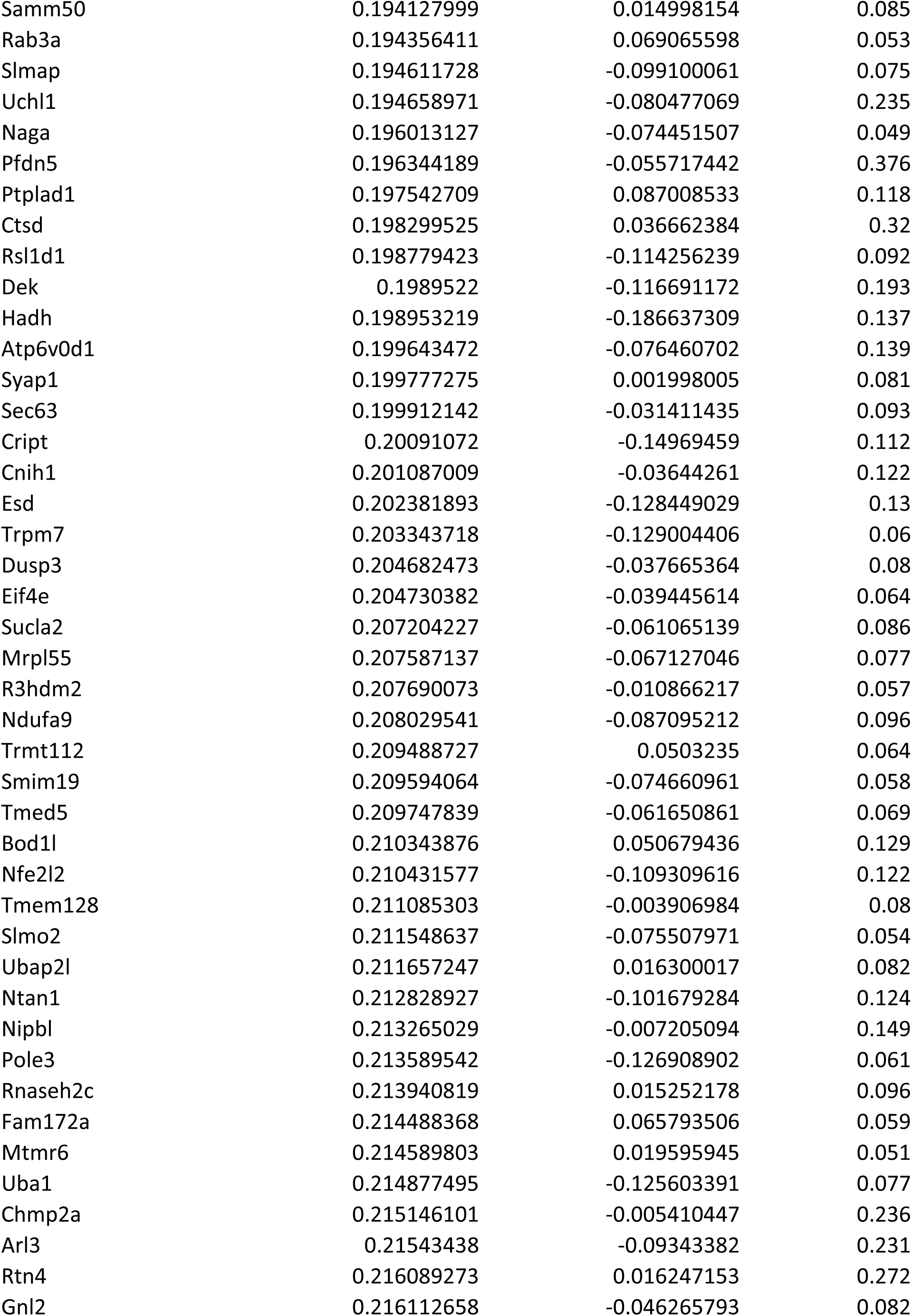
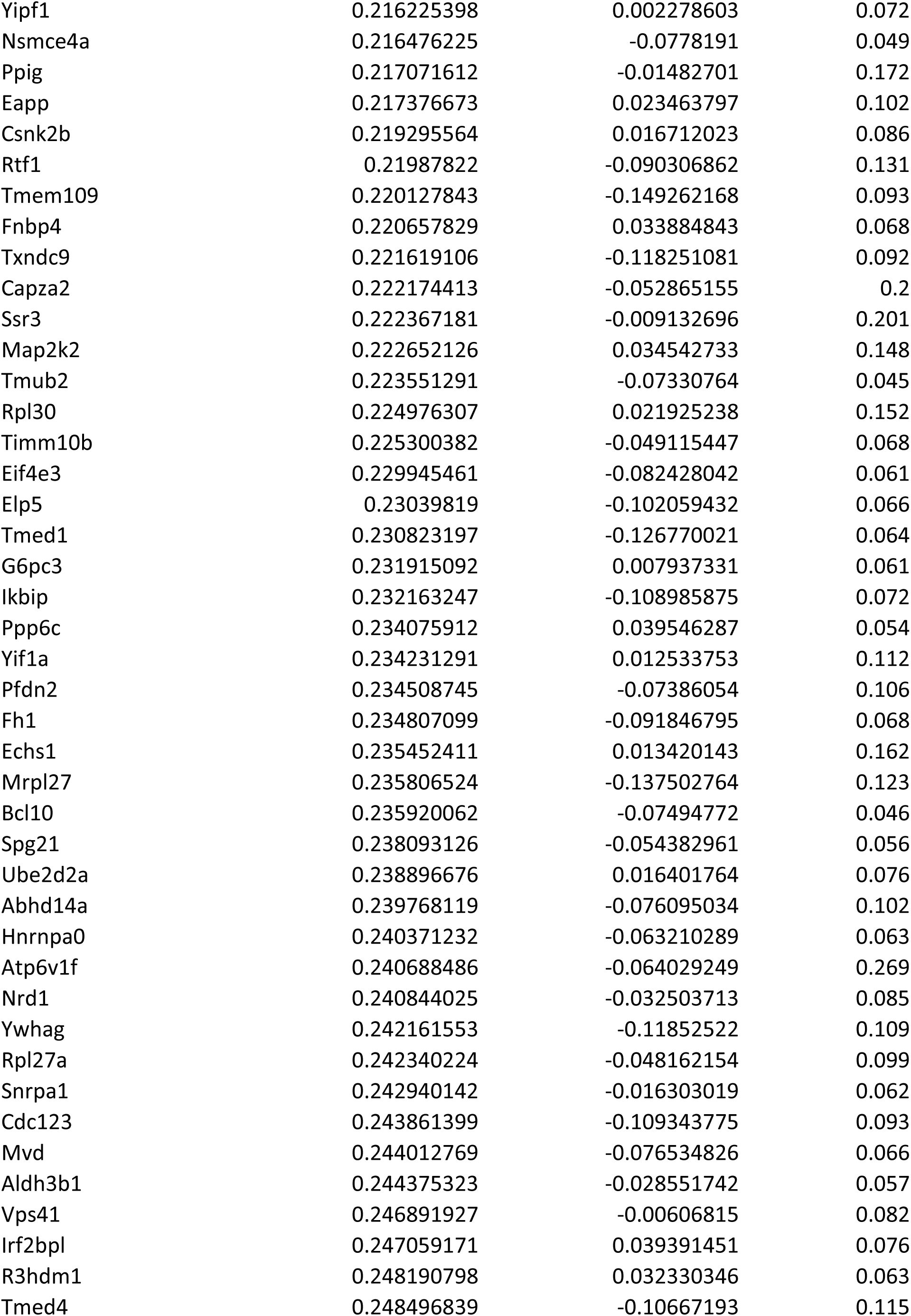
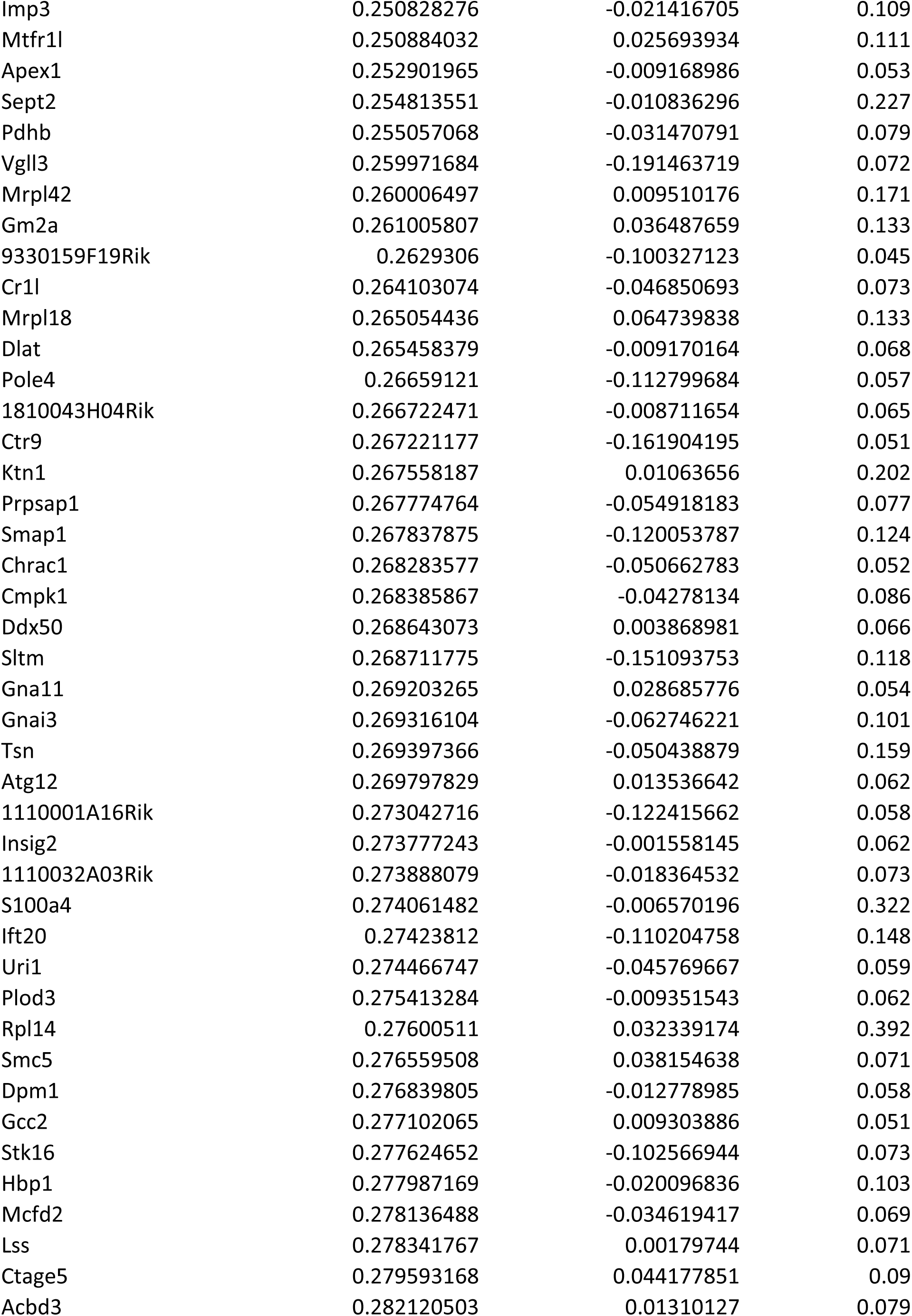
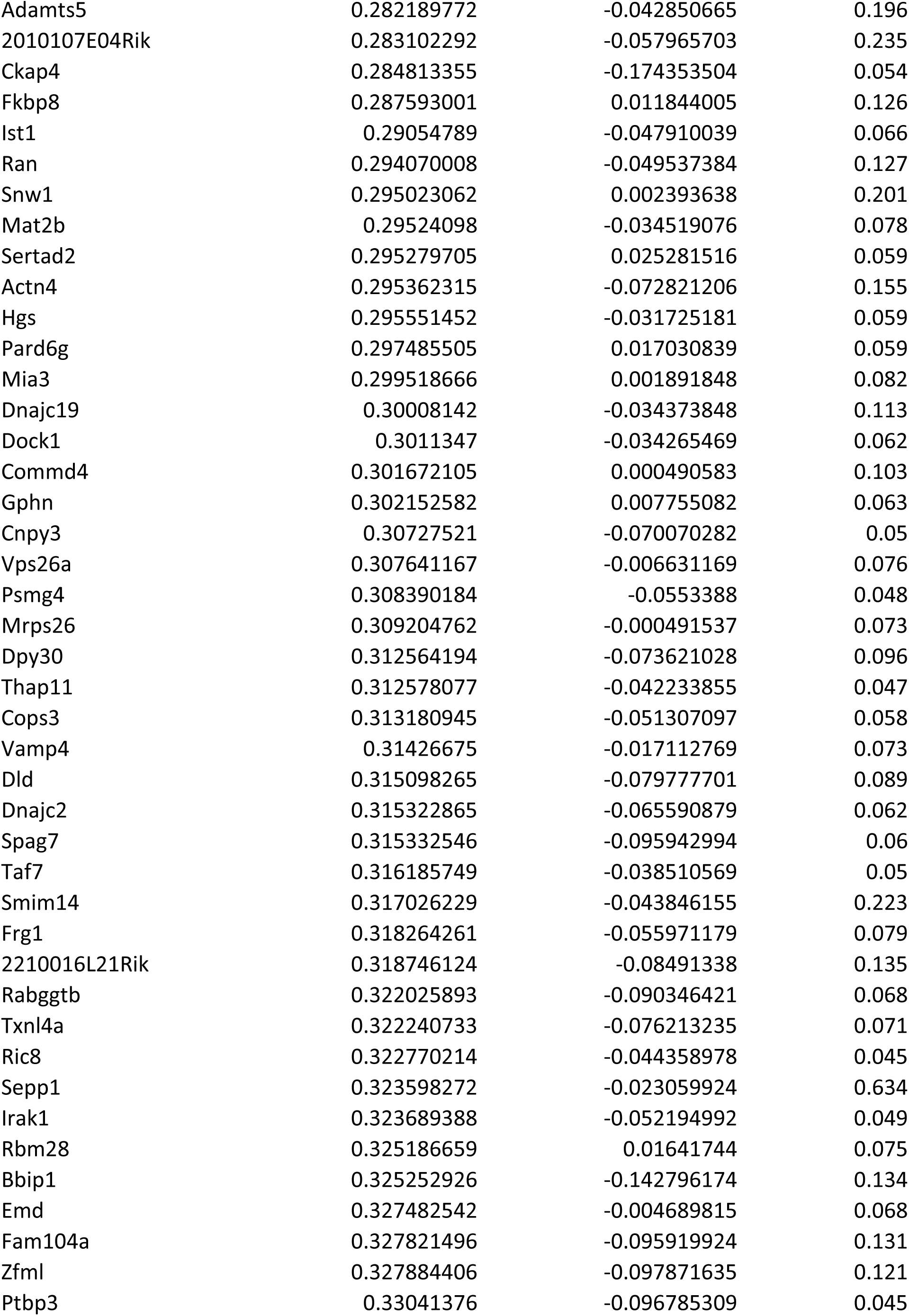
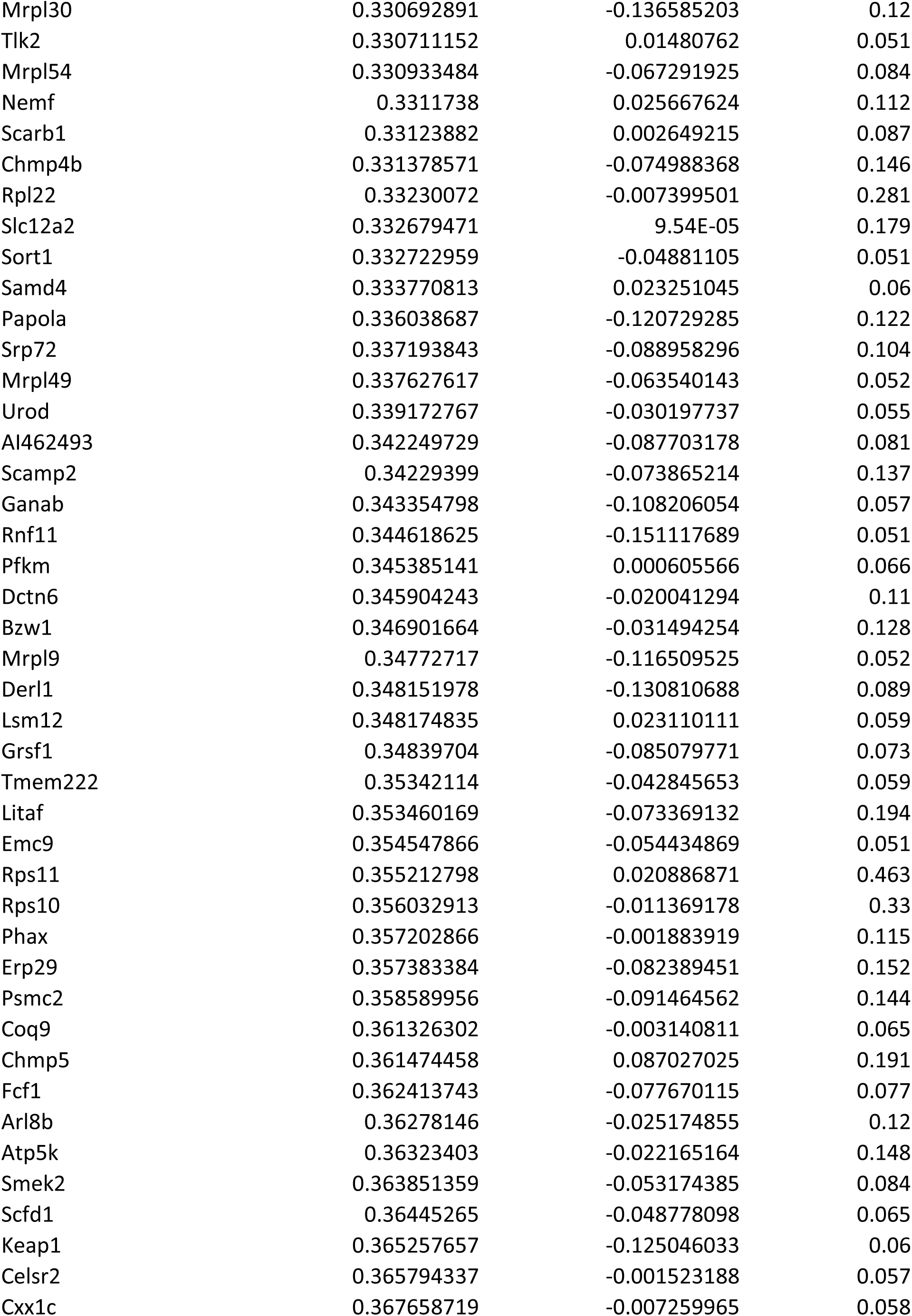
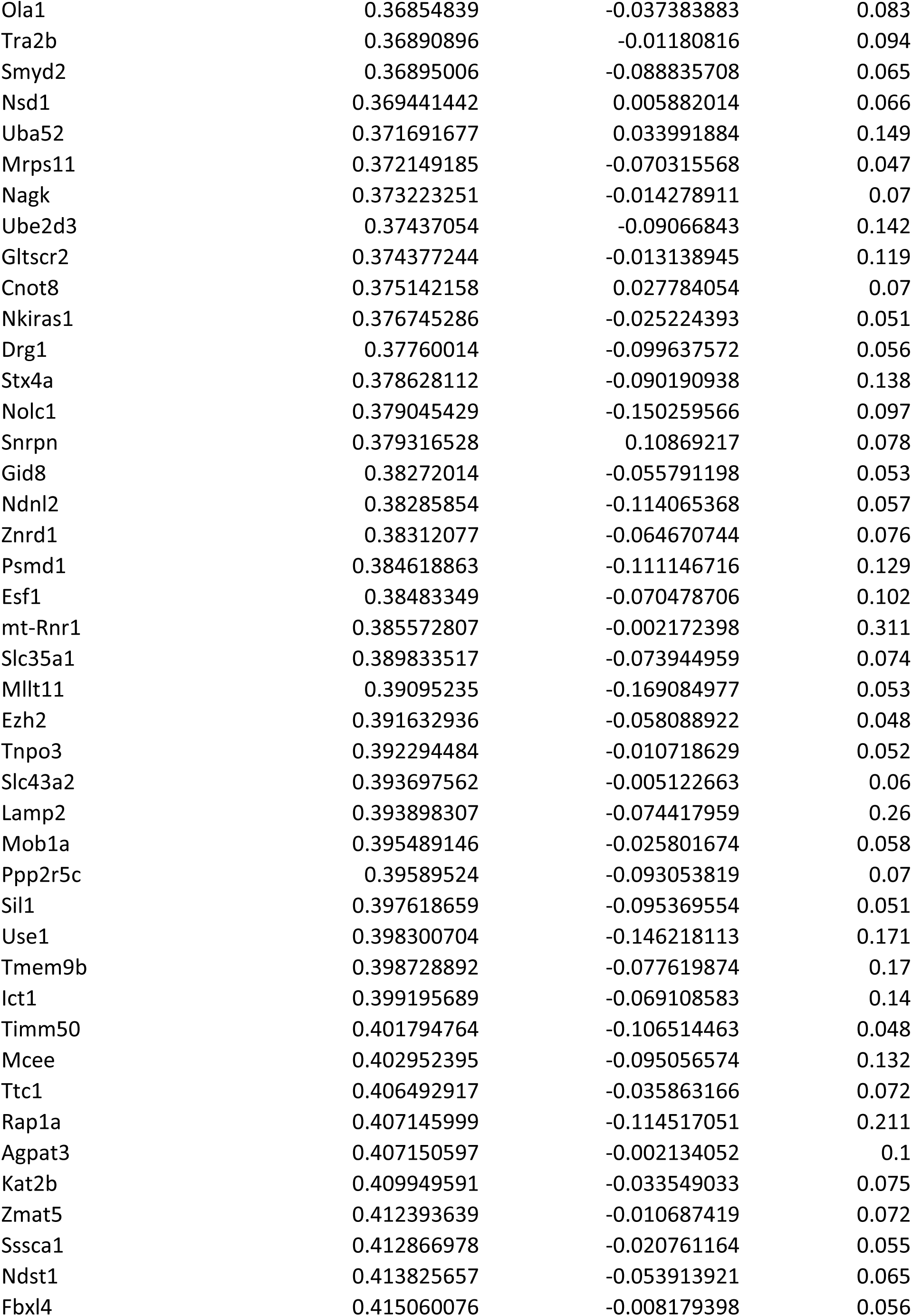
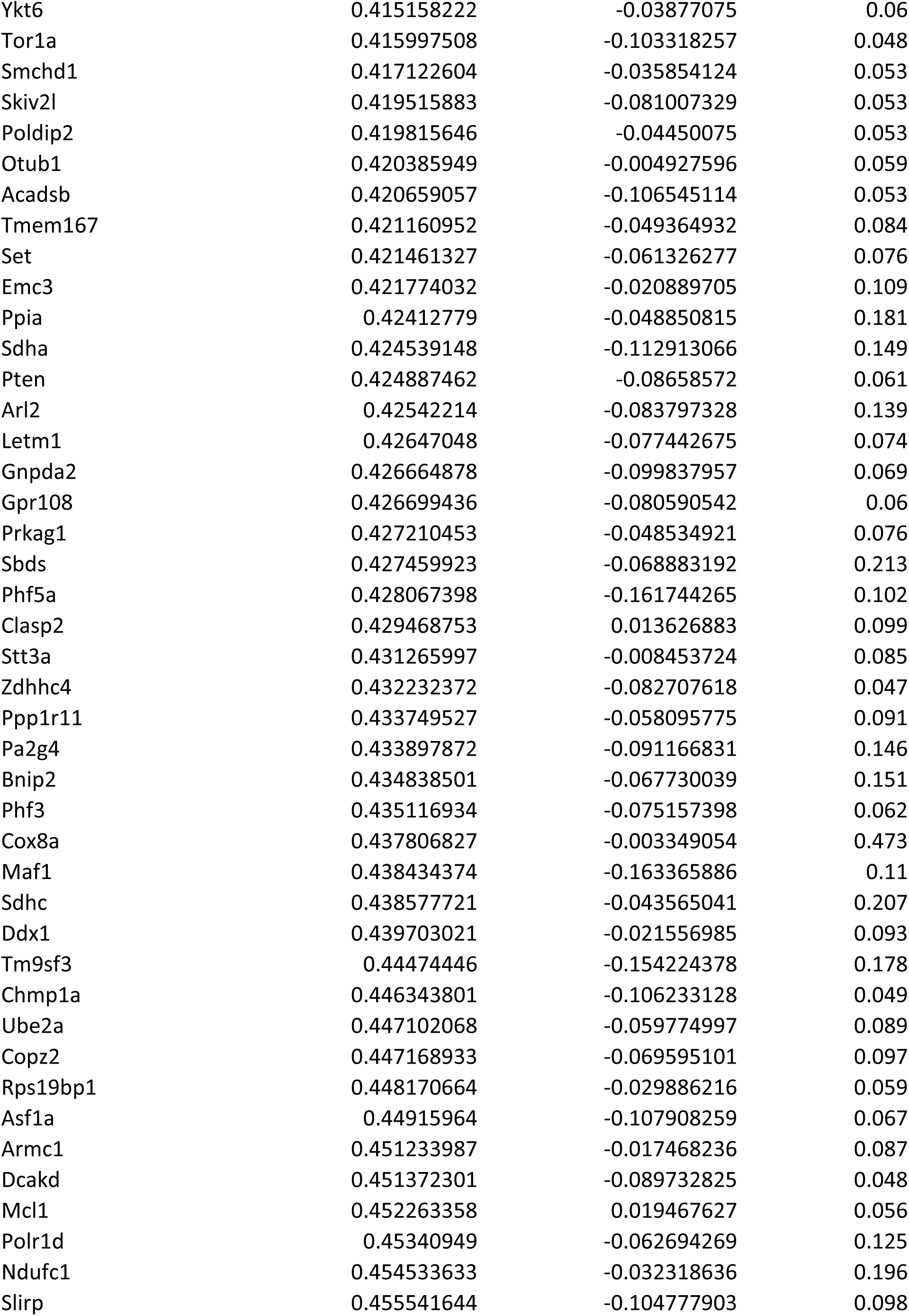
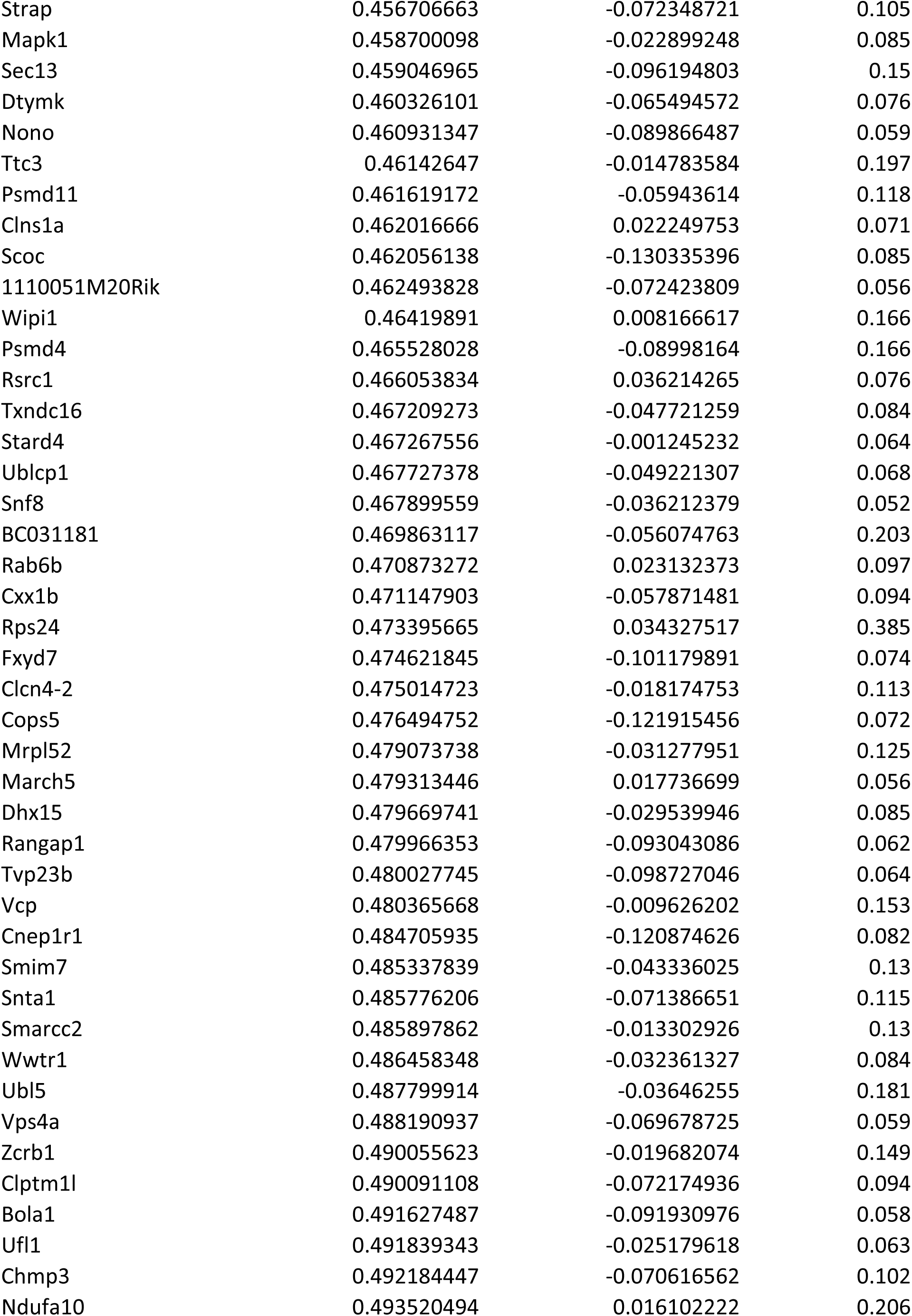
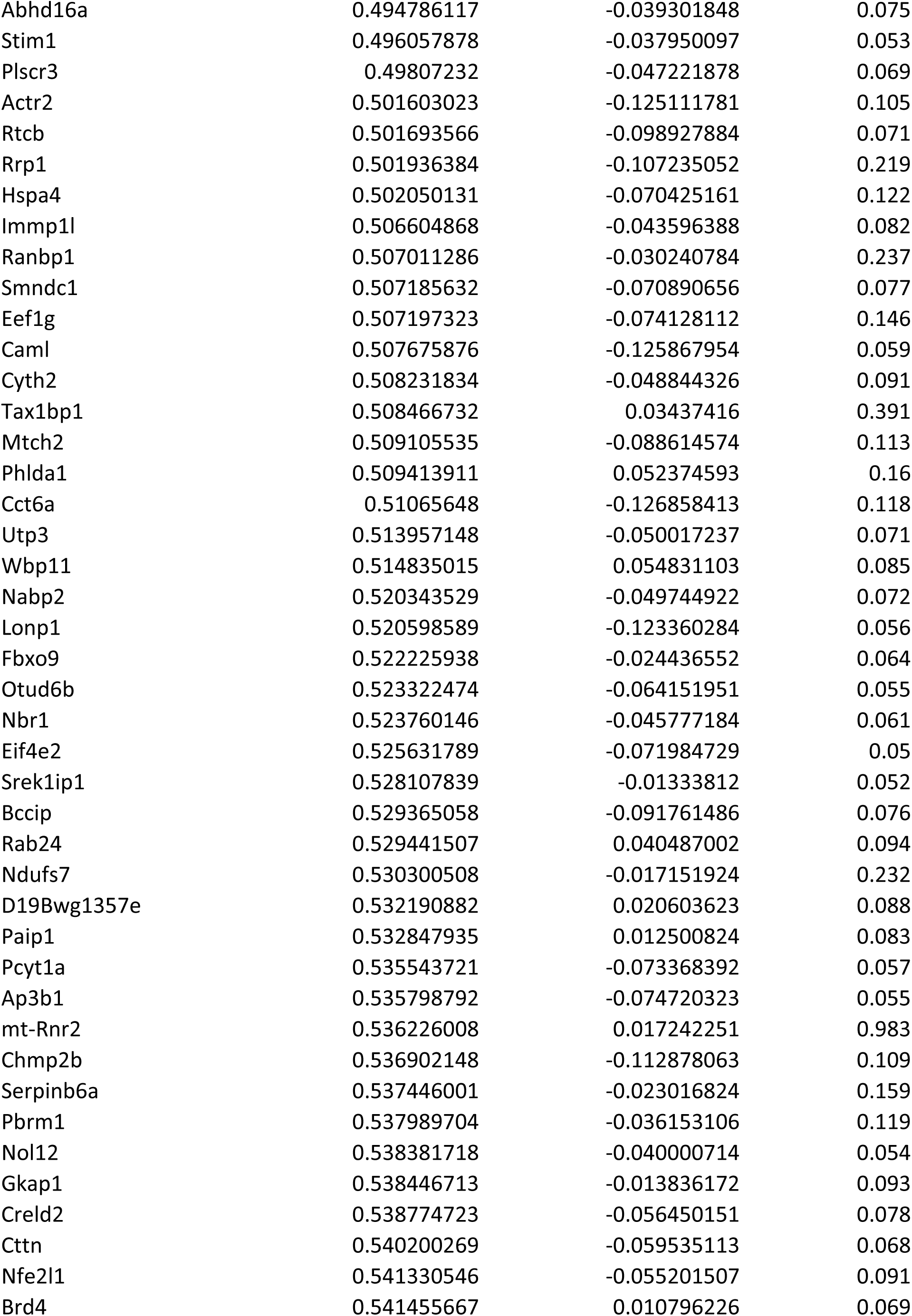
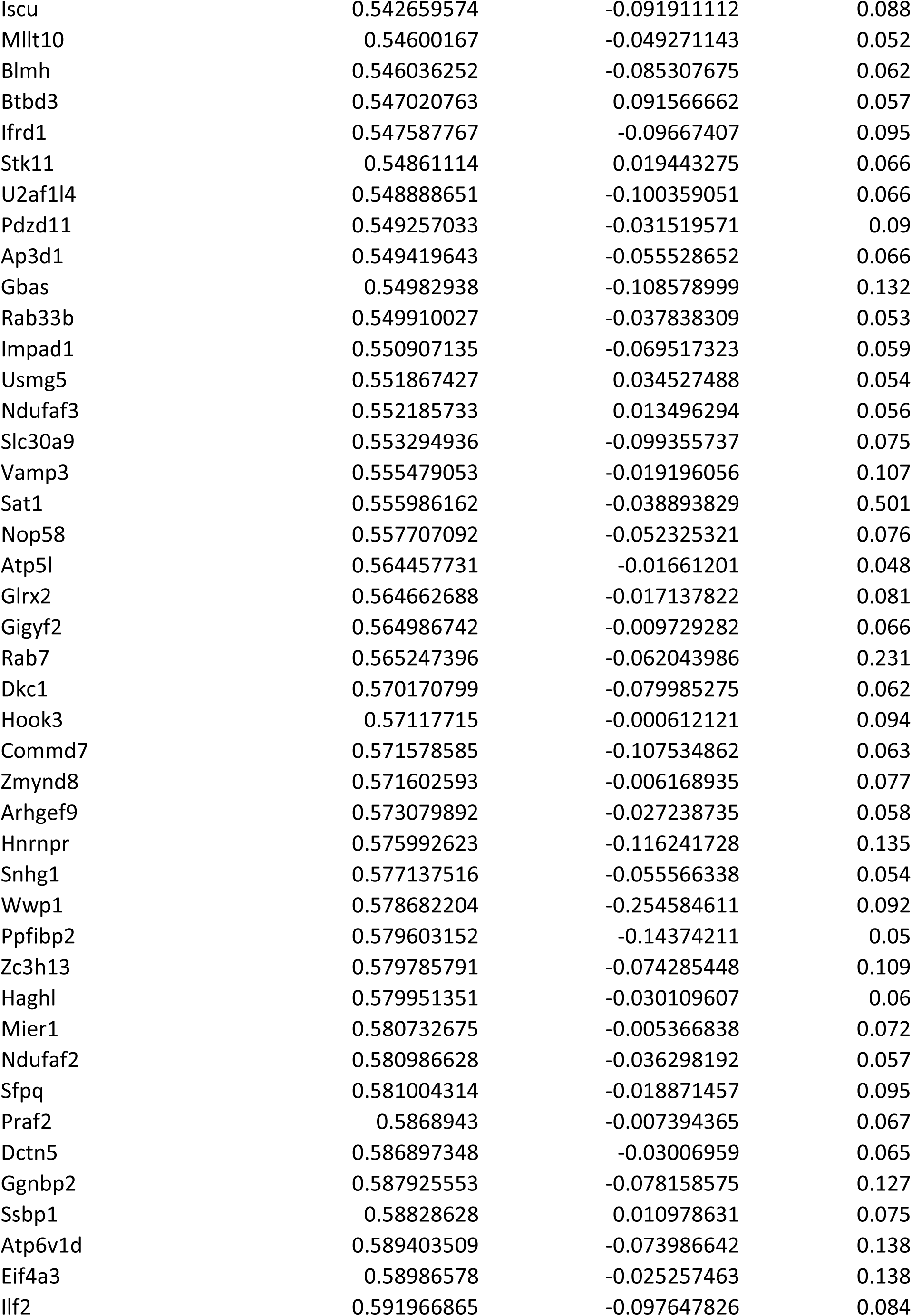
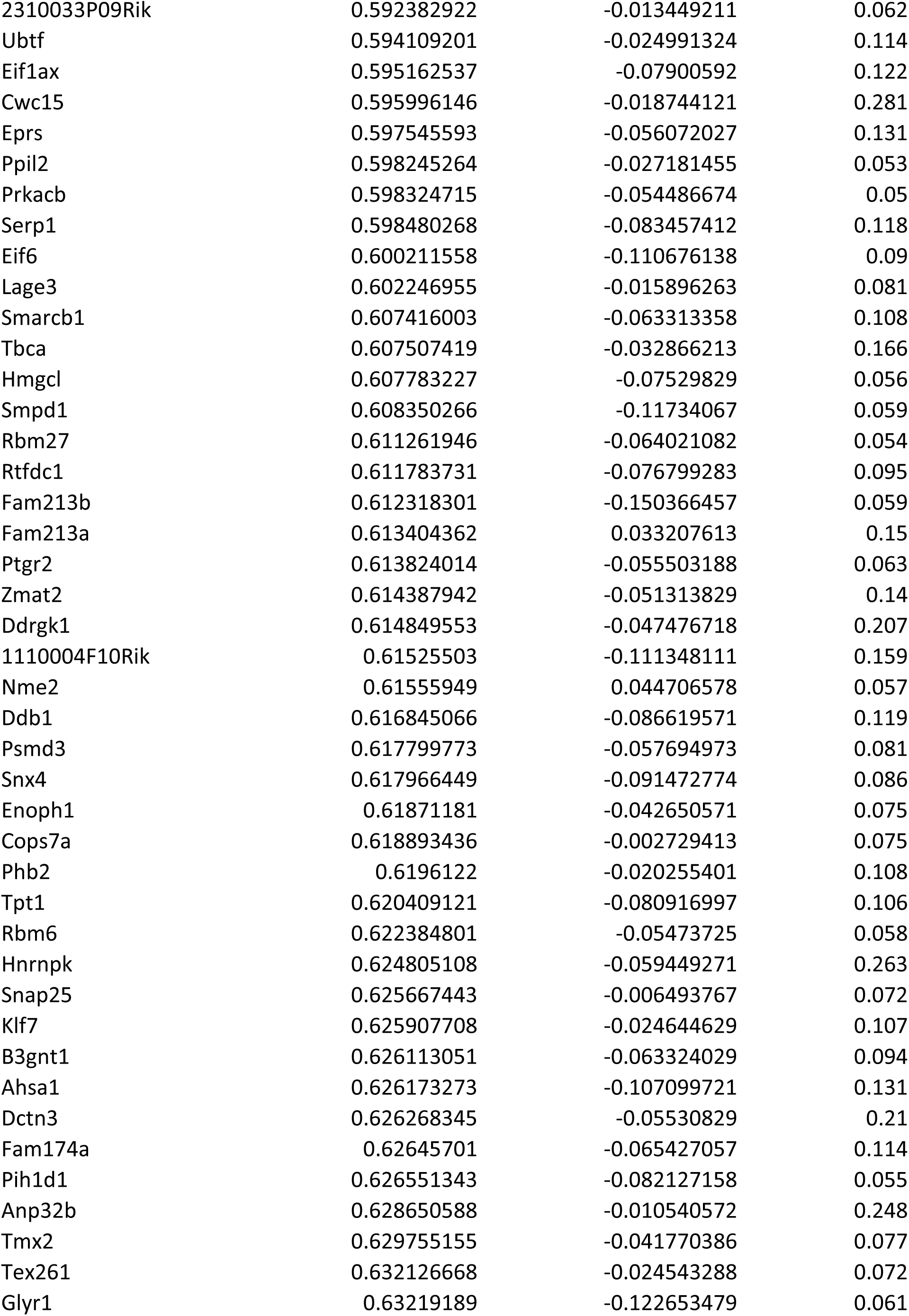
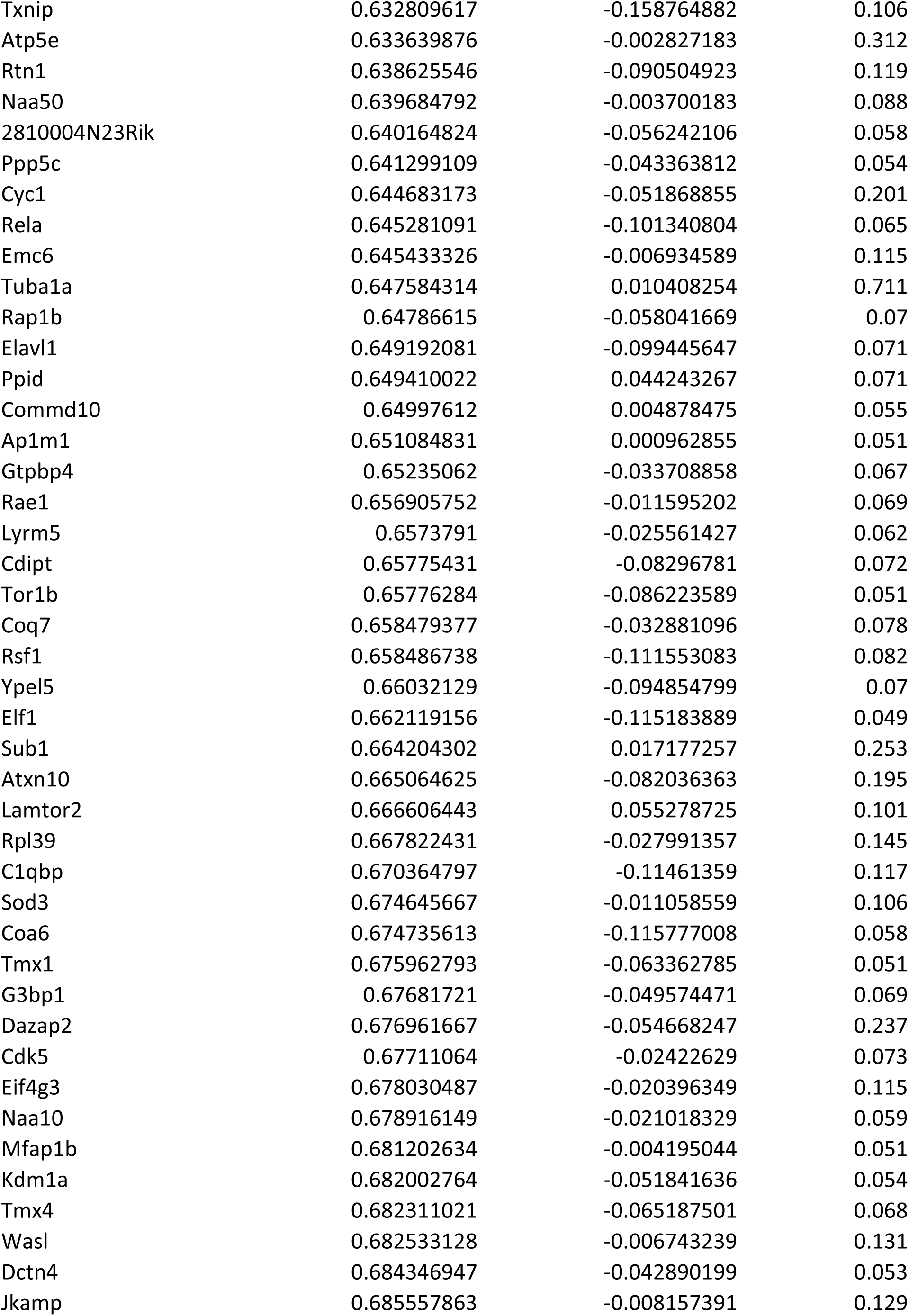
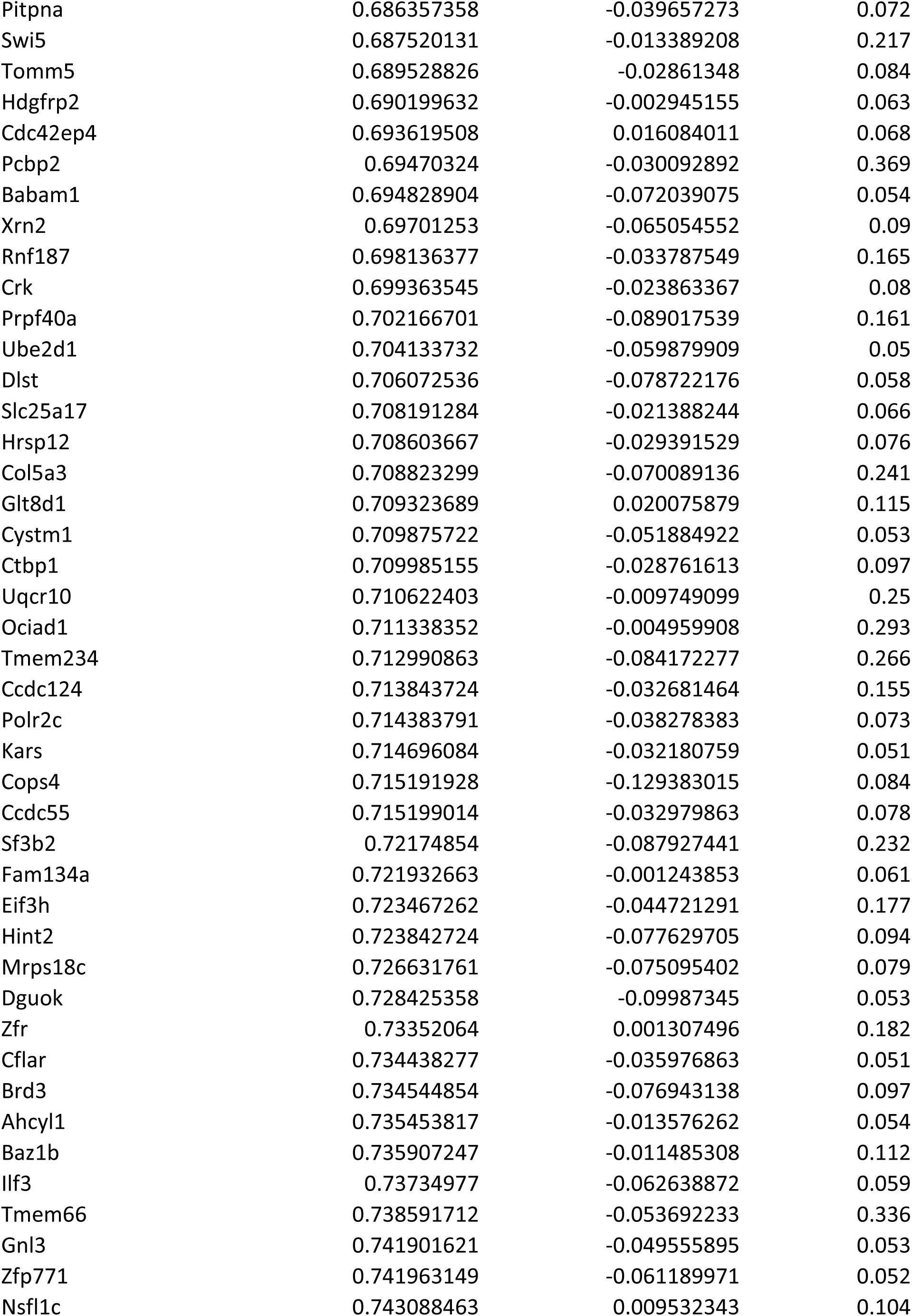
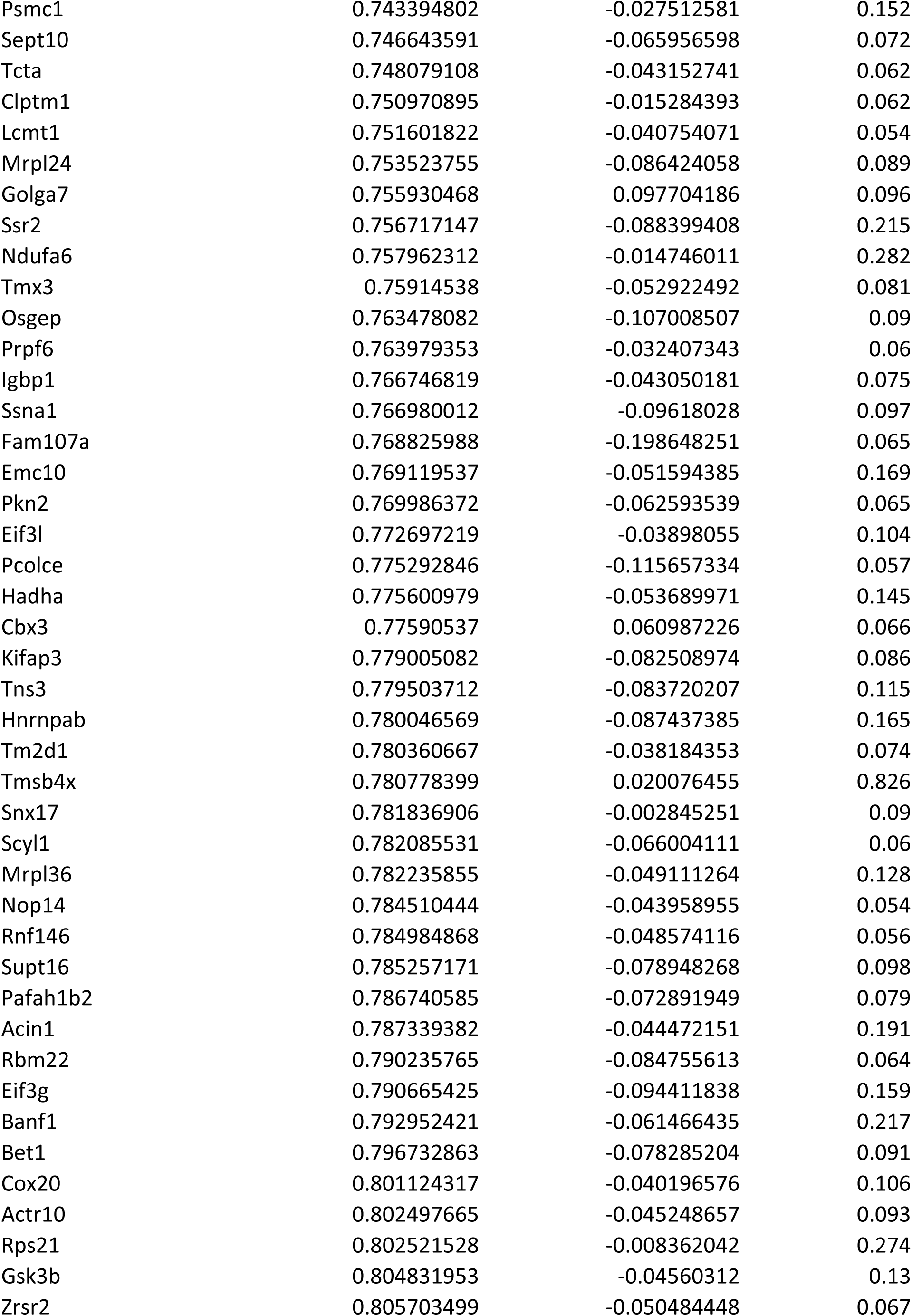
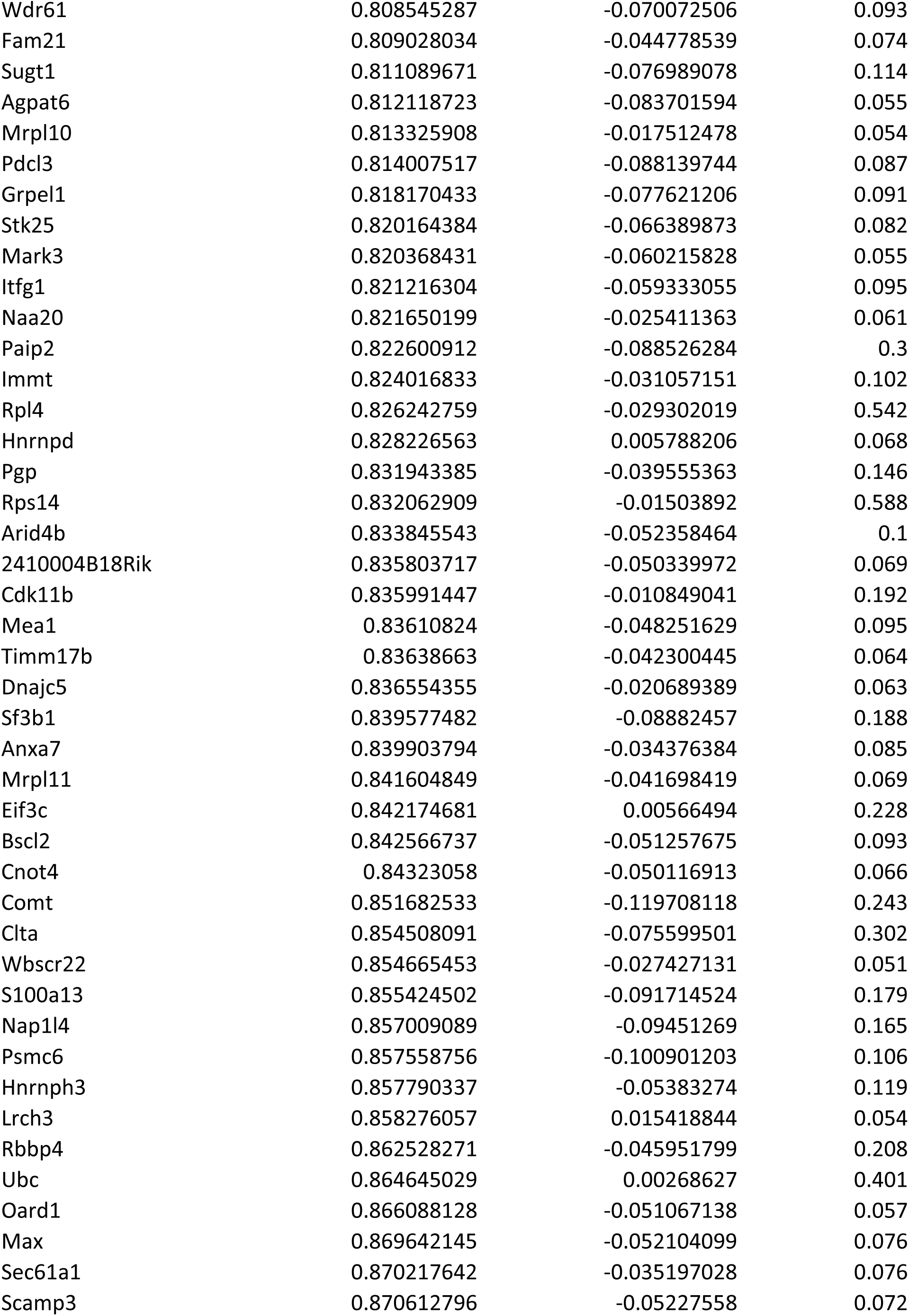
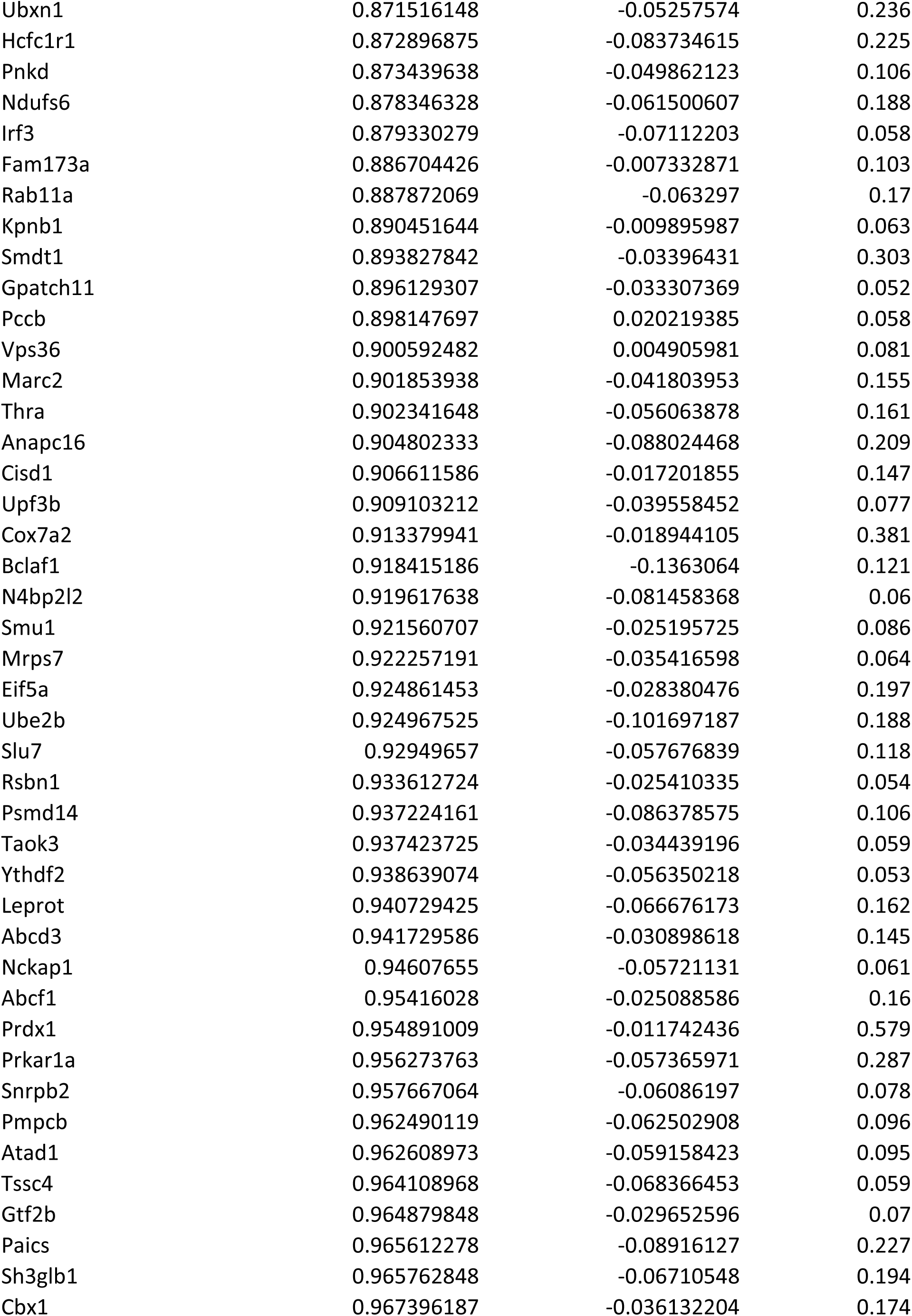
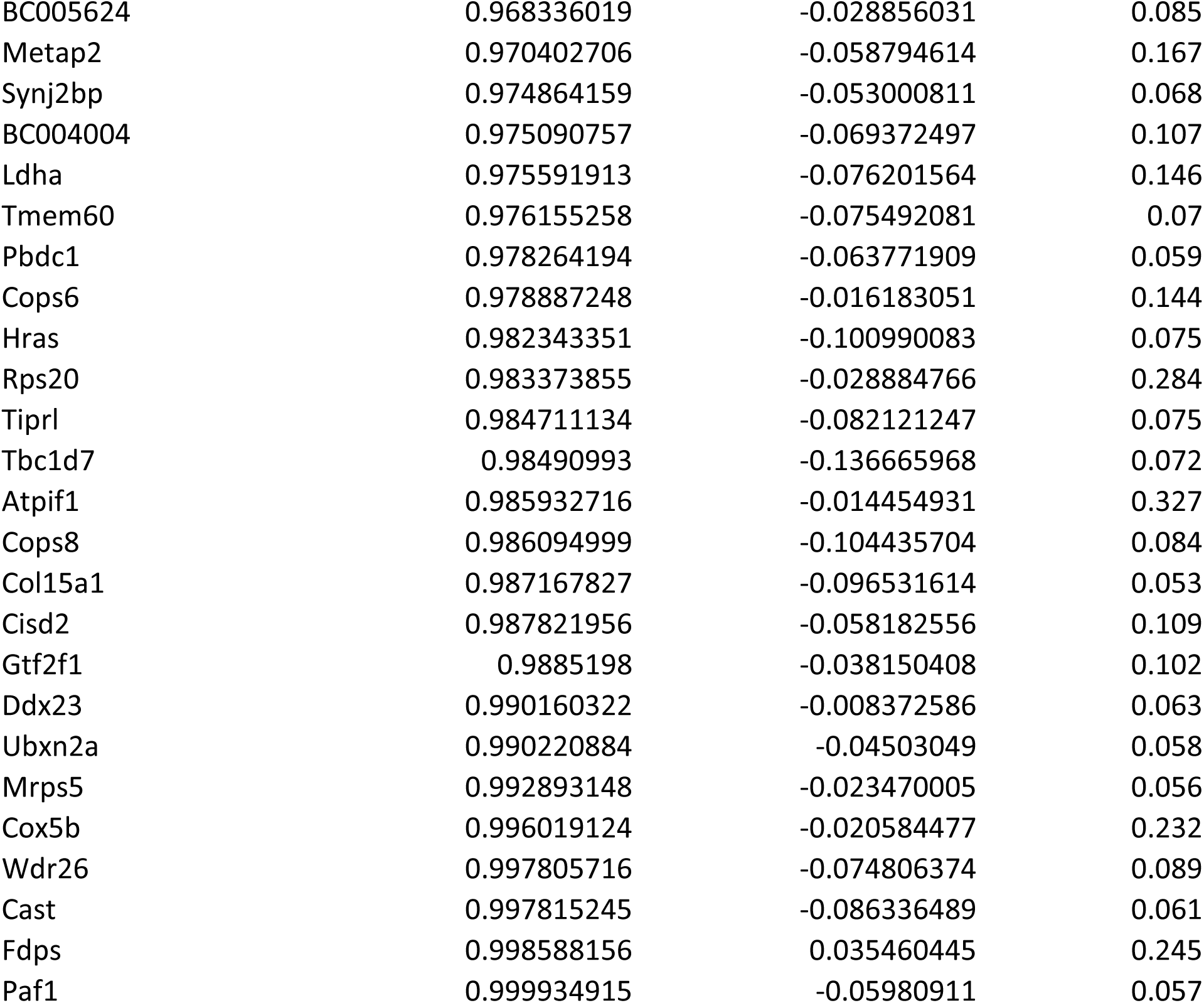

| pct.2 | p_val_adj |
| --- | --- |
| 0.006 | 0 |
| 0.603 | 0 |
| 0.541 | 0 |
| 0.02 | 0 |
| 0.011 | 0 |
| 0.013 | 0 |
| 0.01 | 0 |
| 0.679 | 0 |
| 0.021 | 0 |
| 0.166 | 0 |
| 0.101 | 0 |
| 0.004 | 0 |
| 0.132 | 0 |
| 0.006 | 0 |
| 0.004 | 0 |
| 0.044 | 0 |
| 0.017 | 0 |
| 0.002 | 0 |
| 0.017 | 0 |
| 0.003 | 0 |
| 0.032 | 0 |
| 0.008 | 0 |
| 0.001 | 0 |
| 0.002 | 0 |
| 0.059 | 0 |
| 0.004 | 0 |
| 0.094 | 0 |
| 0.033 | 0 |
| 0.03 | 0 |
| 0.05 | 0 |
| 0.002 | 0 |
| 0.005 | 0 |
| 0.004 | 0 |
| 0.002 | 0 |
| 0.006 | 0 |
| 0.143 | 0 |
| 0.019 | 0 |
| 0.041 | 0 |
| 0.008 | 0 |
| 0.244 | 0 |
| 0.051 | 0 |
| 0.022 | 0 |
| 0.058 | 0 |
| 0.02 | 0 |
| 0.007 | 0 |
| 0.43 | 0 |
| 0.012 | 0 |
| 0.005 | 0 |
| 0.006 | 0 |
| 0.844 | 0 |
| 0.518 | 0 |
| 0.537 | 0 |
| 0.004 | 0 |
| 0.755 | 0 |
| 0.002 | 0 |
| 0.005 | 0 |
| 0.003 | 0 |
| 0.003 | 0 |
| 0.003 | 0 |
| 0.66 | 0 |
| 0.001 | 0 |
| 0.004 | 0 |
| 0.006 | 0 |
| 0 | 0 |
| 0.001 | 0 |
| 0.921 | 0 |
| 0.982 | 0 |
| 0.697 | 0 |
| 0.663 | 0 |
| 0.919 | 0 |
| 0.826 | 0 |
| 0.993 | 0 |
| 0.832 | 0 |
| 0.003 | 7.00E-301 |
| 0.551 | 1.43E-294 |
| 0.11 | 7.86E-289 |
| 0.554 | 2.94E-287 |
| 0.072 | 4.50E-286 |
| 0.615 | 5.22E-286 |
| 0.176 | 9.63E-285 |
| 0.618 | 1.53E-284 |
| 0.041 | 1.68E-275 |
| 0.501 | 2.14E-275 |
| 0.004 | 6.80E-271 |
| 0.093 | 2.95E-270 |
| 0.153 | 4.81E-269 |
| 0.054 | 1.47E-267 |
| 0.163 | 6.95E-264 |
| 0.016 | 1.04E-263 |
| 0.595 | 3.79E-263 |
| 0.016 | 1.51E-258 |
| 0.517 | 1.75E-258 |
| 0.012 | 3.01E-256 |
| 0.019 | 2.27E-252 |
| 0.226 | 5.76E-252 |
| 0.002 | 2.19E-251 |
| 0.16 | 6.94E-251 |
| 0.335 | 3.51E-246 |
| 0.577 | 9.27E-242 |
| 0.524 | 6.88E-241 |
| 0.443 | 1.13E-233 |
| 0.166 | 1.18E-233 |
| 0.061 | 3.41E-233 |
| 0.005 | 5.96E-232 |
| 0.446 | 2.63E-231 |
| 0.566 | 7.30E-227 |
| 0.114 | 1.75E-226 |
| 0.007 | 6.04E-225 |
| 0.003 | 1.59E-222 |
| 0.549 | 9.90E-222 |
| 0.032 | 1.64E-220 |
| 0.002 | 1.31E-219 |
| 0.418 | 1.49E-216 |
| 0.015 | 1.97E-216 |
| 0.945 | 7.26E-216 |
| 0.063 | 1.76E-215 |
| 0 | 1.01E-214 |
| 0.014 | 2.53E-209 |
| 0.632 | 7.78E-208 |
| 0.458 | 1.26E-207 |
| 0.002 | 1.21E-206 |
| 0.014 | 1.43E-206 |
| 0.017 | 2.88E-203 |
| 0.002 | 1.99E-202 |
| 0.408 | 9.15E-202 |
| 0.117 | 1.89E-201 |
| 0.007 | 5.68E-198 |
| 0.056 | 4.86E-196 |
| 0.02 | 1.13E-193 |
| 0.391 | 2.63E-193 |
| 0.051 | 7.02E-190 |
| 0.092 | 1.69E-188 |
| 0.025 | 2.64E-188 |
| 0.678 | 1.39E-186 |
| 0.051 | 8.28E-185 |
| 0.594 | 4.91E-184 |
| 0.021 | 4.14E-183 |
| 0.004 | 1.99E-182 |
| 0.579 | 4.14E-181 |
| 0 | 7.36E-181 |
| 0.923 | 7.34E-179 |
| 0.375 | 9.97E-177 |
| 0.05 | 1.42E-176 |
| 0.033 | 1.99E-176 |
| 0.368 | 2.28E-176 |
| 0.772 | 2.98E-175 |
| 0 | 1.71E-174 |
| 0.398 | 1.50E-173 |
| 0.029 | 1.32E-172 |
| 0.026 | 3.63E-172 |
| 0.033 | 1.18E-171 |
| 0.224 | 1.82E-171 |
| 0.032 | 2.07E-171 |
| 0.002 | 1.97E-169 |
| 0.866 | 2.96E-169 |
| 0.213 | 4.18E-169 |
| 0.631 | 8.22E-169 |
| 0.001 | 1.39E-168 |
| 0.011 | 3.41E-168 |
| 0.464 | 5.84E-168 |
| 0.104 | 2.46E-167 |
| 0.842 | 1.36E-166 |
| 0.665 | 2.09E-166 |
| 0.05 | 6.82E-166 |
| 0.049 | 1.29E-165 |
| 0.005 | 3.81E-165 |
| 0.004 | 1.18E-164 |
| 0.001 | 1.22E-164 |
| 0.015 | 1.48E-164 |
| 0.426 | 2.03E-164 |
| 0.075 | 3.73E-164 |
| 0.007 | 4.92E-163 |
| 0.511 | 5.23E-161 |
| 0.003 | 2.21E-160 |
| 0.397 | 7.18E-159 |
| 0.049 | 2.00E-158 |
| 0.103 | 3.59E-158 |
| 0.096 | 4.01E-158 |
| 0.847 | 6.38E-158 |
| 0.547 | 8.46E-158 |
| 0.136 | 1.74E-157 |
| 0.484 | 2.48E-156 |
| 0.519 | 2.92E-156 |
| 0.029 | 4.24E-155 |
| 0.335 | 2.47E-153 |
| 0.008 | 3.18E-152 |
| 0.391 | 3.76E-152 |
| 0.343 | 5.97E-152 |
| 0.002 | 1.12E-151 |
| 0 | 5.59E-150 |
| 0 | 3.93E-147 |
| 0.083 | 1.04E-146 |
| 0.002 | 1.65E-146 |
| 0.021 | 1.96E-146 |
| 0.076 | 9.13E-146 |
| 0.335 | 3.53E-144 |
| 0.889 | 6.16E-144 |
| 0.013 | 1.90E-143 |
| 0.003 | 4.53E-143 |
| 0.056 | 5.70E-143 |
| 0.013 | 2.06E-142 |
| 0.001 | 2.78E-142 |
| 0.001 | 3.71E-142 |
| 0.002 | 1.12E-140 |
| 0.332 | 4.52E-140 |
| 0.637 | 7.10E-139 |
| 0.198 | 6.22E-138 |
| 0.005 | 7.02E-137 |
| 0.059 | 2.60E-136 |
| 0.026 | 1.12E-135 |
| 0.006 | 3.90E-135 |
| 0.008 | 9.57E-135 |
| 0.583 | 5.88E-133 |
| 0.033 | 1.23E-132 |
| 0.326 | 2.27E-132 |
| 0.325 | 1.48E-131 |
| 0.318 | 4.70E-131 |
| 0.001 | 8.82E-131 |
| 0.465 | 5.07E-130 |
| 0.006 | 8.44E-130 |
| 0.314 | 1.05E-129 |
| 0.221 | 1.88E-129 |
| 0.211 | 1.91E-129 |
| 0.091 | 2.44E-129 |
| 0.044 | 6.53E-129 |
| 0.088 | 2.18E-128 |
| 0.671 | 6.99E-128 |
| 0.022 | 9.53E-128 |
| 0.045 | 1.23E-127 |
| 0.365 | 2.67E-127 |
| 0.383 | 5.63E-127 |
| 0.632 | 6.00E-127 |
| 0.117 | 6.04E-127 |
| 0.092 | 1.05E-126 |
| 0.654 | 1.43E-126 |
| 0.019 | 7.94E-124 |
| 0.047 | 1.09E-122 |
| 0.178 | 2.06E-122 |
| 0.078 | 4.03E-122 |
| 0.002 | 4.96E-122 |
| 0.486 | 8.62E-122 |
| 0.277 | 2.03E-121 |
| 0.389 | 2.40E-121 |
| 0.012 | 3.41E-120 |
| 0.283 | 7.94E-120 |
| 0.284 | 2.15E-119 |
| 0.014 | 1.93E-116 |
| 0.034 | 2.34E-116 |
| 0.029 | 2.58E-116 |
| 0.123 | 1.43E-115 |
| 0.049 | 4.73E-114 |
| 0.015 | 7.68E-114 |
| 0.008 | 9.89E-114 |
| 0.274 | 8.42E-113 |
| 0.005 | 1.57E-112 |
| 0.022 | 2.24E-112 |
| 0.105 | 1.35E-110 |
| 0.363 | 3.28E-110 |
| 0.248 | 5.31E-109 |
| 0.028 | 1.13E-108 |
| 0.396 | 1.76E-108 |
| 0.263 | 2.10E-108 |
| 0.008 | 1.20E-107 |
| 0.253 | 4.39E-107 |
| 0.203 | 1.03E-106 |
| 0.06 | 1.28E-106 |
| 0.159 | 3.61E-106 |
| 0.019 | 1.10E-105 |
| 0.019 | 1.71E-105 |
| 0.022 | 6.72E-105 |
| 0.307 | 7.93E-105 |
| 0.032 | 9.35E-105 |
| 0.009 | 1.57E-104 |
| 0.002 | 2.66E-104 |
| 0.072 | 2.83E-104 |
| 0.248 | 6.35E-104 |
| 0.178 | 3.67E-103 |
| 0.119 | 5.14E-103 |
| 0.482 | 5.55E-103 |
| 0.006 | 6.75E-102 |
| 0.275 | 1.15E-100 |
| 0.008 | 1.10E-99 |
| 0.088 | 3.65E-99 |
| 0.239 | 4.70E-99 |
| 0.271 | 1.13E-98 |
| 0.232 | 1.41E-98 |
| 0.139 | 5.45E-97 |
| 0.009 | 1.34E-96 |
| 0.75 | 1.67E-96 |
| 0.051 | 1.24E-95 |
| 0.233 | 5.13E-95 |
| 0.017 | 1.93E-94 |
| 0.04 | 8.12E-93 |
| 0.033 | 1.00E-92 |
| 0.225 | 1.97E-92 |
| 0.005 | 2.94E-92 |
| 0.058 | 2.41E-91 |
| 0.006 | 7.28E-91 |
| 0.079 | 1.41E-90 |
| 0.39 | 1.54E-90 |
| 0.307 | 6.20E-90 |
| 0.233 | 8.59E-90 |
| 0.31 | 1.55E-89 |
| 0.224 | 1.81E-89 |
| 0.054 | 2.02E-89 |
| 0.217 | 4.83E-89 |
| 0.099 | 6.92E-88 |
| 0.242 | 9.27E-88 |
| 0.022 | 3.35E-87 |
| 0.02 | 3.91E-87 |
| 0.005 | 4.82E-87 |
| 0.014 | 1.34E-85 |
| 0.055 | 2.12E-85 |
| 0.017 | 1.01E-84 |
| 0.019 | 1.92E-84 |
| 0.161 | 2.31E-84 |
| 0.597 | 5.87E-84 |
| 0.212 | 2.75E-83 |
| 0.682 | 3.18E-82 |
| 0.38 | 7.73E-82 |
| 0.014 | 1.06E-81 |
| 0.335 | 1.07E-81 |
| 0.031 | 3.65E-81 |
| 0.027 | 2.39E-80 |
| 0.048 | 3.38E-80 |
| 0.401 | 5.20E-80 |
| 0.035 | 8.17E-80 |
| 0.033 | 1.27E-77 |
| 0.058 | 4.80E-77 |
| 0.074 | 5.02E-77 |
| 0.126 | 7.51E-77 |
| 0.004 | 8.35E-77 |
| 0.32 | 8.89E-77 |
| 0.02 | 2.53E-76 |
| 0.049 | 3.89E-76 |
| 0.129 | 1.76E-75 |
| 0.456 | 5.54E-75 |
| 0.2 | 6.49E-75 |
| 0.517 | 3.35E-74 |
| 0.081 | 5.06E-74 |
| 0.114 | 8.52E-74 |
| 0.014 | 9.97E-74 |
| 0.208 | 1.18E-73 |
| 0.122 | 8.33E-73 |
| 0.071 | 3.01E-72 |
| 0.006 | 4.44E-72 |
| 0.007 | 7.61E-72 |
| 0.025 | 1.71E-71 |
| 0.204 | 1.73E-71 |
| 0.049 | 2.33E-71 |
| 0.049 | 4.69E-71 |
| 0.02 | 7.39E-71 |
| 0.005 | 4.90E-70 |
| 0.212 | 5.56E-70 |
| 0.022 | 6.09E-70 |
| 0.018 | 1.60E-69 |
| 0.008 | 1.18E-68 |
| 0.174 | 1.50E-68 |
| 0.171 | 4.01E-68 |
| 0.067 | 6.41E-68 |
| 0.075 | 1.26E-67 |
| 0.016 | 1.72E-67 |
| 0.109 | 4.63E-67 |
| 0.034 | 5.24E-67 |
| 0.017 | 1.89E-66 |
| 0.174 | 6.99E-66 |
| 0.021 | 1.54E-65 |
| 0.346 | 1.82E-65 |
| 0.123 | 6.10E-65 |
| 0.212 | 7.27E-65 |
| 0.085 | 8.89E-65 |
| 0.168 | 1.51E-64 |
| 0.049 | 1.79E-64 |
| 0.143 | 1.82E-64 |
| 0.068 | 1.93E-64 |
| 0.008 | 2.00E-64 |
| 0.173 | 5.98E-64 |
| 0.16 | 8.86E-64 |
| 0.008 | 1.27E-63 |
| 0.11 | 1.65E-63 |
| 0.012 | 1.86E-63 |
| 0.143 | 2.00E-63 |
| 0.022 | 2.62E-63 |
| 0.114 | 3.87E-63 |
| 0.22 | 1.02E-62 |
| 0.163 | 1.03E-62 |
| 0.122 | 1.06E-62 |
| 0.089 | 2.43E-62 |
| 0.236 | 3.20E-62 |
| 0.254 | 7.42E-62 |
| 0.208 | 1.23E-61 |
| 0.367 | 1.80E-61 |
| 0.028 | 4.21E-61 |
| 0.456 | 5.13E-61 |
| 0.044 | 1.86E-60 |
| 0.013 | 1.93E-60 |
| 0.151 | 2.27E-60 |
| 0.289 | 2.29E-60 |
| 0.595 | 5.41E-60 |
| 0.281 | 6.61E-60 |
| 0.17 | 8.34E-60 |
| 0.026 | 1.20E-59 |
| 0.064 | 2.23E-59 |
| 0.606 | 2.83E-59 |
| 0.348 | 3.42E-59 |
| 0.801 | 6.68E-59 |
| 0.022 | 8.02E-59 |
| 0.036 | 1.31E-58 |
| 0.154 | 1.55E-58 |
| 0.3 | 1.57E-58 |
| 0.408 | 1.73E-58 |
| 0.161 | 1.95E-58 |
| 0.013 | 3.34E-58 |
| 0.7 | 5.02E-58 |
| 0.032 | 1.03E-57 |
| 0.043 | 1.07E-57 |
| 0.038 | 1.84E-57 |
| 0.648 | 2.46E-57 |
| 0.195 | 2.71E-57 |
| 0.008 | 3.44E-57 |
| 0.159 | 3.53E-57 |
| 0.137 | 5.74E-57 |
| 0.184 | 1.33E-56 |
| 0.013 | 4.38E-56 |
| 0.516 | 4.45E-56 |
| 0.156 | 4.90E-56 |
| 0.007 | 1.95E-55 |
| 0.015 | 2.82E-55 |
| 0.28 | 3.86E-55 |
| 0.51 | 5.50E-55 |
| 0.02 | 1.05E-54 |
| 0.189 | 1.15E-54 |
| 0.614 | 1.64E-54 |
| 0.012 | 1.99E-54 |
| 0.019 | 3.30E-54 |
| 0.16 | 6.01E-54 |
| 0.529 | 1.06E-53 |
| 0.392 | 1.58E-53 |
| 0.111 | 1.70E-53 |
| 0.038 | 1.75E-53 |
| 0.017 | 2.26E-53 |
| 0.019 | 4.52E-53 |
| 0.247 | 2.61E-52 |
| 0.227 | 3.32E-52 |
| 0.151 | 3.47E-52 |
| 0.138 | 4.54E-52 |
| 0.157 | 5.83E-52 |
| 0.193 | 6.70E-52 |
| 0.106 | 8.54E-52 |
| 0.144 | 9.74E-52 |
| 0.137 | 6.03E-51 |
| 0.051 | 8.42E-51 |
| 0.184 | 1.02E-50 |
| 0.189 | 1.12E-50 |
| 0.183 | 1.18E-50 |
| 0.176 | 1.75E-50 |
| 0.041 | 1.86E-50 |
| 0.162 | 2.31E-50 |
| 0.588 | 2.46E-50 |
| 0.031 | 2.93E-50 |
| 0.296 | 4.20E-50 |
| 0.143 | 5.45E-50 |
| 0.221 | 9.29E-50 |
| 0.142 | 9.57E-50 |
| 0.202 | 2.14E-49 |
| 0.472 | 2.25E-49 |
| 0.432 | 2.83E-49 |
| 0.153 | 3.17E-49 |
| 0.012 | 4.01E-49 |
| 0.18 | 4.52E-49 |
| 0.16 | 1.82E-48 |
| 0.019 | 2.09E-48 |
| 0.227 | 3.69E-48 |
| 0.348 | 5.86E-48 |
| 0.516 | 1.24E-47 |
| 0.024 | 1.89E-47 |
| 0.061 | 1.97E-47 |
| 0.224 | 2.17E-47 |
| 0.175 | 2.56E-47 |
| 0.17 | 3.72E-47 |
| 0.04 | 4.65E-47 |
| 0.015 | 6.77E-47 |
| 0.011 | 2.62E-46 |
| 0.144 | 3.74E-46 |
| 0.039 | 5.93E-46 |
| 0.023 | 8.08E-46 |
| 0.142 | 8.12E-46 |
| 0.177 | 9.82E-46 |
| 0.048 | 1.15E-45 |
| 0.264 | 2.06E-45 |
| 0.159 | 3.27E-45 |
| 0.285 | 4.41E-45 |
| 0.213 | 5.13E-45 |
| 0.129 | 1.14E-44 |
| 0.234 | 1.58E-44 |
| 0.162 | 1.78E-44 |
| 0.044 | 3.38E-44 |
| 0.075 | 4.22E-44 |
| 0.225 | 4.60E-44 |
| 0.05 | 6.38E-44 |
| 0.171 | 7.60E-44 |
| 0.12 | 1.54E-43 |
| 0.013 | 2.03E-43 |
| 0.035 | 4.33E-43 |
| 0.012 | 5.52E-43 |
| 0.155 | 6.72E-43 |
| 0.027 | 6.90E-43 |
| 0.299 | 1.20E-42 |
| 0.03 | 1.50E-42 |
| 0.024 | 1.61E-42 |
| 0.226 | 1.79E-42 |
| 0.899 | 1.93E-42 |
| 0.092 | 2.02E-42 |
| 0.178 | 2.68E-42 |
| 0.096 | 5.49E-42 |
| 0.356 | 6.38E-42 |
| 0.097 | 6.74E-42 |
| 0.179 | 8.04E-42 |
| 0.125 | 1.20E-41 |
| 0.302 | 1.25E-41 |
| 0.343 | 1.44E-41 |
| 0.119 | 1.73E-41 |
| 0.125 | 1.84E-41 |
| 0.016 | 2.54E-41 |
| 0.012 | 2.81E-41 |
| 0.114 | 3.72E-41 |
| 0.543 | 3.96E-41 |
| 0.154 | 6.15E-41 |
| 0.191 | 6.49E-41 |
| 0.126 | 7.75E-41 |
| 0.215 | 8.08E-41 |
| 0.491 | 1.07E-40 |
| 0.165 | 1.16E-40 |
| 0.156 | 1.64E-40 |
| 0.155 | 1.83E-40 |
| 0.011 | 2.02E-40 |
| 0.122 | 2.44E-40 |
| 0.148 | 3.01E-40 |
| 0.055 | 8.36E-40 |
| 0.025 | 9.41E-40 |
| 0.337 | 1.20E-39 |
| 0.064 | 1.75E-39 |
| 0.012 | 2.07E-39 |
| 0.029 | 2.25E-39 |
| 0.247 | 4.18E-39 |
| 0.022 | 4.60E-39 |
| 0.041 | 5.16E-39 |
| 0.167 | 6.02E-39 |
| 0.112 | 9.13E-39 |
| 0.133 | 9.97E-39 |
| 0.274 | 1.26E-38 |
| 0.222 | 1.27E-38 |
| 0.01 | 2.28E-38 |
| 0.213 | 4.16E-38 |
| 0.02 | 5.70E-38 |
| 0.223 | 7.20E-38 |
| 0.237 | 7.39E-38 |
| 0.039 | 8.80E-38 |
| 0.12 | 1.01E-37 |
| 0.107 | 1.22E-37 |
| 0.112 | 1.38E-37 |
| 0.122 | 1.74E-37 |
| 0.151 | 3.13E-37 |
| 0.261 | 3.42E-37 |
| 0.035 | 8.39E-37 |
| 0.647 | 1.01E-36 |
| 0.052 | 1.37E-36 |
| 0.458 | 2.43E-36 |
| 0.021 | 2.84E-36 |
| 0.48 | 6.62E-36 |
| 0.227 | 8.07E-36 |
| 0.19 | 1.23E-35 |
| 0.093 | 1.34E-35 |
| 0.165 | 1.58E-35 |
| 0.1 | 1.69E-35 |
| 0.187 | 2.67E-35 |
| 0.127 | 2.84E-35 |
| 0.065 | 3.09E-35 |
| 0.105 | 4.88E-35 |
| 0.01 | 5.53E-35 |
| 0.221 | 5.64E-35 |
| 0.107 | 7.93E-35 |
| 0.036 | 8.28E-35 |
| 0.103 | 8.93E-35 |
| 0.022 | 1.87E-34 |
| 0.075 | 2.44E-34 |
| 0.438 | 2.98E-34 |
| 0.096 | 6.39E-34 |
| 0.233 | 8.48E-34 |
| 0.159 | 1.07E-33 |
| 0.395 | 1.67E-33 |
| 0.292 | 1.71E-33 |
| 0.022 | 1.98E-33 |
| 0.169 | 2.06E-33 |
| 0.189 | 4.27E-33 |
| 0.044 | 8.34E-33 |
| 0.033 | 9.89E-33 |
| 0.116 | 1.02E-32 |
| 0.106 | 1.08E-32 |
| 0.102 | 1.50E-32 |
| 0.039 | 1.69E-32 |
| 0.342 | 5.22E-32 |
| 0.095 | 6.93E-32 |
| 0.024 | 9.59E-32 |
| 0.46 | 1.03E-31 |
| 0.1 | 1.07E-31 |
| 0.625 | 1.14E-31 |
| 0.248 | 1.59E-31 |
| 0.013 | 1.76E-31 |
| 0.035 | 1.80E-31 |
| 0.3 | 1.86E-31 |
| 0.12 | 1.92E-31 |
| 0.094 | 2.09E-31 |
| 0.054 | 2.12E-31 |
| 0.365 | 2.96E-31 |
| 0.034 | 3.54E-31 |
| 0.097 | 3.89E-31 |
| 0.091 | 5.10E-31 |
| 0.112 | 5.60E-31 |
| 0.126 | 7.71E-31 |
| 0.032 | 1.00E-30 |
| 0.018 | 2.02E-30 |
| 0.053 | 2.25E-30 |
| 0.247 | 3.34E-30 |
| 0.067 | 4.53E-30 |
| 0.089 | 4.57E-30 |
| 0.987 | 4.94E-30 |
| 0.084 | 4.98E-30 |
| 0.031 | 5.38E-30 |
| 0.444 | 6.32E-30 |
| 0.161 | 7.27E-30 |
| 0.165 | 8.11E-30 |
| 0.357 | 8.93E-30 |
| 0.185 | 9.94E-30 |
| 0.039 | 1.08E-29 |
| 0.014 | 1.21E-29 |
| 0.117 | 2.02E-29 |
| 0.402 | 2.10E-29 |
| 0.045 | 2.42E-29 |
| 0.036 | 2.53E-29 |
| 0.191 | 3.43E-29 |
| 0.146 | 4.55E-29 |
| 0.206 | 5.40E-29 |
| 0.353 | 5.67E-29 |
| 0.087 | 5.69E-29 |
| 0.061 | 7.09E-29 |
| 0.047 | 1.08E-28 |
| 0.02 | 1.09E-28 |
| 0.1 | 1.13E-28 |
| 0.053 | 1.35E-28 |
| 0.083 | 1.37E-28 |
| 0.087 | 1.50E-28 |
| 0.025 | 1.61E-28 |
| 0.106 | 1.72E-28 |
| 0.012 | 2.11E-28 |
| 0.107 | 2.67E-28 |
| 0.419 | 4.20E-28 |
| 0.014 | 5.29E-28 |
| 0.096 | 5.84E-28 |
| 0.025 | 6.70E-28 |
| 0.411 | 7.08E-28 |
| 0.09 | 7.57E-28 |
| 0.143 | 8.85E-28 |
| 0.303 | 1.02E-27 |
| 0.133 | 1.03E-27 |
| 0.172 | 1.24E-27 |
| 0.111 | 1.55E-27 |
| 0.088 | 1.89E-27 |
| 0.019 | 2.04E-27 |
| 0.156 | 2.28E-27 |
| 0.047 | 2.31E-27 |
| 0.012 | 2.48E-27 |
| 0.038 | 2.52E-27 |
| 0.083 | 2.59E-27 |
| 0.673 | 3.13E-27 |
| 0.014 | 3.47E-27 |
| 0.055 | 3.76E-27 |
| 0.09 | 3.94E-27 |
| 0.12 | 4.34E-27 |
| 0.092 | 4.38E-27 |
| 0.015 | 7.00E-27 |
| 0.163 | 7.22E-27 |
| 0.101 | 9.43E-27 |
| 0.189 | 1.36E-26 |
| 0.064 | 1.41E-26 |
| 0.164 | 1.74E-26 |
| 0.022 | 1.85E-26 |
| 0.024 | 2.13E-26 |
| 0.022 | 2.42E-26 |
| 0.066 | 2.44E-26 |
| 0.103 | 2.93E-26 |
| 0.105 | 2.98E-26 |
| 0.022 | 3.23E-26 |
| 0.188 | 3.72E-26 |
| 0.093 | 4.04E-26 |
| 0.195 | 4.09E-26 |
| 0.287 | 4.27E-26 |
| 0.156 | 4.34E-26 |
| 0.174 | 4.85E-26 |
| 0.188 | 5.29E-26 |
| 0.064 | 9.89E-26 |
| 0.046 | 2.01E-25 |
| 0.076 | 2.32E-25 |
| 0.625 | 2.54E-25 |
| 0.863 | 2.80E-25 |
| 0.095 | 3.17E-25 |
| 0.039 | 3.28E-25 |
| 0.015 | 3.90E-25 |
| 0.038 | 5.85E-25 |
| 0.232 | 6.00E-25 |
| 0.127 | 6.09E-25 |
| 0.018 | 6.87E-25 |
| 0.364 | 7.17E-25 |
| 0.08 | 7.78E-25 |
| 0.193 | 8.12E-25 |
| 0.093 | 8.68E-25 |
| 0.079 | 9.80E-25 |
| 0.193 | 1.18E-24 |
| 0.079 | 1.23E-24 |
| 0.018 | 2.19E-24 |
| 0.086 | 2.25E-24 |
| 0.122 | 2.40E-24 |
| 0.098 | 2.54E-24 |
| 0.052 | 2.59E-24 |
| 0.138 | 2.76E-24 |
| 0.097 | 3.94E-24 |
| 0.194 | 4.11E-24 |
| 0.046 | 5.02E-24 |
| 0.015 | 8.10E-24 |
| 0.17 | 8.16E-24 |
| 0.17 | 8.21E-24 |
| 0.098 | 1.53E-23 |
| 0.074 | 1.61E-23 |
| 0.077 | 1.95E-23 |
| 0.21 | 1.99E-23 |
| 0.111 | 2.06E-23 |
| 0.114 | 2.26E-23 |
| 0.067 | 2.74E-23 |
| 0.039 | 2.76E-23 |
| 0.182 | 3.40E-23 |
| 0.032 | 3.68E-23 |
| 0.152 | 4.37E-23 |
| 0.327 | 5.27E-23 |
| 0.067 | 5.81E-23 |
| 0.079 | 6.96E-23 |
| 0.126 | 8.60E-23 |
| 0.095 | 8.83E-23 |
| 0.301 | 9.09E-23 |
| 0.113 | 1.06E-22 |
| 0.019 | 2.09E-22 |
| 0.181 | 2.71E-22 |
| 0.226 | 2.79E-22 |
| 0.068 | 3.11E-22 |
| 0.023 | 3.12E-22 |
| 0.024 | 3.52E-22 |
| 0.115 | 3.61E-22 |
| 0.077 | 3.62E-22 |
| 0.087 | 3.76E-22 |
| 0.031 | 4.45E-22 |
| 0.277 | 4.58E-22 |
| 0.053 | 4.77E-22 |
| 0.086 | 5.16E-22 |
| 0.073 | 5.60E-22 |
| 0.023 | 5.64E-22 |
| 0.067 | 5.65E-22 |
| 0.141 | 6.59E-22 |
| 0.207 | 6.87E-22 |
| 0.265 | 8.30E-22 |
| 0.019 | 8.82E-22 |
| 0.145 | 8.95E-22 |
| 0.051 | 1.19E-21 |
| 0.02 | 1.28E-21 |
| 0.084 | 1.46E-21 |
| 0.018 | 1.60E-21 |
| 0.39 | 1.68E-21 |
| 0.069 | 1.79E-21 |
| 0.086 | 1.82E-21 |
| 0.245 | 1.91E-21 |
| 0.034 | 2.82E-21 |
| 0.138 | 3.64E-21 |
| 0.019 | 3.77E-21 |
| 0.049 | 5.28E-21 |
| 0.077 | 5.43E-21 |
| 0.016 | 5.49E-21 |
| 0.112 | 6.49E-21 |
| 0.184 | 1.12E-20 |
| 0.338 | 1.14E-20 |
| 0.115 | 1.17E-20 |
| 0.077 | 1.22E-20 |
| 0.08 | 1.42E-20 |
| 0.134 | 1.48E-20 |
| 0.064 | 1.73E-20 |
| 0.397 | 1.82E-20 |
| 0.231 | 2.01E-20 |
| 0.109 | 2.11E-20 |
| 0.175 | 2.13E-20 |
| 0.072 | 2.56E-20 |
| 0.024 | 2.62E-20 |
| 0.142 | 3.18E-20 |
| 0.05 | 3.35E-20 |
| 0.103 | 3.63E-20 |
| 0.152 | 3.77E-20 |
| 0.059 | 4.45E-20 |
| 0.039 | 4.64E-20 |
| 0.109 | 4.97E-20 |
| 0.024 | 4.99E-20 |
| 0.126 | 7.24E-20 |
| 0.171 | 7.96E-20 |
| 0.052 | 1.00E-19 |
| 0.35 | 1.06E-19 |
| 0.064 | 1.18E-19 |
| 0.229 | 1.49E-19 |
| 0.094 | 1.55E-19 |
| 0.059 | 1.78E-19 |
| 0.088 | 1.98E-19 |
| 0.019 | 2.24E-19 |
| 0.087 | 2.25E-19 |
| 0.25 | 2.35E-19 |
| 0.364 | 2.37E-19 |
| 0.079 | 2.63E-19 |
| 0.059 | 2.86E-19 |
| 0.062 | 2.87E-19 |
| 0.058 | 3.38E-19 |
| 0.176 | 3.40E-19 |
| 0.048 | 3.49E-19 |
| 0.224 | 3.64E-19 |
| 0.111 | 3.71E-19 |
| 0.126 | 4.18E-19 |
| 0.075 | 6.35E-19 |
| 0.063 | 6.93E-19 |
| 0.035 | 7.01E-19 |
| 0.171 | 7.19E-19 |
| 0.024 | 7.51E-19 |
| 0.47 | 8.55E-19 |
| 0.044 | 8.67E-19 |
| 0.419 | 8.78E-19 |
| 0.126 | 9.88E-19 |
| 0.049 | 9.94E-19 |
| 0.02 | 1.22E-18 |
| 0.353 | 1.28E-18 |
| 0.062 | 1.32E-18 |
| 0.063 | 1.42E-18 |
| 0.1 | 1.79E-18 |
| 0.069 | 1.80E-18 |
| 0.035 | 2.17E-18 |
| 0.109 | 2.61E-18 |
| 0.067 | 2.85E-18 |
| 0.207 | 2.98E-18 |
| 0.258 | 3.28E-18 |
| 0.072 | 3.59E-18 |
| 0.025 | 3.75E-18 |
| 0.024 | 3.85E-18 |
| 0.054 | 3.93E-18 |
| 0.115 | 3.96E-18 |
| 0.117 | 4.21E-18 |
| 0.02 | 4.22E-18 |
| 0.116 | 4.54E-18 |
| 0.125 | 5.01E-18 |
| 0.301 | 5.17E-18 |
| 0.086 | 5.23E-18 |
| 0.301 | 6.03E-18 |
| 0.262 | 7.28E-18 |
| 0.224 | 7.38E-18 |
| 0.055 | 8.92E-18 |
| 0.062 | 1.00E-17 |
| 0.141 | 1.07E-17 |
| 0.08 | 1.39E-17 |
| 0.055 | 1.70E-17 |
| 0.056 | 2.02E-17 |
| 0.155 | 2.13E-17 |
| 0.058 | 2.29E-17 |
| 0.081 | 2.81E-17 |
| 0.172 | 3.11E-17 |
| 0.089 | 3.18E-17 |
| 0.101 | 3.30E-17 |
| 0.069 | 3.31E-17 |
| 0.343 | 3.33E-17 |
| 0.063 | 3.55E-17 |
| 0.189 | 3.67E-17 |
| 0.035 | 3.86E-17 |
| 0.176 | 4.20E-17 |
| 0.077 | 4.57E-17 |
| 0.104 | 4.59E-17 |
| 0.169 | 5.26E-17 |
| 0.083 | 5.45E-17 |
| 0.069 | 6.68E-17 |
| 0.062 | 7.01E-17 |
| 0.113 | 7.24E-17 |
| 0.141 | 7.86E-17 |
| 0.02 | 8.00E-17 |
| 0.059 | 8.15E-17 |
| 0.052 | 1.12E-16 |
| 0.097 | 1.28E-16 |
| 0.165 | 1.30E-16 |
| 0.02 | 1.75E-16 |
| 0.11 | 2.70E-16 |
| 0.028 | 2.71E-16 |
| 0.183 | 2.76E-16 |
| 0.293 | 2.79E-16 |
| 0.048 | 3.20E-16 |
| 0.087 | 3.72E-16 |
| 0.095 | 4.28E-16 |
| 0.032 | 4.58E-16 |
| 0.022 | 4.70E-16 |
| 0.085 | 5.19E-16 |
| 0.069 | 5.93E-16 |
| 0.165 | 6.12E-16 |
| 0.581 | 6.41E-16 |
| 0.186 | 6.53E-16 |
| 0.053 | 7.46E-16 |
| 0.061 | 7.67E-16 |
| 0.048 | 8.89E-16 |
| 0.025 | 9.03E-16 |
| 0.376 | 9.08E-16 |
| 0.113 | 9.55E-16 |
| 0.107 | 9.90E-16 |
| 0.245 | 1.19E-15 |
| 0.203 | 1.29E-15 |
| 0.051 | 1.29E-15 |
| 0.171 | 1.44E-15 |
| 0.104 | 1.48E-15 |
| 0.022 | 1.48E-15 |
| 0.055 | 1.48E-15 |
| 0.107 | 1.52E-15 |
| 0.104 | 1.72E-15 |
| 0.02 | 1.83E-15 |
| 0.254 | 2.13E-15 |
| 0.032 | 2.16E-15 |
| 0.116 | 2.19E-15 |
| 0.032 | 2.31E-15 |
| 0.036 | 2.55E-15 |
| 0.037 | 2.68E-15 |
| 0.126 | 2.77E-15 |
| 0.021 | 2.92E-15 |
| 0.102 | 3.12E-15 |
| 0.584 | 3.13E-15 |
| 0.179 | 3.29E-15 |
| 0.103 | 3.39E-15 |
| 0.15 | 3.54E-15 |
| 0.063 | 4.38E-15 |
| 0.059 | 4.47E-15 |
| 0.412 | 4.75E-15 |
| 0.045 | 5.41E-15 |
| 0.1 | 5.53E-15 |
| 0.036 | 5.89E-15 |
| 0.043 | 6.48E-15 |
| 0.264 | 7.09E-15 |
| 0.061 | 7.13E-15 |
| 0.035 | 7.36E-15 |
| 0.099 | 8.36E-15 |
| 0.201 | 8.92E-15 |
| 0.142 | 9.91E-15 |
| 0.04 | 1.03E-14 |
| 0.202 | 1.16E-14 |
| 0.151 | 1.18E-14 |
| 0.084 | 1.23E-14 |
| 0.205 | 1.25E-14 |
| 0.03 | 1.28E-14 |
| 0.096 | 1.35E-14 |
| 0.018 | 1.36E-14 |
| 0.073 | 1.55E-14 |
| 0.045 | 1.59E-14 |
| 0.431 | 1.61E-14 |
| 0.048 | 1.64E-14 |
| 0.053 | 2.22E-14 |
| 0.058 | 2.34E-14 |
| 0.042 | 2.89E-14 |
| 0.056 | 2.99E-14 |
| 0.22 | 3.04E-14 |
| 0.058 | 3.23E-14 |
| 0.101 | 3.34E-14 |
| 0.019 | 3.36E-14 |
| 0.324 | 3.54E-14 |
| 0.101 | 3.68E-14 |
| 0.106 | 3.76E-14 |
| 0.164 | 3.94E-14 |
| 0.176 | 4.39E-14 |
| 0.143 | 4.40E-14 |
| 0.127 | 4.69E-14 |
| 0.155 | 5.07E-14 |
| 0.085 | 5.53E-14 |
| 0.116 | 6.09E-14 |
| 0.198 | 6.13E-14 |
| 0.14 | 7.09E-14 |
| 0.298 | 7.73E-14 |
| 0.027 | 8.96E-14 |
| 0.213 | 9.61E-14 |
| 0.119 | 1.16E-13 |
| 0.077 | 1.17E-13 |
| 0.072 | 1.18E-13 |
| 0.06 | 1.18E-13 |
| 0.22 | 1.32E-13 |
| 0.063 | 1.37E-13 |
| 0.255 | 1.50E-13 |
| 0.036 | 1.56E-13 |
| 0.25 | 1.69E-13 |
| 0.051 | 1.84E-13 |
| 0.096 | 2.18E-13 |
| 0.056 | 2.19E-13 |
| 0.093 | 2.19E-13 |
| 0.465 | 2.23E-13 |
| 0.198 | 2.23E-13 |
| 0.019 | 2.40E-13 |
| 0.028 | 2.48E-13 |
| 0.083 | 2.62E-13 |
| 0.233 | 2.72E-13 |
| 0.112 | 2.73E-13 |
| 0.247 | 2.73E-13 |
| 0.039 | 2.80E-13 |
| 0.027 | 2.89E-13 |
| 0.019 | 2.93E-13 |
| 0.045 | 3.07E-13 |
| 0.056 | 3.15E-13 |
| 0.079 | 3.82E-13 |
| 0.182 | 3.87E-13 |
| 0.165 | 4.17E-13 |
| 0.113 | 4.45E-13 |
| 0.78 | 4.63E-13 |
| 0.384 | 4.63E-13 |
| 0.046 | 5.59E-13 |
| 0.084 | 5.95E-13 |
| 0.157 | 6.45E-13 |
| 0.342 | 6.54E-13 |
| 0.028 | 6.66E-13 |
| 0.654 | 7.76E-13 |
| 0.046 | 8.01E-13 |
| 0.052 | 9.17E-13 |
| 0.061 | 9.46E-13 |
| 0.05 | 1.03E-12 |
| 0.111 | 1.11E-12 |
| 0.036 | 1.14E-12 |
| 0.079 | 1.17E-12 |
| 0.083 | 1.19E-12 |
| 0.131 | 1.23E-12 |
| 0.213 | 1.28E-12 |
| 0.079 | 1.28E-12 |
| 0.029 | 1.47E-12 |
| 0.103 | 1.97E-12 |
| 0.081 | 2.02E-12 |
| 0.02 | 2.09E-12 |
| 0.055 | 2.11E-12 |
| 0.028 | 2.18E-12 |
| 0.274 | 2.25E-12 |
| 0.022 | 2.49E-12 |
| 0.053 | 2.67E-12 |
| 0.143 | 2.88E-12 |
| 0.028 | 3.02E-12 |
| 0.036 | 3.40E-12 |
| 0.129 | 3.53E-12 |
| 0.08 | 3.66E-12 |
| 0.027 | 3.99E-12 |
| 0.019 | 4.07E-12 |
| 0.089 | 4.40E-12 |
| 0.114 | 4.66E-12 |
| 0.031 | 5.03E-12 |
| 0.587 | 5.48E-12 |
| 0.043 | 6.19E-12 |
| 0.075 | 6.19E-12 |
| 0.03 | 6.25E-12 |
| 0.024 | 6.52E-12 |
| 0.049 | 6.73E-12 |
| 0.03 | 7.32E-12 |
| 0.157 | 7.37E-12 |
| 0.044 | 7.39E-12 |
| 0.188 | 7.80E-12 |
| 0.039 | 7.99E-12 |
| 0.106 | 8.18E-12 |
| 0.059 | 8.38E-12 |
| 0.058 | 8.59E-12 |
| 0.164 | 9.07E-12 |
| 0.122 | 9.65E-12 |
| 0.153 | 1.01E-11 |
| 0.055 | 1.04E-11 |
| 0.174 | 1.08E-11 |
| 0.054 | 1.15E-11 |
| 0.052 | 1.15E-11 |
| 0.06 | 1.17E-11 |
| 0.07 | 1.33E-11 |
| 0.122 | 1.44E-11 |
| 0.388 | 1.47E-11 |
| 0.025 | 1.58E-11 |
| 0.111 | 1.62E-11 |
| 0.113 | 1.66E-11 |
| 0.028 | 1.67E-11 |
| 0.148 | 1.90E-11 |
| 0.279 | 1.94E-11 |
| 0.06 | 2.13E-11 |
| 0.052 | 2.14E-11 |
| 0.09 | 2.27E-11 |
| 0.299 | 2.45E-11 |
| 0.123 | 2.60E-11 |
| 0.04 | 2.97E-11 |
| 0.034 | 3.13E-11 |
| 0.116 | 3.20E-11 |
| 0.582 | 3.33E-11 |
| 0.361 | 3.45E-11 |
| 0.091 | 3.82E-11 |
| 0.048 | 3.83E-11 |
| 0.16 | 4.03E-11 |
| 0.157 | 4.46E-11 |
| 0.027 | 5.86E-11 |
| 0.328 | 5.95E-11 |
| 0.273 | 5.96E-11 |
| 0.04 | 6.46E-11 |
| 0.061 | 6.72E-11 |
| 0.069 | 6.75E-11 |
| 0.083 | 6.81E-11 |
| 0.135 | 9.83E-11 |
| 0.024 | 1.01E-10 |
| 0.065 | 1.14E-10 |
| 0.081 | 1.17E-10 |
| 0.028 | 1.29E-10 |
| 0.253 | 1.30E-10 |
| 0.031 | 1.36E-10 |
| 0.104 | 1.60E-10 |
| 0.155 | 1.81E-10 |
| 0.045 | 1.88E-10 |
| 0.19 | 2.07E-10 |
| 0.261 | 2.12E-10 |
| 0.16 | 2.14E-10 |
| 0.109 | 2.17E-10 |
| 0.031 | 2.25E-10 |
| 0.028 | 2.26E-10 |
| 0.051 | 2.43E-10 |
| 0.06 | 2.63E-10 |
| 0.251 | 3.16E-10 |
| 0.038 | 3.77E-10 |
| 0.187 | 3.89E-10 |
| 0.066 | 4.50E-10 |
| 0.122 | 4.50E-10 |
| 0.117 | 4.61E-10 |
| 0.105 | 4.62E-10 |
| 0.061 | 5.35E-10 |
| 0.145 | 5.74E-10 |
| 0.065 | 6.06E-10 |
| 0.08 | 6.59E-10 |
| 0.088 | 6.67E-10 |
| 0.161 | 6.71E-10 |
| 0.409 | 6.99E-10 |
| 0.023 | 7.27E-10 |
| 0.121 | 7.45E-10 |
| 0.193 | 7.71E-10 |
| 0.098 | 7.97E-10 |
| 0.341 | 8.41E-10 |
| 0.088 | 8.53E-10 |
| 0.201 | 8.77E-10 |
| 0.197 | 9.17E-10 |
| 0.083 | 9.91E-10 |
| 0.061 | 1.17E-09 |
| 0.041 | 1.40E-09 |
| 0.049 | 1.57E-09 |
| 0.067 | 1.60E-09 |
| 0.441 | 1.72E-09 |
| 0.037 | 1.75E-09 |
| 0.122 | 1.79E-09 |
| 0.061 | 1.80E-09 |
| 0.099 | 1.81E-09 |
| 0.028 | 1.86E-09 |
| 0.092 | 1.87E-09 |
| 0.213 | 1.90E-09 |
| 0.141 | 1.93E-09 |
| 0.052 | 2.19E-09 |
| 0.037 | 2.29E-09 |
| 0.057 | 2.37E-09 |
| 0.053 | 2.56E-09 |
| 0.027 | 2.56E-09 |
| 0.068 | 2.63E-09 |
| 0.132 | 2.74E-09 |
| 0.081 | 2.95E-09 |
| 0.266 | 2.96E-09 |
| 0.059 | 3.04E-09 |
| 0.051 | 3.17E-09 |
| 0.074 | 3.29E-09 |
| 0.072 | 3.61E-09 |
| 0.195 | 4.02E-09 |
| 0.107 | 4.06E-09 |
| 0.028 | 4.32E-09 |
| 0.151 | 4.45E-09 |
| 0.055 | 4.73E-09 |
| 0.091 | 4.92E-09 |
| 0.221 | 4.95E-09 |
| 0.032 | 5.39E-09 |
| 0.09 | 5.47E-09 |
| 0.645 | 5.87E-09 |
| 0.12 | 6.17E-09 |
| 0.039 | 6.37E-09 |
| 0.264 | 6.67E-09 |
| 0.182 | 7.51E-09 |
| 0.221 | 7.86E-09 |
| 0.071 | 8.38E-09 |
| 0.188 | 8.39E-09 |
| 0.335 | 8.51E-09 |
| 0.051 | 8.99E-09 |
| 0.143 | 9.15E-09 |
| 0.112 | 9.18E-09 |
| 0.109 | 9.51E-09 |
| 0.124 | 9.54E-09 |
| 0.225 | 9.59E-09 |
| 0.078 | 9.90E-09 |
| 0.035 | 1.02E-08 |
| 0.027 | 1.02E-08 |
| 0.035 | 1.15E-08 |
| 0.088 | 1.27E-08 |
| 0.075 | 1.28E-08 |
| 0.043 | 1.34E-08 |
| 0.034 | 1.44E-08 |
| 0.19 | 1.58E-08 |
| 0.086 | 1.63E-08 |
| 0.082 | 1.64E-08 |
| 0.196 | 1.65E-08 |
| 0.357 | 1.69E-08 |
| 0.293 | 1.71E-08 |
| 0.071 | 1.79E-08 |
| 0.529 | 1.88E-08 |
| 0.053 | 1.96E-08 |
| 0.068 | 2.06E-08 |
| 0.056 | 2.13E-08 |
| 0.052 | 2.14E-08 |
| 0.037 | 2.16E-08 |
| 0.082 | 2.18E-08 |
| 0.058 | 2.29E-08 |
| 0.075 | 2.30E-08 |
| 0.061 | 2.32E-08 |
| 0.42 | 2.41E-08 |
| 0.03 | 2.50E-08 |
| 0.102 | 2.69E-08 |
| 0.307 | 2.73E-08 |
| 0.052 | 2.76E-08 |
| 0.108 | 2.83E-08 |
| 0.083 | 3.06E-08 |
| 0.089 | 3.09E-08 |
| 0.094 | 3.24E-08 |
| 0.062 | 3.48E-08 |
| 0.065 | 3.61E-08 |
| 0.036 | 3.87E-08 |
| 0.072 | 3.95E-08 |
| 0.069 | 4.06E-08 |
| 0.17 | 4.10E-08 |
| 0.076 | 4.58E-08 |
| 0.414 | 4.63E-08 |
| 0.052 | 4.81E-08 |
| 0.11 | 4.93E-08 |
| 0.04 | 4.95E-08 |
| 0.038 | 5.00E-08 |
| 0.056 | 5.51E-08 |
| 0.155 | 5.51E-08 |
| 0.051 | 5.60E-08 |
| 0.304 | 6.18E-08 |
| 0.443 | 6.34E-08 |
| 0.367 | 6.53E-08 |
| 0.089 | 6.57E-08 |
| 0.061 | 6.82E-08 |
| 0.228 | 7.37E-08 |
| 0.069 | 7.43E-08 |
| 0.196 | 7.89E-08 |
| 0.157 | 8.22E-08 |
| 0.161 | 8.65E-08 |
| 0.07 | 9.15E-08 |
| 0.092 | 9.23E-08 |
| 0.155 | 9.66E-08 |
| 0.027 | 9.92E-08 |
| 0.191 | 1.05E-07 |
| 0.022 | 1.11E-07 |
| 0.047 | 1.12E-07 |
| 0.165 | 1.12E-07 |
| 0.093 | 1.16E-07 |
| 0.024 | 1.16E-07 |
| 0.079 | 1.18E-07 |
| 0.025 | 1.24E-07 |
| 0.183 | 1.33E-07 |
| 0.085 | 1.43E-07 |
| 0.11 | 1.44E-07 |
| 0.068 | 1.45E-07 |
| 0.048 | 1.48E-07 |
| 0.065 | 1.57E-07 |
| 0.058 | 1.61E-07 |
| 0.03 | 1.62E-07 |
| 0.119 | 1.63E-07 |
| 0.365 | 1.68E-07 |
| 0.331 | 1.74E-07 |
| 0.106 | 1.74E-07 |
| 0.038 | 1.76E-07 |
| 0.184 | 1.81E-07 |
| 0.031 | 1.82E-07 |
| 0.125 | 1.85E-07 |
| 0.09 | 2.02E-07 |
| 0.024 | 2.15E-07 |
| 0.056 | 2.17E-07 |
| 0.347 | 2.22E-07 |
| 0.101 | 2.26E-07 |
| 0.035 | 2.36E-07 |
| 0.112 | 2.52E-07 |
| 0.039 | 2.64E-07 |
| 0.069 | 2.65E-07 |
| 0.05 | 2.73E-07 |
| 0.11 | 2.76E-07 |
| 0.049 | 2.77E-07 |
| 0.025 | 2.99E-07 |
| 0.055 | 3.22E-07 |
| 0.42 | 3.26E-07 |
| 0.035 | 3.45E-07 |
| 0.088 | 3.54E-07 |
| 0.248 | 3.59E-07 |
| 0.234 | 3.68E-07 |
| 0.079 | 4.17E-07 |
| 0.124 | 4.18E-07 |
| 0.077 | 4.36E-07 |
| 0.053 | 4.40E-07 |
| 0.113 | 4.75E-07 |
| 0.065 | 4.79E-07 |
| 0.22 | 4.88E-07 |
| 0.033 | 4.89E-07 |
| 0.208 | 4.94E-07 |
| 0.104 | 5.19E-07 |
| 0.236 | 5.73E-07 |
| 0.2 | 6.10E-07 |
| 0.027 | 6.21E-07 |
| 0.133 | 6.34E-07 |
| 0.034 | 6.40E-07 |
| 0.026 | 6.57E-07 |
| 0.12 | 6.97E-07 |
| 0.169 | 7.59E-07 |
| 0.052 | 7.79E-07 |
| 0.197 | 7.84E-07 |
| 0.285 | 7.98E-07 |
| 0.161 | 8.05E-07 |
| 0.033 | 8.52E-07 |
| 0.111 | 8.92E-07 |
| 0.701 | 9.11E-07 |
| 0.292 | 9.44E-07 |
| 0.033 | 9.61E-07 |
| 0.04 | 1.01E-06 |
| 0.026 | 1.01E-06 |
| 0.158 | 1.02E-06 |
| 0.059 | 1.04E-06 |
| 0.199 | 1.05E-06 |
| 0.04 | 1.07E-06 |
| 0.969 | 1.11E-06 |
| 0.139 | 1.20E-06 |
| 0.072 | 1.25E-06 |
| 0.101 | 1.29E-06 |
| 0.028 | 1.38E-06 |
| 0.113 | 1.50E-06 |
| 0.035 | 1.51E-06 |
| 0.086 | 1.55E-06 |
| 0.097 | 1.59E-06 |
| 0.03 | 1.60E-06 |
| 0.029 | 1.63E-06 |
| 0.114 | 1.65E-06 |
| 0.063 | 1.76E-06 |
| 0.255 | 1.80E-06 |
| 0.107 | 1.91E-06 |
| 0.059 | 1.96E-06 |
| 0.04 | 2.01E-06 |
| 0.07 | 2.20E-06 |
| 0.093 | 2.51E-06 |
| 0.032 | 2.86E-06 |
| 0.044 | 2.87E-06 |
| 0.123 | 2.93E-06 |
| 0.072 | 2.94E-06 |
| 0.046 | 2.96E-06 |
| 0.036 | 3.00E-06 |
| 0.033 | 3.09E-06 |
| 0.141 | 3.33E-06 |
| 0.085 | 3.75E-06 |
| 0.053 | 3.80E-06 |
| 0.167 | 3.88E-06 |
| 0.114 | 4.21E-06 |
| 0.119 | 4.27E-06 |
| 0.101 | 4.28E-06 |
| 0.088 | 4.42E-06 |
| 0.061 | 4.50E-06 |
| 0.026 | 4.65E-06 |
| 0.032 | 4.69E-06 |
| 0.12 | 5.10E-06 |
| 0.153 | 5.37E-06 |
| 0.031 | 5.46E-06 |
| 0.487 | 5.72E-06 |
| 0.041 | 5.76E-06 |
| 0.146 | 5.77E-06 |
| 0.04 | 5.83E-06 |
| 0.025 | 5.91E-06 |
| 0.072 | 5.91E-06 |
| 0.196 | 6.42E-06 |
| 0.124 | 6.59E-06 |
| 0.027 | 6.89E-06 |
| 0.075 | 7.43E-06 |
| 0.057 | 7.79E-06 |
| 0.13 | 7.89E-06 |
| 0.156 | 7.93E-06 |
| 0.049 | 8.19E-06 |
| 0.056 | 8.55E-06 |
| 0.057 | 8.61E-06 |
| 0.058 | 8.63E-06 |
| 0.042 | 8.80E-06 |
| 0.103 | 8.89E-06 |
| 0.065 | 8.95E-06 |
| 0.074 | 9.26E-06 |
| 0.064 | 9.55E-06 |
| 0.125 | 1.21E-05 |
| 0.087 | 1.24E-05 |
| 0.129 | 1.25E-05 |
| 0.119 | 1.27E-05 |
| 0.051 | 1.28E-05 |
| 0.052 | 1.30E-05 |
| 0.097 | 1.31E-05 |
| 0.527 | 1.32E-05 |
| 0.191 | 1.39E-05 |
| 0.222 | 1.42E-05 |
| 0.249 | 1.51E-05 |
| 0.229 | 1.55E-05 |
| 0.103 | 1.60E-05 |
| 0.102 | 1.69E-05 |
| 0.258 | 1.72E-05 |
| 0.295 | 1.74E-05 |
| 0.533 | 1.81E-05 |
| 0.164 | 1.82E-05 |
| 0.204 | 1.85E-05 |
| 0.033 | 1.88E-05 |
| 0.078 | 1.90E-05 |
| 0.14 | 1.97E-05 |
| 0.301 | 1.98E-05 |
| 0.089 | 2.08E-05 |
| 0.061 | 2.15E-05 |
| 0.099 | 2.18E-05 |
| 0.153 | 2.20E-05 |
| 0.178 | 2.21E-05 |
| 0.09 | 2.49E-05 |
| 0.038 | 2.59E-05 |
| 0.125 | 2.66E-05 |
| 0.025 | 2.92E-05 |
| 0.202 | 2.99E-05 |
| 0.378 | 3.00E-05 |
| 0.138 | 3.03E-05 |
| 0.032 | 3.11E-05 |
| 0.101 | 3.22E-05 |
| 0.065 | 3.37E-05 |
| 0.071 | 3.42E-05 |
| 0.094 | 3.50E-05 |
| 0.141 | 3.63E-05 |
| 0.194 | 3.69E-05 |
| 0.167 | 3.79E-05 |
| 0.056 | 3.84E-05 |
| 0.037 | 3.92E-05 |
| 0.062 | 4.02E-05 |
| 0.076 | 4.10E-05 |
| 0.129 | 4.16E-05 |
| 0.126 | 4.41E-05 |
| 0.321 | 4.41E-05 |
| 0.226 | 4.48E-05 |
| 0.155 | 4.66E-05 |
| 0.045 | 4.67E-05 |
| 0.066 | 4.68E-05 |
| 0.204 | 4.71E-05 |
| 0.203 | 4.82E-05 |
| 0.157 | 4.95E-05 |
| 0.03 | 5.25E-05 |
| 0.156 | 5.37E-05 |
| 0.104 | 5.53E-05 |
| 0.066 | 5.62E-05 |
| 0.294 | 5.70E-05 |
| 0.075 | 5.74E-05 |
| 0.027 | 5.74E-05 |
| 0.155 | 5.83E-05 |
| 0.173 | 6.04E-05 |
| 0.064 | 6.32E-05 |
| 0.032 | 6.42E-05 |
| 0.042 | 6.78E-05 |
| 0.107 | 6.92E-05 |
| 0.077 | 7.28E-05 |
| 0.063 | 7.36E-05 |
| 0.049 | 7.62E-05 |
| 0.156 | 7.66E-05 |
| 0.059 | 7.86E-05 |
| 0.115 | 7.94E-05 |
| 0.052 | 8.05E-05 |
| 0.132 | 8.09E-05 |
| 0.12 | 8.55E-05 |
| 0.06 | 8.59E-05 |
| 0.052 | 8.77E-05 |
| 0.053 | 8.84E-05 |
| 0.035 | 9.62E-05 |
| 0.492 | 0.000100431 |
| 0.157 | 0.000104013 |
| 0.126 | 0.000106265 |
| 0.11 | 0.000106347 |
| 0.292 | 0.000106723 |
| 0.068 | 0.000107934 |
| 0.063 | 0.000109293 |
| 0.045 | 0.000111048 |
| 0.079 | 0.000113227 |
| 0.09 | 0.000115553 |
| 0.075 | 0.000115688 |
| 0.168 | 0.000115789 |
| 0.185 | 0.000117943 |
| 0.09 | 0.00012173 |
| 0.304 | 0.000123115 |
| 0.305 | 0.000125466 |
| 0.04 | 0.000131469 |
| 0.169 | 0.0001317 |
| 0.054 | 0.000137092 |
| 0.146 | 0.00013915 |
| 0.031 | 0.000143518 |
| 0.265 | 0.000145152 |
| 0.029 | 0.00014569 |
| 0.211 | 0.000150414 |
| 0.222 | 0.000154773 |
| 0.036 | 0.000165689 |
| 0.104 | 0.000170815 |
| 0.031 | 0.000181234 |
| 0.128 | 0.000210404 |
| 0.168 | 0.000216813 |
| 0.077 | 0.000227287 |
| 0.269 | 0.000235986 |
| 0.264 | 0.000236946 |
| 0.169 | 0.000242362 |
| 0.071 | 0.000244552 |
| 0.073 | 0.000251406 |
| 0.141 | 0.000270339 |
| 0.063 | 0.000288823 |
| 0.191 | 0.000289299 |
| 0.097 | 0.000292783 |
| 0.096 | 0.000304139 |
| 0.117 | 0.000304243 |
| 0.195 | 0.000315507 |
| 0.079 | 0.000324638 |
| 0.122 | 0.000337024 |
| 0.192 | 0.000340022 |
| 0.045 | 0.000351632 |
| 0.048 | 0.000365617 |
| 0.032 | 0.00040767 |
| 0.034 | 0.000414546 |
| 0.326 | 0.000417535 |
| 0.086 | 0.000417876 |
| 0.736 | 0.000418766 |
| 0.066 | 0.000431596 |
| 0.359 | 0.000432316 |
| 0.152 | 0.000435667 |
| 0.044 | 0.000464157 |
| 0.101 | 0.000471147 |
| 0.068 | 0.000487764 |
| 0.167 | 0.000510137 |
| 0.34 | 0.000518598 |
| 0.033 | 0.000525585 |
| 0.225 | 0.000548378 |
| 0.093 | 0.000554861 |
| 0.048 | 0.000586958 |
| 0.155 | 0.000590876 |
| 0.033 | 0.000596955 |
| 0.051 | 0.000598196 |
| 0.131 | 0.000615759 |
| 0.044 | 0.000626344 |
| 0.265 | 0.000628248 |
| 0.063 | 0.000629237 |
| 0.183 | 0.00063167 |
| 0.03 | 0.000643232 |
| 0.508 | 0.000649538 |
| 0.038 | 0.000659013 |
| 0.028 | 0.000667766 |
| 0.047 | 0.00068092 |
| 0.198 | 0.000685558 |
| 0.111 | 0.000710823 |
| 0.027 | 0.000713058 |
| 0.086 | 0.000766121 |
| 0.032 | 0.000790355 |
| 0.231 | 0.000792459 |
| 0.229 | 0.000812539 |
| 0.147 | 0.000828544 |
| 0.2 | 0.000832903 |
| 0.064 | 0.000834191 |
| 0.048 | 0.000843459 |
| 0.311 | 0.000863669 |
| 0.067 | 0.000882728 |
| 0.077 | 0.000896851 |
| 0.067 | 0.000902977 |
| 0.082 | 0.000904781 |
| 0.029 | 0.000906698 |
| 0.041 | 0.000940166 |
| 0.08 | 0.000953002 |
| 0.308 | 0.001024622 |
| 0.232 | 0.001054284 |
| 0.04 | 0.00106631 |
| 0.147 | 0.001070158 |
| 0.098 | 0.001089029 |
| 0.405 | 0.001102338 |
| 0.125 | 0.001105349 |
| 0.045 | 0.001120365 |
| 0.137 | 0.001254003 |
| 0.054 | 0.001255164 |
| 0.032 | 0.001261574 |
| 0.081 | 0.001277048 |
| 0.073 | 0.001294147 |
| 0.183 | 0.001299673 |
| 0.07 | 0.00130783 |
| 0.077 | 0.001312058 |
| 0.052 | 0.001315621 |
| 0.066 | 0.001331684 |
| 0.031 | 0.001351043 |
| 0.119 | 0.001351999 |
| 0.159 | 0.001389042 |
| 0.579 | 0.001408333 |
| 0.125 | 0.001413287 |
| 0.032 | 0.001418334 |
| 0.044 | 0.001446428 |
| 0.076 | 0.001523397 |
| 0.088 | 0.001549552 |
| 0.064 | 0.001586827 |
| 0.034 | 0.001590738 |
| 0.033 | 0.001592763 |
| 0.062 | 0.001622889 |
| 0.071 | 0.001623302 |
| 0.073 | 0.001645364 |
| 0.048 | 0.001663799 |
| 0.157 | 0.001701114 |
| 0.311 | 0.001746509 |
| 0.342 | 0.001789042 |
| 0.056 | 0.001802695 |
| 0.175 | 0.001832827 |
| 0.093 | 0.001872053 |
| 0.068 | 0.001873633 |
| 0.057 | 0.001881855 |
| 0.027 | 0.001894474 |
| 0.091 | 0.001917251 |
| 0.13 | 0.001935899 |
| 0.484 | 0.001977944 |
| 0.036 | 0.002075591 |
| 0.228 | 0.002206604 |
| 0.2 | 0.002275262 |
| 0.069 | 0.002455708 |
| 0.057 | 0.002522228 |
| 0.368 | 0.002573405 |
| 0.166 | 0.002597044 |
| 0.179 | 0.002696791 |
| 0.061 | 0.00271298 |
| 0.053 | 0.002778135 |
| 0.062 | 0.002834121 |
| 0.049 | 0.002846765 |
| 0.036 | 0.002929154 |
| 0.063 | 0.002959286 |
| 0.132 | 0.003004044 |
| 0.06 | 0.003017731 |
| 0.074 | 0.003035592 |
| 0.101 | 0.003067745 |
| 0.635 | 0.003177289 |
| 0.108 | 0.003326377 |
| 0.092 | 0.00343239 |
| 0.073 | 0.003569371 |
| 0.043 | 0.003592761 |
| 0.047 | 0.003673782 |
| 0.067 | 0.003795515 |
| 0.133 | 0.003817167 |
| 0.111 | 0.003935785 |
| 0.147 | 0.003950847 |
| 0.066 | 0.003971653 |
| 0.1 | 0.003992026 |
| 0.058 | 0.004259398 |
| 0.089 | 0.004347205 |
| 0.027 | 0.004412589 |
| 0.098 | 0.004539397 |
| 0.205 | 0.004699131 |
| 0.146 | 0.004837795 |
| 0.212 | 0.004841984 |
| 0.035 | 0.004901557 |
| 0.205 | 0.005006111 |
| 0.051 | 0.005013748 |
| 0.429 | 0.005098774 |
| 0.068 | 0.005327989 |
| 0.059 | 0.005383729 |
| 0.057 | 0.005490953 |
| 0.072 | 0.005493851 |
| 0.103 | 0.005519399 |
| 0.051 | 0.005671808 |
| 0.205 | 0.006053179 |
| 0.12 | 0.00614095 |
| 0.066 | 0.006141908 |
| 0.099 | 0.006189354 |
| 0.287 | 0.006421977 |
| 0.056 | 0.006430843 |
| 0.346 | 0.006898352 |
| 0.068 | 0.006959706 |
| 0.216 | 0.007057035 |
| 0.074 | 0.007260943 |
| 0.367 | 0.007477946 |
| 0.17 | 0.00780383 |
| 0.087 | 0.007862819 |
| 0.028 | 0.00797275 |
| 0.149 | 0.008154748 |
| 0.049 | 0.00817715 |
| 0.048 | 0.008183833 |
| 0.116 | 0.00824488 |
| 0.052 | 0.008336629 |
| 0.164 | 0.008494859 |
| 0.11 | 0.008497173 |
| 0.164 | 0.008618132 |
| 0.124 | 0.008824073 |
| 0.26 | 0.008898248 |
| 0.038 | 0.008940047 |
| 0.044 | 0.009154035 |
| 0.155 | 0.009216473 |
| 0.055 | 0.009463387 |
| 0.145 | 0.009633478 |
| 0.032 | 0.009683149 |
| 0.105 | 0.009748376 |
| 0.289 | 0.00982646 |
| 0.123 | 0.010113874 |
| 0.3 | 0.01028586 |
| 0.077 | 0.010399808 |
| 0.967 | 0.010561294 |
| 0.175 | 0.010608298 |
| 0.058 | 0.010988022 |
| 0.057 | 0.011349198 |
| 0.062 | 0.011685319 |
| 0.056 | 0.011700837 |
| 0.262 | 0.011800837 |
| 0.079 | 0.011837386 |
| 0.099 | 0.011871468 |
| 0.05 | 0.011950078 |
| 0.325 | 0.012037231 |
| 0.096 | 0.012048724 |
| 0.22 | 0.01205009 |
| 0.084 | 0.012084977 |
| 0.037 | 0.012156852 |
| 0.076 | 0.012288 |
| 0.031 | 0.012426975 |
| 0.077 | 0.012900429 |
| 0.046 | 0.013619689 |
| 0.052 | 0.01396704 |
| 0.03 | 0.014096582 |
| 0.051 | 0.014299326 |
| 0.051 | 0.015353412 |
| 0.162 | 0.015637117 |
| 0.249 | 0.015730777 |
| 0.246 | 0.016017346 |
| 0.052 | 0.016236362 |
| 0.035 | 0.016321312 |
| 0.057 | 0.016712093 |
| 0.212 | 0.016768247 |
| 0.105 | 0.016884232 |
| 0.101 | 0.018835605 |
| 0.069 | 0.019385196 |
| 0.112 | 0.019389329 |
| 0.117 | 0.019643634 |
| 0.179 | 0.019663132 |
| 0.065 | 0.020298873 |
| 0.036 | 0.020669748 |
| 0.058 | 0.020907162 |
| 0.033 | 0.021260191 |
| 0.038 | 0.021769527 |
| 0.087 | 0.022503184 |
| 0.131 | 0.022807386 |
| 0.068 | 0.023766386 |
| 0.169 | 0.024063188 |
| 0.123 | 0.024248771 |
| 0.067 | 0.02560409 |
| 0.037 | 0.025905143 |
| 0.19 | 0.025956027 |
| 0.132 | 0.026477224 |
| 0.044 | 0.026599265 |
| 0.029 | 0.026676741 |
| 0.034 | 0.027069679 |
| 0.194 | 0.027512968 |
| 0.09 | 0.027600195 |
| 0.06 | 0.028132295 |
| 0.035 | 0.028571033 |
| 0.363 | 0.0291773 |
| 0.043 | 0.029322333 |
| 0.119 | 0.029419308 |
| 0.084 | 0.029740309 |
| 0.218 | 0.031085273 |
| 0.204 | 0.031179922 |
| 0.057 | 0.031978855 |
| 0.125 | 0.032742771 |
| 0.056 | 0.032775357 |
| 0.065 | 0.033214657 |
| 0.081 | 0.033371135 |
| 0.41 | 0.033567616 |
| 0.083 | 0.034223794 |
| 0.041 | 0.03447069 |
| 0.256 | 0.034711052 |
| 0.175 | 0.035264531 |
| 0.12 | 0.035431069 |
| 0.056 | 0.038013183 |
| 0.077 | 0.038257237 |
| 0.172 | 0.039511258 |
| 0.162 | 0.040103251 |
| 0.288 | 0.040421188 |
| 0.037 | 0.042139137 |
| 0.29 | 0.042326463 |
| 0.059 | 0.043601499 |
| 0.039 | 0.04380178 |
| 0.038 | 0.044065341 |
| 0.066 | 0.044852847 |
| 0.088 | 0.04535613 |
| 0.169 | 0.045684748 |
| 0.045 | 0.047139041 |
| 0.092 | 0.049549751 |
| 0.086 | 0.050292167 |
| 0.208 | 0.050517376 |
| 0.056 | 0.051629975 |
| 0.057 | 0.05363333 |
| 0.134 | 0.054592917 |
| 0.133 | 0.055595832 |
| 0.249 | 0.0582441 |
| 0.327 | 0.058531444 |
| 0.128 | 0.058846045 |
| 0.121 | 0.058858399 |
| 0.412 | 0.059155994 |
| 0.219 | 0.059423964 |
| 0.053 | 0.060163646 |
| 0.039 | 0.061336195 |
| 0.07 | 0.063161851 |
| 0.206 | 0.064309467 |
| 0.114 | 0.066324702 |
| 0.148 | 0.067937779 |
| 0.235 | 0.068412435 |
| 0.143 | 0.068580059 |
| 0.086 | 0.06926733 |
| 0.05 | 0.070074716 |
| 0.128 | 0.070199922 |
| 0.394 | 0.07029881 |
| 0.09 | 0.070972286 |
| 0.166 | 0.071888125 |
| 0.133 | 0.071919683 |
| 0.136 | 0.074059064 |
| 0.197 | 0.075074671 |
| 0.134 | 0.075689686 |
| 0.063 | 0.078897657 |
| 0.458 | 0.07973888 |
| 0.233 | 0.080805596 |
| 0.05 | 0.081421505 |
| 0.116 | 0.08359635 |
| 0.462 | 0.084788433 |
| 0.086 | 0.08495322 |
| 0.211 | 0.085098897 |
| 0.103 | 0.085396268 |
| 0.573 | 0.090313173 |
| 0.279 | 0.090667397 |
| 0.088 | 0.091073354 |
| 0.036 | 0.091147227 |
| 0.235 | 0.091259795 |
| 0.039 | 0.093806431 |
| 0.034 | 0.096261316 |
| 0.031 | 0.096262315 |
| 0.1 | 0.097567256 |
| 0.064 | 0.1052384 |
| 0.166 | 0.107201879 |
| 0.363 | 0.109298932 |
| 0.247 | 0.110222022 |
| 0.092 | 0.11036295 |
| 0.073 | 0.111114437 |
| 0.041 | 0.112263348 |
| 0.069 | 0.112537469 |
| 0.303 | 0.112990116 |
| 0.167 | 0.114820628 |
| 0.121 | 0.115385404 |
| 0.126 | 0.119256531 |
| 0.057 | 0.120805357 |
| 0.091 | 0.120952274 |
| 0.083 | 0.121642885 |
| 0.087 | 0.121682441 |
| 0.035 | 0.123869185 |
| 0.057 | 0.128995734 |
| 0.038 | 0.129388911 |
| 0.058 | 0.130014343 |
| 0.03 | 0.130877434 |
| 0.118 | 0.13278053 |
| 0.059 | 0.132798551 |
| 0.049 | 0.13296643 |
| 0.197 | 0.136377491 |
| 0.231 | 0.140292415 |
| 0.062 | 0.142445354 |
| 0.141 | 0.142833665 |
| 0.057 | 0.145597216 |
| 0.353 | 0.147541511 |
| 0.167 | 0.149203792 |
| 0.052 | 0.150778924 |
| 0.045 | 0.152127234 |
| 0.127 | 0.153150958 |
| 0.171 | 0.15487575 |
| 0.139 | 0.156170121 |
| 0.295 | 0.156612702 |
| 0.835 | 0.159928539 |
| 0.095 | 0.162917473 |
| 0.068 | 0.166457154 |
| 0.291 | 0.166655377 |
| 0.105 | 0.167061897 |
| 0.069 | 0.167119751 |
| 0.158 | 0.169914786 |
| 0.072 | 0.172345787 |
| 0.254 | 0.174035478 |
| 0.107 | 0.174674771 |
| 0.175 | 0.176411523 |
| 0.082 | 0.176757546 |
| 0.05 | 0.177314103 |
| 0.102 | 0.178076958 |
| 0.034 | 0.181125317 |
| 0.075 | 0.182097067 |
| 0.069 | 0.185010171 |
| 0.087 | 0.185365593 |
| 0.088 | 0.185555763 |
| 0.149 | 0.187500107 |
| 0.181 | 0.1893937 |
| 0.201 | 0.191817981 |
| 0.056 | 0.193990478 |
| 0.05 | 0.195764018 |
| 0.225 | 0.196227157 |
| 0.148 | 0.200731807 |
| 0.05 | 0.203270905 |
| 0.034 | 0.204373165 |
| 0.106 | 0.205468582 |
| 0.087 | 0.205684332 |
| 0.059 | 0.206189968 |
| 0.173 | 0.214317545 |
| 0.037 | 0.219155054 |
| 0.076 | 0.220733731 |
| 0.04 | 0.2237917 |
| 0.035 | 0.226883109 |
| 0.043 | 0.229682096 |
| 0.092 | 0.230314788 |
| 0.194 | 0.231921395 |
| 0.051 | 0.232805213 |
| 0.04 | 0.233120677 |
| 0.031 | 0.240498655 |
| 0.164 | 0.241308258 |
| 0.052 | 0.242536344 |
| 0.039 | 0.244897962 |
| 0.067 | 0.256080285 |
| 0.127 | 0.25644365 |
| 0.072 | 0.263078284 |
| 0.112 | 0.263452874 |
| 0.12 | 0.271257773 |
| 0.053 | 0.274090895 |
| 0.077 | 0.284770099 |
| 0.055 | 0.290373607 |
| 0.076 | 0.304241562 |
| 0.241 | 0.306163404 |
| 0.037 | 0.308240757 |
| 0.173 | 0.319252193 |
| 0.041 | 0.322492586 |
| 0.124 | 0.325642814 |
| 0.104 | 0.33388653 |
| 0.115 | 0.335773028 |
| 0.09 | 0.336434275 |
| 0.032 | 0.336792273 |
| 0.161 | 0.337889369 |
| 0.06 | 0.340319207 |
| 0.086 | 0.34045826 |
| 0.074 | 0.34157917 |
| 0.119 | 0.34671855 |
| 0.063 | 0.347913156 |
| 0.038 | 0.350734025 |
| 0.191 | 0.351623914 |
| 0.064 | 0.35172105 |
| 0.233 | 0.35578583 |
| 0.298 | 0.358856861 |
| 0.046 | 0.360077642 |
| 0.074 | 0.361251893 |
| 0.036 | 0.362787187 |
| 0.055 | 0.366492964 |
| 0.066 | 0.369099399 |
| 0.067 | 0.369463116 |
| 0.048 | 0.37191161 |
| 0.064 | 0.380465793 |
| 0.072 | 0.408311031 |
| 0.152 | 0.409994069 |
| 0.116 | 0.412630806 |
| 0.102 | 0.416470719 |
| 0.061 | 0.433770983 |
| 0.071 | 0.434941391 |
| 0.053 | 0.441214696 |
| 0.121 | 0.446462774 |
| 0.057 | 0.454147417 |
| 0.397 | 0.455350534 |
| 0.084 | 0.471656976 |
| 0.062 | 0.477638566 |
| 0.292 | 0.480036979 |
| 0.054 | 0.489358144 |
| 0.043 | 0.492782986 |
| 0.14 | 0.495287542 |
| 0.093 | 0.503201963 |
| 0.059 | 0.503315066 |
| 0.036 | 0.519176782 |
| 0.051 | 0.52031547 |
| 0.078 | 0.530592755 |
| 0.061 | 0.560405616 |
| 0.316 | 0.564257355 |
| 0.087 | 0.577978158 |
| 0.143 | 0.582096865 |
| 0.104 | 0.594037083 |
| 0.094 | 0.603841525 |
| 0.069 | 0.610994868 |
| 0.051 | 0.622464736 |
| 0.083 | 0.622559039 |
| 0.104 | 0.624767568 |
| 0.345 | 0.639272559 |
| 0.053 | 0.650271588 |
| 0.076 | 0.659812055 |
| 0.42 | 0.663333148 |
| 0.238 | 0.677902645 |
| 0.084 | 0.686876222 |
| 0.51 | 0.707743661 |
| 0.129 | 0.723517534 |
| 0.056 | 0.731188809 |
| 0.11 | 0.732600037 |
| 0.077 | 0.73300718 |
| 0.083 | 0.743045297 |
| 0.054 | 0.745136859 |
| 0.092 | 0.774577766 |
| 0.117 | 0.777394623 |
| 0.063 | 0.779394645 |
| 0.066 | 0.794991306 |
| 0.108 | 0.825989839 |
| 0.104 | 0.826222161 |
| 0.35 | 0.827663758 |
| 0.135 | 0.830800748 |
| 0.152 | 0.840069608 |
| 0.037 | 0.843625219 |
| 0.089 | 0.859768278 |
| 0.054 | 0.876998943 |
| 0.039 | 0.884056263 |
| 0.069 | 0.89046312 |
| 0.155 | 0.891277819 |
| 0.084 | 0.892520161 |
| 0.119 | 0.91403713 |
| 0.054 | 0.930356748 |
| 0.08 | 0.943797553 |
| 0.06 | 0.948194453 |
| 0.352 | 0.987298355 |
| 0.096 | 1 |
| 0.034 | 1 |
| 0.081 | 1 |
| 0.066 | 1 |
| 0.115 | 1 |
| 0.136 | 1 |
| 0.06 | 1 |
| 0.124 | 1 |
| 0.073 | 1 |
| 0.038 | 1 |
| 0.054 | 1 |
| 0.048 | 1 |
| 0.084 | 1 |
| 0.033 | 1 |
| 0.087 | 1 |
| 0.038 | 1 |
| 0.118 | 1 |
| 0.038 | 1 |
| 0.096 | 1 |
| 0.108 | 1 |
| 0.082 | 1 |
| 0.222 | 1 |
| 0.174 | 1 |
| 0.052 | 1 |
| 0.055 | 1 |
| 0.141 | 1 |
| 0.135 | 1 |
| 0.124 | 1 |
| 0.134 | 1 |
| 0.056 | 1 |
| 0.28 | 1 |
| 0.036 | 1 |
| 0.174 | 1 |
| 0.062 | 1 |
| 0.033 | 1 |
| 0.073 | 1 |
| 0.062 | 1 |
| 0.506 | 1 |
| 0.146 | 1 |
| 0.035 | 1 |
| 0.041 | 1 |
| 0.149 | 1 |
| 0.166 | 1 |
| 0.642 | 1 |
| 0.04 | 1 |
| 0.174 | 1 |
| 0.163 | 1 |
| 0.071 | 1 |
| 0.046 | 1 |
| 0.062 | 1 |
| 0.059 | 1 |
| 0.069 | 1 |
| 0.056 | 1 |
| 0.118 | 1 |
| 0.146 | 1 |
| 0.092 | 1 |
| 0.033 | 1 |
| 0.061 | 1 |
| 0.199 | 1 |
| 0.048 | 1 |
| 0.074 | 1 |
| 0.096 | 1 |
| 0.065 | 1 |
| 0.093 | 1 |
| 0.045 | 1 |
| 0.125 | 1 |
| 0.078 | 1 |
| 0.198 | 1 |
| 0.124 | 1 |
| 0.115 | 1 |
| 0.149 | 1 |
| 0.097 | 1 |
| 0.315 | 1 |
| 0.05 | 1 |
| 0.052 | 1 |
| 0.104 | 1 |
| 0.038 | 1 |
| 0.102 | 1 |
| 0.125 | 1 |
| 0.189 | 1 |
| 0.149 | 1 |
| 0.113 | 1 |
| 0.073 | 1 |
| 0.256 | 1 |
| 0.145 | 1 |
| 0.053 | 1 |
| 0.178 | 1 |
| 0.087 | 1 |
| 0.071 | 1 |
| 0.041 | 1 |
| 0.052 | 1 |
| 0.071 | 1 |
| 0.033 | 1 |
| 0.119 | 1 |
| 0.117 | 1 |
| 0.106 | 1 |
| 0.048 | 1 |
| 0.069 | 1 |
| 0.068 | 1 |
| 0.062 | 1 |
| 0.062 | 1 |
| 0.101 | 1 |
| 0.05 | 1 |
| 0.053 | 1 |
| 0.04 | 1 |
| 0.062 | 1 |
| 0.366 | 1 |
| 0.251 | 1 |
| 0.107 | 1 |
| 0.055 | 1 |
| 0.039 | 1 |
| 0.071 | 1 |
| 0.141 | 1 |
| 0.035 | 1 |
| 0.049 | 1 |
| 0.164 | 1 |
| 0.033 | 1 |
| 0.058 | 1 |
| 0.117 | 1 |
| 0.041 | 1 |
| 0.06 | 1 |
| 0.042 | 1 |
| 0.225 | 1 |
| 0.156 | 1 |
| 0.058 | 1 |
| 0.124 | 1 |
| 0.059 | 1 |
| 0.208 | 1 |
| 0.229 | 1 |
| 0.061 | 1 |
| 0.108 | 1 |
| 0.075 | 1 |
| 0.1 | 1 |
| 0.086 | 1 |
| 0.061 | 1 |
| 0.045 | 1 |
| 0.045 | 1 |
| 0.039 | 1 |
| 0.045 | 1 |
| 0.116 | 1 |
| 0.034 | 1 |
| 0.161 | 1 |
| 0.109 | 1 |
| 0.12 | 1 |
| 0.055 | 1 |
| 0.072 | 1 |
| 0.069 | 1 |
| 0.073 | 1 |
| 0.065 | 1 |
| 0.128 | 1 |
| 0.069 | 1 |
| 0.393 | 1 |
| 0.089 | 1 |
| 0.323 | 1 |
| 0.068 | 1 |
| 0.057 | 1 |
| 0.061 | 1 |
| 0.112 | 1 |
| 0.177 | 1 |
| 0.045 | 1 |
| 0.049 | 1 |
| 0.091 | 1 |
| 0.115 | 1 |
| 0.129 | 1 |
| 0.269 | 1 |
| 0.121 | 1 |
| 0.115 | 1 |
| 0.061 | 1 |
| 0.041 | 1 |
| 0.077 | 1 |
| 0.042 | 1 |
| 0.053 | 1 |
| 0.132 | 1 |
| 0.185 | 1 |
| 0.217 | 1 |
| 0.163 | 1 |
| 0.068 | 1 |
| 0.317 | 1 |
| 0.06 | 1 |
| 0.168 | 1 |
| 0.077 | 1 |
| 0.057 | 1 |
| 0.053 | 1 |
| 0.068 | 1 |
| 0.091 | 1 |
| 0.073 | 1 |
| 0.057 | 1 |
| 0.092 | 1 |
| 0.041 | 1 |
| 0.05 | 1 |
| 0.15 | 1 |
| 0.356 | 1 |
| 0.051 | 1 |
| 0.051 | 1 |
| 0.088 | 1 |
| 0.076 | 1 |
| 0.118 | 1 |
| 0.062 | 1 |
| 0.065 | 1 |
| 0.175 | 1 |
| 0.066 | 1 |
| 0.113 | 1 |
| 0.165 | 1 |
| 0.077 | 1 |
| 0.129 | 1 |
| 0.137 | 1 |
| 0.267 | 1 |
| 0.035 | 1 |
| 0.064 | 1 |
| 0.035 | 1 |
| 0.154 | 1 |
| 0.077 | 1 |
| 0.106 | 1 |
| 0.267 | 1 |
| 0.197 | 1 |
| 0.125 | 1 |
| 0.039 | 1 |
| 0.061 | 1 |
| 0.077 | 1 |
| 0.064 | 1 |
| 0.044 | 1 |
| 0.121 | 1 |
| 0.057 | 1 |
| 0.053 | 1 |
| 0.06 | 1 |
| 0.167 | 1 |
| 0.189 | 1 |
| 0.332 | 1 |
| 0.124 | 1 |
| 0.085 | 1 |
| 0.071 | 1 |
| 0.19 | 1 |
| 0.272 | 1 |
| 0.036 | 1 |
| 0.046 | 1 |
| 0.093 | 1 |
| 0.084 | 1 |
| 0.125 | 1 |
| 0.234 | 1 |
| 0.162 | 1 |
| 0.209 | 1 |
| 0.055 | 1 |
| 0.076 | 1 |
| 0.204 | 1 |
| 0.04 | 1 |
| 0.088 | 1 |
| 0.07 | 1 |
| 0.086 | 1 |
| 0.123 | 1 |
| 0.055 | 1 |
| 0.147 | 1 |
| 0.08 | 1 |
| 0.062 | 1 |
| 0.131 | 1 |
| 0.107 | 1 |
| 0.108 | 1 |
| 0.265 | 1 |
| 0.067 | 1 |
| 0.054 | 1 |
| 0.134 | 1 |
| 0.082 | 1 |
| 0.166 | 1 |
| 0.07 | 1 |
| 0.217 | 1 |
| 0.076 | 1 |
| 0.065 | 1 |
| 0.179 | 1 |
| 0.295 | 1 |
| 0.098 | 1 |
| 0.238 | 1 |
| 0.061 | 1 |
| 0.073 | 1 |
| 0.051 | 1 |
| 0.137 | 1 |
| 0.255 | 1 |
| 0.052 | 1 |
| 0.199 | 1 |
| 0.134 | 1 |
| 0.041 | 1 |
| 0.157 | 1 |
| 0.037 | 1 |
| 0.221 | 1 |
| 0.062 | 1 |
| 0.166 | 1 |
| 0.068 | 1 |
| 0.056 | 1 |
| 0.187 | 1 |
| 0.249 | 1 |
| 0.04 | 1 |
| 0.055 | 1 |
| 0.132 | 1 |
| 0.112 | 1 |
| 0.038 | 1 |
| 0.126 | 1 |
| 0.058 | 1 |
| 0.32 | 1 |
| 0.051 | 1 |
| 0.041 | 1 |
| 0.138 | 1 |
| 0.1 | 1 |
| 0.061 | 1 |
| 0.165 | 1 |
| 0.045 | 1 |
| 0.139 | 1 |
| 0.071 | 1 |
| 0.241 | 1 |
| 0.331 | 1 |
| 0.062 | 1 |
| 0.087 | 1 |
| 0.04 | 1 |
| 0.082 | 1 |
| 0.103 | 1 |
| 0.035 | 1 |
| 0.136 | 1 |
| 0.097 | 1 |
| 0.051 | 1 |
| 0.053 | 1 |
| 0.109 | 1 |
| 0.069 | 1 |
| 0.128 | 1 |
| 0.202 | 1 |
| 0.05 | 1 |
| 0.066 | 1 |
| 0.042 | 1 |
| 0.147 | 1 |
| 0.054 | 1 |
| 0.049 | 1 |
| 0.04 | 1 |
| 0.368 | 1 |
| 0.078 | 1 |
| 0.101 | 1 |
| 0.201 | 1 |
| 0.068 | 1 |
| 0.28 | 1 |
| 0.055 | 1 |
| 0.142 | 1 |
| 0.038 | 1 |
| 0.25 | 1 |
| 0.225 | 1 |
| 0.146 | 1 |
| 0.114 | 1 |
| 0.056 | 1 |
| 0.162 | 1 |
| 0.155 | 1 |
| 0.07 | 1 |
| 0.122 | 1 |
| 0.075 | 1 |
| 0.064 | 1 |
| 0.063 | 1 |
| 0.203 | 1 |
| 0.055 | 1 |
| 0.152 | 1 |
| 0.073 | 1 |
| 0.18 | 1 |
| 0.052 | 1 |
| 0.079 | 1 |
| 0.104 | 1 |
| 0.092 | 1 |
| 0.152 | 1 |
| 0.426 | 1 |
| 0.053 | 1 |
| 0.054 | 1 |
| 0.055 | 1 |
| 0.07 | 1 |
| 0.105 | 1 |
| 0.255 | 1 |
| 0.039 | 1 |
| 0.071 | 1 |
| 0.288 | 1 |
| 0.088 | 1 |
| 0.154 | 1 |
| 0.065 | 1 |
| 0.067 | 1 |
| 0.294 | 1 |
| 0.057 | 1 |
| 0.063 | 1 |
| 0.08 | 1 |
| 0.133 | 1 |
| 0.112 | 1 |
| 0.055 | 1 |
| 0.049 | 1 |
| 0.04 | 1 |
| 0.062 | 1 |
| 0.058 | 1 |
| 0.108 | 1 |
| 0.038 | 1 |
| 0.084 | 1 |
| 0.078 | 1 |
| 0.06 | 1 |
| 0.061 | 1 |
| 0.091 | 1 |
| 0.094 | 1 |
| 0.167 | 1 |
| 0.073 | 1 |
| 0.137 | 1 |
| 0.185 | 1 |
| 0.051 | 1 |
| 0.039 | 1 |
| 0.084 | 1 |
| 0.127 | 1 |
| 0.054 | 1 |
| 0.28 | 1 |
| 0.055 | 1 |
| 0.052 | 1 |
| 0.246 | 1 |
| 0.103 | 1 |
| 0.038 | 1 |
| 0.152 | 1 |
| 0.115 | 1 |
| 0.041 | 1 |
| 0.045 | 1 |
| 0.088 | 1 |
| 0.097 | 1 |
| 0.132 | 1 |
| 0.084 | 1 |
| 0.076 | 1 |
| 0.101 | 1 |
| 0.042 | 1 |
| 0.246 | 1 |
| 0.071 | 1 |
| 0.041 | 1 |
| 0.357 | 1 |
| 0.071 | 1 |
| 0.084 | 1 |
| 0.094 | 1 |
| 0.07 | 1 |
| 0.048 | 1 |
| 0.101 | 1 |
| 0.134 | 1 |
| 0.039 | 1 |
| 0.044 | 1 |
| 0.1 | 1 |
| 0.165 | 1 |
| 0.067 | 1 |
| 0.071 | 1 |
| 0.086 | 1 |
| 0.074 | 1 |
| 0.041 | 1 |
| 0.073 | 1 |
| 0.085 | 1 |
| 0.109 | 1 |
| 0.04 | 1 |
| 0.038 | 1 |
| 0.092 | 1 |
| 0.051 | 1 |
| 0.06 | 1 |
| 0.065 | 1 |
| 0.28 | 1 |
| 0.047 | 1 |
| 0.064 | 1 |
| 0.057 | 1 |
| 0.149 | 1 |
| 0.048 | 1 |
| 0.056 | 1 |
| 0.046 | 1 |
| 0.296 | 1 |
| 0.078 | 1 |
| 0.126 | 1 |
| 0.06 | 1 |
| 0.06 | 1 |
| 0.124 | 1 |
| 0.071 | 1 |
| 0.085 | 1 |
| 0.159 | 1 |
| 0.241 | 1 |
| 0.217 | 1 |
| 0.114 | 1 |
| 0.122 | 1 |
| 0.051 | 1 |
| 0.189 | 1 |
| 0.047 | 1 |
| 0.096 | 1 |
| 0.112 | 1 |
| 0.071 | 1 |
| 0.076 | 1 |
| 0.052 | 1 |
| 0.079 | 1 |
| 0.048 | 1 |
| 0.076 | 1 |
| 0.038 | 1 |
| 0.085 | 1 |
| 0.176 | 1 |
| 0.048 | 1 |
| 0.109 | 1 |
| 0.21 | 1 |
| 0.04 | 1 |
| 0.065 | 1 |
| 0.623 | 1 |
| 0.082 | 1 |
| 0.085 | 1 |
| 0.121 | 1 |
| 0.057 | 1 |
| 0.052 | 1 |
| 0.157 | 1 |
| 0.096 | 1 |
| 0.05 | 1 |
| 0.4 | 1 |
| 0.065 | 1 |
| 0.108 | 1 |
| 0.057 | 1 |
| 0.116 | 1 |
| 0.039 | 1 |
| 0.183 | 1 |
| 0.117 | 1 |
| 0.109 | 1 |
| 0.042 | 1 |
| 0.037 | 1 |
| 0.174 | 1 |
| 0.085 | 1 |
| 0.065 | 1 |
| 0.059 | 1 |
| 0.181 | 1 |
| 0.045 | 1 |
| 0.075 | 1 |
| 0.077 | 1 |
| 0.037 | 1 |
| 0.2 | 1 |
| 0.056 | 1 |
| 0.044 | 1 |
| 0.275 | 1 |
| 0.128 | 1 |
| 0.039 | 1 |
| 0.048 | 1 |
| 0.089 | 1 |
| 0.071 | 1 |
| 0.058 | 1 |
| 0.113 | 1 |
| 0.052 | 1 |
| 0.068 | 1 |
| 0.078 | 1 |
| 0.082 | 1 |
| 0.142 | 1 |
| 0.038 | 1 |
| 0.048 | 1 |
| 0.065 | 1 |
| 0.118 | 1 |
| 0.073 | 1 |
| 0.376 | 1 |
| 0.089 | 1 |
| 0.055 | 1 |
| 0.115 | 1 |
| 0.058 | 1 |
| 0.216 | 1 |
| 0.078 | 1 |
| 0.185 | 1 |
| 0.061 | 1 |
| 0.054 | 1 |
| 0.068 | 1 |
| 0.081 | 1 |
| 0.145 | 1 |
| 0.108 | 1 |
| 0.182 | 1 |
| 0.065 | 1 |
| 0.056 | 1 |
| 0.235 | 1 |
| 0.065 | 1 |
| 0.045 | 1 |
| 0.112 | 1 |
| 0.241 | 1 |
| 0.156 | 1 |
| 0.039 | 1 |
| 0.15 | 1 |
| 0.074 | 1 |
| 0.088 | 1 |
| 0.18 | 1 |
| 0.081 | 1 |
| 0.048 | 1 |
| 0.267 | 1 |
| 0.09 | 1 |
| 0.235 | 1 |
| 0.116 | 1 |
| 0.198 | 1 |
| 0.057 | 1 |
| 0.122 | 1 |
| 0.123 | 1 |
| 0.051 | 1 |
| 0.04 | 1 |
| 0.094 | 1 |
| 0.067 | 1 |
| 0.074 | 1 |
| 0.077 | 1 |
| 0.049 | 1 |
| 0.15 | 1 |
| 0.602 | 1 |
| 0.235 | 1 |
| 0.376 | 1 |
| 0.155 | 1 |
| 0.053 | 1 |
| 0.173 | 1 |
| 0.402 | 1 |
| 0.051 | 1 |
| 0.046 | 1 |
| 0.062 | 1 |
| 0.053 | 1 |
| 0.054 | 1 |
| 0.077 | 1 |
| 0.06 | 1 |
| 0.14 | 1 |
| 0.062 | 1 |
| 0.164 | 1 |
| 0.088 | 1 |
| 0.07 | 1 |
| 0.076 | 1 |
| 0.058 | 1 |
| 0.067 | 1 |
| 0.201 | 1 |
| 0.257 | 1 |
| 0.073 | 1 |
| 0.099 | 1 |
| 0.268 | 1 |
| 0.072 | 1 |
| 0.038 | 1 |
| 0.21 | 1 |
| 0.24 | 1 |
| 0.038 | 1 |
| 0.101 | 1 |
| 0.089 | 1 |
| 0.162 | 1 |
| 0.081 | 1 |
| 0.232 | 1 |
| 0.088 | 1 |
| 0.17 | 1 |
| 0.11 | 1 |
| 0.052 | 1 |
| 0.191 | 1 |
| 0.253 | 1 |
| 0.148 | 1 |
| 0.049 | 1 |
| 0.061 | 1 |
| 0.071 | 1 |
| 0.124 | 1 |
| 0.088 | 1 |
| 0.168 | 1 |
| 0.328 | 1 |
| 0.044 | 1 |
| 0.186 | 1 |
| 0.062 | 1 |
| 0.261 | 1 |
| 0.113 | 1 |
| 0.106 | 1 |
| 0.082 | 1 |
| 0.144 | 1 |
| 0.203 | 1 |
| 0.051 | 1 |
| 0.175 | 1 |
| 0.043 | 1 |
| 0.066 | 1 |
| 0.04 | 1 |
| 0.092 | 1 |
| 0.108 | 1 |
| 0.108 | 1 |
| 0.13 | 1 |
| 0.041 | 1 |
| 0.04 | 1 |
| 0.315 | 1 |
| 0.057 | 1 |
| 0.093 | 1 |
| 0.249 | 1 |
| 0.2 | 1 |
| 0.047 | 1 |
| 0.056 | 1 |
| 0.127 | 1 |
| 0.069 | 1 |
| 0.056 | 1 |
| 0.064 | 1 |
| 0.04 | 1 |
| 0.117 | 1 |
| 0.091 | 1 |
| 0.199 | 1 |
| 0.057 | 1 |
| 0.075 | 1 |
| 0.097 | 1 |
| 0.153 | 1 |
| 0.09 | 1 |
| 0.061 | 1 |
| 0.063 | 1 |
| 0.142 | 1 |
| 0.057 | 1 |
| 0.139 | 1 |
| 0.301 | 1 |
| 0.121 | 1 |
| 0.053 | 1 |
| 0.116 | 1 |
| 0.087 | 1 |
| 0.085 | 1 |
| 0.041 | 1 |
| 0.062 | 1 |
| 0.06 | 1 |
| 0.097 | 1 |
| 0.153 | 1 |
| 0.094 | 1 |
| 0.038 | 1 |
| 0.047 | 1 |
| 0.086 | 1 |
| 0.06 | 1 |
| 0.105 | 1 |
| 0.159 | 1 |
| 0.086 | 1 |
| 0.069 | 1 |
| 0.151 | 1 |
| 0.066 | 1 |
| 0.089 | 1 |
| 0.054 | 1 |
| 0.067 | 1 |
| 0.075 | 1 |
| 0.086 | 1 |
| 0.245 | 1 |
| 0.065 | 1 |
| 0.191 | 1 |
| 0.049 | 1 |
| 0.069 | 1 |
| 0.178 | 1 |
| 0.043 | 1 |
| 0.118 | 1 |
| 0.068 | 1 |
| 0.136 | 1 |
| 0.157 | 1 |
| 0.146 | 1 |
| 0.067 | 1 |
| 0.082 | 1 |
| 0.055 | 1 |
| 0.076 | 1 |
| 0.279 | 1 |
| 0.062 | 1 |
| 0.111 | 1 |
| 0.087 | 1 |
| 0.132 | 1 |
| 0.067 | 1 |
| 0.081 | 1 |
| 0.09 | 1 |
| 0.053 | 1 |
| 0.085 | 1 |
| 0.139 | 1 |
| 0.041 | 1 |
| 0.04 | 1 |
| 0.202 | 1 |
| 0.149 | 1 |
| 0.679 | 1 |
| 0.09 | 1 |
| 0.114 | 1 |
| 0.044 | 1 |
| 0.051 | 1 |
| 0.208 | 1 |
| 0.073 | 1 |
| 0.11 | 1 |
| 0.093 | 1 |
| 0.055 | 1 |
| 0.064 | 1 |
| 0.064 | 1 |
| 0.07 | 1 |
| 0.196 | 1 |
| 0.174 | 1 |
| 0.096 | 1 |
| 0.145 | 1 |
| 0.128 | 1 |
| 0.056 | 1 |
| 0.064 | 1 |
| 0.082 | 1 |
| 0.067 | 1 |
| 0.101 | 1 |
| 0.088 | 1 |
| 0.05 | 1 |
| 0.053 | 1 |
| 0.11 | 1 |
| 0.068 | 1 |
| 0.055 | 1 |
| 0.056 | 1 |
| 0.095 | 1 |
| 0.075 | 1 |
| 0.112 | 1 |
| 0.067 | 1 |
| 0.052 | 1 |
| 0.08 | 1 |
| 0.096 | 1 |
| 0.051 | 1 |
| 0.142 | 1 |
| 0.129 | 1 |
| 0.069 | 1 |
| 0.054 | 1 |
| 0.146 | 1 |
| 0.527 | 1 |
| 0.05 | 1 |
| 0.049 | 1 |
| 0.23 | 1 |
| 0.237 | 1 |
| 0.076 | 1 |
| 0.053 | 1 |
| 0.175 | 1 |
| 0.161 | 1 |
| 0.062 | 1 |
| 0.097 | 1 |
| 0.05 | 1 |
| 0.043 | 1 |
| 0.047 | 1 |
| 0.175 | 1 |
| 0.091 | 1 |
| 0.126 | 1 |
| 0.071 | 1 |
| 0.07 | 1 |
| 0.078 | 1 |
| 0.077 | 1 |
| 0.052 | 1 |
| 0.05 | 1 |
| 0.044 | 1 |
| 0.134 | 1 |
| 0.041 | 1 |
| 0.062 | 1 |
| 0.04 | 1 |
| 0.074 | 1 |
| 0.053 | 1 |
| 0.04 | 1 |
| 0.229 | 1 |
| 0.05 | 1 |
| 0.074 | 1 |
| 0.043 | 1 |
| 0.111 | 1 |
| 0.045 | 1 |
| 0.116 | 1 |
| 0.107 | 1 |
| 0.177 | 1 |
| 0.146 | 1 |
| 0.042 | 1 |
| 0.055 | 1 |
| 0.042 | 1 |
| 0.051 | 1 |
| 0.048 | 1 |
| 0.131 | 1 |
| 0.082 | 1 |
| 0.106 | 1 |
| 0.065 | 1 |
| 0.088 | 1 |
| 0.065 | 1 |
| 0.134 | 1 |
| 0.129 | 1 |
| 0.063 | 1 |
| 0.071 | 1 |
| 0.118 | 1 |
| 0.047 | 1 |
| 0.07 | 1 |
| 0.078 | 1 |
| 0.108 | 1 |
| 0.083 | 1 |
| 0.082 | 1 |
| 0.107 | 1 |
| 0.152 | 1 |
| 0.136 | 1 |
| 0.074 | 1 |
| 0.066 | 1 |
| 0.095 | 1 |
| 0.061 | 1 |
| 0.117 | 1 |
| 0.055 | 1 |
| 0.048 | 1 |
| 0.056 | 1 |
| 0.065 | 1 |
| 0.134 | 1 |
| 0.156 | 1 |
| 0.229 | 1 |
| 0.051 | 1 |
| 0.074 | 1 |
| 0.113 | 1 |
| 0.064 | 1 |
| 0.043 | 1 |
| 0.057 | 1 |
| 0.081 | 1 |
| 0.063 | 1 |
| 0.051 | 1 |
| 0.058 | 1 |
| 0.139 | 1 |
| 0.18 | 1 |
| 0.1 | 1 |
| 0.076 | 1 |
| 0.06 | 1 |
| 0.087 | 1 |
| 0.05 | 1 |
| 0.087 | 1 |
| 0.192 | 1 |
| 0.075 | 1 |
| 0.242 | 1 |
| 0.076 | 1 |
| 0.114 | 1 |
| 0.121 | 1 |
| 0.082 | 1 |
| 0.149 | 1 |
| 0.127 | 1 |
| 0.087 | 1 |
| 0.057 | 1 |
| 0.145 | 1 |
| 0.059 | 1 |
| 0.141 | 1 |
| 0.083 | 1 |
| 0.05 | 1 |
| 0.085 | 1 |
| 0.051 | 1 |
| 0.105 | 1 |
| 0.091 | 1 |
| 0.042 | 1 |
| 0.057 | 1 |
| 0.054 | 1 |
| 0.064 | 1 |
| 0.071 | 1 |
| 0.287 | 1 |
| 0.065 | 1 |
| 0.158 | 1 |
| 0.046 | 1 |
| 0.269 | 1 |
| 0.059 | 1 |
| 0.054 | 1 |
| 0.083 | 1 |
| 0.106 | 1 |
| 0.126 | 1 |
| 0.141 | 1 |
| 0.052 | 1 |
| 0.145 | 1 |
| 0.114 | 1 |
| 0.04 | 1 |
| 0.233 | 1 |
| 0.063 | 1 |
| 0.042 | 1 |
| 0.054 | 1 |
| 0.133 | 1 |
| 0.092 | 1 |
| 0.059 | 1 |
| 0.074 | 1 |
| 0.086 | 1 |
| 0.083 | 1 |
| 0.062 | 1 |
| 0.051 | 1 |
| 0.094 | 1 |
| 0.1 | 1 |
| 0.109 | 1 |
| 0.293 | 1 |
| 0.097 | 1 |
| 0.056 | 1 |
| 0.097 | 1 |
| 0.129 | 1 |
| 0.049 | 1 |
| 0.113 | 1 |
| 0.152 | 1 |
| 0.074 | 1 |
| 0.263 | 1 |
| 0.063 | 1 |
| 0.088 | 1 |
| 0.069 | 1 |
| 0.056 | 1 |
| 0.071 | 1 |
| 0.186 | 1 |
| 0.052 | 1 |
| 0.102 | 1 |
| 0.065 | 1 |
| 0.056 | 1 |
| 0.057 | 1 |
| 0.398 | 1 |
| 0.072 | 1 |
| 0.077 | 1 |
| 0.088 | 1 |
| 0.069 | 1 |
| 0.065 | 1 |
| 0.064 | 1 |
| 0.091 | 1 |
| 0.088 | 1 |
| 0.049 | 1 |
| 0.077 | 1 |
| 0.047 | 1 |
| 0.12 | 1 |
| 0.096 | 1 |
| 0.147 | 1 |
| 0.054 | 1 |
| 0.065 | 1 |
| 0.043 | 1 |
| 0.077 | 1 |
| 0.099 | 1 |
| 0.051 | 1 |
| 0.209 | 1 |
| 0.124 | 1 |
| 0.093 | 1 |
| 0.108 | 1 |
| 0.094 | 1 |
| 0.115 | 1 |
| 0.101 | 1 |
| 0.063 | 1 |
| 0.041 | 1 |
| 0.055 | 1 |
| 0.154 | 1 |
| 0.097 | 1 |
| 0.095 | 1 |
| 0.043 | 1 |
| 0.056 | 1 |
| 0.07 | 1 |
| 0.057 | 1 |
| 0.206 | 1 |
| 0.101 | 1 |
| 0.13 | 1 |
| 0.071 | 1 |
| 0.076 | 1 |
| 0.042 | 1 |
| 0.187 | 1 |
| 0.055 | 1 |
| 0.146 | 1 |
| 0.105 | 1 |
| 0.048 | 1 |
| 0.31 | 1 |
| 0.1 | 1 |
| 0.167 | 1 |
| 0.05 | 1 |
| 0.056 | 1 |
| 0.043 | 1 |
| 0.102 | 1 |
| 0.134 | 1 |
| 0.077 | 1 |
| 0.055 | 1 |
| 0.072 | 1 |
| 0.089 | 1 |
| 0.127 | 1 |
| 0.056 | 1 |
| 0.053 | 1 |
| 0.062 | 1 |
| 0.085 | 1 |
| 0.109 | 1 |
| 0.244 | 1 |
| 0.115 | 1 |
| 0.294 | 1 |
| 0.091 | 1 |
| 0.135 | 1 |
| 0.143 | 1 |
| 0.043 | 1 |
| 0.135 | 1 |
| 0.061 | 1 |
| 0.041 | 1 |
| 0.184 | 1 |
| 0.061 | 1 |
| 0.1 | 1 |
| 0.069 | 1 |
| 0.068 | 1 |
| 0.045 | 1 |
| 0.099 | 1 |
| 0.051 | 1 |
| 0.493 | 1 |
| 0.106 | 1 |
| 0.091 | 1 |
| 0.098 | 1 |
| 0.089 | 1 |
| 0.116 | 1 |
| 0.072 | 1 |
| 0.042 | 1 |
| 0.091 | 1 |
| 0.095 | 1 |
| 0.042 | 1 |
| 0.055 | 1 |
| 0.044 | 1 |
| 0.082 | 1 |
| 0.113 | 1 |
| 0.056 | 1 |
| 0.069 | 1 |
| 0.206 | 1 |
| 0.109 | 1 |
| 0.059 | 1 |
| 0.054 | 1 |
| 0.137 | 1 |
| 0.078 | 1 |
| 0.051 | 1 |
| 0.076 | 1 |
| 0.057 | 1 |
| 0.059 | 1 |
| 0.054 | 1 |
| 0.257 | 1 |
| 0.086 | 1 |
| 0.085 | 1 |
| 0.101 | 1 |
| 0.089 | 1 |
| 0.069 | 1 |
| 0.144 | 1 |
| 0.411 | 1 |
| 0.149 | 1 |
| 0.13 | 1 |
| 0.096 | 1 |
| 0.058 | 1 |
| 0.114 | 1 |
| 0.085 | 1 |
| 0.07 | 1 |
| 0.06 | 1 |
| 0.199 | 1 |
| 0.082 | 1 |
| 0.063 | 1 |
| 0.067 | 1 |
| 0.078 | 1 |
| 0.127 | 1 |
| 0.077 | 1 |
| 0.06 | 1 |
| 0.063 | 1 |
| 0.062 | 1 |
| 0.207 | 1 |
| 0.095 | 1 |
| 0.049 | 1 |
| 0.158 | 1 |
| 0.047 | 1 |
| 0.046 | 1 |
| 0.154 | 1 |
| 0.057 | 1 |
| 0.061 | 1 |
| 0.068 | 1 |
| 0.075 | 1 |
| 0.133 | 1 |
| 0.059 | 1 |
| 0.065 | 1 |
| 0.135 | 1 |
| 0.187 | 1 |
| 0.088 | 1 |
| 0.049 | 1 |
| 0.052 | 1 |
| 0.191 | 1 |
| 0.049 | 1 |
| 0.218 | 1 |
| 0.072 | 1 |
| 0.058 | 1 |
| 0.059 | 1 |
| 0.16 | 1 |
| 0.046 | 1 |
| 0.104 | 1 |
| 0.045 | 1 |
| 0.44 | 1 |
| 0.183 | 1 |
| 0.17 | 1 |
| 0.093 | 1 |
| 0.05 | 1 |
| 0.16 | 1 |
| 0.057 | 1 |
| 0.056 | 1 |
| 0.052 | 1 |
| 0.042 | 1 |
| 0.082 | 1 |
| 0.049 | 1 |
| 0.049 | 1 |
| 0.099 | 1 |
| 0.098 | 1 |
| 0.062 | 1 |
| 0.044 | 1 |
| 0.056 | 1 |
| 0.568 | 1 |
| 0.059 | 1 |
| 0.043 | 1 |
| 0.075 | 1 |
| 0.149 | 1 |
| 0.106 | 1 |
| 0.123 | 1 |
| 0.061 | 1 |
| 0.201 | 1 |
| 0.725 | 1 |
| 0.148 | 1 |
| 0.047 | 1 |
| 0.365 | 1 |
| 0.092 | 1 |
| 0.049 | 1 |
| 0.121 | 1 |
| 0.191 | 1 |
| 0.131 | 1 |
| 0.071 | 1 |
| 0.048 | 1 |
| 0.114 | 1 |
| 0.161 | 1 |
| 0.057 | 1 |
| 0.059 | 1 |
| 0.089 | 1 |
| 0.224 | 1 |
| 0.115 | 1 |
| 0.058 | 1 |
| 0.082 | 1 |
| 0.186 | 1 |
| 0.313 | 1 |
| 0.055 | 1 |
| 0.043 | 1 |
| 0.069 | 1 |
| 0.363 | 1 |
| 0.135 | 1 |
| 0.195 | 1 |
| 0.128 | 1 |
| 0.064 | 1 |
| 0.086 | 1 |
| 0.074 | 1 |
| 0.043 | 1 |
| 0.066 | 1 |
| 0.269 | 1 |
| 0.066 | 1 |
| 0.081 | 1 |
| 0.05 | 1 |
| 0.205 | 1 |
| 0.042 | 1 |
| 0.072 | 1 |
| 0.218 | 1 |
| 0.1 | 1 |
| 0.067 | 1 |
| 0.055 | 1 |
| 0.232 | 1 |
| 0.079 | 1 |
| 0.147 | 1 |
| 0.084 | 1 |
| 0.043 | 1 |
| 0.164 | 1 |
| 0.083 | 1 |
| 0.188 | 1 |
| 0.123 | 1 |
| 0.043 | 1 |
| 0.081 | 1 |
| 0.055 | 1 |
| 0.075 | 1 |
| 0.123 | 1 |
| 0.076 | 1 |
| 0.054 | 1 |
| 0.083 | 1 |
| 0.229 | 1 |
| 0.09 | 1 |
| 0.067 | 1 |
| 0.109 | 1 |
| 0.209 | 1 |
| 0.077 | 1 |
| 0.091 | 1 |
| 0.275 | 1 |
| 0.096 | 1 |
| 0.105 | 1 |
| 0.412 | 1 |
| 0.095 | 1 |
| 0.068 | 1 |
| 0.07 | 1 |
| 0.095 | 1 |
| 0.056 | 1 |
| 0.066 | 1 |
| 0.125 | 1 |
| 0.093 | 1 |
| 0.129 | 1 |
| 0.072 | 1 |
| 0.063 | 1 |
| 0.389 | 1 |
| 0.074 | 1 |
| 0.14 | 1 |
| 0.052 | 1 |
| 0.066 | 1 |
| 0.09 | 1 |
| 0.193 | 1 |
| 0.071 | 1 |
| 0.144 | 1 |
| 0.382 | 1 |
| 0.08 | 1 |
| 0.182 | 1 |
| 0.118 | 1 |
| 0.351 | 1 |
| 0.167 | 1 |
| 0.171 | 1 |
| 0.092 | 1 |
| 0.082 | 1 |
| 0.105 | 1 |
| 0.291 | 1 |
| 0.052 | 1 |
| 0.067 | 1 |
| 0.064 | 1 |
| 0.058 | 1 |
| 0.048 | 1 |
| 0.161 | 1 |
| 0.043 | 1 |
| 0.047 | 1 |
| 0.088 | 1 |
| 0.287 | 1 |
| 0.097 | 1 |
| 0.1 | 1 |
| 0.065 | 1 |
| 0.088 | 1 |
| 0.198 | 1 |
| 0.129 | 1 |
| 0.073 | 1 |
| 0.065 | 1 |
| 0.059 | 1 |
| 0.23 | 1 |
| 0.083 | 1 |
| 0.104 | 1 |
| 0.09 | 1 |
| 0.088 | 1 |
| 0.153 | 1 |
| 0.1 | 1 |
| 0.078 | 1 |
| 0.153 | 1 |
| 0.131 | 1 |
| 0.085 | 1 |
| 0.129 | 1 |
| 0.163 | 1 |
| 0.066 | 1 |
| 0.067 | 1 |
| 0.06 | 1 |
| 0.055 | 1 |
| 0.068 | 1 |
| 0.189 | 1 |
| 0.109 | 1 |
| 0.261 | 1 |
| 0.104 | 1 |
| 0.084 | 1 |
| 0.182 | 1 |
| 0.065 | 1 |
| 0.052 | 1 |
| 0.05 | 1 |
| 0.09 | 1 |
| 0.055 | 1 |
| 0.07 | 1 |
| 0.375 | 1 |
| 0.128 | 1 |
| 0.094 | 1 |
| 0.121 | 1 |
| 0.125 | 1 |
| 0.048 | 1 |
| 0.155 | 1 |
| 0.334 | 1 |
| 0.06 | 1 |
| 0.15 | 1 |
| 0.073 | 1 |
| 0.055 | 1 |
| 0.247 | 1 |
| 0.107 | 1 |
| 0.075 | 1 |
| 0.045 | 1 |
| 0.083 | 1 |
| 0.246 | 1 |
| 0.056 | 1 |
| 0.386 | 1 |
| 0.108 | 1 |
| 0.3 | 1 |
| 0.101 | 1 |
| 0.201 | 1 |
| 0.144 | 1 |
| 0.149 | 1 |
| 0.071 | 1 |
| 0.082 | 1 |
| 0.12 | 1 |
| 0.132 | 1 |
| 0.138 | 1 |
| 0.068 | 1 |
| 0.07 | 1 |
| 0.072 | 1 |
| 0.095 | 1 |
| 0.086 | 1 |
| 0.05 | 1 |
| 0.105 | 1 |
| 0.056 | 1 |
| 0.066 | 1 |
| 0.077 | 1 |
| 0.117 | 1 |
| 0.131 | 1 |
| 0.071 | 1 |
| 0.062 | 1 |
| 0.073 | 1 |
| 0.133 | 1 |
| 0.136 | 1 |
| 0.068 | 1 |
| 0.086 | 1 |
| 0.051 | 1 |
| 0.044 | 1 |
| 0.085 | 1 |
| 0.217 | 1 |
| 0.238 | 1 |
| 0.254 | 1 |
| 0.073 | 1 |
| 0.063 | 1 |
| 0.056 | 1 |
| 0.181 | 1 |
| 0.092 | 1 |
| 0.077 | 1 |
| 0.141 | 1 |
| 0.101 | 1 |
| 0.06 | 1 |
| 0.1 | 1 |
| 0.182 | 1 |
| 0.185 | 1 |
| 0.135 | 1 |
| 0.052 | 1 |
| 0.14 | 1 |
| 0.076 | 1 |
| 0.053 | 1 |
| 0.073 | 1 |
| 0.071 | 1 |
| 0.054 | 1 |
| 0.08 | 1 |
| 0.047 | 1 |
| 0.102 | 1 |
| 0.095 | 1 |
| 0.075 | 1 |
| 0.149 | 1 |
| 0.13 | 1 |
| 0.053 | 1 |
| 0.063 | 1 |
| 0.069 | 1 |
| 0.111 | 1 |
| 0.055 | 1 |
| 0.279 | 1 |
| 0.076 | 1 |
| 0.117 | 1 |
| 0.089 | 1 |
| 0.054 | 1 |
| 0.101 | 1 |
| 0.074 | 1 |
| 0.05 | 1 |
| 0.073 | 1 |
| 0.068 | 1 |
| 0.055 | 1 |
| 0.123 | 1 |
| 0.098 | 1 |
| 0.101 | 1 |
| 0.046 | 1 |
| 0.21 | 1 |
| 0.071 | 1 |
| 0.062 | 1 |
| 0.159 | 1 |
| 0.123 | 1 |
| 0.051 | 1 |
| 0.065 | 1 |
| 0.123 | 1 |
| 0.06 | 1 |
| 0.063 | 1 |
| 0.057 | 1 |
| 0.057 | 1 |
| 0.188 | 1 |
| 0.085 | 1 |
| 0.131 | 1 |
| 0.059 | 1 |
| 0.077 | 1 |
| 0.058 | 1 |
| 0.125 | 1 |
| 0.048 | 1 |
| 0.109 | 1 |
| 0.168 | 1 |
| 0.055 | 1 |
| 0.064 | 1 |
| 0.055 | 1 |
| 0.066 | 1 |
| 0.302 | 1 |
| 0.156 | 1 |
| 0.051 | 1 |
| 0.055 | 1 |
| 0.372 | 1 |
| 0.064 | 1 |
| 0.051 | 1 |
| 0.045 | 1 |
| 0.08 | 1 |
| 0.093 | 1 |
| 0.061 | 1 |
| 0.079 | 1 |
| 0.082 | 1 |
| 0.071 | 1 |
| 0.206 | 1 |
| 0.245 | 1 |
| 0.059 | 1 |
| 0.116 | 1 |
| 0.059 | 1 |
| 0.135 | 1 |
| 0.187 | 1 |
| 0.07 | 1 |
| 0.052 | 1 |
| 0.163 | 1 |
| 0.052 | 1 |
| 0.066 | 1 |
| 0.075 | 1 |
| 0.121 | 1 |
| 0.055 | 1 |
| 0.095 | 1 |
| 0.056 | 1 |
| 0.055 | 1 |
| 0.068 | 1 |
| 0.054 | 1 |
| 0.066 | 1 |
| 0.103 | 1 |
| 0.053 | 1 |
| 0.064 | 1 |
| 0.066 | 1 |
| 0.096 | 1 |
| 0.068 | 1 |
| 0.066 | 1 |
| 0.056 | 1 |
| 0.206 | 1 |
| 0.086 | 1 |
| 0.142 | 1 |
| 0.074 | 1 |
| 0.078 | 1 |
| 0.051 | 1 |
| 0.622 | 1 |
| 0.054 | 1 |
| 0.068 | 1 |
| 0.14 | 1 |
| 0.061 | 1 |
| 0.139 | 1 |
| 0.128 | 1 |
| 0.051 | 1 |
| 0.126 | 1 |
| 0.046 | 1 |
| 0.09 | 1 |
| 0.103 | 1 |
| 0.08 | 1 |
| 0.133 | 1 |
| 0.265 | 1 |
| 0.188 | 1 |
| 0.056 | 1 |
| 0.054 | 1 |
| 0.129 | 1 |
| 0.111 | 1 |
| 0.057 | 1 |
| 0.061 | 1 |
| 0.087 | 1 |
| 0.143 | 1 |
| 0.062 | 1 |
| 0.055 | 1 |
| 0.073 | 1 |
| 0.101 | 1 |
| 0.118 | 1 |
| 0.057 | 1 |
| 0.094 | 1 |
| 0.053 | 1 |
| 0.079 | 1 |
| 0.065 | 1 |
| 0.178 | 1 |
| 0.057 | 1 |
| 0.442 | 1 |
| 0.313 | 1 |
| 0.107 | 1 |
| 0.139 | 1 |
| 0.15 | 1 |
| 0.059 | 1 |
| 0.181 | 1 |
| 0.083 | 1 |
| 0.111 | 1 |
| 0.156 | 1 |
| 0.077 | 1 |
| 0.058 | 1 |
| 0.065 | 1 |
| 0.051 | 1 |
| 0.052 | 1 |
| 0.076 | 1 |
| 0.087 | 1 |
| 0.058 | 1 |
| 0.06 | 1 |
| 0.141 | 1 |
| 0.052 | 1 |
| 0.064 | 1 |
| 0.147 | 1 |
| 0.111 | 1 |
| 0.065 | 1 |
| 0.046 | 1 |
| 0.06 | 1 |
| 0.144 | 1 |
| 0.102 | 1 |
| 0.072 | 1 |
| 0.058 | 1 |
| 0.062 | 1 |
| 0.082 | 1 |
| 0.135 | 1 |
| 0.108 | 1 |
| 0.294 | 1 |
| 0.067 | 1 |
| 0.057 | 1 |
| 0.053 | 1 |
| 0.047 | 1 |
| 0.054 | 1 |
| 0.263 | 1 |
| 0.053 | 1 |
| 0.075 | 1 |
| 0.055 | 1 |
| 0.176 | 1 |
| 0.175 | 1 |
| 0.146 | 1 |
| 0.053 | 1 |
| 0.138 | 1 |
| 0.078 | 1 |
| 0.214 | 1 |
| 0.093 | 1 |
| 0.069 | 1 |
| 0.066 | 1 |
| 0.05 | 1 |
| 0.059 | 1 |
| 0.05 | 1 |
| 0.055 | 1 |
| 0.053 | 1 |
| 0.048 | 1 |
| 0.058 | 1 |
| 0.048 | 1 |
| 0.064 | 1 |
| 0.057 | 1 |
| 0.089 | 1 |
| 0.081 | 1 |
| 0.102 | 1 |
| 0.186 | 1 |
| 0.154 | 1 |
| 0.066 | 1 |
| 0.144 | 1 |
| 0.068 | 1 |
| 0.074 | 1 |
| 0.064 | 1 |
| 0.07 | 1 |
| 0.219 | 1 |
| 0.106 | 1 |
| 0.092 | 1 |
| 0.079 | 1 |
| 0.051 | 1 |
| 0.097 | 1 |
| 0.151 | 1 |
| 0.156 | 1 |
| 0.067 | 1 |
| 0.448 | 1 |
| 0.113 | 1 |
| 0.213 | 1 |
| 0.086 | 1 |
| 0.18 | 1 |
| 0.053 | 1 |
| 0.094 | 1 |
| 0.09 | 1 |
| 0.064 | 1 |
| 0.071 | 1 |
| 0.081 | 1 |
| 0.052 | 1 |
| 0.052 | 1 |
| 0.117 | 1 |
| 0.201 | 1 |
| 0.103 | 1 |
| 0.11 | 1 |
| 0.079 | 1 |
| 0.154 | 1 |
| 0.081 | 1 |
| 0.063 | 1 |
| 0.187 | 1 |
| 0.124 | 1 |
| 0.066 | 1 |
| 0.089 | 1 |
| 0.06 | 1 |
| 0.156 | 1 |
| 0.171 | 1 |
| 0.071 | 1 |
| 0.078 | 1 |
| 0.059 | 1 |
| 0.073 | 1 |
| 0.056 | 1 |
| 0.207 | 1 |
| 0.092 | 1 |
| 0.088 | 1 |
| 0.372 | 1 |
| 0.078 | 1 |
| 0.106 | 1 |
| 0.076 | 1 |
| 0.131 | 1 |
| 0.051 | 1 |
| 0.079 | 1 |
| 0.066 | 1 |
| 0.068 | 1 |
| 0.144 | 1 |
| 0.086 | 1 |
| 0.136 | 1 |
| 0.107 | 1 |
| 0.124 | 1 |
| 0.078 | 1 |
| 0.186 | 1 |
| 0.063 | 1 |
| 0.141 | 1 |
| 0.087 | 1 |
| 0.062 | 1 |
| 0.058 | 1 |
| 0.106 | 1 |
| 0.198 | 1 |
| 0.079 | 1 |
| 0.049 | 1 |
| 0.064 | 1 |
| 0.108 | 1 |
| 0.075 | 1 |
| 0.22 | 1 |
| 0.114 | 1 |
| 0.087 | 1 |
| 0.223 | 1 |
| 0.081 | 1 |
| 0.15 | 1 |
| 0.063 | 1 |
| 0.085 | 1 |
| 0.379 | 1 |
| 0.117 | 1 |
| 0.166 | 1 |
| 0.121 | 1 |
| 0.075 | 1 |
| 0.08 | 1 |
| 0.076 | 1 |
| 0.06 | 1 |
| 0.059 | 1 |
| 0.059 | 1 |
| 0.057 | 1 |
| 0.053 | 1 |
| 0.048 | 1 |
| 0.08 | 1 |
| 0.089 | 1 |
| 0.238 | 1 |
| 0.083 | 1 |
| 0.078 | 1 |
| 0.061 | 1 |
| 0.051 | 1 |
| 0.995 | 1 |
| 0.112 | 1 |
| 0.15 | 1 |
| 0.113 | 1 |
| 0.05 | 1 |
| 0.098 | 1 |
| 0.072 | 1 |
| 0.071 | 1 |
| 0.095 | 1 |
| 0.064 | 1 |
| 0.092 | 1 |
| 0.048 | 1 |
| 0.065 | 1 |
| 0.054 | 1 |
| 0.098 | 1 |
| 0.062 | 1 |
| 0.07 | 1 |
| 0.084 | 1 |
| 0.07 | 1 |
| 0.135 | 1 |
| 0.049 | 1 |
| 0.062 | 1 |
| 0.05 | 1 |
| 0.052 | 1 |
| 0.07 | 1 |
| 0.101 | 1 |
| 0.482 | 1 |
| 0.08 | 1 |
| 0.051 | 1 |
| 0.084 | 1 |
| 0.069 | 1 |
| 0.219 | 1 |
| 0.065 | 1 |
| 0.089 | 1 |
| 0.066 | 1 |
| 0.073 | 1 |
| 0.054 | 1 |
| 0.138 | 1 |
| 0.05 | 1 |
| 0.093 | 1 |
| 0.053 | 1 |
| 0.113 | 1 |
| 0.056 | 1 |
| 0.068 | 1 |
| 0.061 | 1 |
| 0.09 | 1 |
| 0.063 | 1 |
| 0.061 | 1 |
| 0.131 | 1 |
| 0.071 | 1 |
| 0.141 | 1 |
| 0.131 | 1 |
| 0.086 | 1 |
| 0.058 | 1 |
| 0.108 | 1 |
| 0.126 | 1 |
| 0.27 | 1 |
| 0.134 | 1 |
| 0.049 | 1 |
| 0.052 | 1 |
| 0.121 | 1 |
| 0.093 | 1 |
| 0.077 | 1 |
| 0.102 | 1 |
| 0.158 | 1 |
| 0.058 | 1 |
| 0.061 | 1 |
| 0.057 | 1 |
| 0.09 | 1 |
| 0.061 | 1 |
| 0.145 | 1 |
| 0.059 | 1 |
| 0.142 | 1 |
| 0.196 | 1 |
| 0.161 | 1 |
| 0.055 | 1 |
| 0.121 | 1 |
| 0.084 | 1 |
| 0.088 | 1 |
| 0.071 | 1 |
| 0.071 | 1 |
| 0.112 | 1 |
| 0.1 | 1 |
| 0.055 | 1 |
| 0.264 | 1 |
| 0.076 | 1 |
| 0.102 | 1 |
| 0.089 | 1 |
| 0.133 | 1 |
| 0.201 | 1 |
| 0.108 | 1 |
| 0.051 | 1 |
| 0.24 | 1 |
| 0.08 | 1 |
| 0.069 | 1 |
| 0.063 | 1 |
| 0.1 | 1 |
| 0.298 | 1 |
| 0.113 | 1 |
| 0.084 | 1 |
| 0.061 | 1 |
| 0.051 | 1 |
| 0.203 | 1 |
| 0.067 | 1 |
| 0.111 | 1 |
| 0.693 | 1 |
| 0.067 | 1 |
| 0.067 | 1 |
| 0.068 | 1 |
| 0.052 | 1 |
| 0.048 | 1 |
| 0.07 | 1 |
| 0.066 | 1 |
| 0.065 | 1 |
| 0.074 | 1 |
| 0.053 | 1 |
| 0.075 | 1 |
| 0.084 | 1 |
| 0.072 | 1 |
| 0.051 | 1 |
| 0.246 | 1 |
| 0.196 | 1 |
| 0.098 | 1 |
| 0.148 | 1 |
| 0.118 | 1 |
| 0.109 | 1 |
| 0.06 | 1 |
| 0.054 | 1 |
| 0.072 | 1 |
| 0.228 | 1 |
| 0.069 | 1 |
| 0.119 | 1 |
| 0.056 | 1 |
| 0.048 | 1 |
| 0.056 | 1 |
| 0.07 | 1 |
| 0.126 | 1 |
| 0.055 | 1 |
| 0.125 | 1 |
| 0.074 | 1 |
| 0.211 | 1 |
| 0.086 | 1 |
| 0.06 | 1 |
| 0.066 | 1 |
| 0.367 | 1 |
| 0.055 | 1 |
| 0.091 | 1 |
| 0.158 | 1 |
| 0.077 | 1 |
| 0.162 | 1 |
| 0.052 | 1 |
| 0.055 | 1 |
| 0.069 | 1 |
| 0.079 | 1 |
| 0.24 | 1 |
| 0.111 | 1 |
| 0.05 | 1 |
| 0.093 | 1 |
| 0.244 | 1 |
| 0.283 | 1 |
| 0.263 | 1 |
| 0.156 | 1 |
| 0.07 | 1 |
| 0.053 | 1 |
| 0.085 | 1 |
| 0.081 | 1 |
| 0.229 | 1 |
| 0.059 | 1 |
| 0.171 | 1 |
| 0.095 | 1 |
| 0.08 | 1 |
| 0.054 | 1 |
| 0.178 | 1 |
| 0.048 | 1 |
| 0.098 | 1 |
| 0.052 | 1 |
| 0.108 | 1 |
| 0.056 | 1 |
| 0.331 | 1 |
| 0.05 | 1 |
| 0.054 | 1 |
| 0.101 | 1 |
| 0.154 | 1 |
| 0.068 | 1 |
| 0.06 | 1 |
| 0.064 | 1 |
| 0.055 | 1 |
| 0.09 | 1 |
| 0.095 | 1 |
| 0.214 | 1 |
| 0.272 | 1 |
| 0.078 | 1 |
| 0.086 | 1 |
| 0.057 | 1 |
| 0.073 | 1 |
| 0.093 | 1 |
| 0.061 | 1 |
| 0.163 | 1 |
| 0.066 | 1 |
| 0.1 | 1 |
| 0.054 | 1 |
| 0.14 | 1 |
| 0.069 | 1 |
| 0.087 | 1 |
| 0.116 | 1 |
| 0.164 | 1 |
| 0.075 | 1 |
| 0.812 | 1 |
| 0.087 | 1 |
| 0.061 | 1 |
| 0.124 | 1 |
| 0.055 | 1 |
| 0.057 | 1 |
| 0.095 | 1 |
| 0.08 | 1 |
| 0.186 | 1 |
| 0.065 | 1 |
| 0.158 | 1 |
| 0.211 | 1 |
| 0.088 | 1 |
| 0.107 | 1 |
| 0.094 | 1 |
| 0.266 | 1 |
| 0.131 | 1 |
| 0.068 | 1 |
| 0.09 | 1 |
| 0.075 | 1 |
| 0.11 | 1 |
| 0.056 | 1 |
| 0.055 | 1 |
| 0.084 | 1 |
| 0.088 | 1 |
| 0.083 | 1 |
| 0.056 | 1 |
| 0.092 | 1 |
| 0.059 | 1 |
| 0.291 | 1 |
| 0.099 | 1 |
| 0.53 | 1 |
| 0.069 | 1 |
| 0.142 | 1 |
| 0.591 | 1 |
| 0.098 | 1 |
| 0.067 | 1 |
| 0.187 | 1 |
| 0.093 | 1 |
| 0.065 | 1 |
| 0.062 | 1 |
| 0.186 | 1 |
| 0.086 | 1 |
| 0.067 | 1 |
| 0.222 | 1 |
| 0.09 | 1 |
| 0.067 | 1 |
| 0.231 | 1 |
| 0.293 | 1 |
| 0.05 | 1 |
| 0.178 | 1 |
| 0.164 | 1 |
| 0.102 | 1 |
| 0.116 | 1 |
| 0.053 | 1 |
| 0.201 | 1 |
| 0.397 | 1 |
| 0.055 | 1 |
| 0.075 | 1 |
| 0.075 | 1 |
| 0.073 | 1 |
| 0.229 | 1 |
| 0.22 | 1 |
| 0.103 | 1 |
| 0.182 | 1 |
| 0.057 | 1 |
| 0.102 | 1 |
| 0.167 | 1 |
| 0.061 | 1 |
| 0.298 | 1 |
| 0.051 | 1 |
| 0.057 | 1 |
| 0.08 | 1 |
| 0.154 | 1 |
| 0.159 | 1 |
| 0.205 | 1 |
| 0.144 | 1 |
| 0.075 | 1 |
| 0.372 | 1 |
| 0.119 | 1 |
| 0.059 | 1 |
| 0.085 | 1 |
| 0.063 | 1 |
| 0.192 | 1 |
| 0.183 | 1 |
| 0.116 | 1 |
| 0.053 | 1 |
| 0.104 | 1 |
| 0.058 | 1 |
| 0.052 | 1 |
| 0.158 | 1 |
| 0.142 | 1 |
| 0.06 | 1 |
| 0.159 | 1 |
| 0.564 | 1 |
| 0.28 | 1 |
| 0.076 | 1 |
| 0.094 | 1 |
| 0.093 | 1 |
| 0.058 | 1 |
| 0.069 | 1 |
| 0.221 | 1 |
| 0.189 | 1 |
| 0.172 | 1 |
| 0.084 | 1 |
| 0.164 | 1 |
| 0.067 | 1 |
| 0.105 | 1 |
| 0.143 | 1 |
| 0.069 | 1 |
| 0.058 | 1 |
| 0.143 | 1 |
| 0.074 | 1 |
| 0.279 | 1 |
| 0.074 | 1 |
| 0.071 | 1 |
| 0.321 | 1 |
| 0.083 | 1 |
| 0.052 | 1 |
| 0.107 | 1 |
| 0.101 | 1 |
| 0.062 | 1 |
| 0.057 | 1 |
| 0.055 | 1 |
| 0.229 | 1 |
| 0.088 | 1 |
| 0.06 | 1 |
| 0.243 | 1 |
| 0.056 | 1 |

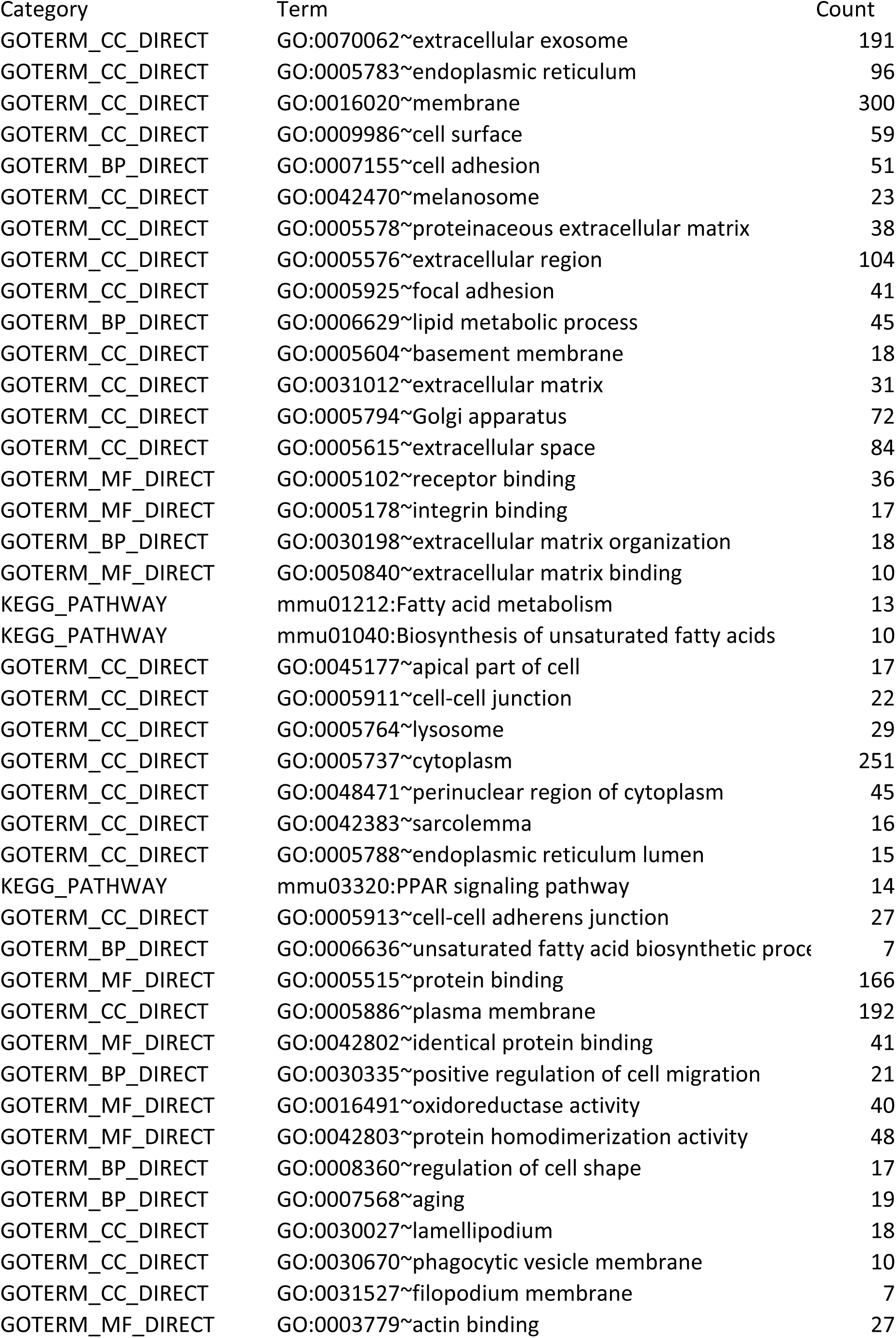
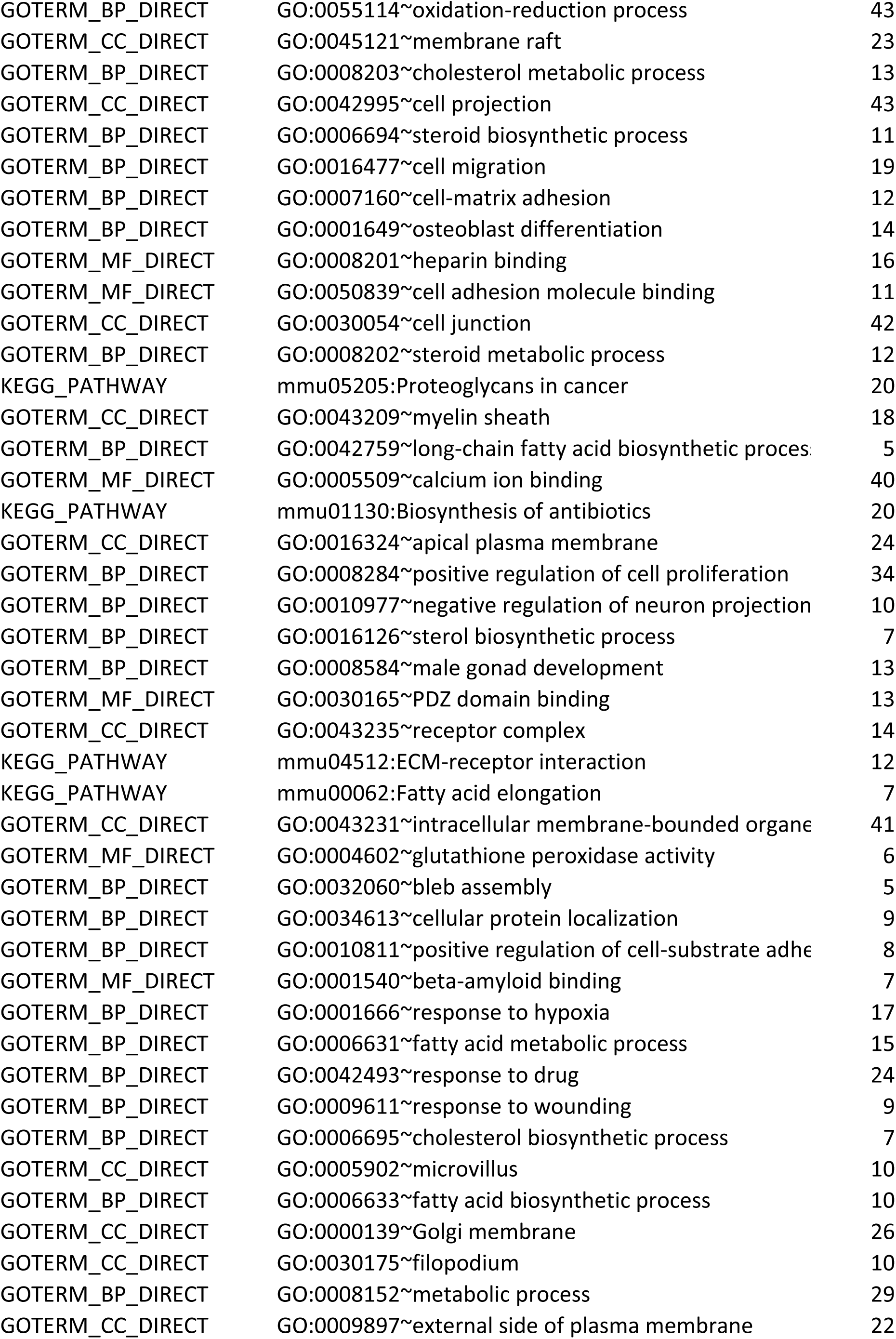
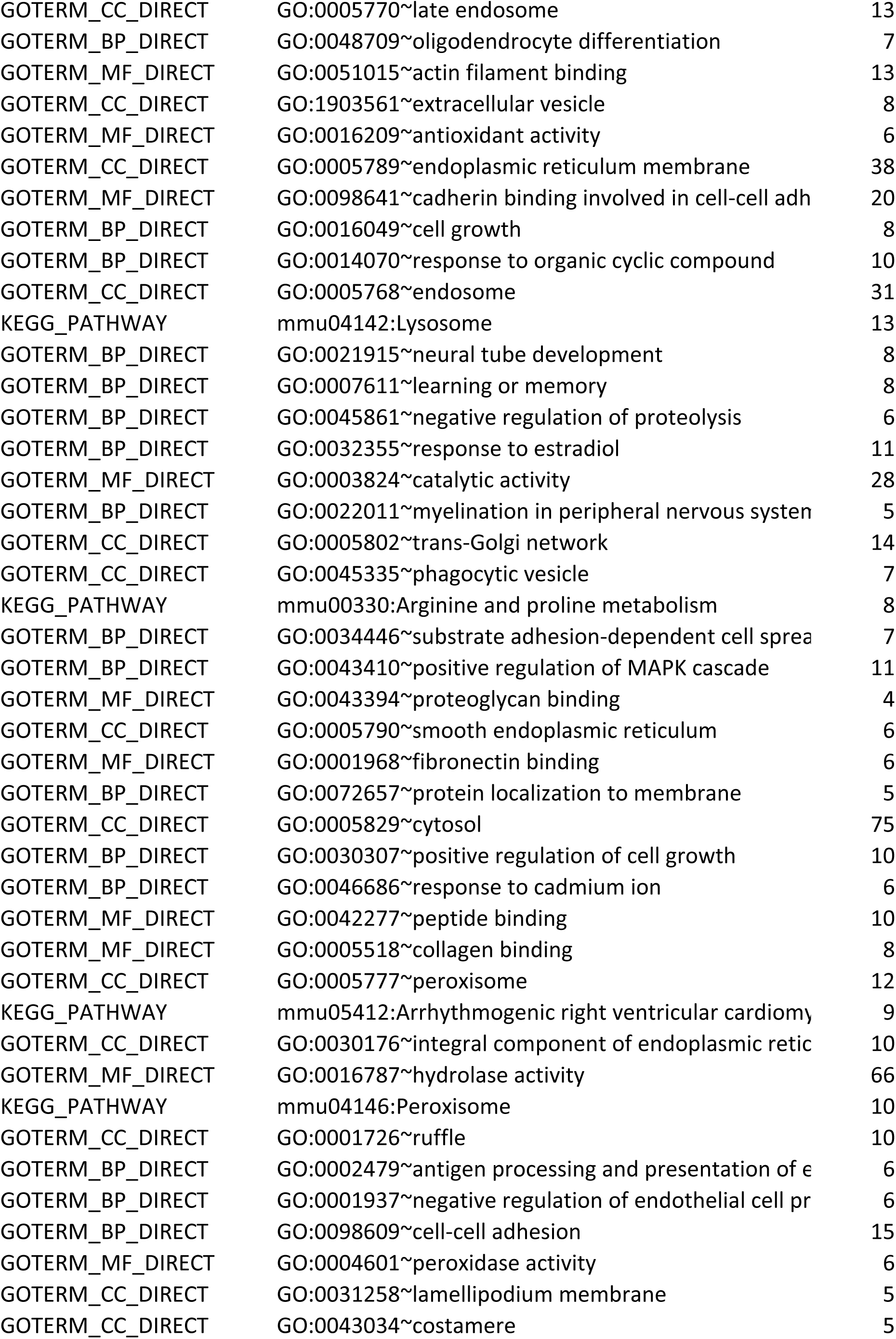
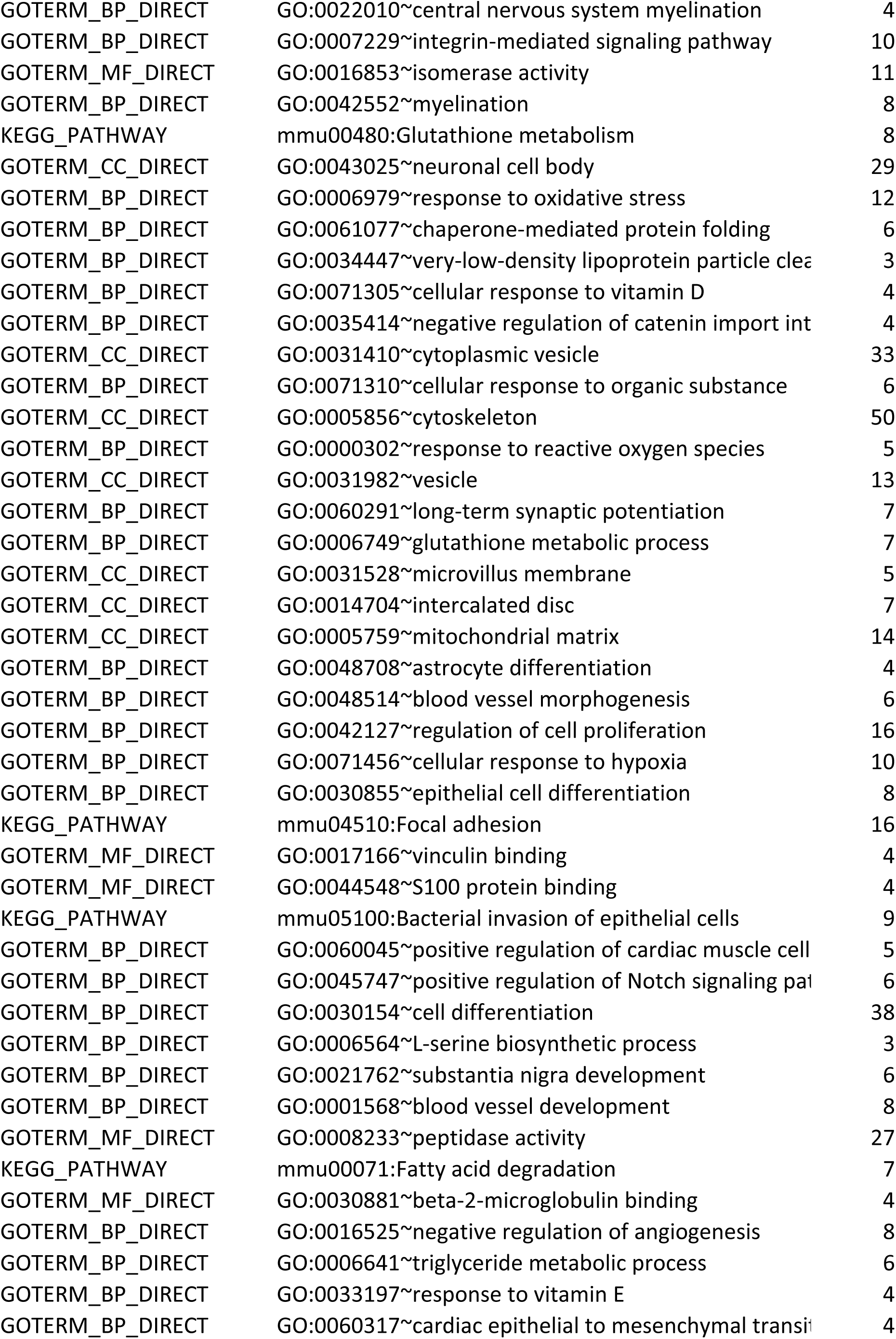
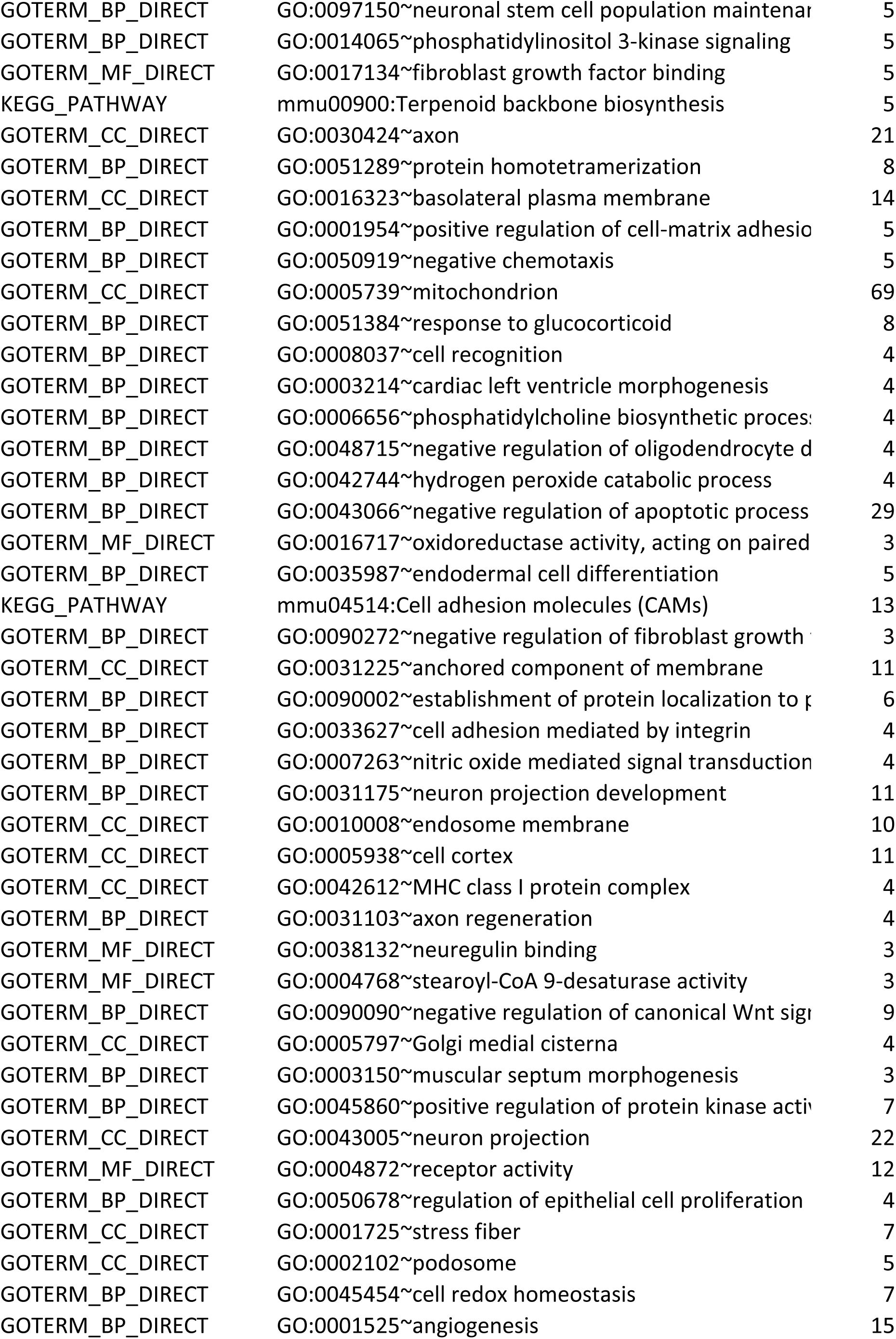
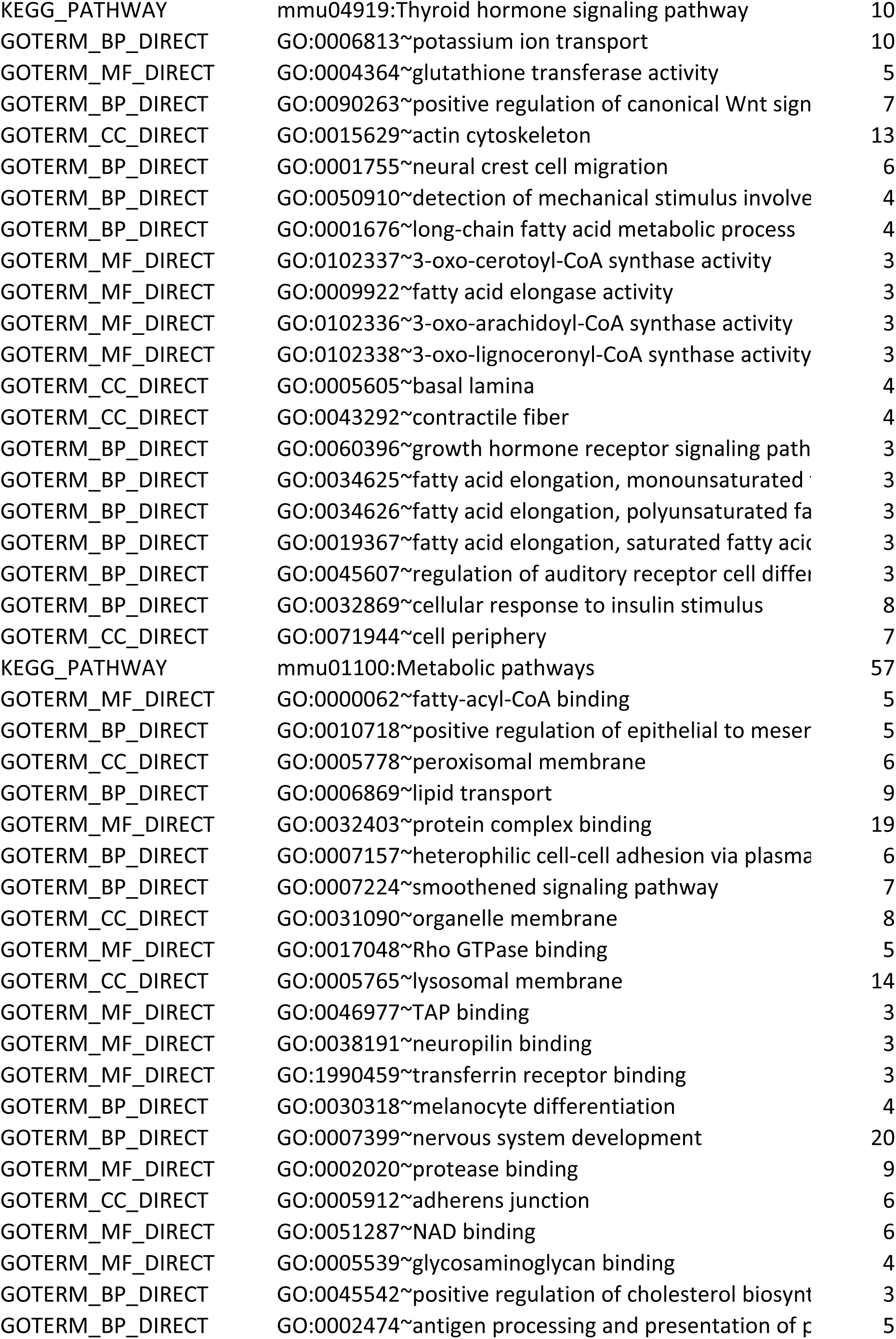
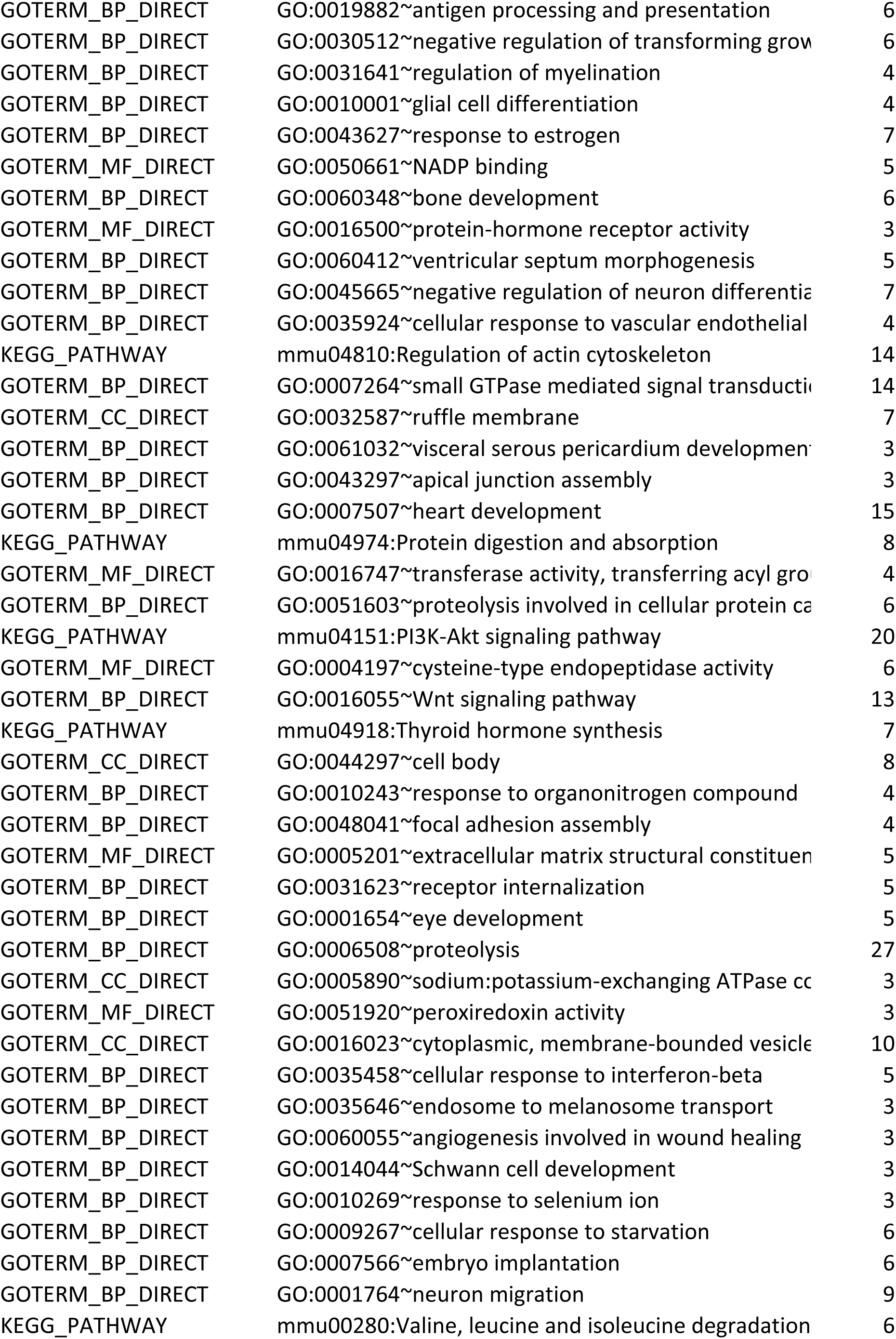
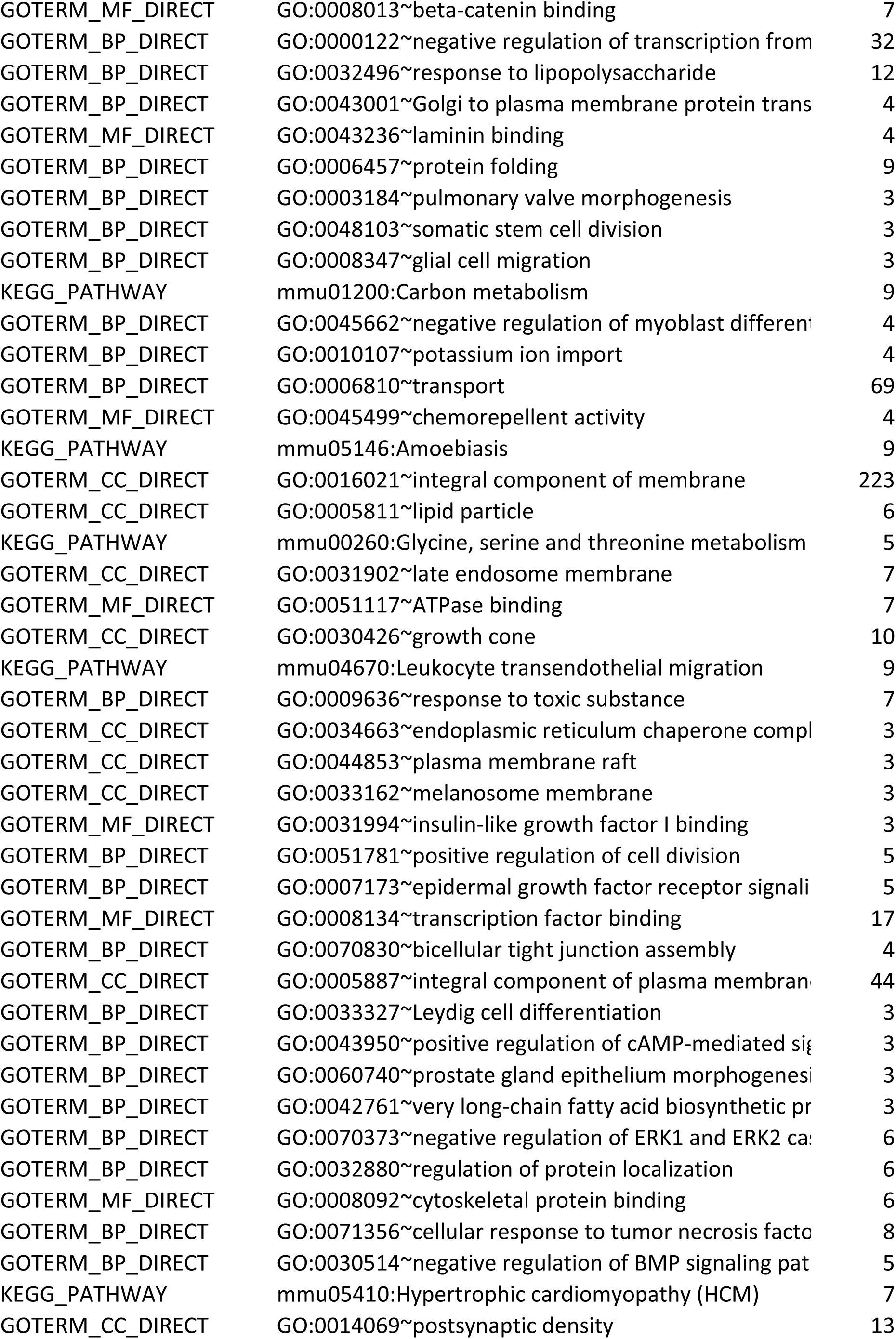
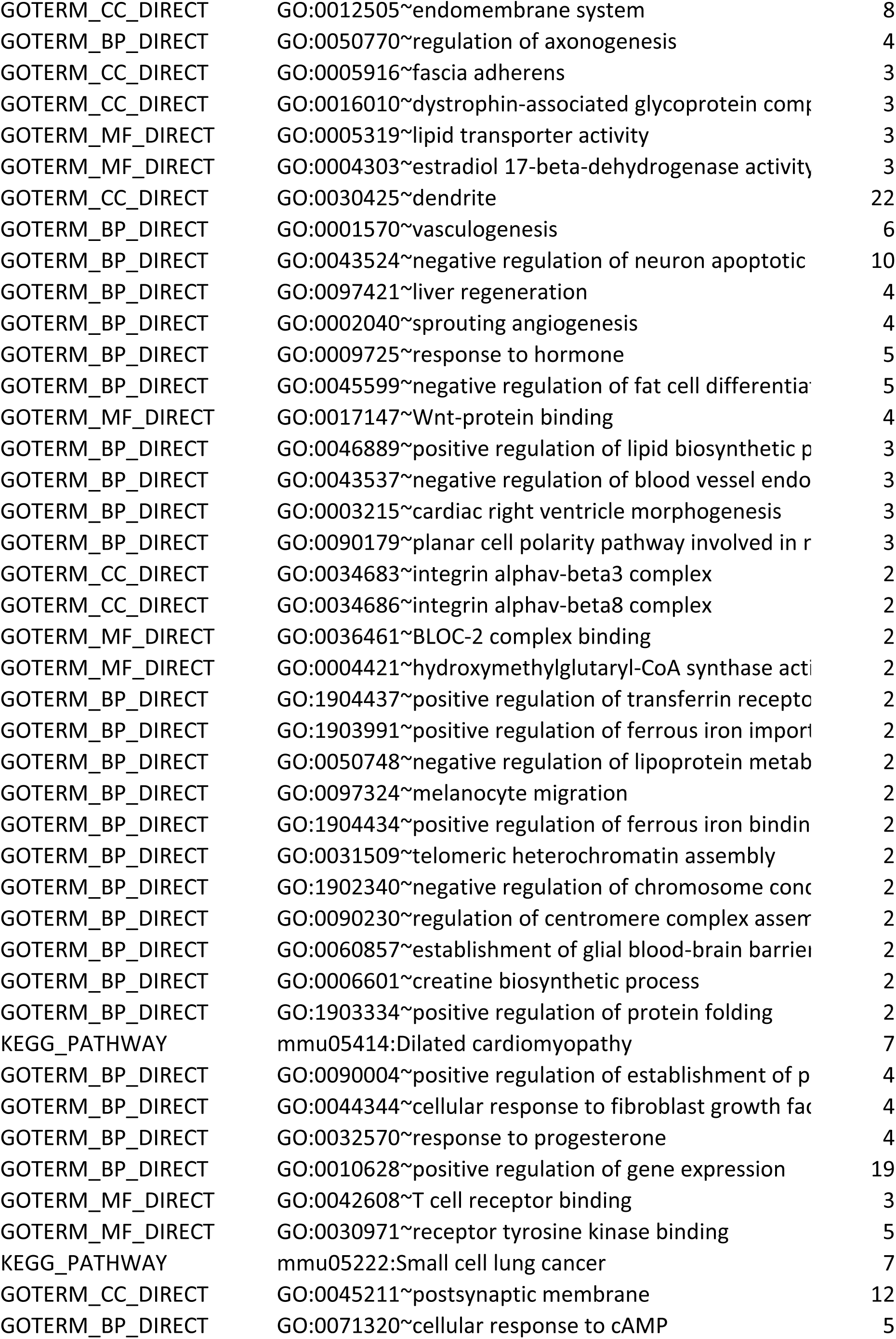
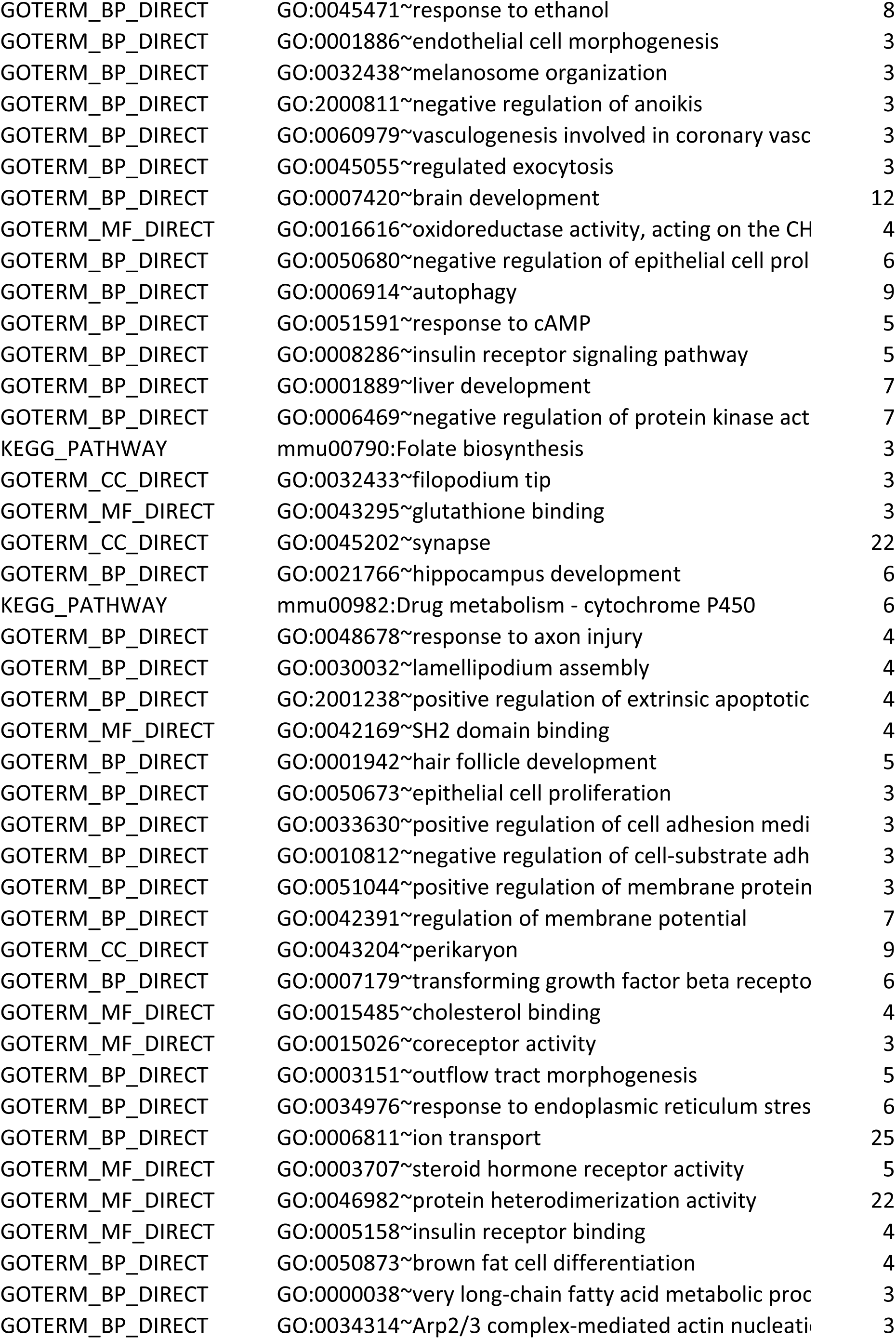
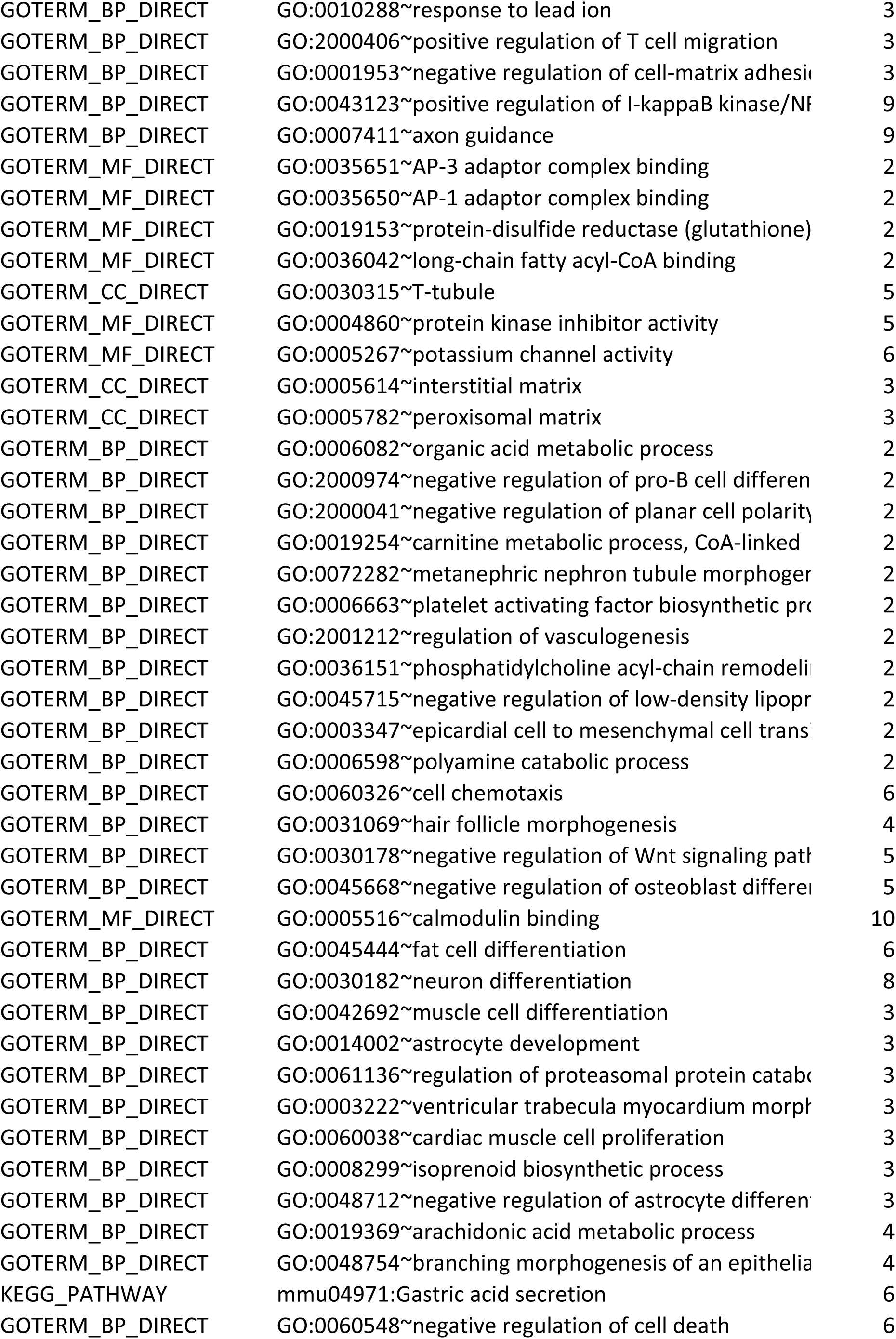
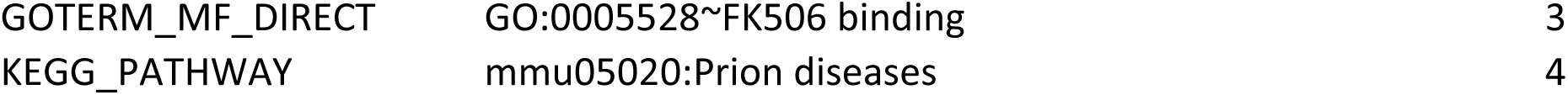

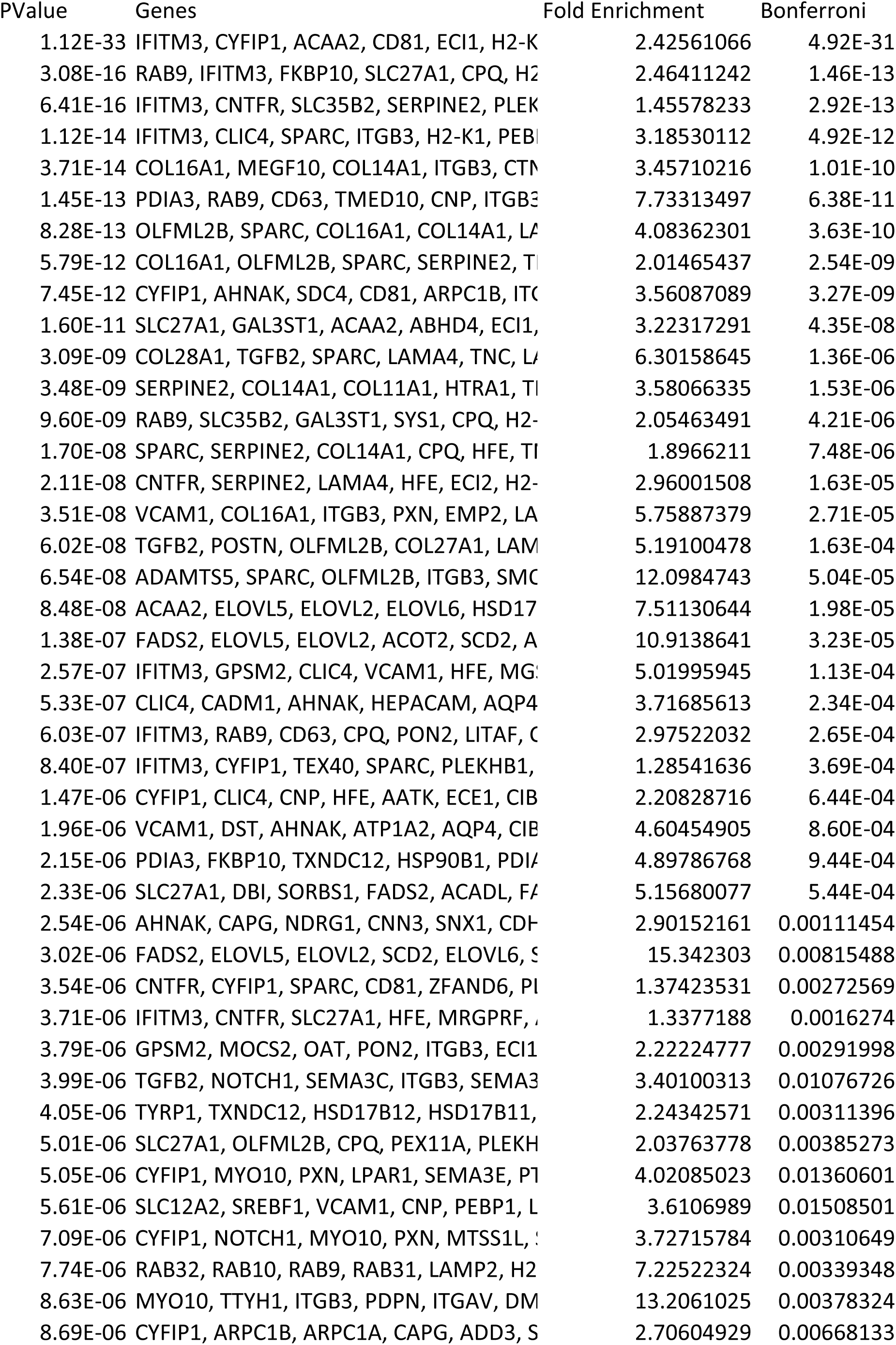
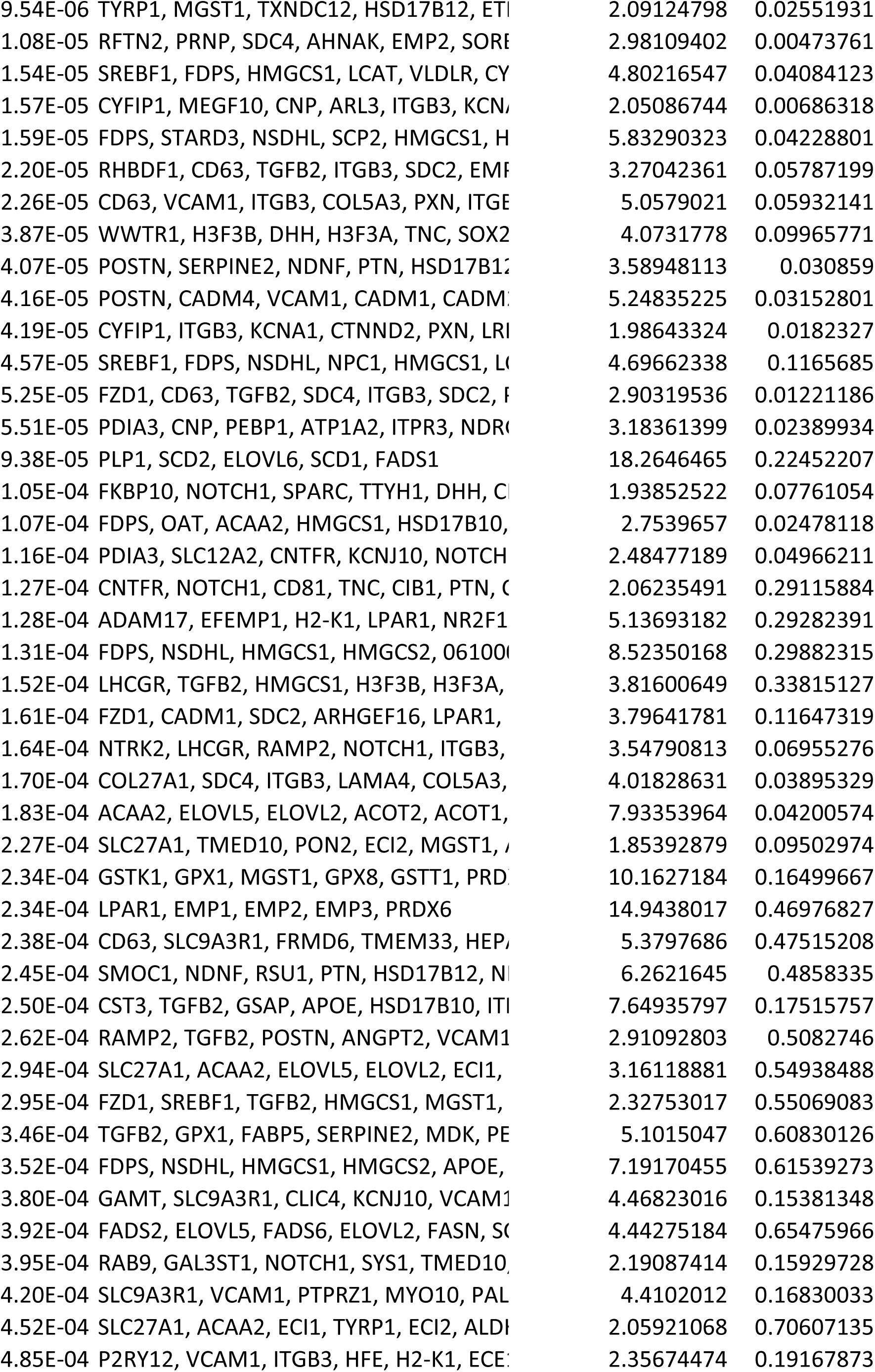
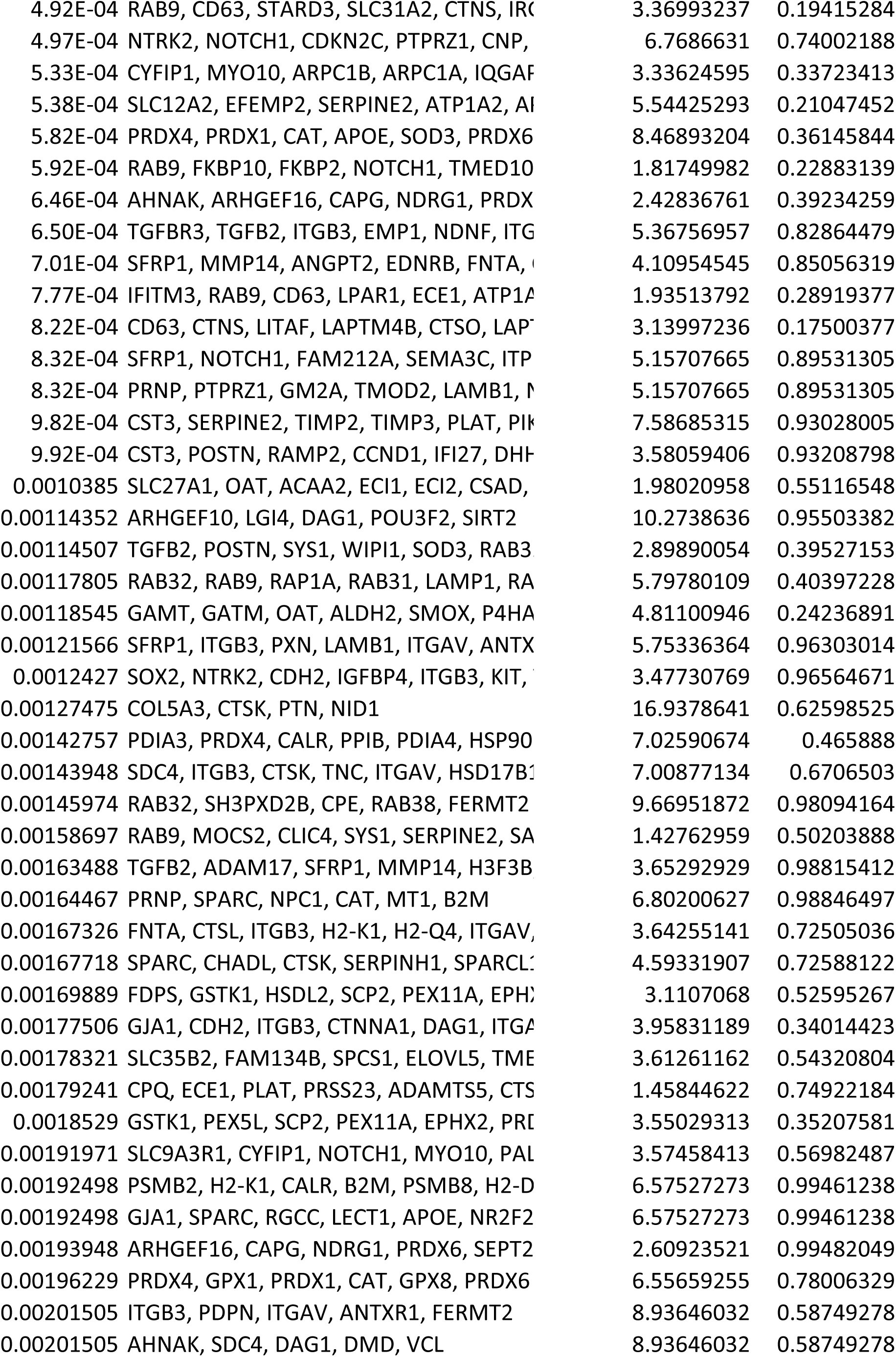
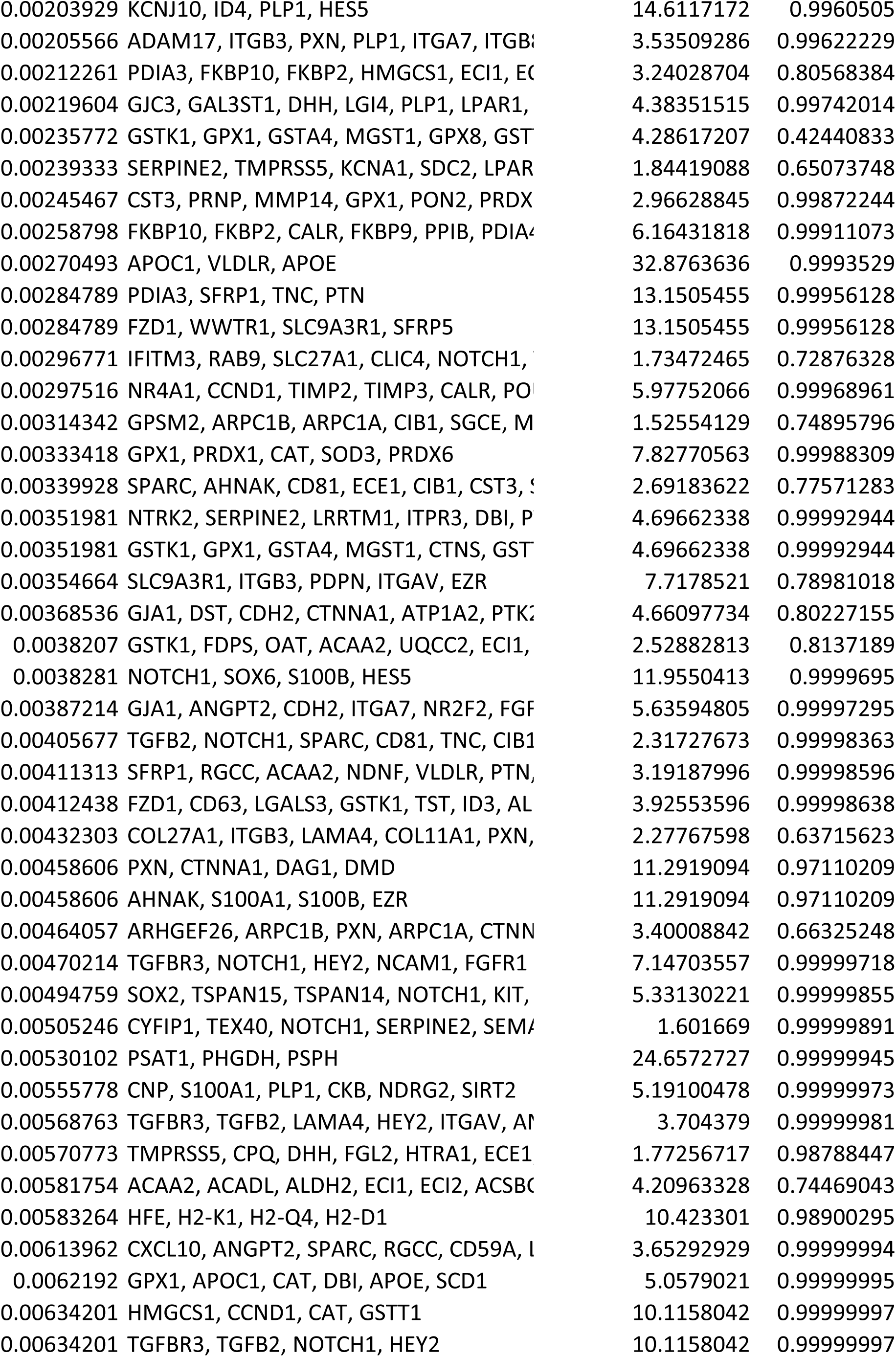
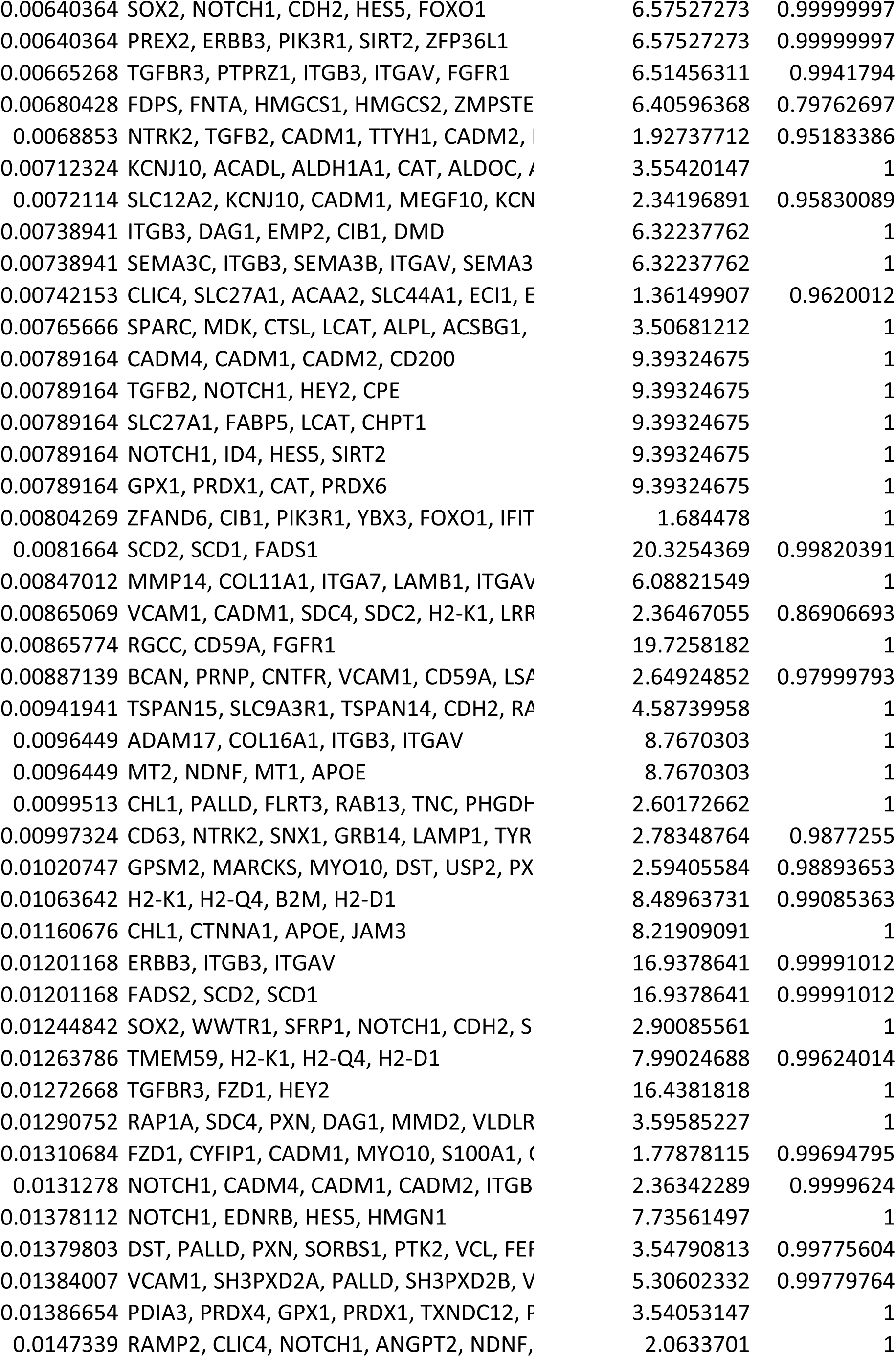
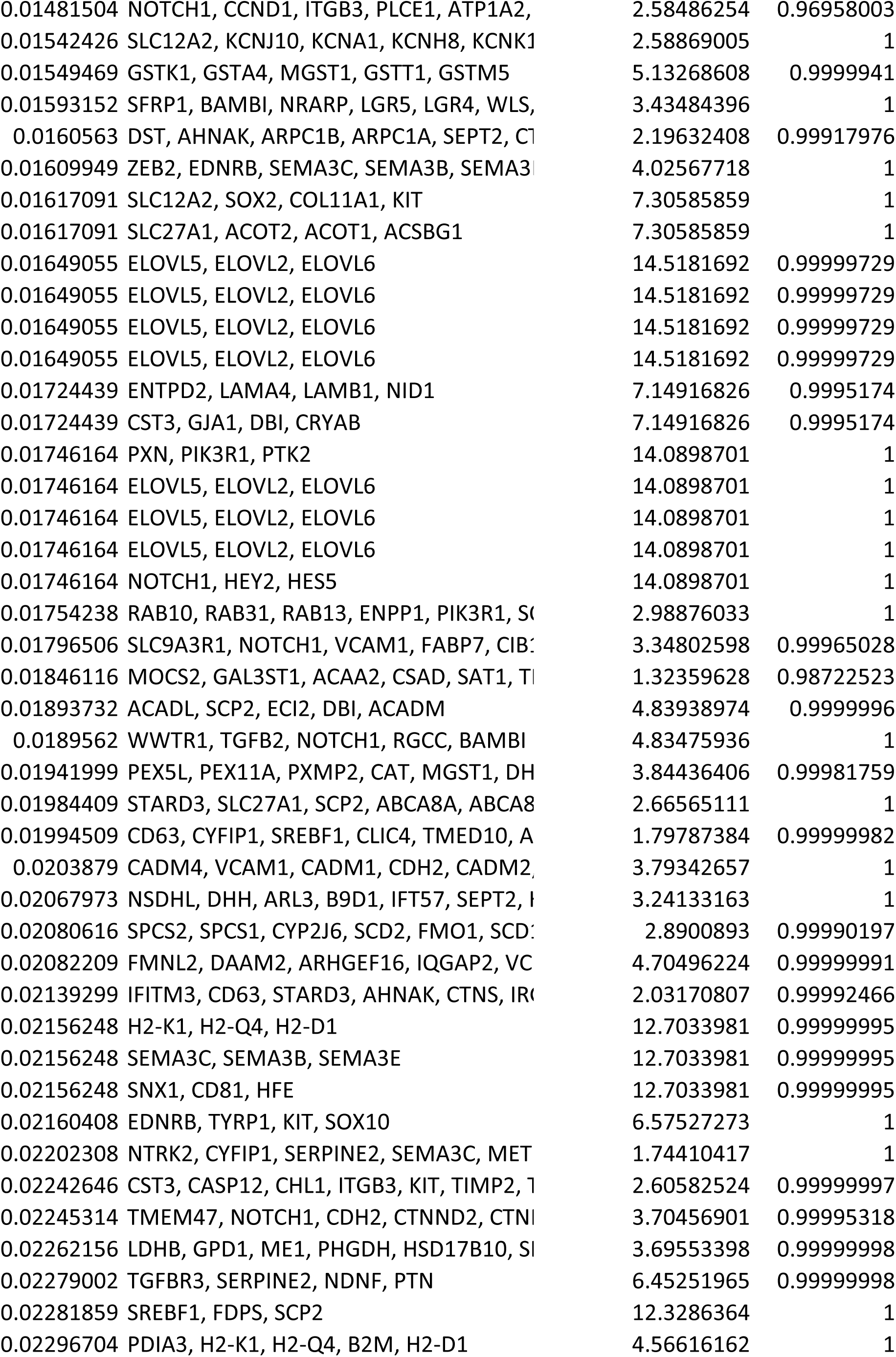
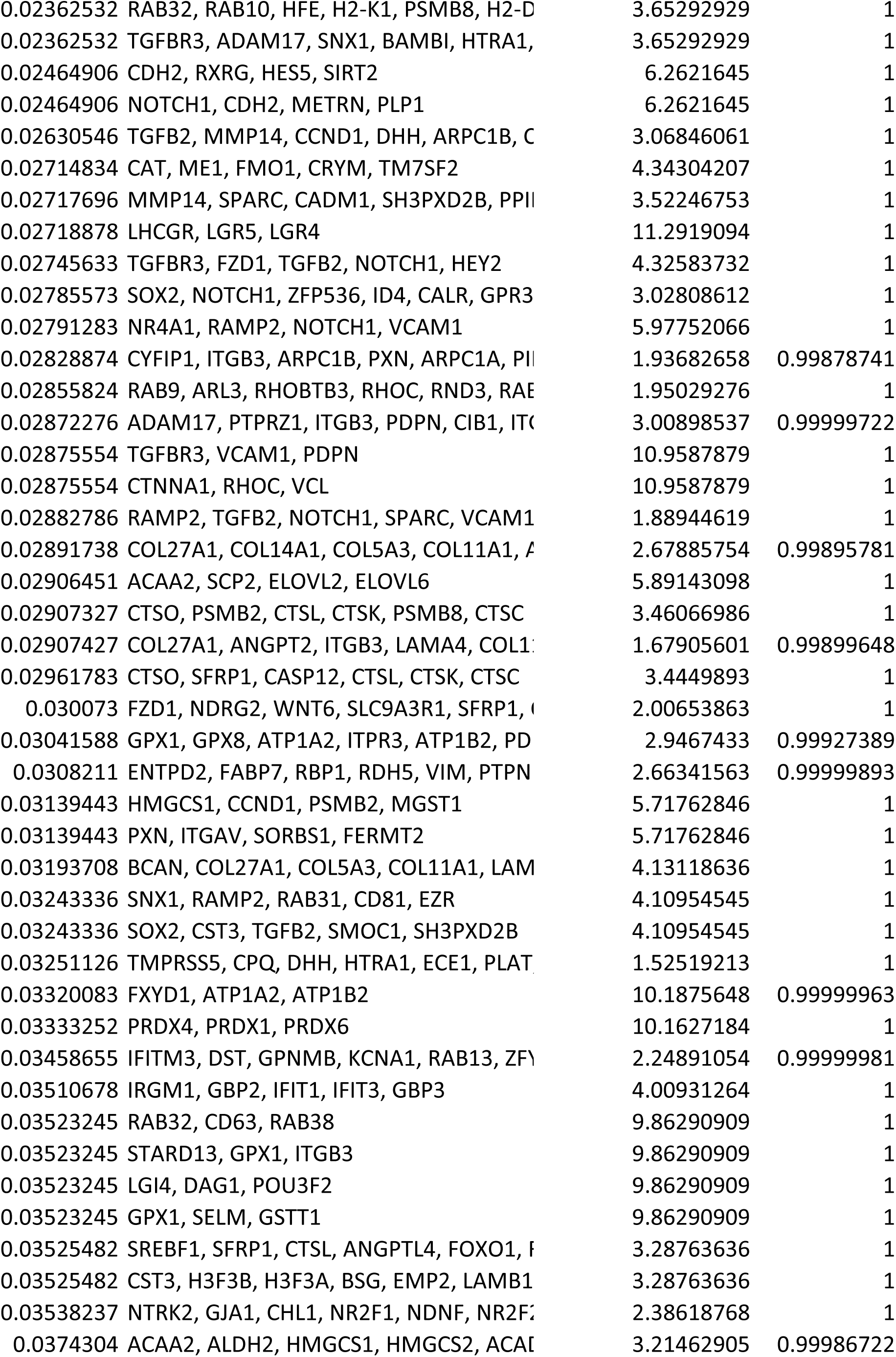
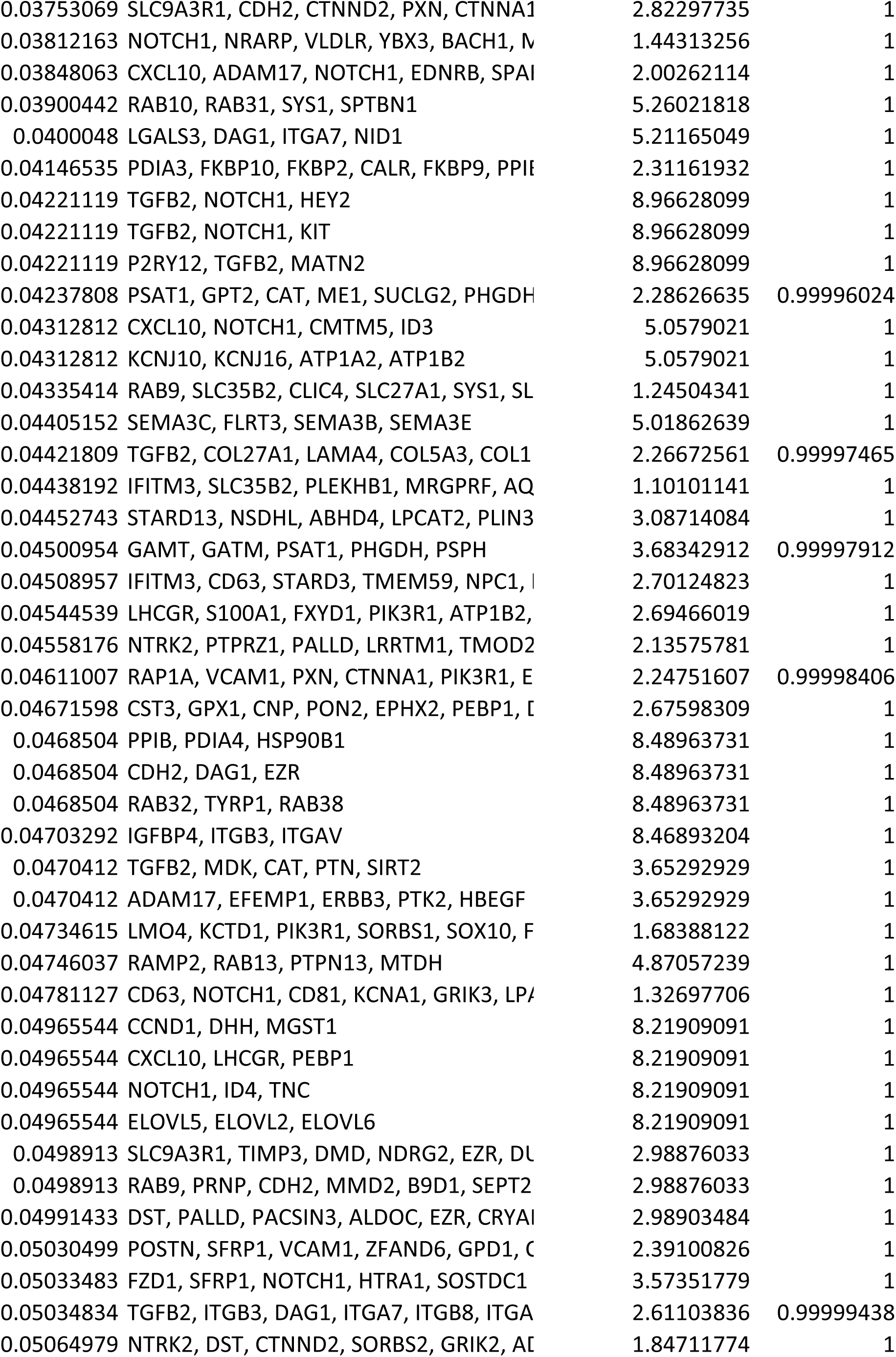
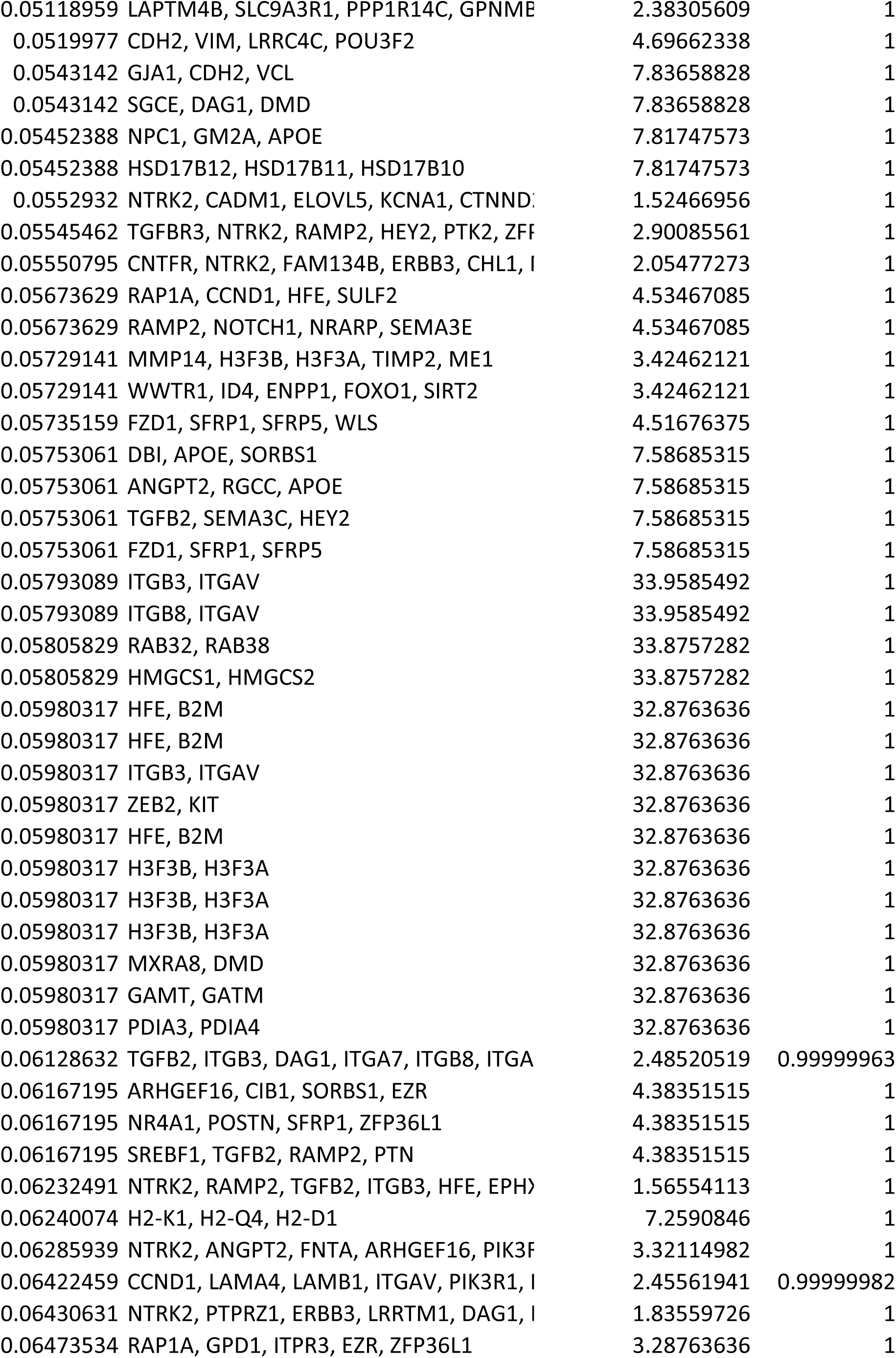
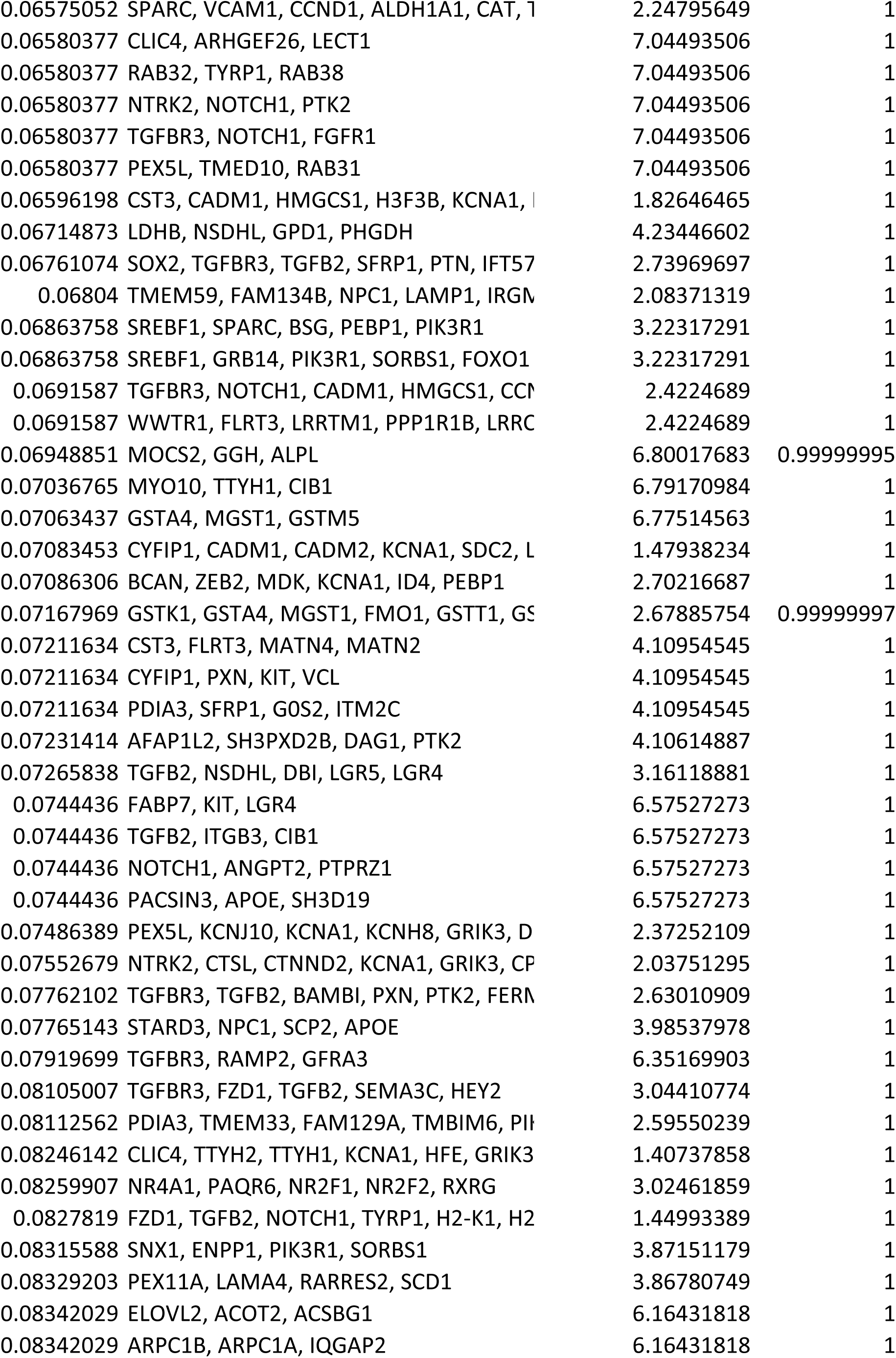
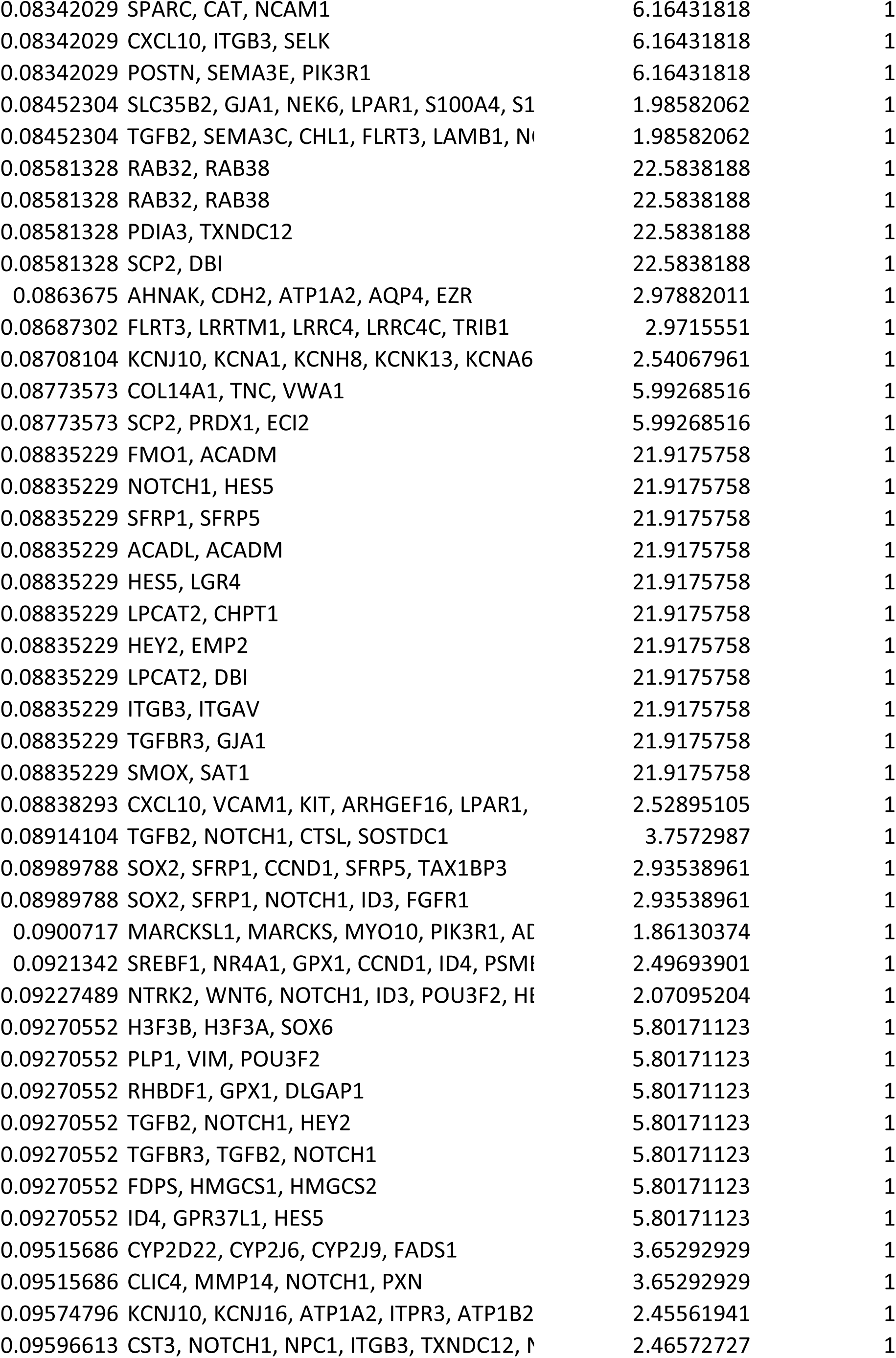
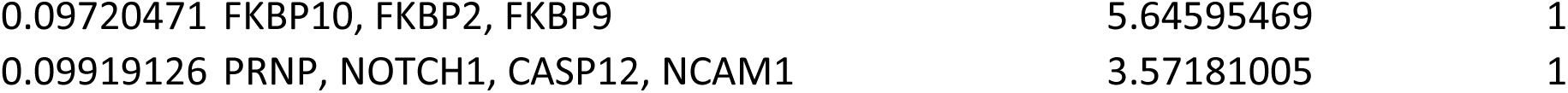

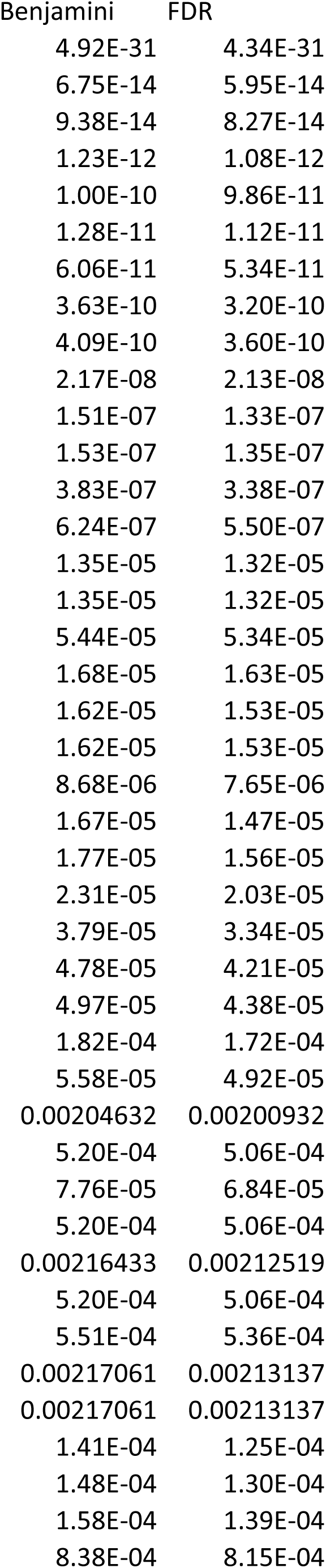
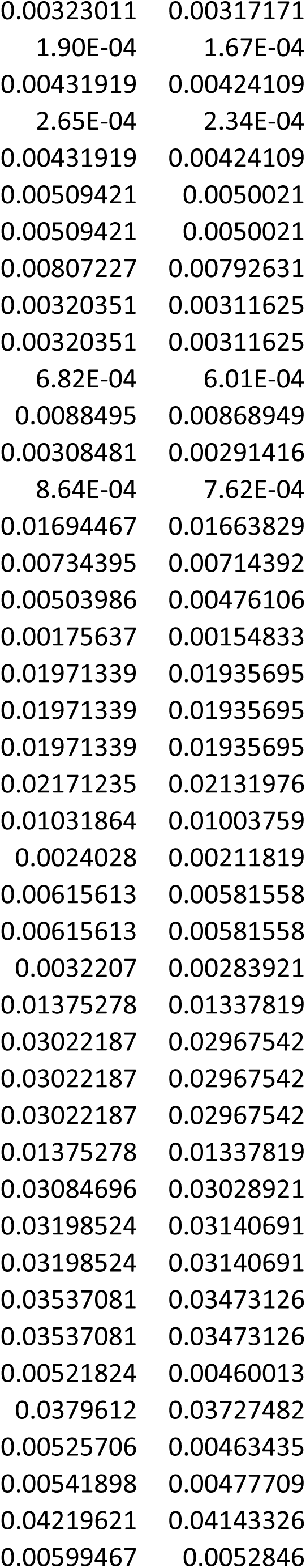
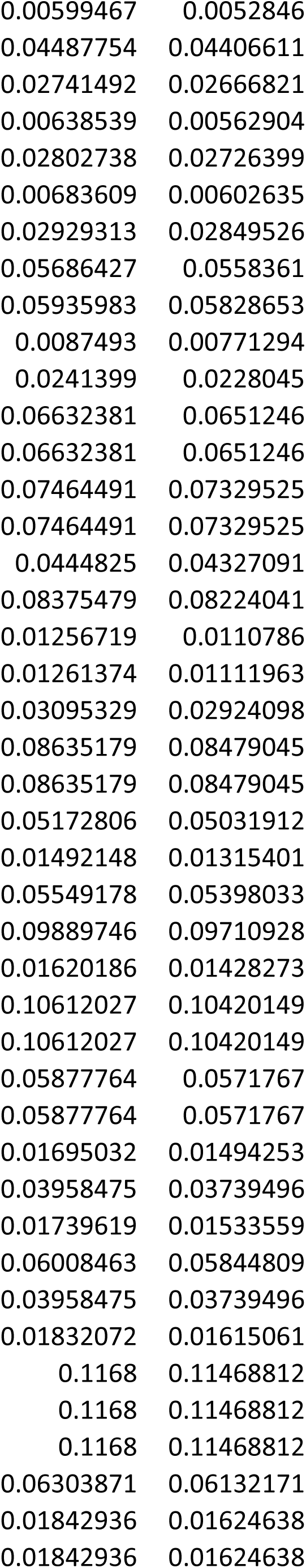
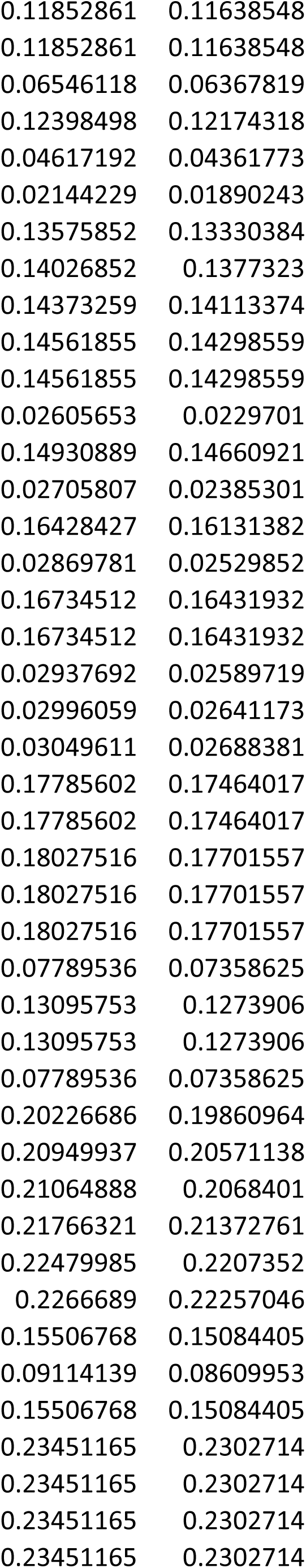
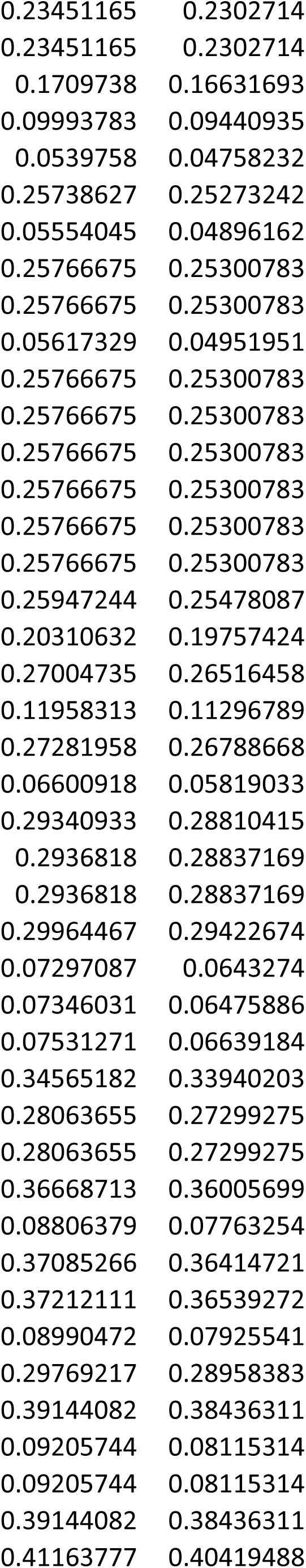
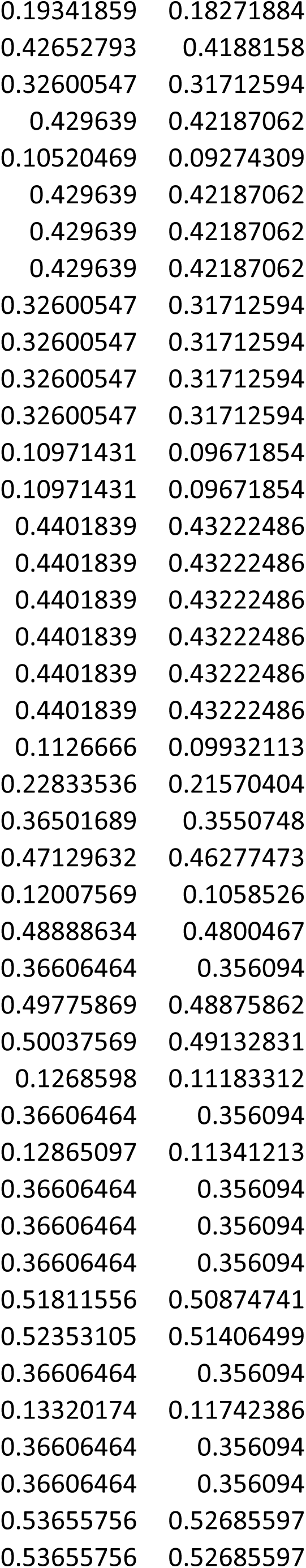
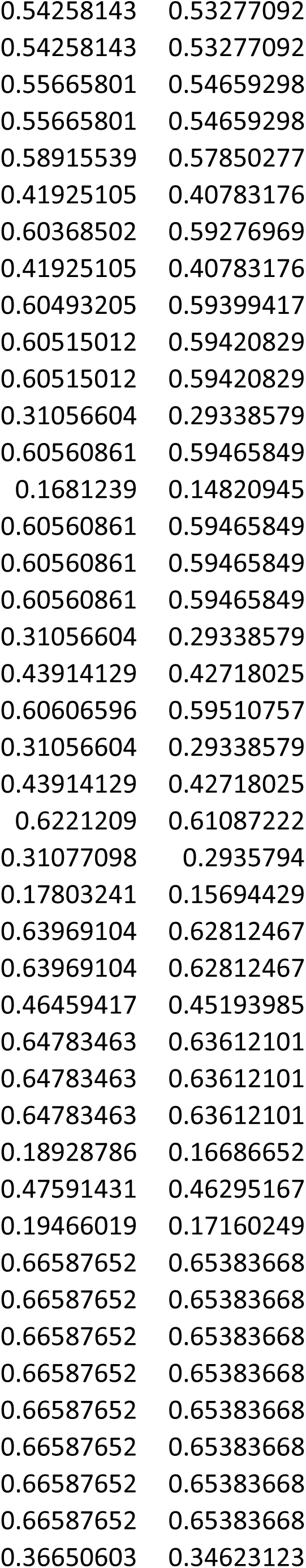
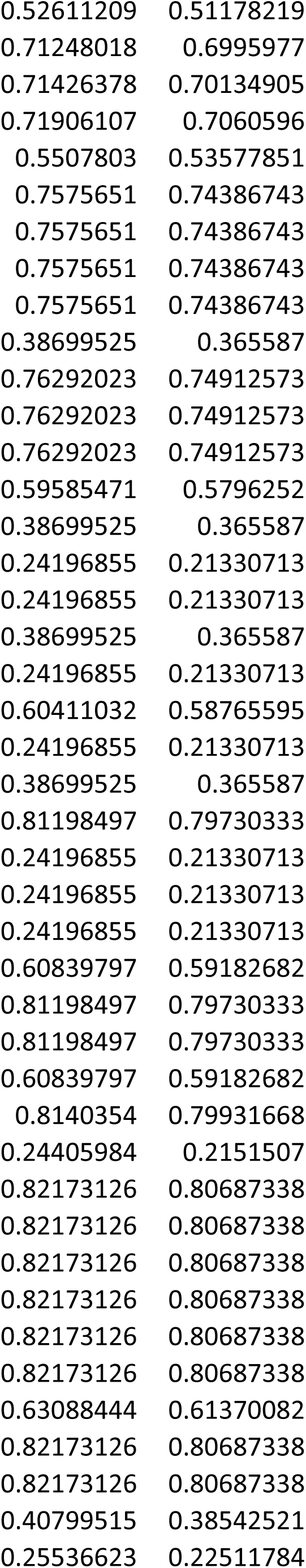
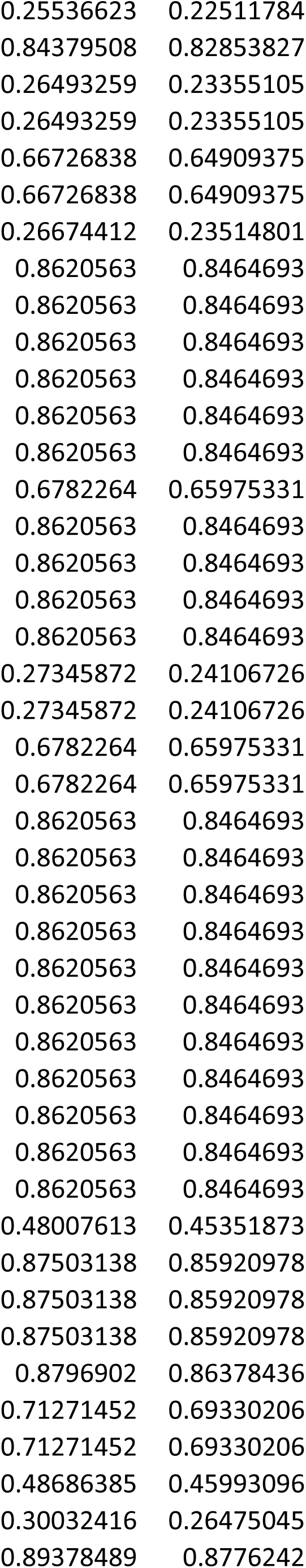
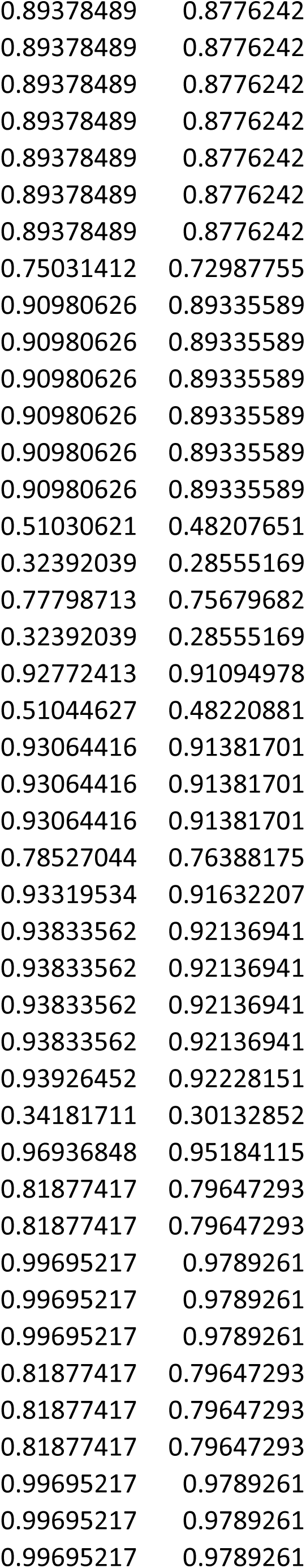
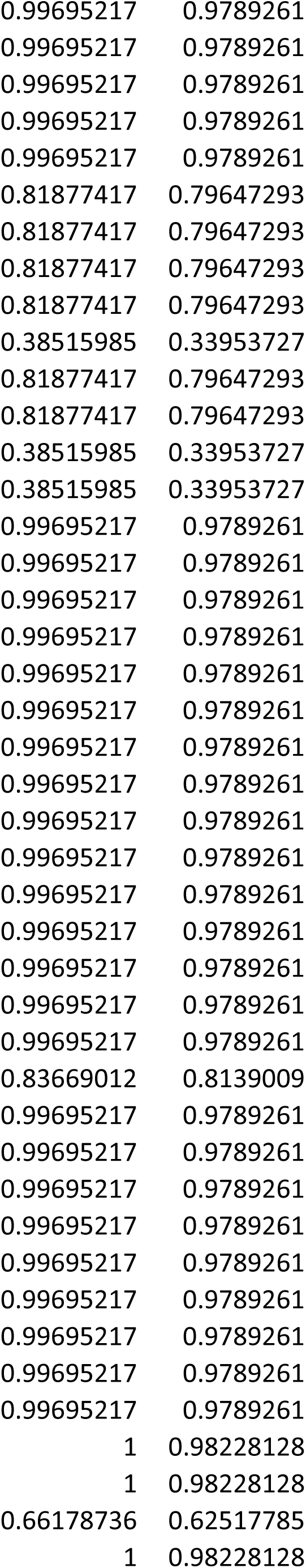
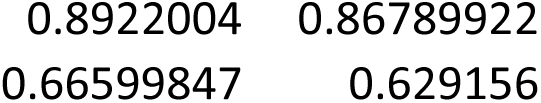

| Category | Term | Count |
| --- | --- | --- |
| GOTERM_CC_DIRECT | GO:0070062~extracellular exosome | 154 |
| GOTERM_CC_DIRECT | GO:0016020~membrane | 286 |
| GOTERM_CC_DIRECT | GO:0005737~cytoplasm | 256 |
| GOTERM_BP_DIRECT | GO:0042552~myelination | 17 |
| GOTERM_CC_DIRECT | GO:0048471~perinuclear region of cytoplasm | 54 |
| GOTERM_CC_DIRECT | GO:0043209~myelin sheath | 26 |
| GOTERM_CC_DIRECT | GO:0005925~focal adhesion | 36 |
| GOTERM_CC_DIRECT | GO:0005856~cytoskeleton | 65 |
| GOTERM_CC_DIRECT | GO:0005913~cell-cell adherens junction | 30 |
| KEGG_PATHWAY | mmu05100:Bacterial invasion of epithelial cells | 15 |
| GOTERM_MF_DIRECT | GO:0003779~actin binding | 30 |
| GOTERM_MF_DIRECT | GO:0098641~cadherin binding involved in cell-cell adh | 26 |
| GOTERM_MF_DIRECT | GO:0051015~actin filament binding | 17 |
| GOTERM_CC_DIRECT | GO:0033270~paranode region of axon | 7 |
| GOTERM_BP_DIRECT | GO:0007155~cell adhesion | 34 |
| KEGG_PATHWAY | mmu04810:Regulation of actin cytoskeleton | 21 |
| GOTERM_CC_DIRECT | GO:0005829~cytosol | 82 |
| GOTERM_MF_DIRECT | GO:0019911~structural constituent of myelin sheath | 5 |
| GOTERM_MF_DIRECT | GO:0005515~protein binding | 158 |
| GOTERM_BP_DIRECT | GO:0098609~cell-cell adhesion | 19 |
| GOTERM_CC_DIRECT | GO:0005783~endoplasmic reticulum | 64 |
| GOTERM_CC_DIRECT | GO:0030054~cell junction | 40 |
| GOTERM_CC_DIRECT | GO:0043220~Schmidt-Lanterman incisure | 6 |
| GOTERM_CC_DIRECT | GO:0030175~filopodium | 11 |
| KEGG_PATHWAY | mmu04360:Axon guidance | 14 |
| KEGG_PATHWAY | mmu04510:Focal adhesion | 18 |
| GOTERM_BP_DIRECT | GO:0006629~lipid metabolic process | 29 |
| GOTERM_BP_DIRECT | GO:0016477~cell migration | 17 |
| KEGG_PATHWAY | mmu04915:Estrogen signaling pathway | 12 |
| GOTERM_MF_DIRECT | GO:0005178~integrin binding | 12 |
| GOTERM_BP_DIRECT | GO:0007399~nervous system development | 25 |
| GOTERM_CC_DIRECT | GO:0030864~cortical actin cytoskeleton | 8 |
| KEGG_PATHWAY | mmu04520:Adherens junction | 10 |
| GOTERM_CC_DIRECT | GO:0030424~axon | 24 |
| GOTERM_CC_DIRECT | GO:0043005~neuron projection | 26 |
| GOTERM_CC_DIRECT | GO:0005794~Golgi apparatus | 54 |
| KEGG_PATHWAY | mmu04015:Rap1 signaling pathway | 17 |
| GOTERM_CC_DIRECT | GO:0015629~actin cytoskeleton | 16 |
| GOTERM_CC_DIRECT | GO:0042470~melanosome | 11 |
| GOTERM_BP_DIRECT | GO:0046627~negative regulation of insulin receptor si | 7 |
| GOTERM_BP_DIRECT | GO:0008202~steroid metabolic process | 10 |
| GOTERM_BP_DIRECT | GO:0030335~positive regulation of cell migration | 16 |
| KEGG_PATHWAY | mmu05205:Proteoglycans in cancer | 16 |
| GOTERM_MF_DIRECT | GO:0008289~lipid binding | 19 |
| GOTERM_BP_DIRECT | GO:0071277~cellular response to calcium ion | 8 |
| GOTERM_BP_DIRECT | GO:0048843~negative regulation of axon extension in | 6 |
| GOTERM_BP_DIRECT | GO:0016126~sterol biosynthetic process | 6 |
| KEGG_PATHWAY | mmu04512:ECM-receptor interaction | 10 |
| GOTERM_BP_DIRECT | GO:0030148~sphingolipid biosynthetic process | 6 |
| GOTERM_CC_DIRECT | GO:0005768~endosome | 29 |
| GOTERM_CC_DIRECT | GO:0005789~endoplasmic reticulum membrane | 35 |
| GOTERM_CC_DIRECT | GO:0009986~cell surface | 32 |
| GOTERM_BP_DIRECT | GO:0007519~skeletal muscle tissue development | 8 |
| KEGG_PATHWAY | mmu04921:Oxytocin signaling pathway | 13 |
| GOTERM_BP_DIRECT | GO:0051726~regulation of cell cycle | 11 |
| GOTERM_CC_DIRECT | GO:0005903~brush border | 9 |
| GOTERM_BP_DIRECT | GO:0042326~negative regulation of phosphorylation | 5 |
| GOTERM_CC_DIRECT | GO:0045202~synapse | 27 |
| GOTERM_CC_DIRECT | GO:0043296~apical junction complex | 5 |
| KEGG_PATHWAY | mmu05200:Pathways in cancer | 23 |
| GOTERM_CC_DIRECT | GO:0030027~lamellipodium | 13 |
| KEGG_PATHWAY | mmu04514:Cell adhesion molecules (CAMs) | 13 |
| GOTERM_CC_DIRECT | GO:0045121~membrane raft | 17 |
| GOTERM_CC_DIRECT | GO:0014069~postsynaptic density | 16 |
| GOTERM_BP_DIRECT | GO:0007264~small GTPase mediated signal transducti | 16 |
| KEGG_PATHWAY | mmu00330:Arginine and proline metabolism | 7 |
| GOTERM_MF_DIRECT | GO:0030215~semaphorin receptor binding | 5 |
| GOTERM_BP_DIRECT | GO:0051017~actin filament bundle assembly | 6 |
| KEGG_PATHWAY | mmu05412:Arrhythmogenic right ventricular cardiomy | 8 |
| GOTERM_BP_DIRECT | GO:0042493~response to drug | 20 |
| KEGG_PATHWAY | mmu04540:Gap junction | 9 |
| GOTERM_CC_DIRECT | GO:0000139~Golgi membrane | 22 |
| GOTERM_CC_DIRECT | GO:0005911~cell-cell junction | 14 |
| GOTERM_BP_DIRECT | GO:0010038~response to metal ion | 4 |
| GOTERM_CC_DIRECT | GO:0030425~dendrite | 25 |
| GOTERM_CC_DIRECT | GO:0045177~apical part of cell | 10 |
| GOTERM_CC_DIRECT | GO:0005764~lysosome | 19 |
| GOTERM_BP_DIRECT | GO:0007229~integrin-mediated signaling pathway | 9 |
| GOTERM_CC_DIRECT | GO:0043218~compact myelin | 3 |
| KEGG_PATHWAY | mmu04725:Cholinergic synapse | 10 |
| KEGG_PATHWAY | mmu04978:Mineral absorption | 6 |
| KEGG_PATHWAY | mmu04010:MAPK signaling pathway | 16 |
| GOTERM_MF_DIRECT | GO:0004860~protein kinase inhibitor activity | 7 |
| GOTERM_BP_DIRECT | GO:0001934~positive regulation of protein phosphory | 13 |
| GOTERM_CC_DIRECT | GO:0005802~trans-Golgi network | 12 |
| GOTERM_BP_DIRECT | GO:0050919~negative chemotaxis | 5 |
| GOTERM_BP_DIRECT | GO:0007160~cell-matrix adhesion | 8 |
| GOTERM_CC_DIRECT | GO:0043025~neuronal cell body | 26 |
| GOTERM_BP_DIRECT | GO:0030036~actin cytoskeleton organization | 11 |
| KEGG_PATHWAY | mmu05132:Salmonella infection | 8 |
| GOTERM_MF_DIRECT | GO:0005096~GTPase activator activity | 15 |
| GOTERM_MF_DIRECT | GO:0045499~chemorepellent activity | 5 |
| GOTERM_CC_DIRECT | GO:0042995~cell projection | 32 |
| GOTERM_CC_DIRECT | GO:0044295~axonal growth cone | 5 |
| GOTERM_BP_DIRECT | GO:0007420~brain development | 14 |
| GOTERM_CC_DIRECT | GO:0033010~paranodal junction | 3 |
| GOTERM_MF_DIRECT | GO:0019904~protein domain specific binding | 17 |
| GOTERM_BP_DIRECT | GO:0043066~negative regulation of apoptotic process | 27 |
| GOTERM_BP_DIRECT | GO:0030308~negative regulation of cell growth | 10 |
| GOTERM_BP_DIRECT | GO:0035556~intracellular signal transduction | 21 |
| GOTERM_BP_DIRECT | GO:0006915~apoptotic process | 27 |
| GOTERM_BP_DIRECT | GO:0043123~positive regulation of I-kappaB kinase/NF | 11 |
| KEGG_PATHWAY | mmu04723:Retrograde endocannabinoid signaling | 9 |
| GOTERM_BP_DIRECT | GO:0007266~Rho protein signal transduction | 6 |
| GOTERM_BP_DIRECT | GO:0051260~protein homooligomerization | 13 |
| GOTERM_CC_DIRECT | GO:0005741~mitochondrial outer membrane | 11 |
| GOTERM_MF_DIRECT | GO:0017049~GTP-Rho binding | 4 |
| KEGG_PATHWAY | mmu04666:Fc gamma R-mediated phagocytosis | 8 |
| KEGG_PATHWAY | mmu05145:Toxoplasmosis | 9 |
| GOTERM_BP_DIRECT | GO:0009636~response to toxic substance | 8 |
| GOTERM_BP_DIRECT | GO:0010942~positive regulation of cell death | 6 |
| GOTERM_BP_DIRECT | GO:0008285~negative regulation of cell proliferation | 20 |
| KEGG_PATHWAY | mmu04720:Long-term potentiation | 7 |
| GOTERM_BP_DIRECT | GO:0035338~long-chain fatty-acyl-CoA biosynthetic pr | 3 |
| GOTERM_BP_DIRECT | GO:0030913~paranodal junction assembly | 3 |
| GOTERM_BP_DIRECT | GO:0001755~neural crest cell migration | 6 |
| GOTERM_BP_DIRECT | GO:0006695~cholesterol biosynthetic process | 5 |
| GOTERM_BP_DIRECT | GO:0008203~cholesterol metabolic process | 8 |
| GOTERM_CC_DIRECT | GO:0005921~gap junction | 5 |
| KEGG_PATHWAY | mmu04912:GnRH signaling pathway | 8 |
| KEGG_PATHWAY | mmu04014:Ras signaling pathway | 14 |
| GOTERM_BP_DIRECT | GO:0051591~response to cAMP | 6 |
| GOTERM_CC_DIRECT | GO:0016459~myosin complex | 6 |
| KEGG_PATHWAY | mmu04530:Tight junction | 8 |
| GOTERM_BP_DIRECT | GO:0071526~semaphorin-plexin signaling pathway | 5 |
| GOTERM_BP_DIRECT | GO:0010976~positive regulation of neuron projection | 10 |
| GOTERM_BP_DIRECT | GO:0016601~Rac protein signal transduction | 4 |
| KEGG_PATHWAY | mmu05218:Melanoma | 7 |
| GOTERM_BP_DIRECT | GO:0019367~fatty acid elongation, saturated fatty acid | 3 |
| GOTERM_BP_DIRECT | GO:0034625~fatty acid elongation, monounsaturated | 3 |
| GOTERM_BP_DIRECT | GO:0034626~fatty acid elongation, polyunsaturated fa | 3 |
| GOTERM_BP_DIRECT | GO:0031175~neuron projection development | 10 |
| GOTERM_MF_DIRECT | GO:0102336~3-oxo-arachidoyl-CoA synthase activity | 3 |
| GOTERM_MF_DIRECT | GO:0102337~3-oxo-cerotoyl-CoA synthase activity | 3 |
| GOTERM_MF_DIRECT | GO:0102338~3-oxo-lignoceronyl-CoA synthase activity | 3 |
| GOTERM_MF_DIRECT | GO:0009922~fatty acid elongase activity | 3 |
| GOTERM_BP_DIRECT | GO:0016192~vesicle-mediated transport | 13 |
| KEGG_PATHWAY | mmu04724:Glutamatergic synapse | 9 |
| GOTERM_CC_DIRECT | GO:0005604~basement membrane | 8 |
| GOTERM_CC_DIRECT | GO:0005938~cell cortex | 10 |
| GOTERM_CC_DIRECT | GO:0005874~microtubule | 17 |
| GOTERM_BP_DIRECT | GO:0022409~positive regulation of cell-cell adhesion | 4 |
| GOTERM_CC_DIRECT | GO:0005902~microvillus | 7 |
| GOTERM_MF_DIRECT | GO:0017124~SH3 domain binding | 9 |
| KEGG_PATHWAY | mmu05133:Pertussis | 7 |
| KEGG_PATHWAY | mmu04670:Leukocyte transendothelial migration | 9 |
| GOTERM_BP_DIRECT | GO:0032287~peripheral nervous system myelin mainte | 3 |
| GOTERM_BP_DIRECT | GO:0071679~commissural neuron axon guidance | 3 |
| GOTERM_BP_DIRECT | GO:0051412~response to corticosterone | 4 |
| GOTERM_CC_DIRECT | GO:0043197~dendritic spine | 10 |
| KEGG_PATHWAY | mmu04713:Circadian entrainment | 8 |
| GOTERM_CC_DIRECT | GO:0043204~perikaryon | 10 |
| KEGG_PATHWAY | mmu04916:Melanogenesis | 8 |
| GOTERM_BP_DIRECT | GO:0031668~cellular response to extracellular stimulu | 4 |
| GOTERM_CC_DIRECT | GO:0035748~myelin sheath abaxonal region | 3 |
| GOTERM_CC_DIRECT | GO:0030056~hemidesmosome | 3 |
| KEGG_PATHWAY | mmu04071:Sphingolipid signaling pathway | 9 |
| KEGG_PATHWAY | mmu05231:Choline metabolism in cancer | 8 |
| GOTERM_BP_DIRECT | GO:0071456~cellular response to hypoxia | 8 |
| GOTERM_BP_DIRECT | GO:0051898~negative regulation of protein kinase B si | 5 |
| GOTERM_MF_DIRECT | GO:0042803~protein homodimerization activity | 33 |
| GOTERM_BP_DIRECT | GO:0042759~long-chain fatty acid biosynthetic proces | 3 |
| GOTERM_BP_DIRECT | GO:0030042~actin filament depolymerization | 3 |
| GOTERM_MF_DIRECT | GO:0047372~acylglycerol lipase activity | 3 |
| GOTERM_BP_DIRECT | GO:0007010~cytoskeleton organization | 8 |
| GOTERM_CC_DIRECT | GO:0030496~midbody | 9 |
| KEGG_PATHWAY | mmu04960:Aldosterone-regulated sodium reabsorptic | 5 |
| GOTERM_CC_DIRECT | GO:0005765~lysosomal membrane | 13 |
| GOTERM_CC_DIRECT | GO:0005770~late endosome | 9 |
| GOTERM_CC_DIRECT | GO:0005923~bicellular tight junction | 9 |
| GOTERM_CC_DIRECT | GO:0031674~I band | 4 |
| KEGG_PATHWAY | mmu04270:Vascular smooth muscle contraction | 9 |
| GOTERM_MF_DIRECT | GO:0048365~Rac GTPase binding | 5 |
| GOTERM_BP_DIRECT | GO:0001822~kidney development | 9 |
| GOTERM_CC_DIRECT | GO:0005886~plasma membrane | 152 |
| GOTERM_BP_DIRECT | GO:0010811~positive regulation of cell-substrate adhe | 5 |
| KEGG_PATHWAY | mmu04730:Long-term depression | 6 |
| GOTERM_MF_DIRECT | GO:0000982~transcription factor activity, RNA polyme | 4 |
| GOTERM_CC_DIRECT | GO:0044754~autolysosome | 3 |
| KEGG_PATHWAY | mmu05414:Dilated cardiomyopathy | 7 |
| GOTERM_CC_DIRECT | GO:0008305~integrin complex | 4 |
| GOTERM_BP_DIRECT | GO:0008542~visual learning | 6 |
| GOTERM_BP_DIRECT | GO:0071404~cellular response to low-density lipoprot | 3 |
| KEGG_PATHWAY | mmu04611:Platelet activation | 9 |
| GOTERM_BP_DIRECT | GO:0045860~positive regulation of protein kinase acti | 6 |
| GOTERM_CC_DIRECT | GO:0031902~late endosome membrane | 7 |
| KEGG_PATHWAY | mmu04144:Endocytosis | 14 |
| GOTERM_BP_DIRECT | GO:0001525~angiogenesis | 13 |
| GOTERM_BP_DIRECT | GO:0043524~negative regulation of neuron apoptotic | 10 |
| GOTERM_BP_DIRECT | GO:0010033~response to organic substance | 6 |
| KEGG_PATHWAY | mmu05210:Colorectal cancer | 6 |
| KEGG_PATHWAY | mmu05140:Leishmaniasis | 6 |
| GOTERM_CC_DIRECT | GO:0031982~vesicle | 10 |
| GOTERM_BP_DIRECT | GO:0051781~positive regulation of cell division | 5 |
| GOTERM_CC_DIRECT | GO:0001725~stress fiber | 6 |
| GOTERM_MF_DIRECT | GO:0019901~protein kinase binding | 20 |
| GOTERM_BP_DIRECT | GO:0016242~negative regulation of macroautophagy | 3 |
| KEGG_PATHWAY | mmu04012:ErbB signaling pathway | 7 |
| KEGG_PATHWAY | mmu04727:GABAergic synapse | 7 |
| GOTERM_MF_DIRECT | GO:0008270~zinc ion binding | 41 |
| GOTERM_CC_DIRECT | GO:0031234~extrinsic component of cytoplasmic side | 6 |
| GOTERM_BP_DIRECT | GO:0031032~actomyosin structure organization | 4 |
| GOTERM_BP_DIRECT | GO:2001243~negative regulation of intrinsic apoptotic | 4 |
| KEGG_PATHWAY | mmu04151:PI3K-Akt signaling pathway | 17 |
| GOTERM_BP_DIRECT | GO:0043525~positive regulation of neuron apoptotic p | 6 |
| GOTERM_BP_DIRECT | GO:0019538~protein metabolic process | 3 |
| GOTERM_BP_DIRECT | GO:0042761~very long-chain fatty acid biosynthetic pr | 3 |
| GOTERM_BP_DIRECT | GO:0008210~estrogen metabolic process | 3 |
| GOTERM_MF_DIRECT | GO:0044548~S100 protein binding | 3 |
| GOTERM_MF_DIRECT | GO:0017166~vinculin binding | 3 |
| GOTERM_MF_DIRECT | GO:0004861~cyclin-dependent protein serine/threonin | 3 |
| GOTERM_MF_DIRECT | GO:0032403~protein complex binding | 17 |
| GOTERM_CC_DIRECT | GO:0071944~cell periphery | 6 |
| GOTERM_MF_DIRECT | GO:0019900~kinase binding | 7 |
| GOTERM_CC_DIRECT | GO:0001726~ruffle | 7 |
| GOTERM_BP_DIRECT | GO:2000378~negative regulation of reactive oxygen sp | 4 |
| GOTERM_BP_DIRECT | GO:0046686~response to cadmium ion | 4 |
| GOTERM_BP_DIRECT | GO:0060291~long-term synaptic potentiation | 5 |
| KEGG_PATHWAY | mmu05216:Thyroid cancer | 4 |
| GOTERM_CC_DIRECT | GO:0010008~endosome membrane | 8 |
| GOTERM_BP_DIRECT | GO:1900181~negative regulation of protein localizatio | 3 |
| GOTERM_CC_DIRECT | GO:0005578~proteinaceous extracellular matrix | 15 |
| GOTERM_MF_DIRECT | GO:0008432~JUN kinase binding | 3 |
| GOTERM_MF_DIRECT | GO:0030898~actin-dependent ATPase activity | 3 |
| KEGG_PATHWAY | mmu04062:Chemokine signaling pathway | 11 |
| GOTERM_BP_DIRECT | GO:0007611~learning or memory | 5 |
| KEGG_PATHWAY | mmu04971:Gastric acid secretion | 6 |
| GOTERM_BP_DIRECT | GO:0006633~fatty acid biosynthetic process | 6 |
| GOTERM_MF_DIRECT | GO:0008017~microtubule binding | 11 |
| KEGG_PATHWAY | mmu05161:Hepatitis B | 9 |
| GOTERM_BP_DIRECT | GO:0007411~axon guidance | 9 |
| GOTERM_CC_DIRECT | GO:0071682~endocytic vesicle lumen | 2 |
| GOTERM_CC_DIRECT | GO:0097038~perinuclear endoplasmic reticulum | 3 |
| GOTERM_CC_DIRECT | GO:0031143~pseudopodium | 3 |
| GOTERM_CC_DIRECT | GO:0043231~intracellular membrane-bounded organe | 29 |
| GOTERM_MF_DIRECT | GO:0001540~beta-amyloid binding | 4 |
| GOTERM_BP_DIRECT | GO:0006682~galactosylceramide biosynthetic process | 2 |
| GOTERM_BP_DIRECT | GO:0006601~creatine biosynthetic process | 2 |
| GOTERM_BP_DIRECT | GO:1902746~regulation of lens fiber cell differentiatio | 2 |
| GOTERM_MF_DIRECT | GO:0008396~oxysterol 7-alpha-hydroxylase activity | 2 |
| GOTERM_BP_DIRECT | GO:0071850~mitotic cell cycle arrest | 3 |
| GOTERM_BP_DIRECT | GO:0070498~interleukin-1-mediated signaling pathwa | 3 |
| GOTERM_BP_DIRECT | GO:0007179~transforming growth factor beta recepto | 6 |
| KEGG_PATHWAY | mmu05213:Endometrial cancer | 5 |
| GOTERM_MF_DIRECT | GO:0008201~heparin binding | 9 |
| GOTERM_BP_DIRECT | GO:0090090~negative regulation of canonical Wnt sigr | 7 |
| GOTERM_CC_DIRECT | GO:0032587~ruffle membrane | 6 |
| GOTERM_BP_DIRECT | GO:0031643~positive regulation of myelination | 3 |
| GOTERM_BP_DIRECT | GO:0033627~cell adhesion mediated by integrin | 3 |
| GOTERM_BP_DIRECT | GO:0006636~unsaturated fatty acid biosynthetic proce | 3 |
| GOTERM_BP_DIRECT | GO:0043149~stress fiber assembly | 3 |
| GOTERM_CC_DIRECT | GO:0005635~nuclear envelope | 9 |
| GOTERM_MF_DIRECT | GO:0016757~transferase activity, transferring glycosyl | 11 |
| GOTERM_BP_DIRECT | GO:0007616~long-term memory | 4 |
| GOTERM_CC_DIRECT | GO:0030426~growth cone | 9 |
| GOTERM_BP_DIRECT | GO:0043410~positive regulation of MAPK cascade | 7 |
| GOTERM_CC_DIRECT | GO:0033268~node of Ranvier | 3 |
| GOTERM_MF_DIRECT | GO:0003774~motor activity | 6 |
| GOTERM_BP_DIRECT | GO:0031103~axon regeneration | 3 |
| GOTERM_BP_DIRECT | GO:0022011~myelination in peripheral nervous system | 3 |
| GOTERM_BP_DIRECT | GO:0009617~response to bacterium | 4 |
| GOTERM_MF_DIRECT | GO:0019899~enzyme binding | 17 |
| KEGG_PATHWAY | mmu05416:Viral myocarditis | 6 |
| KEGG_PATHWAY | mmu05410:Hypertrophic cardiomyopathy (HCM) | 6 |
| GOTERM_BP_DIRECT | GO:0009408~response to heat | 5 |
| GOTERM_BP_DIRECT | GO:0007050~cell cycle arrest | 6 |
| GOTERM_MF_DIRECT | GO:0030544~Hsp70 protein binding | 4 |
| GOTERM_BP_DIRECT | GO:0018105~peptidyl-serine phosphorylation | 8 |
| GOTERM_BP_DIRECT | GO:0006665~sphingolipid metabolic process | 4 |
| GOTERM_CC_DIRECT | GO:0044291~cell-cell contact zone | 3 |
| GOTERM_MF_DIRECT | GO:0017048~Rho GTPase binding | 4 |
| KEGG_PATHWAY | mmu04726:Serotonergic synapse | 8 |
| GOTERM_BP_DIRECT | GO:0007015~actin filament organization | 6 |
| GOTERM_BP_DIRECT | GO:0030048~actin filament-based movement | 3 |
| GOTERM_BP_DIRECT | GO:0048193~Golgi vesicle transport | 3 |
| GOTERM_BP_DIRECT | GO:0042692~muscle cell differentiation | 3 |
| GOTERM_BP_DIRECT | GO:0008654~phospholipid biosynthetic process | 5 |
| KEGG_PATHWAY | mmu01130:Biosynthesis of antibiotics | 11 |
| GOTERM_CC_DIRECT | GO:0019898~extrinsic component of membrane | 6 |
| GOTERM_BP_DIRECT | GO:0043065~positive regulation of apoptotic process | 15 |
| GOTERM_BP_DIRECT | GO:0019065~receptor-mediated endocytosis of virus k | 2 |
| GOTERM_BP_DIRECT | GO:2000544~regulation of endothelial cell chemotaxis | 2 |
| GOTERM_BP_DIRECT | GO:0061084~negative regulation of protein refolding | 2 |
| GOTERM_BP_DIRECT | GO:0032290~peripheral nervous system myelin forma | 2 |
| GOTERM_BP_DIRECT | GO:0006598~polyamine catabolic process | 2 |
| GOTERM_MF_DIRECT | GO:0031871~proteinase activated receptor binding | 2 |
| GOTERM_MF_DIRECT | GO:0035373~chondroitin sulfate proteoglycan binding | 2 |
| GOTERM_CC_DIRECT | GO:0005739~mitochondrion | 57 |
| GOTERM_CC_DIRECT | GO:0043679~axon terminus | 6 |
| GOTERM_MF_DIRECT | GO:0019003~GDP binding | 5 |
| GOTERM_BP_DIRECT | GO:0021762~substantia nigra development | 4 |
| GOTERM_BP_DIRECT | GO:0007605~sensory perception of sound | 8 |
| GOTERM_BP_DIRECT | GO:0006882~cellular zinc ion homeostasis | 3 |
| GOTERM_MF_DIRECT | GO:0051393~alpha-actinin binding | 3 |
| KEGG_PATHWAY | mmu04022:cGMP-PKG signaling pathway | 9 |
| KEGG_PATHWAY | mmu05166:HTLV-I infection | 13 |
| GOTERM_BP_DIRECT | GO:0008360~regulation of cell shape | 8 |
| GOTERM_MF_DIRECT | GO:0030674~protein binding, bridging | 5 |
| GOTERM_BP_DIRECT | GO:0006694~steroid biosynthetic process | 5 |
| GOTERM_BP_DIRECT | GO:0007154~cell communication | 4 |
| GOTERM_CC_DIRECT | GO:0016234~inclusion body | 3 |
| GOTERM_BP_DIRECT | GO:0009612~response to mechanical stimulus | 5 |
| GOTERM_BP_DIRECT | GO:0034599~cellular response to oxidative stress | 5 |
| GOTERM_BP_DIRECT | GO:0031116~positive regulation of microtubule polym | 3 |
| GOTERM_BP_DIRECT | GO:0043409~negative regulation of MAPK cascade | 3 |
| GOTERM_BP_DIRECT | GO:0031589~cell-substrate adhesion | 3 |
| GOTERM_MF_DIRECT | GO:0031434~mitogen-activated protein kinase kinase | 3 |
| KEGG_PATHWAY | mmu00100:Steroid biosynthesis | 3 |
| GOTERM_CC_DIRECT | GO:0016323~basolateral plasma membrane | 10 |

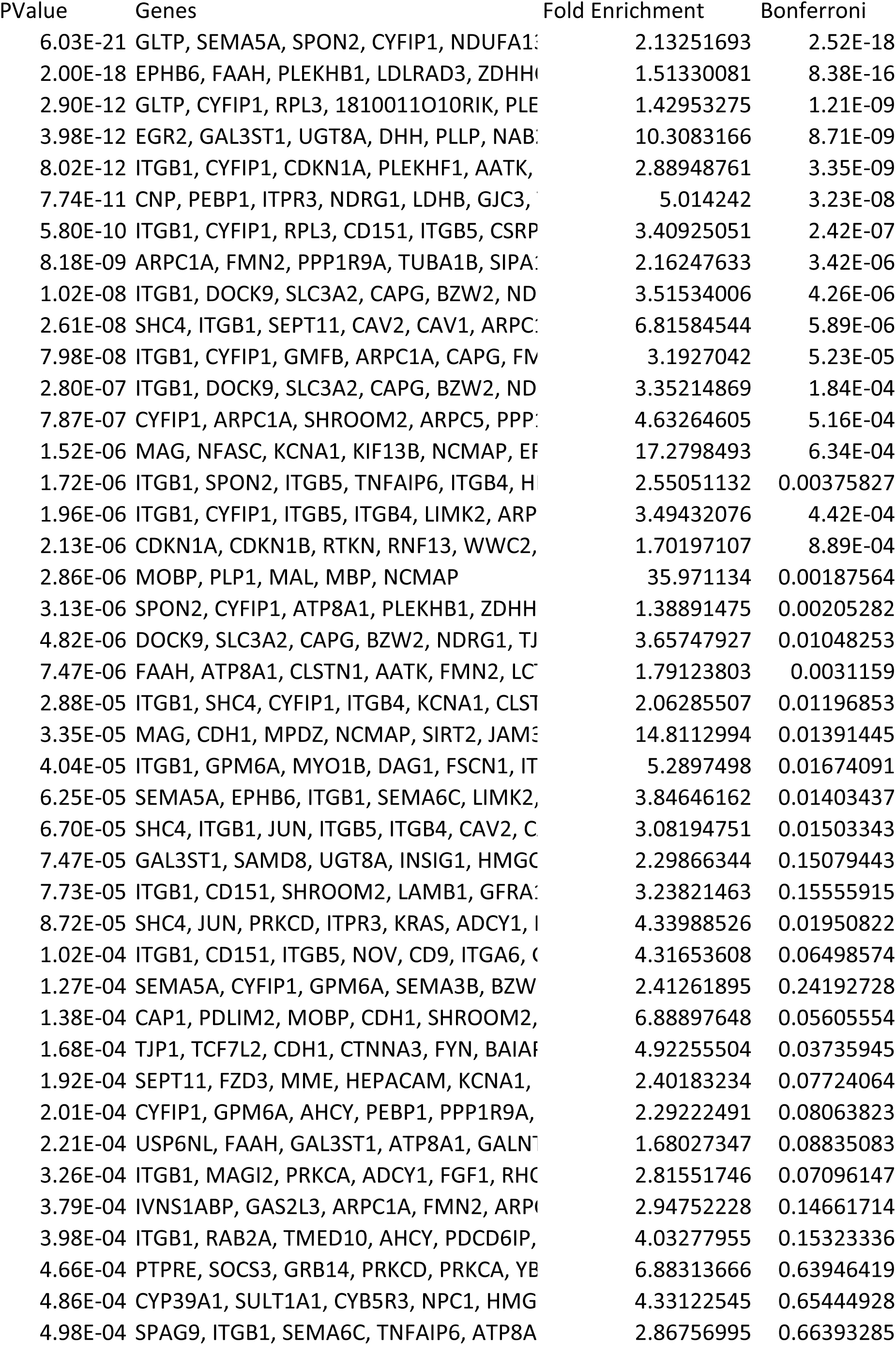
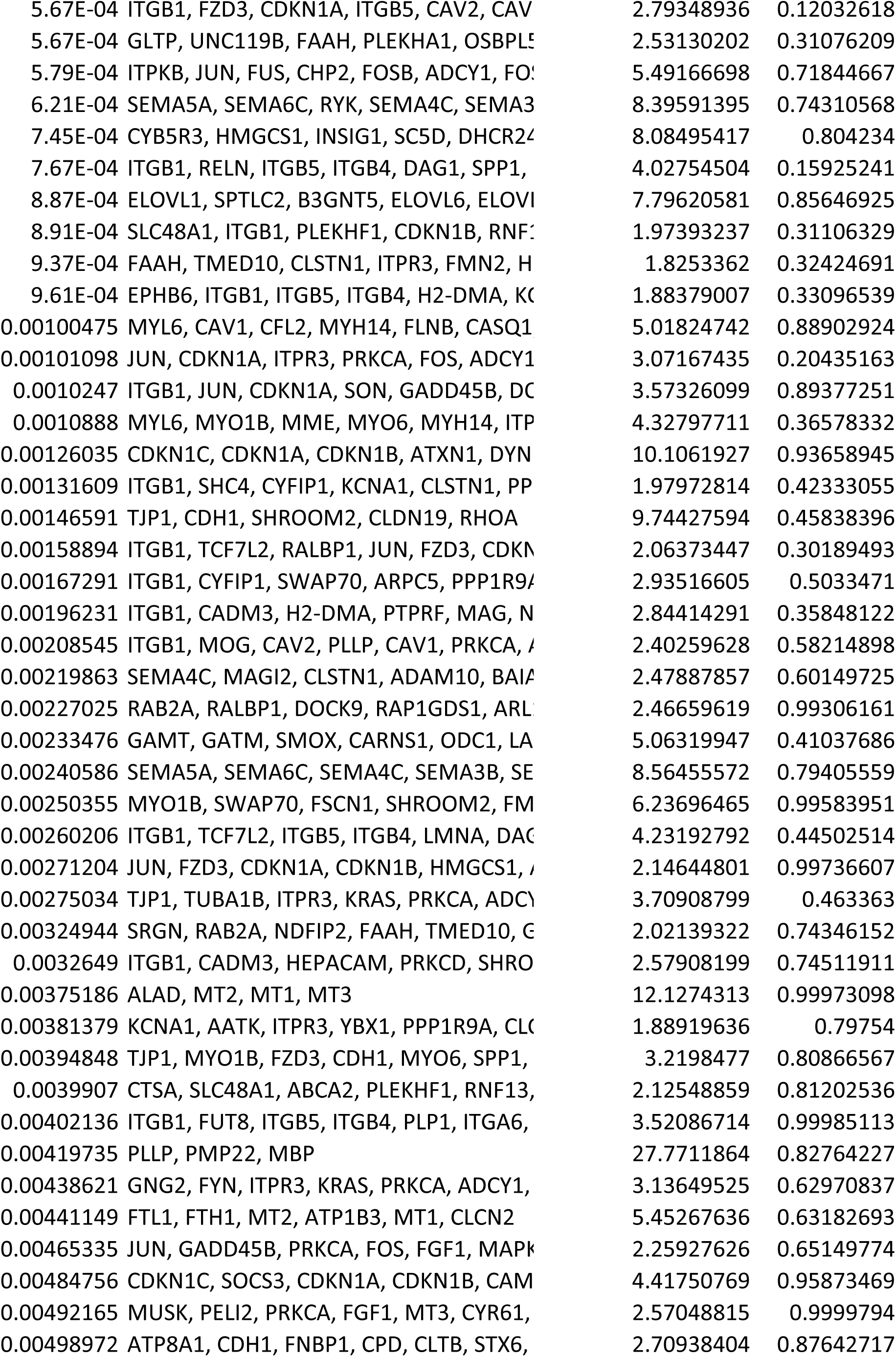
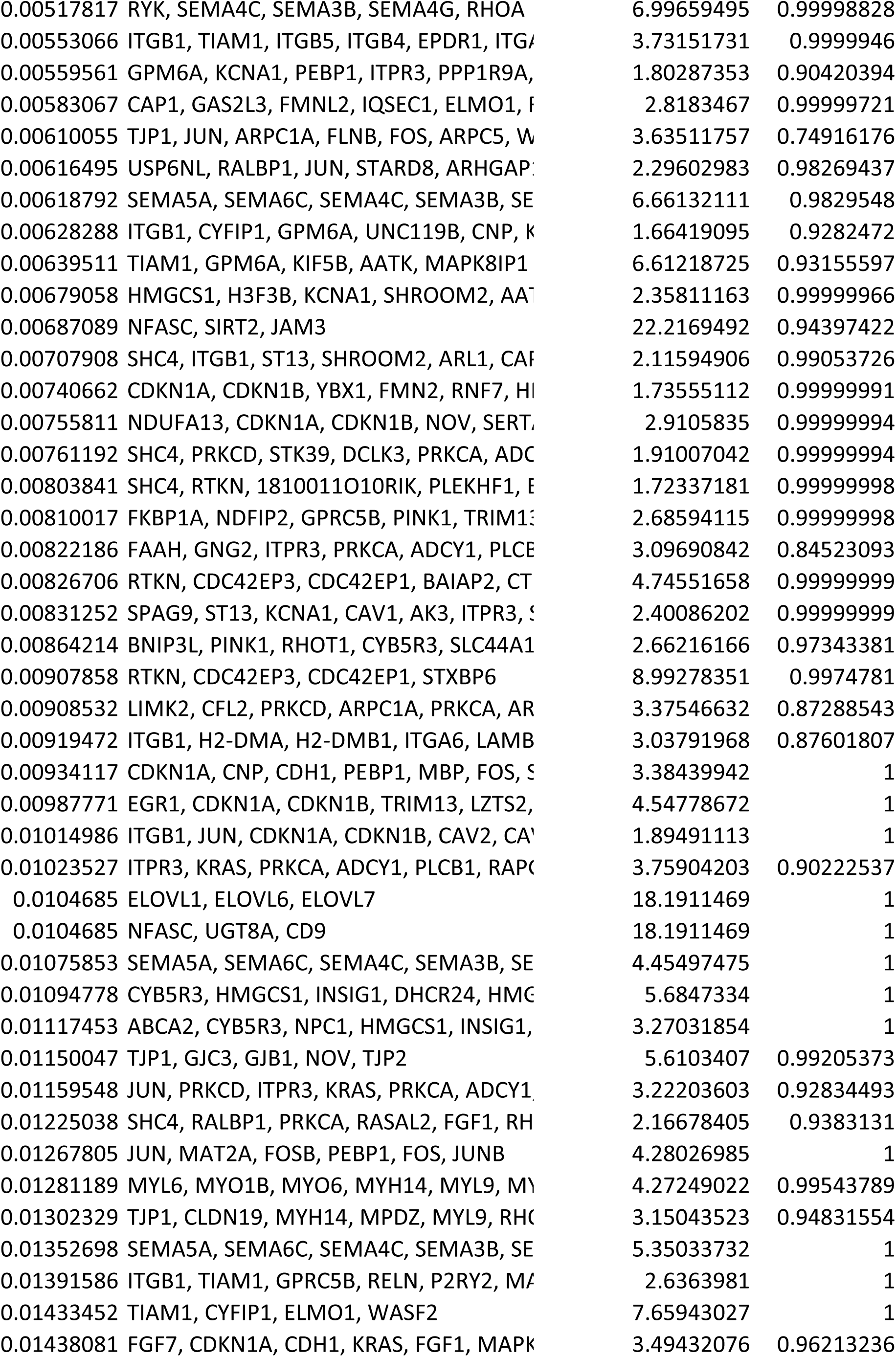
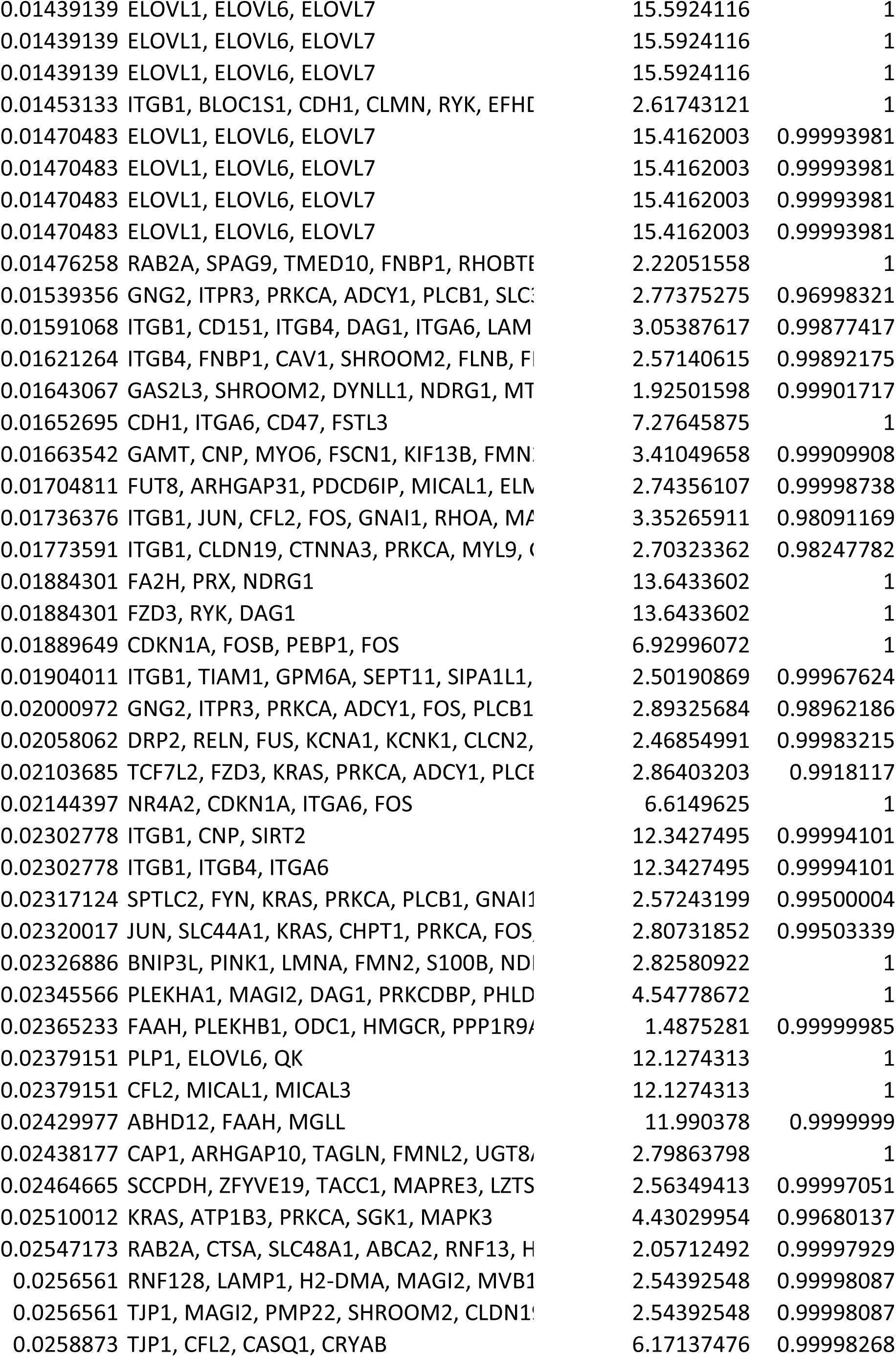
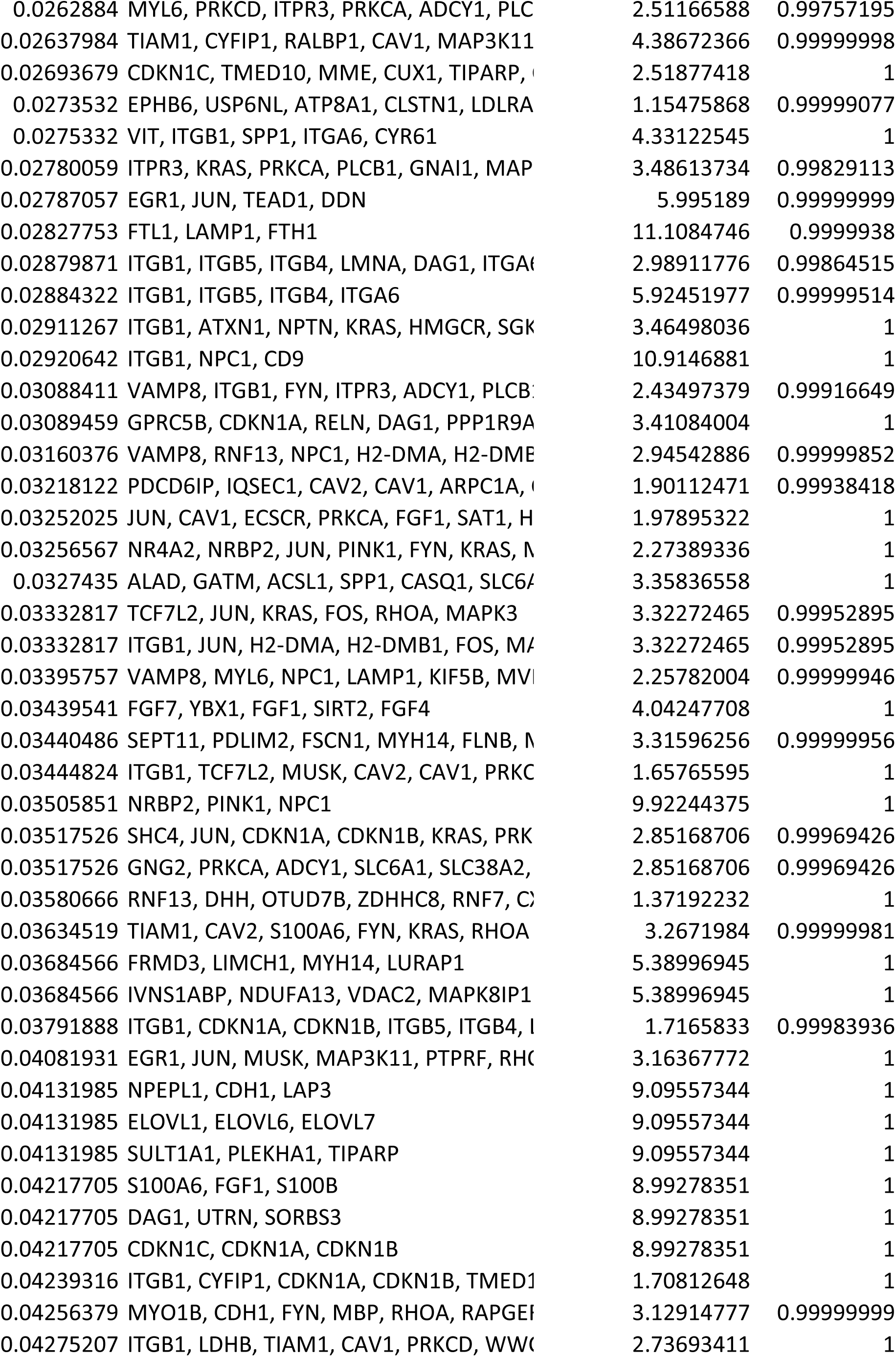
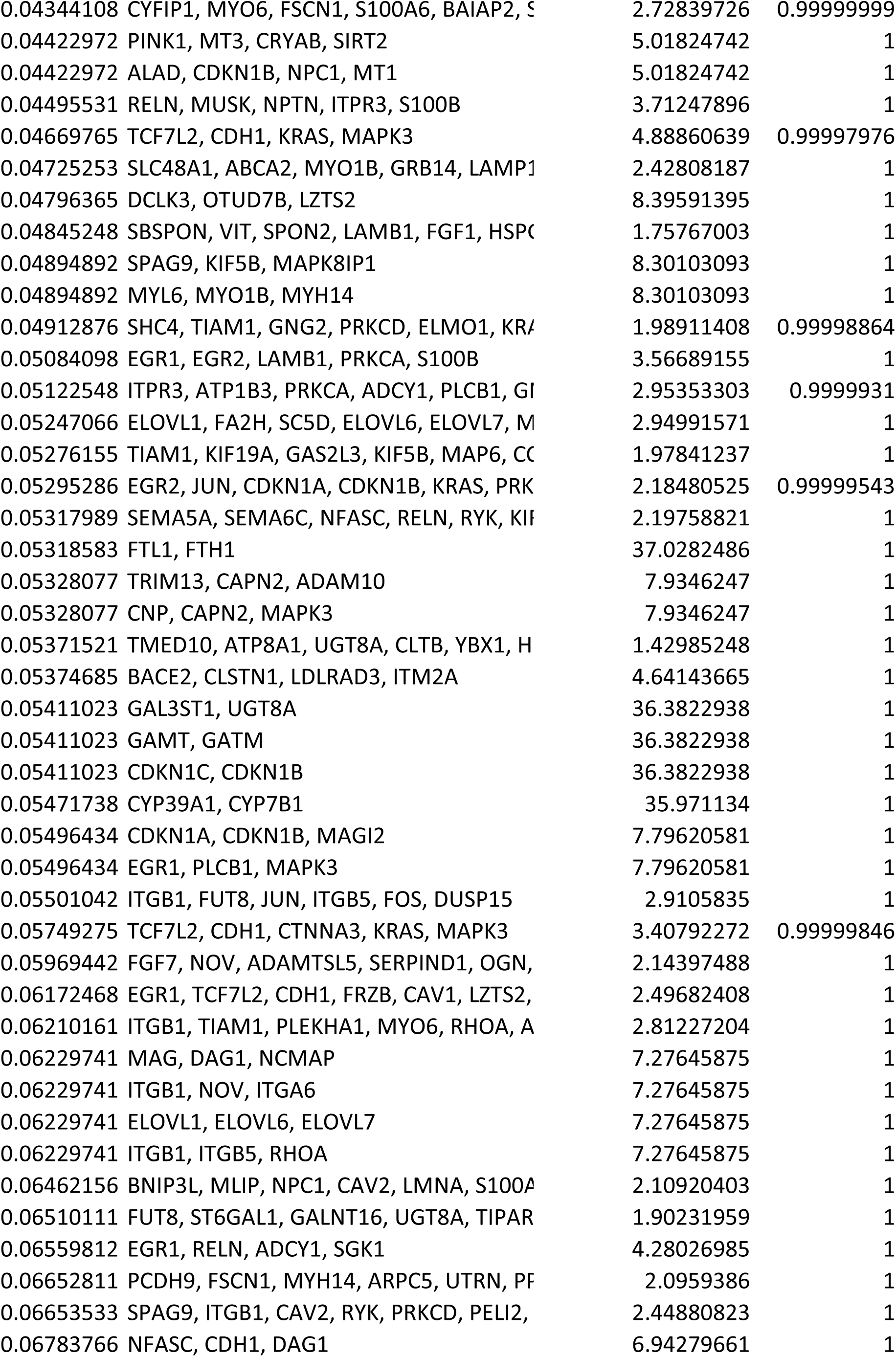
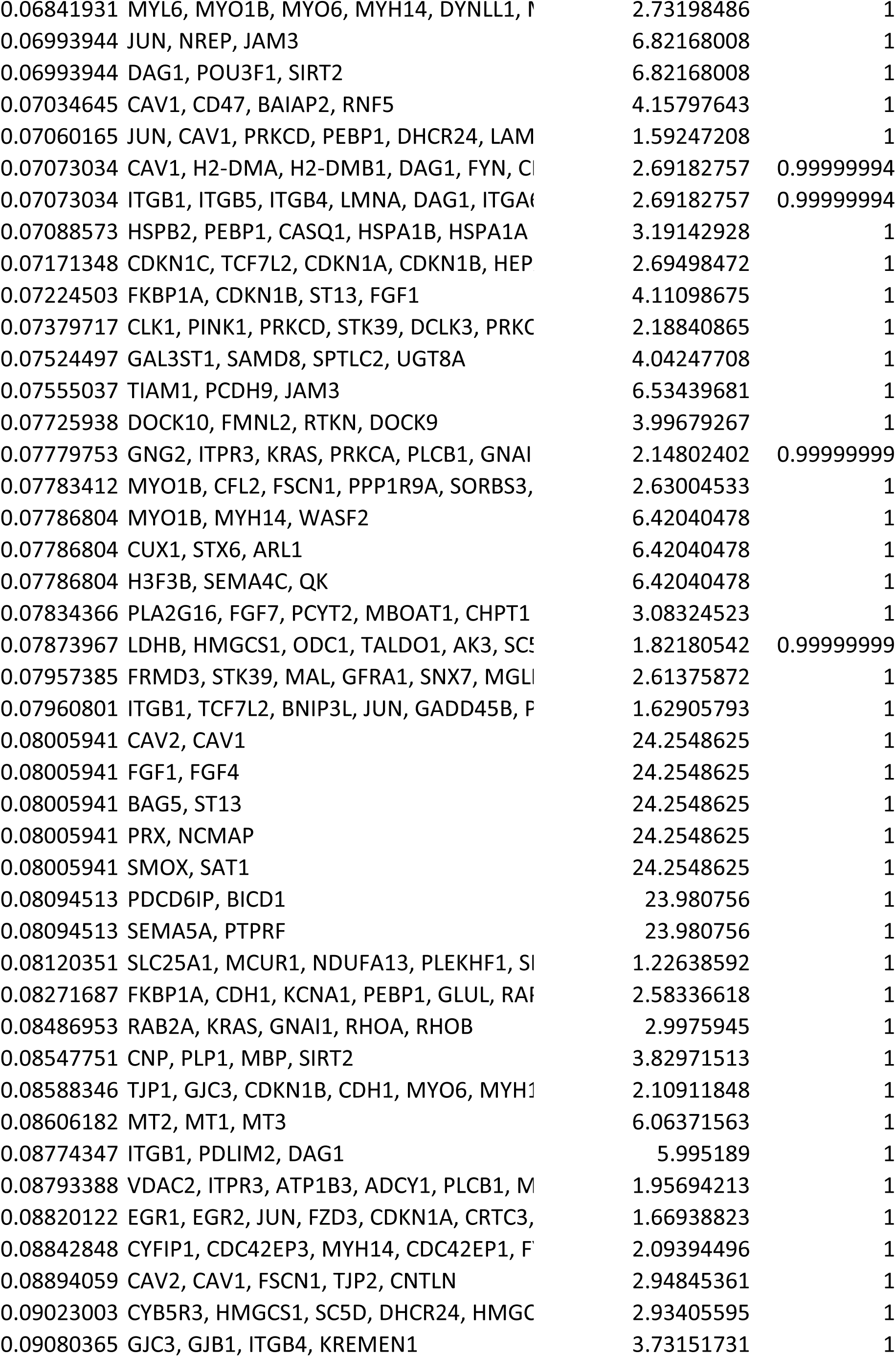
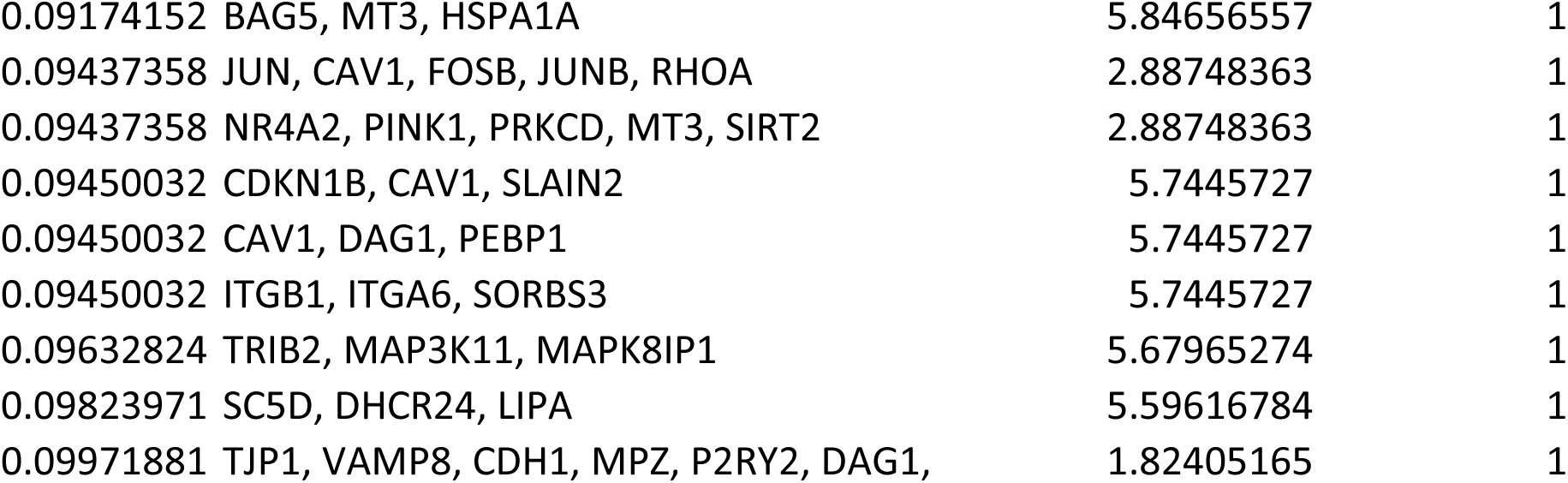

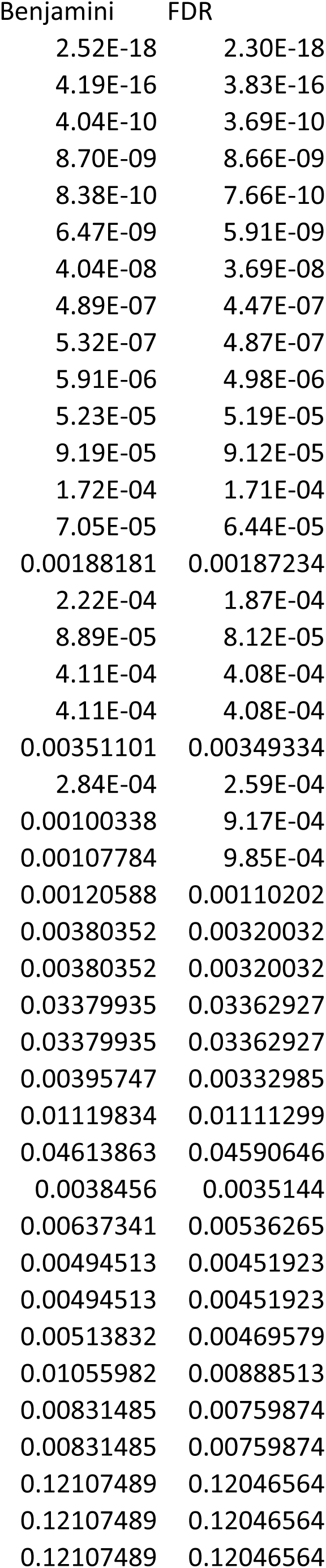
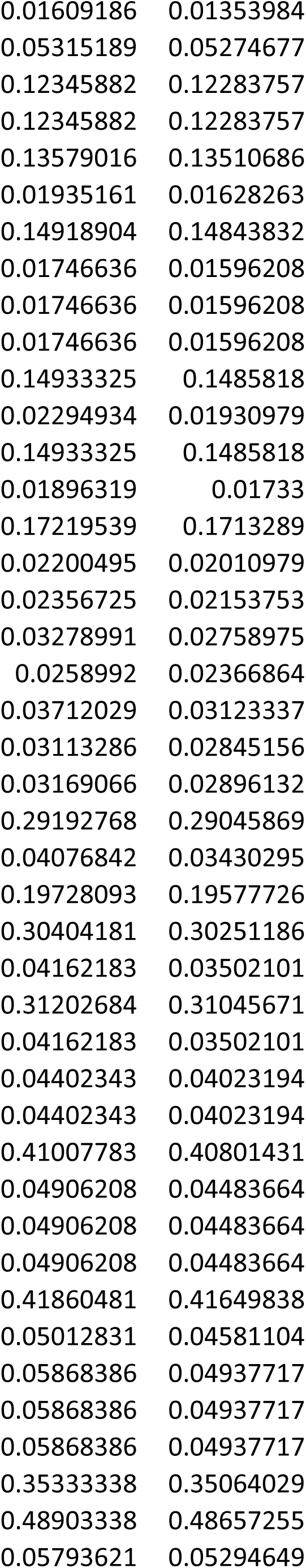
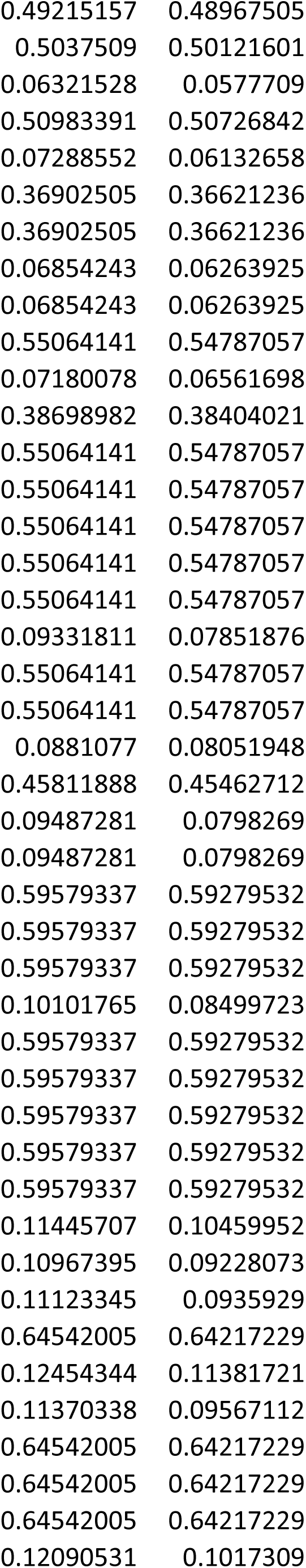
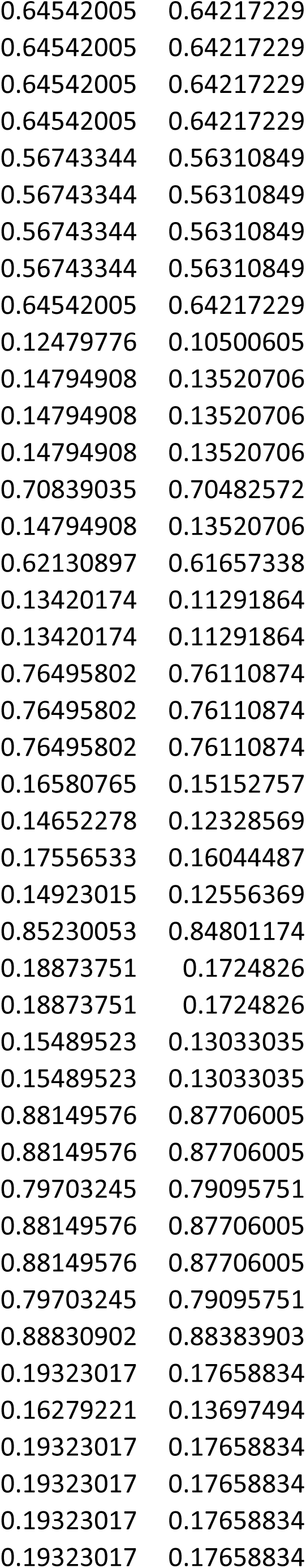
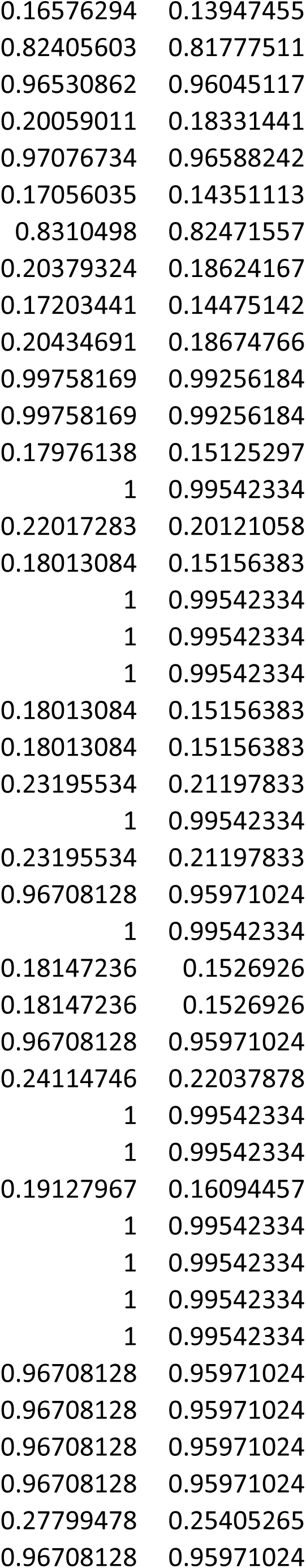
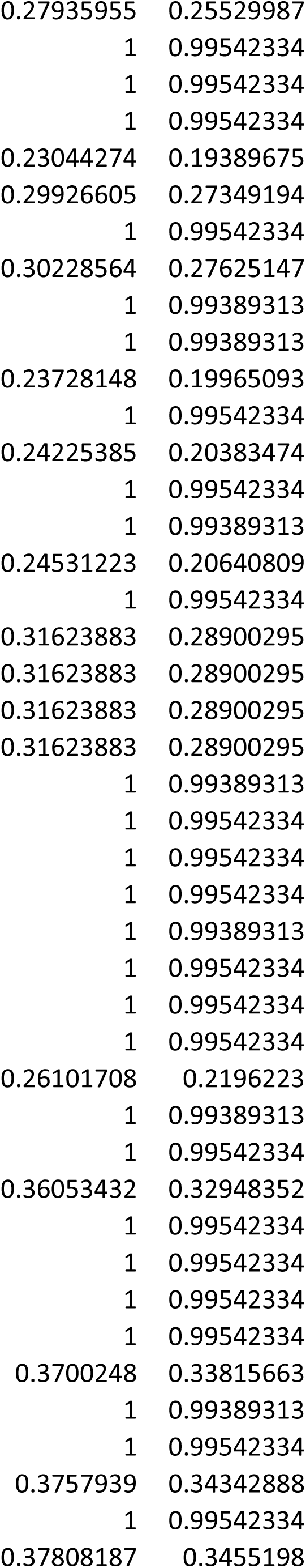
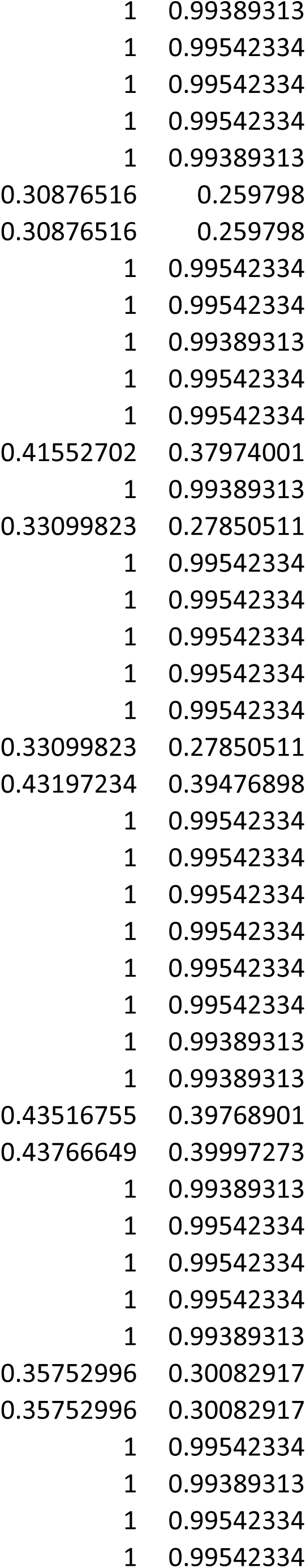
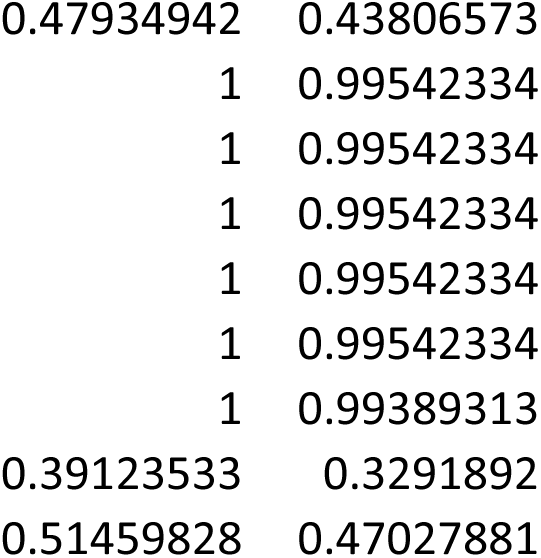

